# Resolving sub-clonal heterogeneity within cell-line growths by single cell sequencing genomic DNA

**DOI:** 10.1101/757211

**Authors:** Enrique I. Velazquez-Villarreal, Shamoni Maheshwari, Jon Sorenson, Ian T. Fiddes, Vijay Kumar, Yifeng Yin, Michelle Webb, Claudia Catalanotti, Mira Grigorova, Paul A. Edwards, John D. Carpten, David W. Craig

## Abstract

We performed shallow single-cell sequencing of genomic DNA across 1,475 cells from a well-studied cell-line, COLO829, to resolve overall tumor complexity and clonality. This melanoma tumor-line has been previously characterized by multiple technologies and provides a benchmark for evaluating somatic alterations, though has exhibited conflicting and indeterminate copy number states. We identified at least four major sub-clones by discriminant analysis of principal components (DAPC) of single cell copy number data. Break-point and loss of heterozygosity (LOH) analysis of aggregated data from sub-clones revealed a complex rearrangement of chromosomes 1, 10 and 18 that was maintained in all but two sub-clones. Likewise, two of the sub-clones were distinguished by loss of 1 copy of chromosome 8. Re-analysis of previous spectral karyotyping data and bulk sequencing data recapitulated these sub-clone hallmark features and explains why the original bulk sequencing experiments generated conflicting copy number results. Overall, our results demonstrate how shallow copy number profiling together with clustering analysis of single cell sequencing can uncover significant hidden insights even in well studied cell-lines.

## INTRODUCTION

Gaining a single cell view of tumor heterogeneity is crucial for improving our understanding of tumor evolution and enabling future advances in cancer research. The standard paradigm is bulk sequencing of genomic DNA derived from millions of heterogeneous cells. In bulk sequencing, the ability to resolve sub-clonality is largely lost except for indirect inference, resulting in an ensemble view dominated by the majority clone^1,2^. While bulk sequencing has provided major insights into tumor biology, low-resolution single cell methods, such as spectral karyotyping, show that dissecting events at single cell resolution is critical to accurately describe the genome of heterogeneous population of cells that underlie sub-clonal complexity and tumor evolution. Here, we used droplet-based shallow genome sequencing of 1,475 single cells from a well characterized melanoma cell line to reveal that it carried at least four different populations with distinct genome-wide copy number profiles.

The melanoma COLO829 and germline COLO829-BL tumor/normal pair have been extensively analyzed using multiple methods and technologies ^1–4^. The first exhaustive sequencing of this tumor/normal pair by Pleasance *et al* described several hallmark events including a homozygous 12kb deletion in PTEN, BRAF 600V/E, and a CDK2NA 2bp deletion. Previous studies using bulk sequencing in the tumor-line COLO829 have focused largely on developing tools, catalogues and standardizations to improve copy number estimation and cancer characterization^2^. While a few of the studies found cell line complexity inconsistent with the assumption of clonality and suggestive of multiple sub-clones, in general, most analyses presumed COLO829 to be a single clone. Of papers looking at copy number, Craig *et al* observed differences among growths in chromosome 1p, and Gusnanto *et al* found evidence for a mixture of clones, but were unable to resolve the individual components using bulk data and methods. Much of the work on this tumor-line highlighted major CNV hallmark events, as well as a series of inconsistent findings that point towards bulk sequencing methods being lossy and unable to resolve the complexity of COLO829^4^.

Beyond the difficulty of resolving clonal mixtures, an additional challenge of bulk sequencing even in the context of a paired normal is that without single-cell resolution there are limited informatic options to resolve relative differences in read-depth to integer copy number states. At some point, most algorithms require an assumption, such as a diploid region or tumor purity, to effectively normalize against and the success in doing so can be dependent on the accuracy of that particular assumption. Likewise, even with a uniform set of algorithms applied on the same cell line variable results were observed across institutions, suggesting that there may be differences among growths with some sub-populations of cells^4^. In this paper we performed shallow single-cell sequencing of genomic DNA across 1,475 cells from the same cell-line, COLO829, and show that it is in fact a complex mixture and identify key structural variants that contribute to its sub-clonal evolution.

## RESULTS

### Copy number profiling at single cell resolution

We sequenced 1,475 cells with 3.044 billion 2×100 paired-end reads, and conducted barcode-aware bioinformatic analysis using the cellranger-dna pipeline to call copy number profiles at single cell resolution (**Fig. 1B**). The library had a median read duplication ratio of ~10% per cell, with on average 1,358,777 effective mapped deduplicated high-quality reads per cell. However, copy number profiling is performed by cellranger-dna at single cell resolution and is thus inferred from an average of 436 reads/Mb (**Supplementary Fig. 2B**). Notably, while the average genome-wide ploidy of single cells was tightly distributed around a median ploidy of 3, the individual CNV profiles were different between cells (**Supplementary Fig. 3 and 10**). For example, cells at similar sequencing depths and average ploidies, cell #596 with 448 reads/Mb and a mean ploidy of 3.03 and cell #415 with 472 reads/Mb and a mean ploidy of 3.02 (**Supplementary Fig. 3**) exhibit extensive copy number changes. At a 2Mb resolution cell #596 has three distinct copy number states in chromosome 1 (3-2-4), two in chromosome 18 (2-3) and a single ploidy of 3 across chromosome 8; in contrast #415 exhibits two copy number states (2-4) in chromosome 1, while chromosome 18 is a single segment at copy number 2 and chromosome 8 is at a different copy number of 4 (**Supplementary Fig. 3**). Thus, despite uniform average genome-wide ploidies single-cell resolution revealed cells having different copy number profiles in a single growth of COLO829 (**Supplementary Fig. 4**).

**Figure 1.**
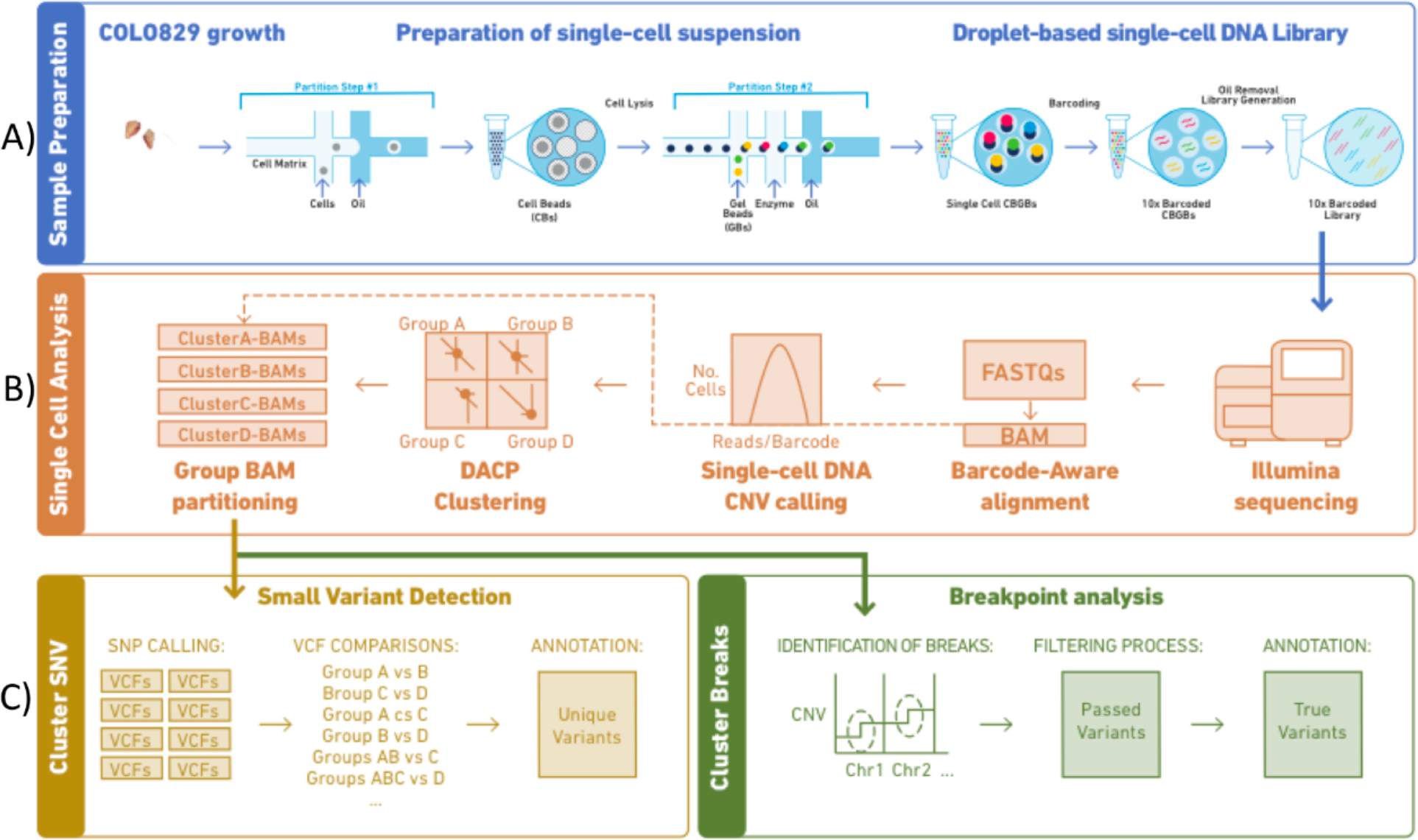
Workflow of the single cells sample and data processing. **A)**Sample preparation of the chromium technology. **B)**Single Cell Analysis. Data processing from sample sequencing to BAM partitioning. **C)**Data analysis including variant detection and breakpoint analyses.

### COLO829 is composed of four major clusters

We leveraged CNV events that were observed in this population to identify sub-clones and cluster single cell CNV data. For this and all other downstream analysis we excluded 6% of the cells that were flagged by the cellranger-dna pipeline as “noisy” and focused on the remaining 1,373 single cells. First, we filtered the raw CNV events identified by cellranger-dna by applying a size cutoff of >2 Mb and a quality cutoff of 15. Next, we derived a binary CNV event matrix tabulating the absence/presence of a CNV event across the 1,373 single cells, see methods -clustering of single cell CNV data. This resulted in 114 CNV events with a majority <100 Mbp with ploidies of 2, 3 and 4 (**Supplementary Fig. 5**). Clustering was performed using the adegenet R package which uses Discriminant Analysis of Principal Components (DAPC), a multivariate method that attempts to identify groups of genetically related individuals by constructing linear combinations of the original alleles (which in our case are CNV events) that have the largest between-group variance and the smallest within-group variance (**Fig. 2A**, **Supplementary Fig. 6**). For this analysis we chose a clustering solution of k=11 guided by a BIC curve with the optimum ranging between 10 to 15 (**Fig. 2B**). Plotting the single cells on the coordinates of two primary DAPC axes revealed four distinct groups (A-D) of cells (**Fig. 2C**), with sub-structure within two of them (**Fig. 2C**).

**Figure 2.**
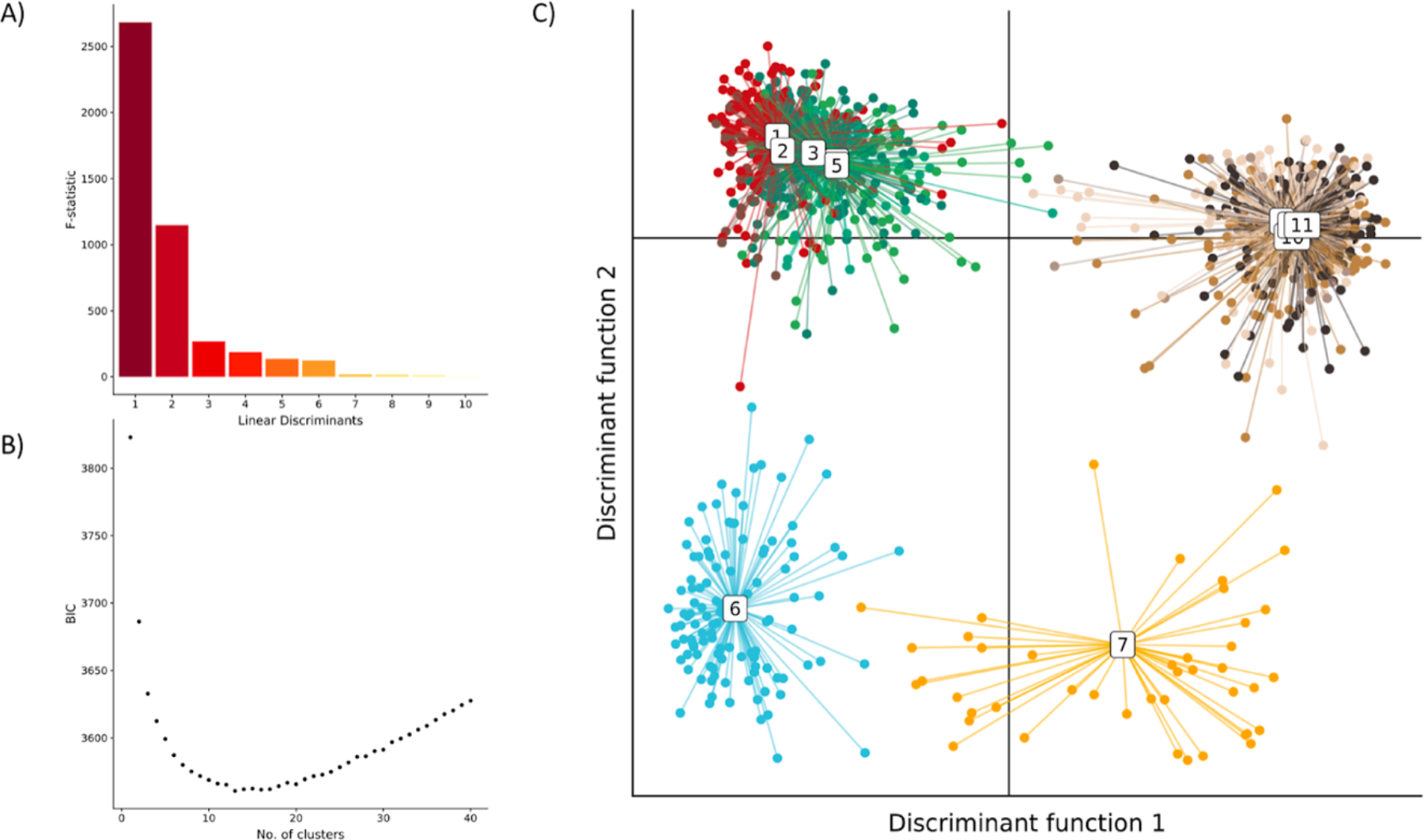
Clustering of COLO829 single cells using Discriminant Analysis of Principal Components (DAPC). **A)**Bar plot of eigenvalues, which correspond to the ratio of the variance between groups over the variance within groups for each discriminant function. **B)**Inference of optimal cluster number using Bayesian Information Criterion (BIC). **C)**Scatterplots showing the inference of population structure in 1,373 cells using the first two principal components of the DAPC analysis. Individual dots represent single cells and the color represents cluster assignment.

Visual inspection of single cell CNV heatmap reveals striking clusters of chromosome-scale differences between cells (**Fig. 3**). Shown in Figure 2, the DAPC clustering analysis indicates four major groups: *Group A* (653 cells), *Group B* (117 cells), *Group C* (43 cells), and *Group D* (560 cells). *Group A* and *Group B* are distinguished by a copy number of 3 on chromosome 8, whereas *Group C* and *Group D* have four copies of chromosome 8. *Group B* and *Group C* showed a loss of chromosome 1pter-1p22.3, chromosome 10p14-p11.22, and chromosome 18. Additional events are evident including on chromosomes 11 and 6, though we focus on the large-scale chromosomal differences evident between these four groups. A previous study, comparing 4 different cell-line growths through bulk sequencing analysis observed a similar 1p loss in one sample (referred to here as the TGen growth). To gain a better understanding of the relationship between these events, we created four group-level BAMs, one per major DAPC group to enable additional bulk-format analysis.

**Figure 3.**
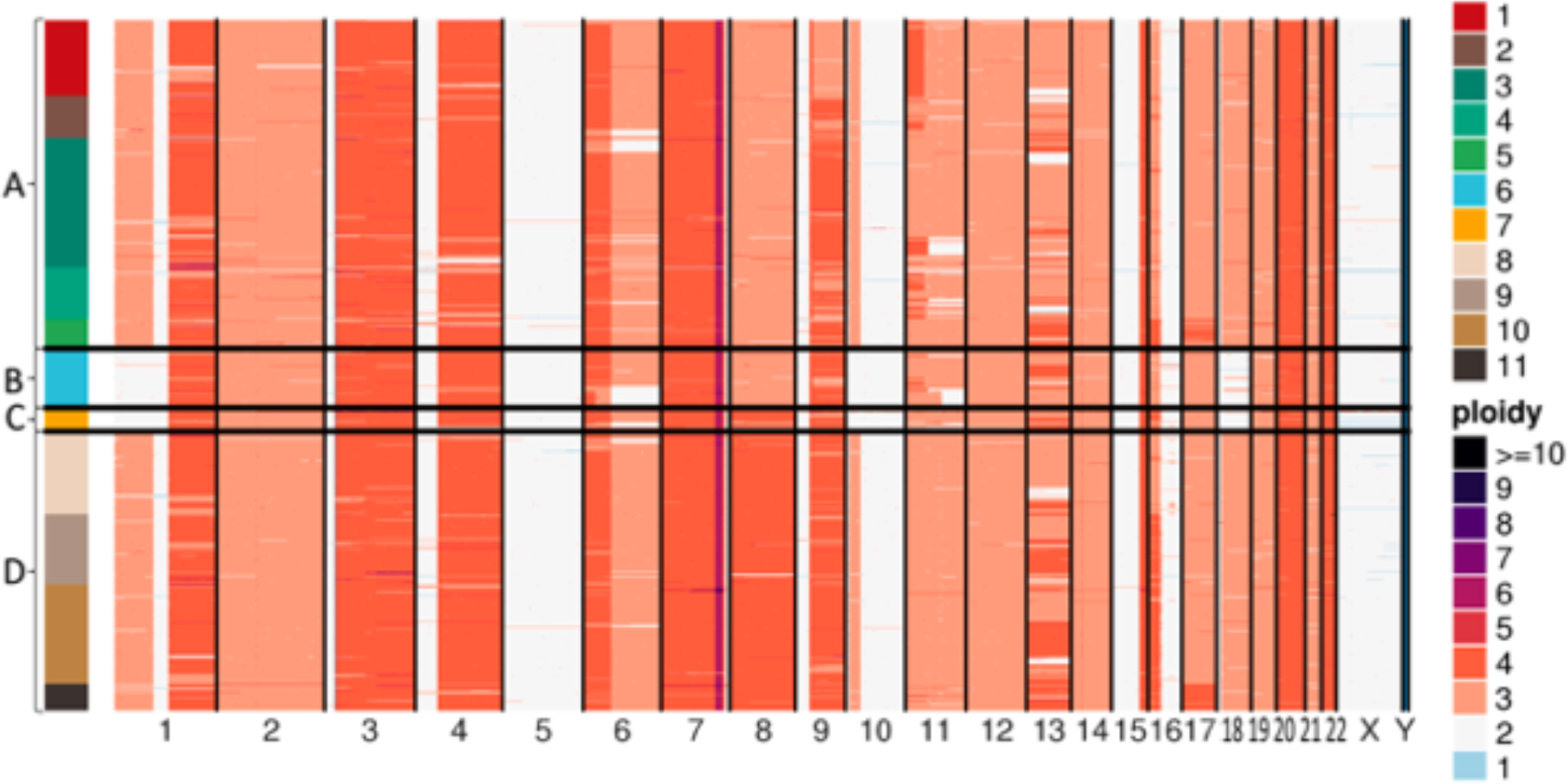
Single cell copy number heatmap of COLO829. Heatmap showing hierarchical clustering of 1,373 single cell CNV profiles at 2 Mb resolution. Each row depicts the whole genome of a single cell, colors (Group A: clusters 1 - 5 [red, brown, greens], Group B: cluster 6 [blue], Group C: cluster 7 [yellow], Group D: clusters 8 - 11 [browns]) represent the called ploidy as specified by the legend on the right and rows are clustered by groups (Group A: 653 cells, Group B: 117 cells, Group C: 43 cells, Group D: 560). Hierarchical clustering was performed calculating the distance between each single cell CNV data to posteriorly join them into groups. The 11 clusters (upper right) were calculated from the inference of optimal cluster number analysis. Ploidy number (lower right) is represented by distinct colors.

### Insight from Loss of Heterozygosity and break-point analysis

LOH and breakpoint analyses were conducted for each group-level single-cell sequencing BAM. For this analysis, we compared all combinations of the four main groups (*Groups: A, B, C & D*) to four bulk sequencing runs from Craig *et al* (TGen, GSC, Illumina and EBI bulk sequencing of growths) to see if hallmark events visible in the bulk sequencing of prior studies were historical events shared by the single cell data, or whether we were identifying newly emerging events.

We first performed breakpoint analysis by mapping anomalous read-pairs in order to identify the structural variants behind copy number changes. Specifically, we expect paired-read mapping to show how copy number segments are joined together. Analysis and identification of clusters of anomalous readsets was determined as described in the methods. To insure adequate power even in groups with low cell counts, we focused on anomalous read-pairs that aligned to regions within 2Mb of the median breakpoint locations of the 114 shared CNV events identified in the above analysis. Break-point analysis on aggregated data showed DAPC group-specific anomalous read patterns that indicated clusters of breakpoints involving chromosomes 1, 10, and 18 indicating translocations of 1p22 (GRCh37 chr1:87,337,015) to 10p14 (GRCh37 chr10:36,119,061), and from 10p11 (GRCh37 chr10:7,634,373) to 18p11 (GRCh37 chr18:9,868,810) (**Supplementary Fig. 10).** In fact, exact breakpoints can be mapped to base-pair level resolution using reads that span the junction boundary (shown in detail in **Supplementary Fig. 10**). These events were observed in *Group A* and *Group D*, but not *Group B* or *Group C*. Examination of the bulk sequencing growth found translocations in a subset of reads for the EBI, GSC and Illumina growths, but not the TGen growth. Considering the location of the copy number breakpoints it is likely derived from an abnormal chromosome 18 containing portions of chromosome 1p and 10p, designated as (der18)(1pter->p22∷10p14->10p11∷18p11->18q). This abnormal chromosome 18 then is lost in both *Group B* and *Group C*.

To expand on this observation, LOH analysis was conducted by examining the allele fractions of germline SNPs known to be heterozygous within COLO829 lymphoblastic cell-line, which has previously sequenced multiple times. LOH provides information about the ratio of parent of origin for copy number events, and importantly is independent of the original clustering analysis. In order to ensure higher quality variants, we used quality filtered heterozygous SNPs known to have a population-based minor allele frequency above 1% used. Shown in **Fig. 4** and **Supplementary Fig. 8**, the allele fraction for each SNP is plotted within the COLO829 DAPC-defined groups and independent growths of the cell-line. As expected, groups with fewer cells have greater noise in their allele fraction. However, by taking the average of multiple SNPs across a segment sub-clone specific patterns of LOH become evident. For example, the p-arm of chromosome 1 is determined to be at a copy number of 3 in *Group A* and exhibits B-allele frequencies of 67/32 consistent with a heterozygous triploid genetic background, while *Group B* for the same region is at a copy number of 2 and B-allele frequencies of 98/1.1 suggesting that chromosome 1p is homozygous diploid for one of the parents. Similar analysis of the data from Craig *et al*, we identified that TGen growth lost an original haplotype where EBI, Illumina, and GSC growth had multiple copies of one of their ploidies and still a single copy of the chromosome 1 translocation. Notably, we observed that *Group B* continues to mirror the TGen lineage. In this example, it is clearly observed evidence from the homozygous deletion in TGen where the clustering was biologically correct **(Supplementary Fig. 10)**.

**Figure 4.**
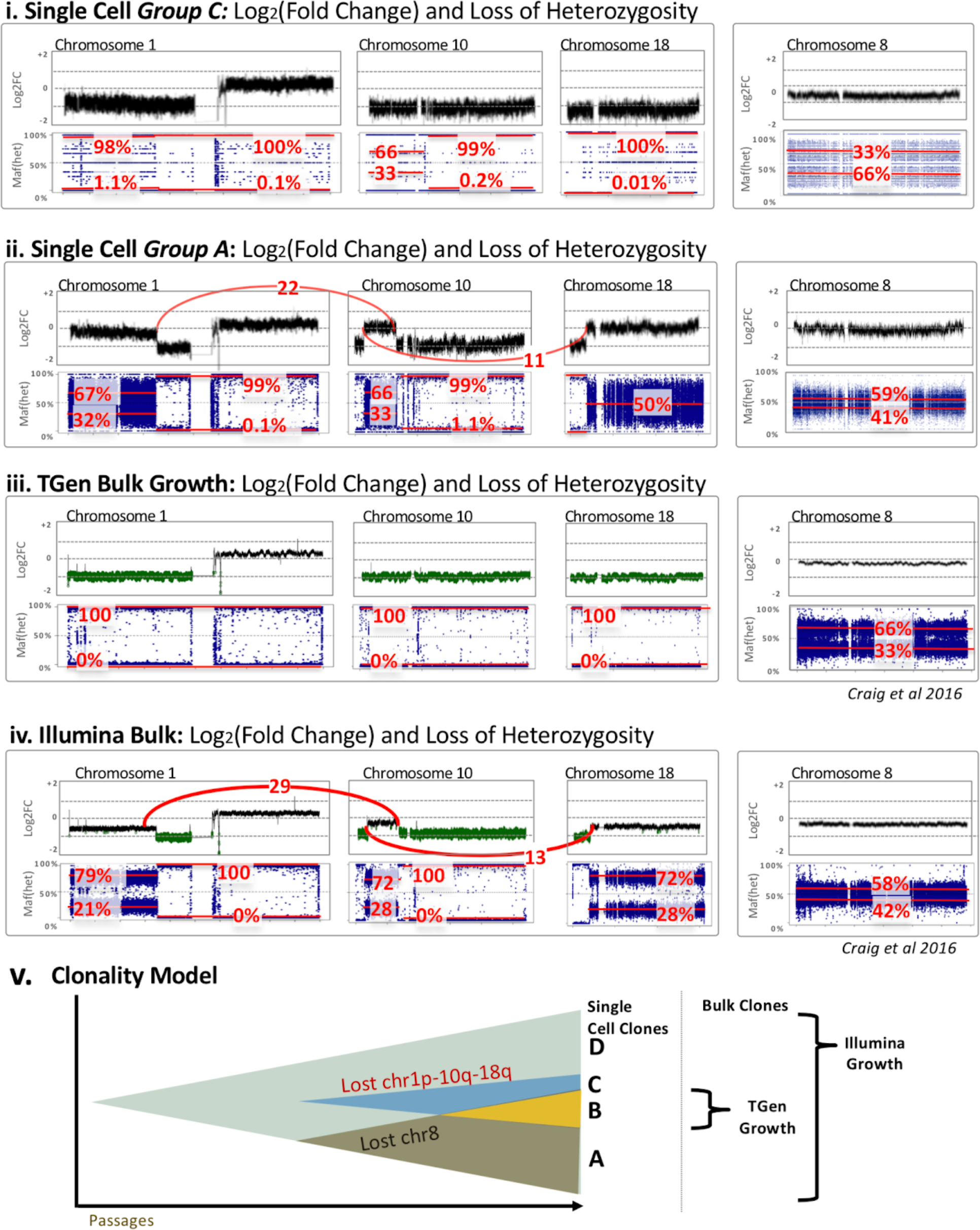
Log_2_Fold*Δ* (upper in each) and Het SNP allele frequency (lower blue in each) for (i) Group C and (ii) Group A, (iii) Illumina Bulk Growth, and (iv) TGen Bulk Growth. The upper graph of each panel provides estimated log2 fold change (noting bulk copy number does not inherently produce copy number estimates). For the chromosome 1, 10, and 18 via (p22.3;10p14) and t(p11.22;18p11.22), counts of anomalous reads supporting the junction are shown in red, whereas this event is absent in Group B and C, as well as the TGen Growth. The lower plots of each panel are the allele fraction of known heterozygous SNPs (identified from previous VCFs in the germline lymphocyte lines) for COLO829. Their deviation from the expected 50% allele fraction provides an indication of Loss of Heterozygosity (LOH), where the relative noise is dependent on the number of reads over a SNP and greater spread is observed in groups with fewer reads. The median major/minor allele fractions are provided for each region in red. (v) Schematic model of the major clones shows a simple model whereby the sub-clones emerge as some cells do not maintain the abnormal chr1p-10q-18q line and/or an additional copy of chromosome 8.

More broadly, it’s evident that the TGen growth has several parallels to one of the single-cell groups identified in this study, both at chromosome 8 and the abnormal (der18)(1pter->p22∷10p14->10p11∷18p11->18q). Referring first to the abnormal chromosome 18, LOH and copy analysis **(Fig. 3 & 4)** demonstrates these regions are 3 copies and not homozygous, while the rest of these chromosomes are at 2 copies with LOH (**Fig. 4**). This model suggests that as the cells are grown, certain groups of cells lose the abnormal chromosome 18, while others lose the less stable chromosome 8.

### Validation from spectral karyotyping

We compared the major features determined from single cell DNA analysis with previously generated karyotyping, of an independent sample, observing remarkable agreement **(Supplementary Fig. 9)**. All the major rearrangements seen in the karyotype correspond to copy number changes and LOH in the single cell analysis. The der(?)t(1p?;18q?) of the karyotype is consistent with the (der18)(1pter->p22∷10p14->10p11∷18p11->18q) postulated from sequencing. The der(?)t(1;3)(q?;p22-24?) × 2 exactly fits the 4 copies of 1q and most of chromosome 3 up to 3p2. The iso(4q) matches the two copies identified in single cell samples. The two copies of der(7)t(7;15)—most of chromosome 7 with small piece of 15 attached to distal 7q— matches the copy number 2 of the tip of 7q (presumably with LOH of the tip) and 4 copies of 15qter, suggesting that it is der(7)t(7q[near end];15q[near ter]), formed before duplication. The 6 copies of the region of 7q adjacent to the break is most likely a duplication of this region in the translocated copies. The 4 metaphases where one eleven is replaced by a der(?)t(11;18), may correspond to a few cells that have lost the 1;10;18 and that have a translocated 11 broken mid 11q. The del(6), del(9), and del(16) are consistent with the copy number losses seen on these chromosomes. The del(9) and del(16) are present in 1 or 2 copies, thus probably formed before duplication and some cells losing one copy. For the del(6), the single cell sequencing detected 3 copies of normal 6, apparently with allele ratios 1:2. This suggests that the deletion occurred after duplication, and some cells subsequently lost a normal 6, as in the karyotype. Overall, combining the single cell sequencing with the karyotype enabled us to construct a plausible evolutionary history of the line. Both the karyotype and single cell sequencing suggest that the cell line duplicated its entire karyotype after most of the rearrangements seen. The translocations—the 1;3, 1;10;18 and 7;15 translocations—fit the Dutrillaux monosomic pattern of karyotype evolution, in which two normal chromosomes are replaced by one unbalanced translocation, resulting in copy number loss and (if before genome duplication) LOH^6,11^.

## DISCUSSION

In this study, we resolve sub-clonal heterogeneity in growths of COLO829 by performing single cell DNA sequencing to identify major sub-clones. These major sub-clones further enabled LOH and breakpoint analysis, providing a clearer picture of clonal heterogeneity.

Numerous sequencing studies have utilized this line as a control, and a few have indicated some evidence for underlying heterogeneity of the line. Most importantly, we identified sub-clones with unique hallmark features that provide a potential explanation of previously reported variability in growths of COLO829. We observe an abnormal chromosome providing an extra copy of a suggestive break on chromosome 18p that is consistently maintained in some daughter cells. Similarly, we observe that in some cells there is also a loss of chromosome 8. Notably, only the single cell methods provide clear insight into the underlying diversity of the cell-line. In addition, it should be remarked that SKY data presented here were available via an online resource and played an essential role in validating the initial findings by single-cell copy number.

A key aspect of the single-cell sequencing of DNA is the efficiency and accuracy of the clustering analysis to create clonal groups of cells, enabling downstream analysis leveraging bulk analysis tools. Many of these bulk methods specific to tumor or clone analysis, such as LOH and breakpoint analysis rely on finding events unique to a clone. If a cell is incorrectly placed in the wrong group, these algorithms become largely unreliable. While not a key role, there is a clear path whereby one could see identifying mutations specific to a clone. This is more likely the case in patient samples, as is frequently shown in heterogeneous cancers such as colon and prostate cancer.

A major general and far-reaching observation is also made of how single-cell shallow sequencing is a starting, enabling further iterative analysis of breakpoints and SNPs, and while not shown in this study mutation specific clones. While in this study we show how LOH analysis enables characterizing clones, it is clear from these studies that much further algorithm development is possible. In general, despite numerous papers identifying some aspects of the sub-clonal heterogeneity within this cell-line, they rendered a fragmented view confounded by the inability to couple disparate structural and mutational events to single clonal genomes. Here, we observe a remarkable agreement between the single cell sequencing samples and SKY analysis with a more comprehensive copy number changes of this cell line. Single cell resolution does precisely that, and allows us to synthesize a bigger picture of what is driving the events.

## Accession codes

Data have been deposited in the following 10X Genomics hosted link: https://support.10xgenomics.com/single-cell-dna/datasets/1.0.0/colo829_G1_1k.

## ACKNOWLEDGEMENTS

This work was supported by 10X Genomics and the Department of Translational Genomics, Keck School of Medicine of University of Southern California.

## AUTHOR CONTRIBUTIONS

E.I.V., S.M. D.W.C, J.S. and C.C. contributed to the experimental design. C.C. and Y.Y. conducted the experiments. S.M., E.I.V., D.W.C, M.W., V.K. and Y.Y. contributed to the data analysis. M.G. and P.A.E. contributed to the Spectral Karyotype analysis. E.I.V., did the writing. E.I.V., S.M., J.S., I.T.F., V.K., Y.Y., M.W., C.C., J.D.C., D.W.C. oversaw all aspects of the manuscript.

## METHODS

### Preparation of single-cell suspension

COLO829 cell line was obtained from American Type Culture Collection (ATCC), Manassas, VA. Cells were cultured in their recommended media conditions at 37°C. Prior to FACS sorting, the cancer cell line was trypsinized, followed by inactivation with FBS and washed by centrifugation at 300g in 1X PBS with 0.04% BSA. Cells were counted and resuspended in recommended media at a final concentration of 1e6/mL in a FACS tube. 2 μL of Vybrant® DyeCycle™ Green stain was added to cell suspension and incubated at 37°C for 30 minutes in the dark. Cells were then analyzed and sorted on a flow cytometer using 488 nm excitation and green emission gating on cells in the G1 phase of the cell cycle (**Supplementary Fig. 1**). Cells were counted post sorting to ensure accurate concentration.

### Single-cell DNA library generation

Single-cell suspension was processed using Chromium Single-Cell CNV Solution (10× Genomics) as described in the user guide to generate a barcoded single cell DNA library (**Fig. 1A**). Single cells were partitioned in a hydrogel matrix by combining with a CB polymer to form Cell beads (CBs) using a microfluidic chip. Post a first encapsulation, CBs are treated to lyse the encapsulated cells and dentaure the genomic DNA (gDNA). The denatured gDNA in the CB is then accessible to amplification and barcoding. A second microfluidic encapsulation step is required to partition the CB with 10× barcode Gel Beads (GBs) to generate an emulsion called GEMs. Immediately after barcoding and amplification, 10× barcoded fragments were pooled and attached to standard Illumina adaptors. Finally, sequencing libraries were quantified by qPCR before sequencing on the Illumina platform using NovaSeq S4 chemistry with 2×100 paired-end reads (**Fig. 1B**).

### Single-cell CNV calling using Cell Ranger DNA

Paired-end reads were processed using version 1.0 of the cellranger-dna pipeline (10× Genomics)^8,9^. As described previously, the pipeline consists of barcode processing, alignment to the (hg19) genome and the identification of cell-associated barcodes. Copy number calling is performed on each barcode separately after masking out regions of the genome with low mappability and normalizing for GC content. 1,475 barcodes were defined as cells, roughly all barcodes with greater than 1/10th the number of reads as the maximum per-barcode read count. Cells flagged as noisy by the pipeline (102 cells, 6.9%) were removed from downstream analysis, leaving behind 1,373 cells.

### Clustering of single cell CNV data

Single cell CNV calls were extracted from a BED file generated by the cellranger-dna pipeline for 1,373 cell barcodes. Events were filtered to include those with a size >= 2Mbp and with a confidence score > 15. Events from different single cells were grouped together if they had identical copy number and shared 90% reciprocal overlap. Next, events present in less than 5% of cancer cells were discarded. This analysis generated a binary CNV mutation matrix with 112 polymorphic events ranging in size from 2.1 to 147.6 Mbp and the custom R script used to perform this analysis is included as **Supplementary File 1**.

To identify clusters we implemented the fast maximum-likelihood (ML) genetic clustering and Bayesian Information Criterion (BIC), subsequently, using the Bioconductor adegenet package (version 2.1.1). BIC curve suggested between 10 to 15 clusters, where 11 (k=11) were selected as the optimal clustering solution (**Fig. 2A & B**). This yielded 4 major groups with sub-structure within two of them (**Fig. 2C**). A list of barcodes per major group was generated (**Supplementary Table 3**).

### Bulk Copy Number and Loss of Heterozygosity analysis

A python script was used to split the BAM file by barcode assignment, generating a BAM file for each sub-clone (**Supplementary File 2**). WIth cellranger-dna version 1.1 this functionality is a new sub-pipeline. The new BAMs only include reads from barcodes that are assigned to that cluster to enable downstream analysis with traditional mutation calling tools developed for bulk-data.

Fold change and Loss of Heterozygosity (LOH) analysis was performed using previously developed tools, tCoNuT(1.0) (https://github.com/tgen/tCoNuT) and DNACopy (version 1.48.0) for both the single cell grouped BAMs, and for previous bulk sequencing of COLO829 growths^2^. The previous bulk sequencing was conducted by 4 groups with independent growths of both COLO829 melanoma line and the paired COLO829BL germline lymphoblastic cell-line. Consistent with prior publications, these are referred to as the TGen, EBI, GSC and Illumina growths. Use of additional copy number analysis tools provided a framework for comparing aggregated sub-clone data to previous bulk sequencing, and added additional analysis capabilities such as loss-of-heterozygosity.

LOH was examined using germline heterozygous SNPs for COLO829 using the companion COLO829BL. Specifically, heterozygous germline variants were identified using GATK’s HaplotypeCaller (version 3.5.0) from previous sequencing data^51^. The VCFs were annotated with dbSNP 147 using snpEff (version 3.5h) and input to the tCoNuT parseMergeVCF script (1.0) to output heterozygous SNPs. Heterozygous SNPs were identified along with reads supporting alternative and reference alleles. Only SNPs within a range of 0.4 to 0.6 allele fractions were utilized from germline 80× whole genome sequencing data. Whereas tCoNuT converts to an absolute minor allele fraction, figures are shown to span allele fractions from 0 to 1, and both minor and major allele fractions are provided in figures. Using allele fractions, variants near 0.5 still retain alleles from both the maternal and paternal haplotypes, whereas as those nearing 0 and 1 have lost this heterozygosity. Intermediate levels can often be interpreted across a range, such as 0.66/0.33 allele fractions with copy number of 3. The median allele fractions for the minor/major allele were obtained across copy number segments within the single cell CNV bed file. These regional LOH values are shown in Figure 4 with red text for the region spanned by the red line.

### Variant detection and breakpoint analysis

For this analysis, we utilized a previously validated script for detection of anomalous read pairs (tgen_somaticSV) to identify clusters of read-pair mappings consistent with translocations, inversions, and other structural variants (https://github.com/davcraig75/tgen_somaticSV) ^5^. The tool defines donor and acceptor regions, and counts read-pairs supporting each, where each donor/acceptor region spans no more than 3× the insert distance and is greater than 10,000bp in seperation. A reference set of reads are required, similar to a tumor/normal set, and for this analysis we utilized the other groups or the COLO829BL reference line. As the key events were in two of the groups, the former method did not yield meaningful results, and the key region was identified in comparison to references. Additional filters included requiring reads supporting both directions, e.g. determined by first read and mapping quality >20. Anomalous paired reads clusters were sorted by number of reads, and examined for reads within 2 megabases of a CNV breakpoint. In some cases, more than one read cluster was evident and in those cases we prioritized those nearest to change in copy number.

We attempted to identify point mutations specific to clones. Briefly, we called unique variants per major group by systematically comparing 4 groups using Strelka2 small variant caller and filtering the output using the following criteria: Filter=PASS and QSS_NT > 30. In addition, we calculated the transition transversion ratio and visually examined variants within IGV, filtering out variants known to be present. Examination of coding variants did not yield any high quality novel point mutations specific any clone and not within the original COLO829 line. The lack of novel mutations in this hyper-mutated line might be because it is a cell-line, rather than each clone resulting for clonal evolution as in the case of patient-derived tumors where the following methods have been found effective.

