## Supplemental files for "Resolving sub-clonal heterogeneity within cell-line growths by single cell sequencing genomic DNA"

### INSTITUTIONS

[1]Department of Translational Genomics, Keck School of Medicine of University of Southern California, Los Angeles CA, United States

[2]10X Genomics, Pleasanton CA, United States

[3]Cancer Research UK Cambridge Institute, Cambridge, United Kingdom

[4]Hutchison-MRC Research Centre, University of Cambridge, Cambridge, United Kingdom

### CORRESPONDING AUTHORS

Enrique I. Velazquez-Villarreal

David W. Craig

**TABLE OF CONTENTS**


| | |
|---|---|
| **Supplementary Figures** | Pages 1 - 11 |
| **Supplementary Tables** | Pages 12 - 64 |
| **Supplementary Documents** | Pages 65 - 67 |

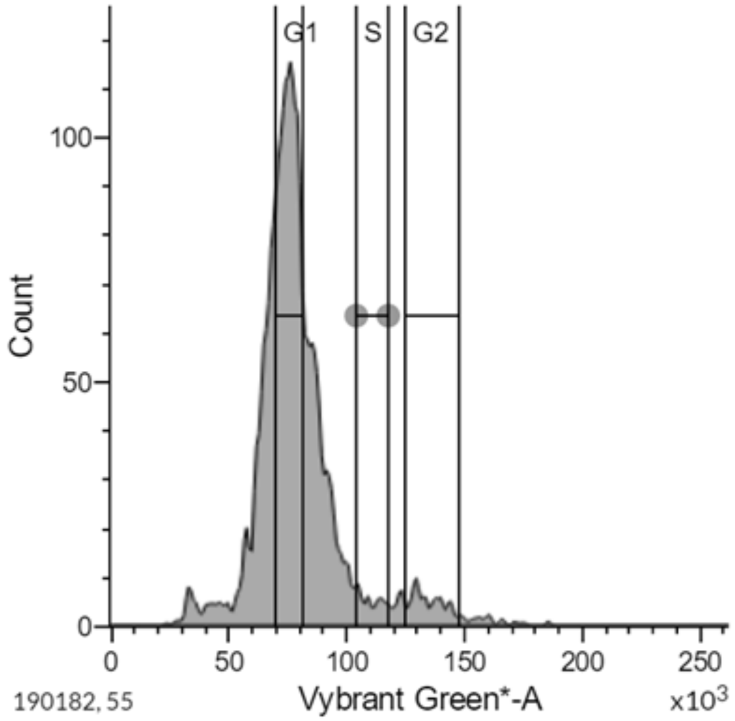


**Supplementary Figure 1 | Distribution of DNA content in COLO829 nuclei by flow cytometry.** Cells in the G1 phase were selected for scCNV library preparation.

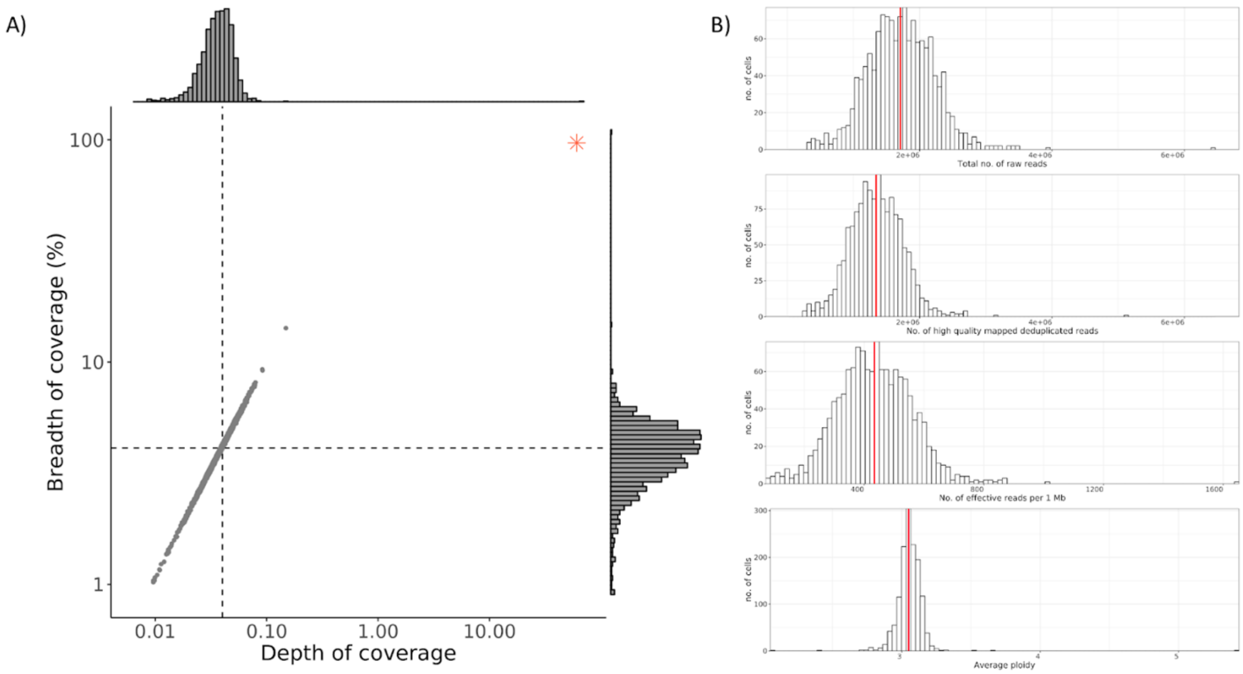


**Supplementary Figure 2 | Sequencing depth . A)** Depth of coverage versus breadth of coverage. A scatter plot depicted the effective depth of coverage (x-axis) versus the percent of the genome with at least 1x coverage (y-axis) for each cell. The dashed lines represent the median single cell values (0.04 depth of coverage, 4.11 breadth of coverage). The red asterix plots the values for aggregated data as a pseudo bulk experiment (60.68x depth of coverage, 95.97% breadth of coverage). **B)** Histogram of summary data quality metrics. Histogram depicting the distribution of per cell summary metrics with the median indicated by the red line: Total number of raw reads (median value 1,709,262); Number of high quality mapped deduplicated reads (median value 1,343,083); Number of effective reads per 1 Mb (median value 434) and Average ploidy (median value 3.048).

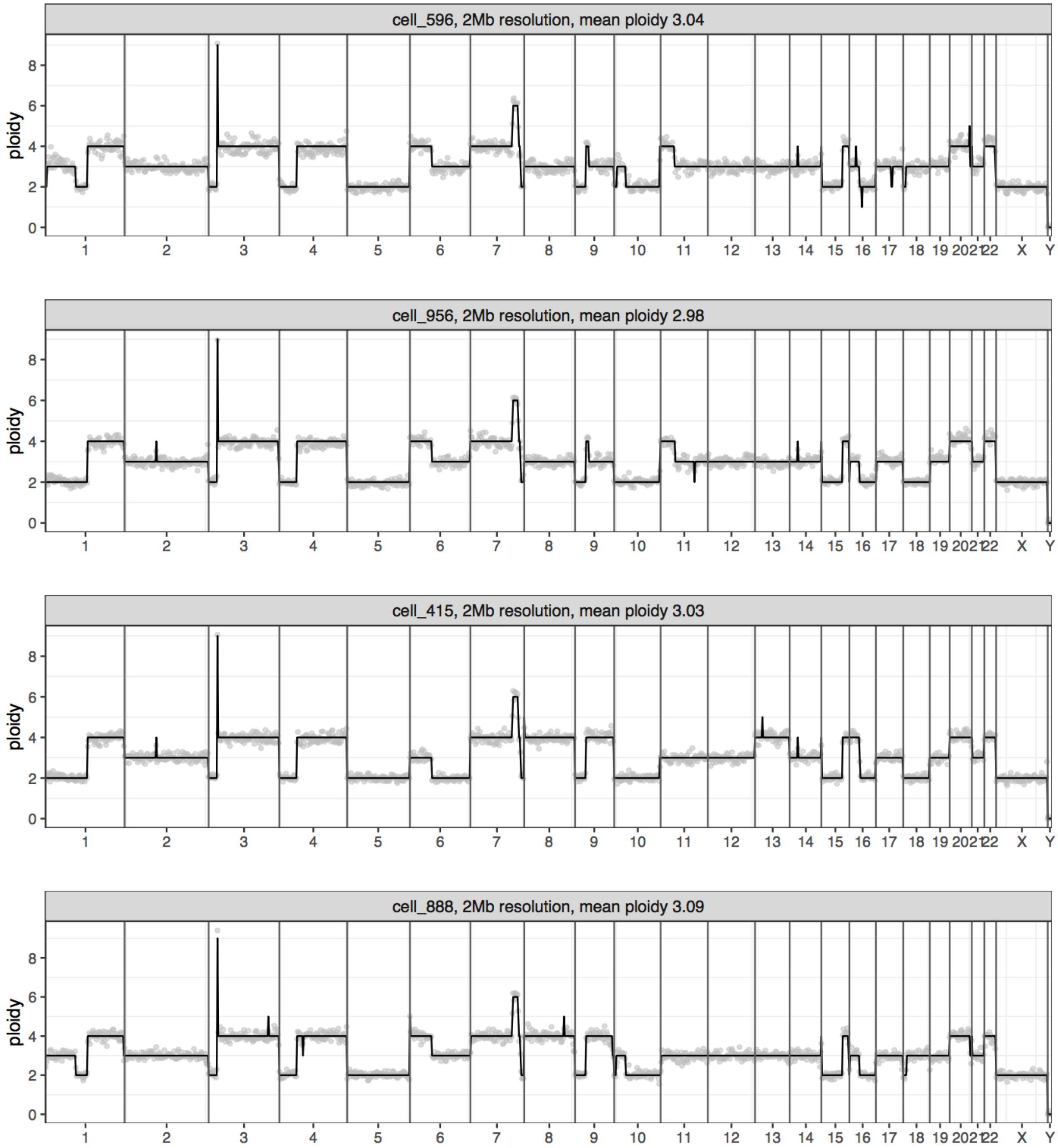


**Supplementary Figure 3 | Four representative single cell ploidy plots.** Copy number profiles plotted at a resolution of 2 Mb bins. Solid black line indicates ploidy call and gray dots indicate raw read counts for each bin.

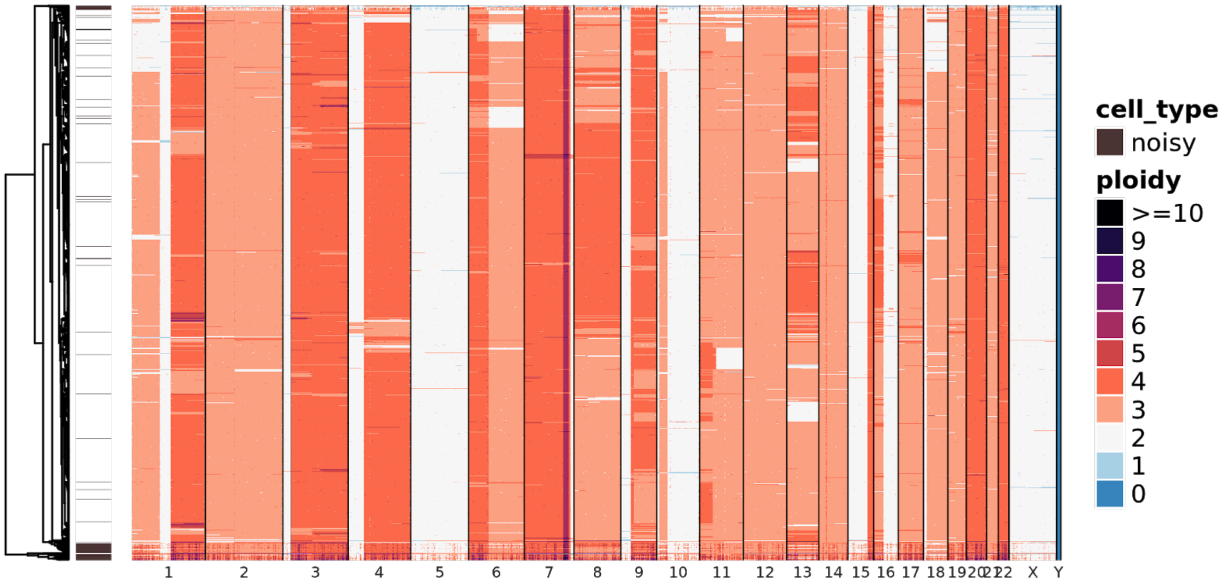


**Supplementary Figure 4 | Heatmap of raw data**. Heatmap showing hierarchical clustering of 1,475 single cell CNV profiles at 2 Mb resolution. Each row depicts the whole genome of a single cell and colors represent the called ploidy as specified by the legend on the right. Color-bar on the right marks single cells labeled by the pipeline as noisy in brown.

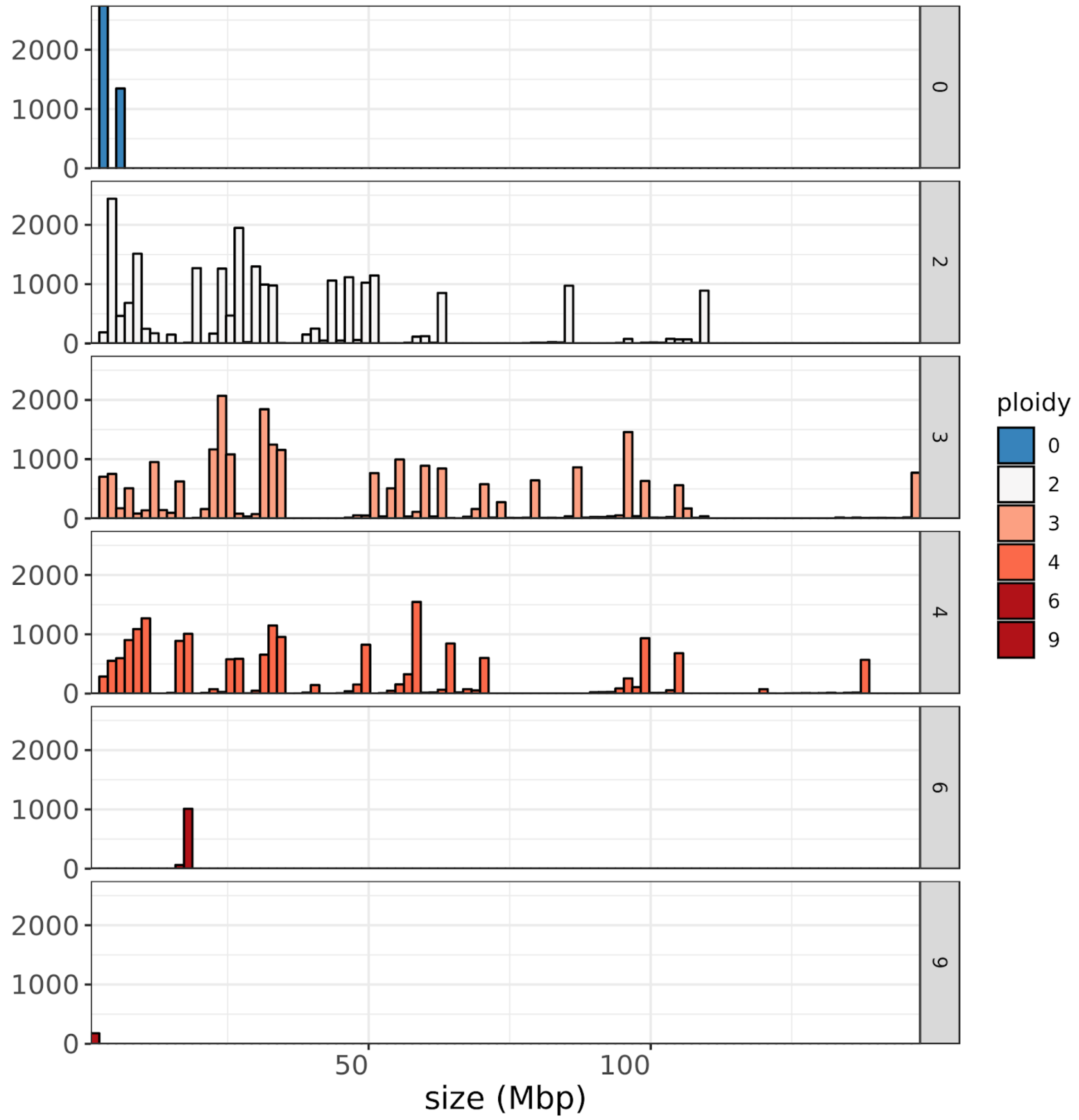


**Supplementary Figure 5 | Size distribution of CNV events by ploidy.** Histogram of the 114 CNV events that passed the following filtering criteria: event quality > 15; event size > 2 Mb and event frequency > 0.05.

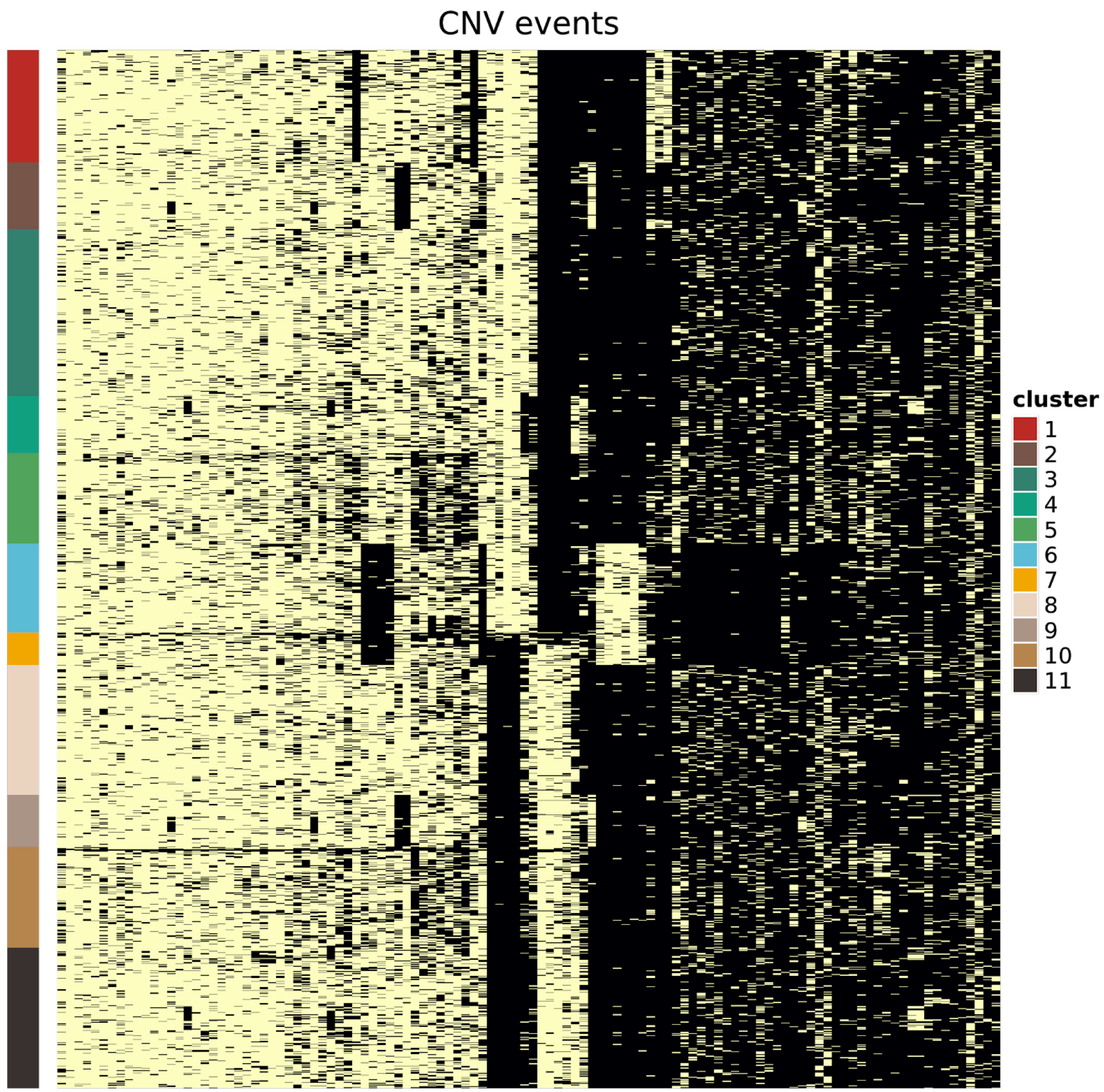


**Supplementary Figure 6 | CNV events within each cluster**. Heatmap with one row for each cell and one column for each of the 122 polymorphic CNV events that passed filtering, ordered by their frequency in the population. The cells are ordered by membership in DAPC clusters. The presence or absence of a mutation is depicted in yellow and black, respectively.

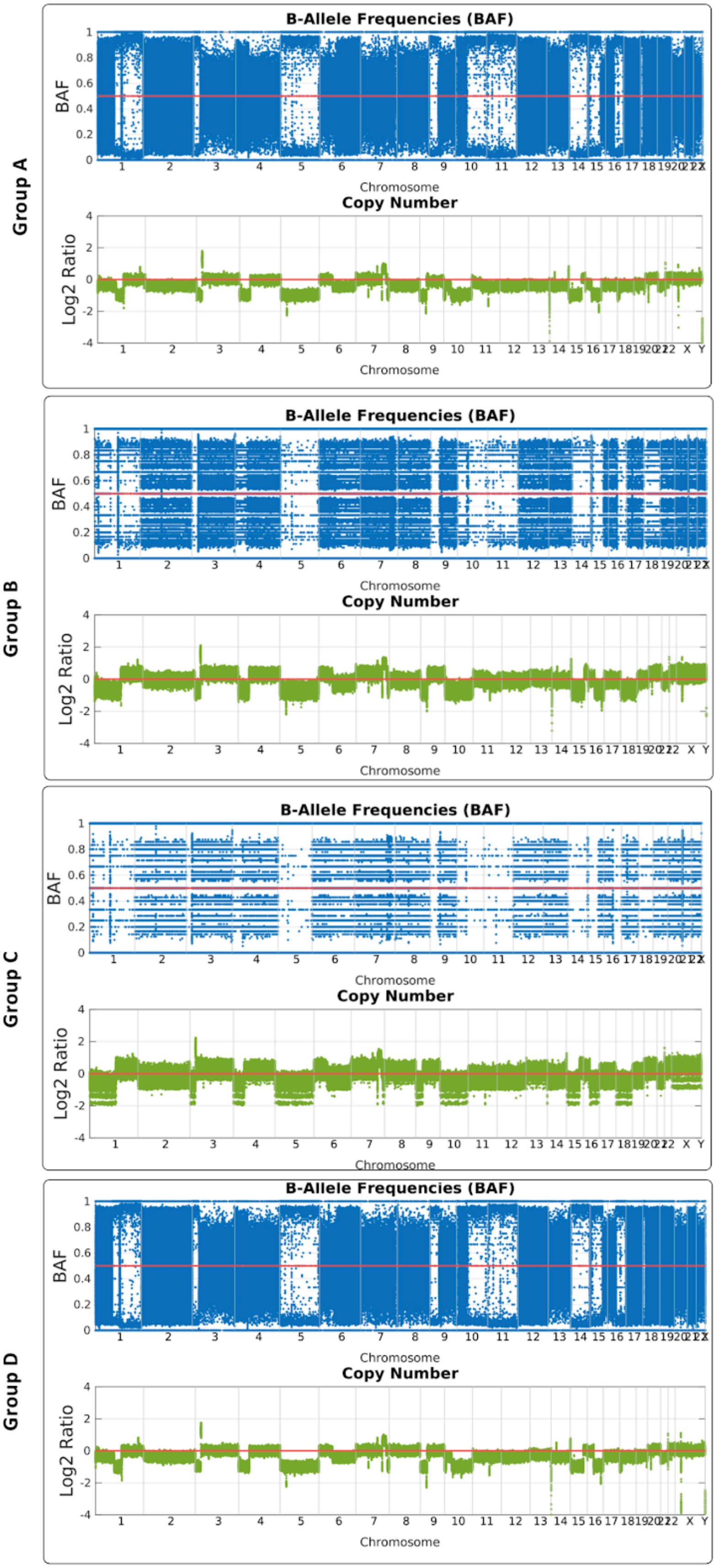


**Supplementary Figure 7 |** Log2(Fold Change) and LOH for each group. The upper graphs show the Log2(Fold Change) using the program tCoNuT, which was used previously for bulk sequencing and produces allele fraction plots. It is important to note that that tCoNuT, like other bulk copy number tools cannot determine copy number natively without assumptions for deconvoluting potential mixtures. Single cell copy numbers as used elsewhere do not require this assumption and absolute ploidy can be better inferred. The lower graphs provide the allele fractions of known heterozygous SNPs determined previously from sequencing on the lymphoblastic germline pair. For example in *Group A* at chr5 when copy number is 1, we see allele fractions partitioning into 0 and 1 allele fractions indicating homozygosity. The relative noise is dependent on number of reads over a SNP, and thus greater spread is shown. SNPs across a region or segment could be averaged together to get a regional LOH, value, though individual SNPs are shown in these examples. In the figure above we observe loss of heterozygosity (LOH) for chromosomes 5, 10, 11 and 14. We also see some chromosomes with partial LOH. Within the main figure 4, chromosome 1, 10, and 18 are shown, whereas this figure shows all chromosomes.

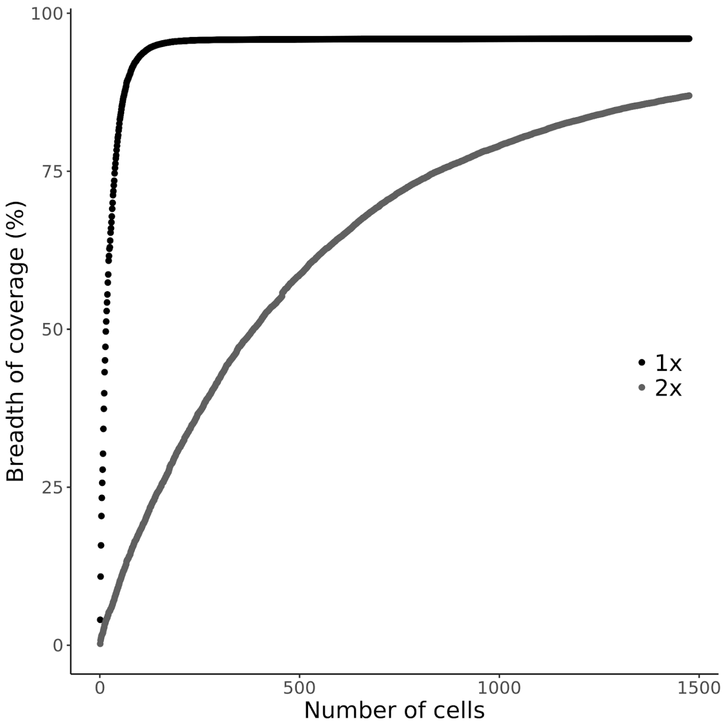


**Supplementary Figure 8 | Cumulative breadth of coverage**. A cumulative plot of genome coverage at two different depths 1x and 2x (black and gray, respectively). Number of cells needed to achieve 75% coverage across the genome is 37 at 1x depth and 845 at 2x depth.

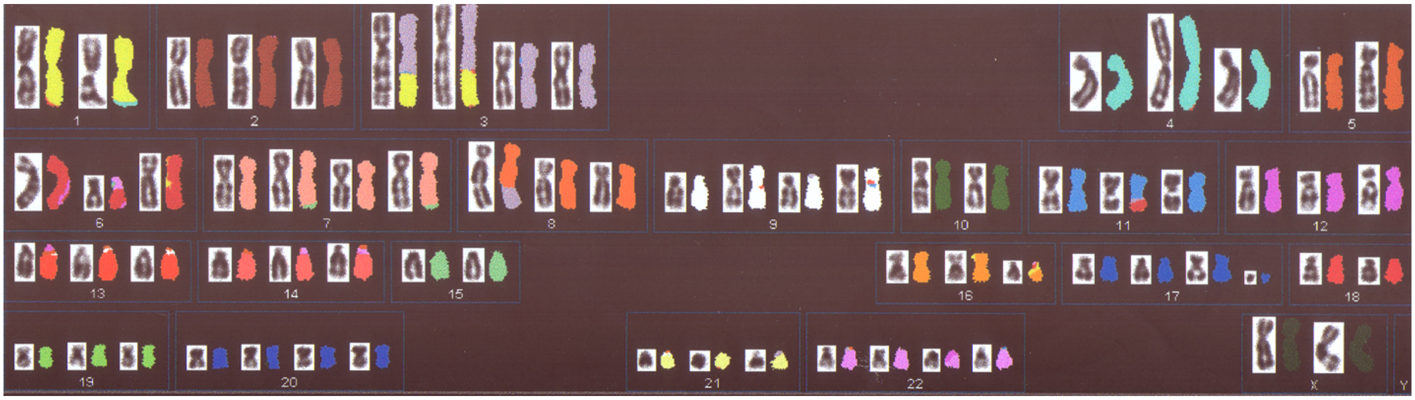

**Supplementary Figure 9 |  Spectral Karyotype (SKY) of COLO829 cell-line**. SKY multicolour-FISH Karyotype from 10 metaphases, performed without knowledge of the single cell sequencing and from a separate sample of the cell line, as described[10] but without any confirmation. The karyotype can be regarded as pseudo-tetraploid with losses. It shows clear signs of endoreduplication, several abnormal chromosomes being present in two copies. There is evidence of Dutrillaux's monosomic pattern of karyotype evolution with endoreduplication - e.g. 2 copies each of  chromosomes 1, 3 and an unbalanced 1;3 translocation; 2 each of 7, 15 and a probable 7;15 translocation. Consensus karyotype: 69 chromosomes (mode 68) 2 x 1, 2 x 3, 3 x 2, der(?)t(1;3)(q?;p22-24?) x 2, 4 x 2, iso(4), 5 x 2, 6 x 2, del(6) x 2, 7 x 2, der(7)t(7;15?*) x 2, 8 x 3, 9 x 2, del(9) x 2, 10 x 2, 11 x 3**, 12 x 3, 13 x 3, 14 x 3, 15 x 2, 16 x 2, del(16), 17 x 3, 18 x 2, 19 x 3, 20 x 4, 21 x 3, 22 x 4, X x 2.

* the fragment is identified by SKY as chromosome 15 but the fragment is very small so this identification is unreliable.

** In 6/10 metaphases there are 3 elevens. In four of these there is an additional unbalanced 1;18 translocation, der(?)t(1p?;18q?), while in the other 4 metaphases one eleven is replaced by a der(?)t(11;18). A possible explanation is that the consensus should include the 1;18 translocation and three elevens, but in some metaphases one eleven and the 1;18 have combined to yield the single der(?)t(11;18).

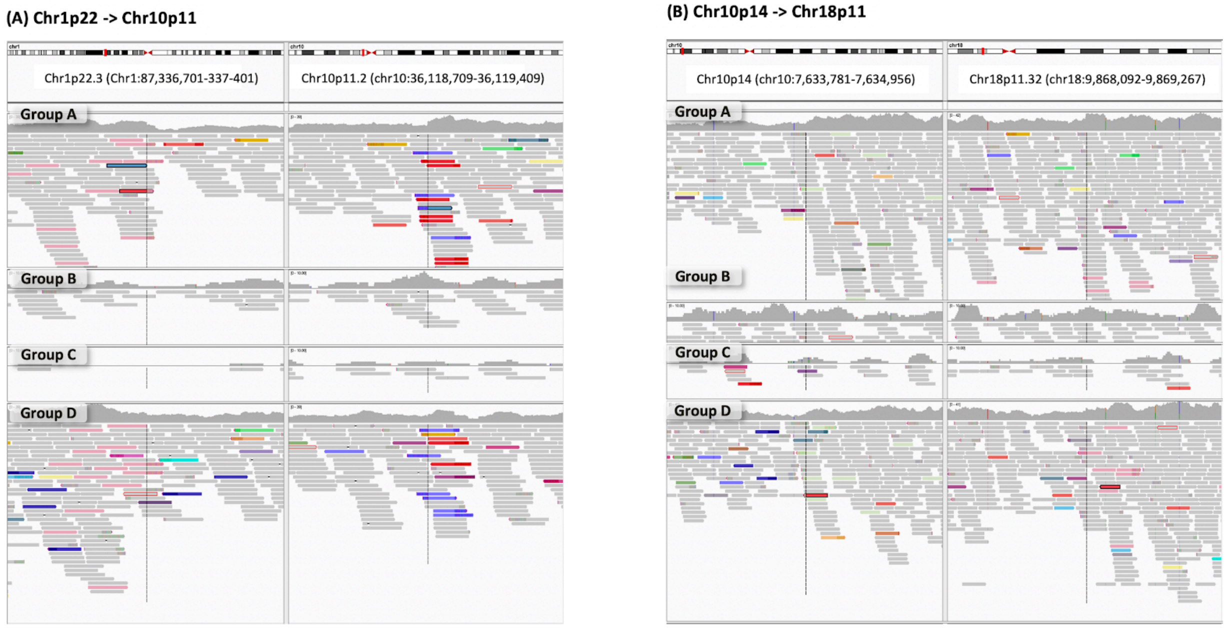


**Supplementary Figure 10 | IGV Traces of anomalous reads spanning translocations.** In left is a split view of Chr1p22 and Chr10p11, showing on the left reads in pink whose parter maps to chr10p11 shown in blue (red reads map 2.7Mb upstream). In the right is a split view of chr10p14 and Chr18p11 where on the partner of light green reads on chromosome 10p11(left) map to Chr18p11.32 (pink reads). Read coloring is formally defined within the IGV manual, and are set to indicate instances where the insert distances between paired reads is beyond 1000bp. Reads mapping within the same chromosome are red, reads mapping to chromosome 18 are in pink, reads mapping to chromosome 10 are blue, reads mapping to chromosome 1 are light green.

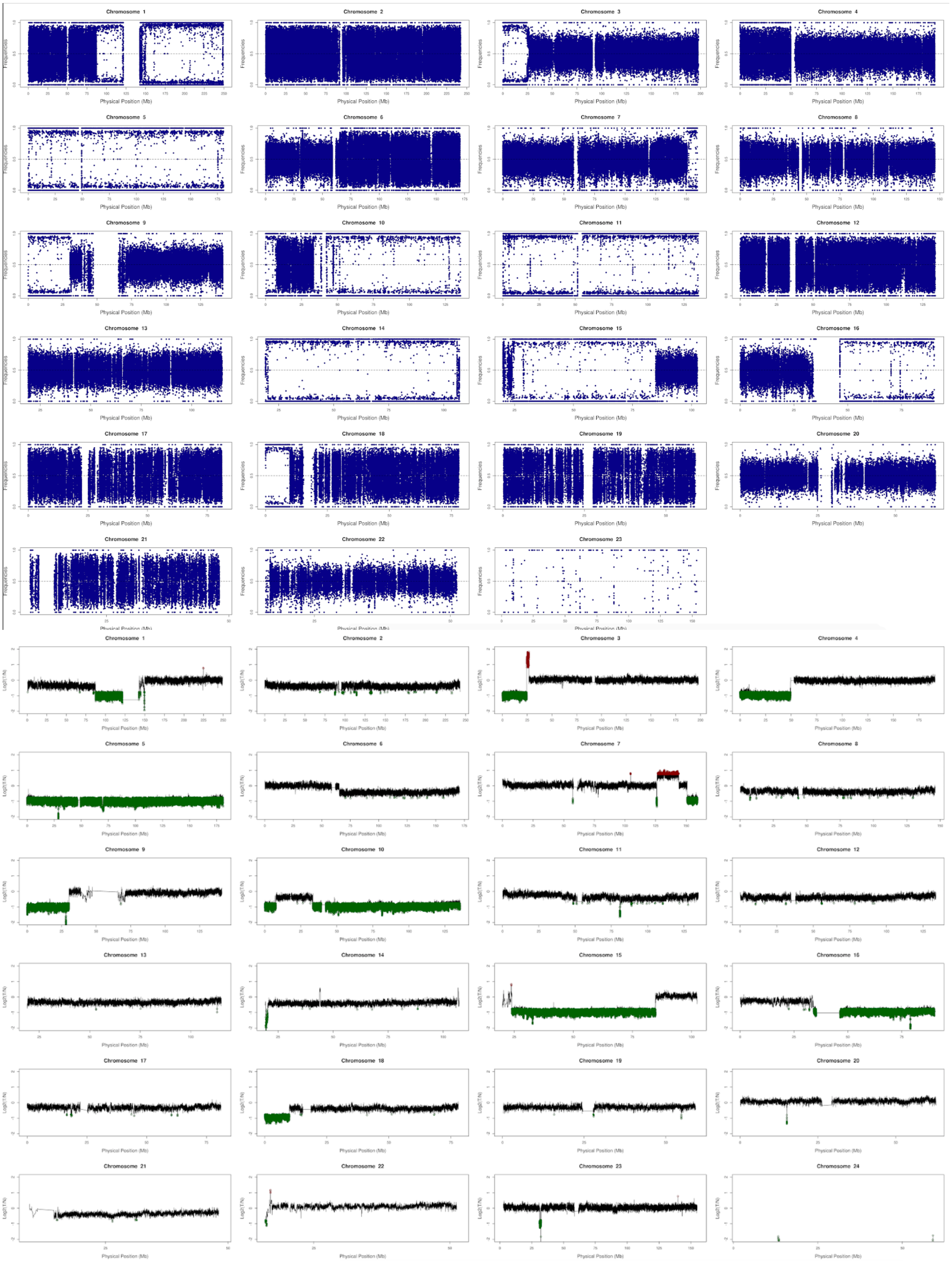


**Supplementary Figure 11 |** Exemplary Loss of Heterozygosity (LOH) broken out by chromosome for Group A (upper) and copy number estimate shown by chromosome.

**Supplementary Table 1 | Aggregate summary metrics**. Summary metrics describing sequencing data quality by group.

| Groups | Number of cells | Number of reads | Number of effective reads | Average ploidy | Median reads per Mb |
|---|---|---|---|---|---|
| **A** | 653 | 1,106,320,742 | 866,039,383 | 3.03 | 428 |
| **B** | 117 | 195,118,990 | 152,584,425 | 2.95 | 416 |
| **C** | 43 | 78,145,878 | 60,875,291 | 3.01 | 450 |
| **D** | 560 | 975,448,264 | 763,375,990 | 3.11 | 435 |

**Supplementary Table 2 | Per cell summary metrics.** Per cell metrics describing sequencing and CNV estimation data quality.

| barcode | total_num_reads | num_unmapped_reads | num_duplicate_reads | num_lowmapq_reads | num_mapped_dedup_reads | effective_reads_per_1Mbp | effective_depth_of_coverage | breadth_of_coverage_percent | mean_ploidy | raw_mapd | normalized_mapd | is_noisy |
|---|---|---|---|---|---|---|---|---|---|---|---|---|
| AAACCTGAGGCTACGA-1 | 1474382 | 14559 | 162572 | 129544 | 1167707 | 377 | 0.034061409 | 3.583452633 | 3.024427318 | 0.097366283 | 0.095937271 | 0 |
| AAACCTGGTCCTTGGG-1 | 2008984 | 34666 | 217429 | 183078 | 1573811 | 508 | 0.045954133 | 4.790426567 | 3.115261147 | 0.085370623 | 0.085370623 | 0 |
| AAACCTGTCACACGGC-1 | 1317000 | 74665 | 138878 | 112449 | 991008 | 320 | 0.028780153 | 3.0426807 | 3.087988182 | 0.105997248 | 0.10452012 | 0 |
| AAACCTGTCGATACAC-1 | 2317026 | 39831 | 253774 | 206942 | 1816479 | 587 | 0.052921028 | 5.471836533 | 3.141717404 | 0.084710744 | 0.081279279 | 0 |
| AAACGGGAGCAGATCG-1 | 1339444 | 63506 | 140596 | 118291 | 1017051 | 329 | 0.029609209 | 3.125955133 | 3.007868648 | 0.101492014 | 0.100216564 | 0 |
| AAACGGGCAAAGTAAC-1 | 1477800 | 26950 | 159618 | 138195 | 1153037 | 372 | 0.033729644 | 3.5298458 | 2.995454545 | 0.100199795 | 0.101997012 | 0 |
| AAACGGGCAACCTCAA-1 | 1595138 | 22541 | 172915 | 143540 | 1256142 | 406 | 0.036628118 | 3.840771333 | 3.093311835 | 0.094287391 | 0.094494211 | 0 |
| AAACGGGGTCTTCGTC-1 | 1421664 | 29372 | 155391 | 126737 | 1110164 | 359 | 0.032357084 | 3.406371133 | 3.097976261 | 0.099142644 | 0.100191291 | 0 |
| AAACGGGTCACGCTCT-1 | 1947700 | 69453 | 203082 | 170985 | 1504180 | 486 | 0.043601468 | 4.5173273 | 3.110392748 | 0.095612311 | 0.095115229 | 0 |
| AAACGGGTCAGCGGCT-1 | 1874782 | 77070 | 194127 | 167465 | 1436120 | 464 | 0.041812765 | 4.379685167 | 3.104920385 | 0.088254406 | 0.088273856 | 0 |
| AAAGATGAGAATTGTG-1 | 1360902 | 18396 | 149620 | 110941 | 1081945 | 350 | 0.031547124 | 3.319917867 | 3.019837493 | 0.102167465 | 0.098182972 | 0 |
| AAAGATGAGCCACCTG-1 | 1401994 | 37826 | 151902 | 114344 | 1097922 | 355 | 0.031966123 | 3.362255867 | 3.295852273 | 0.126780578 | 0.124095547 | 1 |
| AAAGCAAGTGACACGA-1 | 1627784 | 21922 | 177338 | 141815 | 1286709 | 416 | 0.037520406 | 3.930806 | 2.99614311 | 0.093248267 | 0.091966727 | 0 |
| AAAGCAATCGCGGACT-1 | 2535646 | 66346 | 270589 | 221324 | 1977387 | 639 | 0.057652785 | 5.9424639 | 2.924726828 | 0.081466752 | 0.080154264 | 0 |
| AAAGCAATCTCACCTG-1 | 1362442 | 41465 | 145952 | 118863 | 1056162 | 341 | 0.030766899 | 3.2326992 | 3.016306123 | 0.097433428 | 0.097433428 | 0 |
| AAAGTAGTCTCAGCGG-1 | 1588694 | 72155 | 164063 | 139314 | 1213162 | 392 | 0.03534518 | 3.717024933 | 3.071467129 | 0.096109253 | 0.094734635 | 0 |
| AAAGTAGTCTTGCGAA-1 | 1743832 | 21599 | 190415 | 151535 | 1380283 | 446 | 0.040302052 | 4.212429 | 3.109763721 | 0.093638707 | 0.092047768 | 0 |
| AAATGCCAGTGGAGAA-1 | 1887248 | 24398 | 206852 | 159160 | 1496838 | 484 | 0.043653634 | 4.5483168 | 3.217591145 | 0.091792681 | 0.087716553 | 0 |
| AAATGCCGTCGCGCTA-1 | 1270338 | 34801 | 127771 | 116846 | 990920 | 320 | 0.0288885 | 3.060221167 | 3.096571218 | 0.102880182 | 0.105037849 | 0 |
| AACACGTAGGACAGAA-1 | 1715456 | 26131 | 187931 | 141367 | 1360027 | 439 | 0.039683552 | 4.1453637 | 3.039962489 | 0.090117962 | 0.088450146 | 0 |
| AACCATGAGGAATCGC-1 | 1755616 | 29575 | 184714 | 160968 | 1380359 | 446 | 0.040288294 | 4.222308 | 2.9406127 | 0.090392407 | 0.092221475 | 0 |
| AACCATGAGGCTATCT-1 | 2232400 | 56542 | 240451 | 200646 | 1734761 | 560 | 0.05028511 | 5.1617407 | 3.022067702 | 0.090892961 | 0.090312027 | 1 |
| AACCATGCATTCATCT-1 | 2499372 | 69982 | 268612 | 222164 | 1938614 | 626 | 0.056290364 | 5.774783533 | 3.144110356 | 0.083626078 | 0.084799031 | 0 |
| AACCATGTCGACGAAG-1 | 1875222 | 33941 | 202787 | 159999 | 1478495 | 478 | 0.043146902 | 4.502720333 | 3.072046662 | 0.087085148 | 0.088366009 | 0 |
| AACCGCGGTAACAGCG-1 | 1166466 | 48495 | 120434 | 102263 | 895274 | 289 | 0.026036977 | 2.760984133 | 3.05008627 | 0.111992158 | 0.108989662 | 0 |
| AACCGCGGTTTACTTC-1 | 1988752 | 62880 | 205591 | 184180 | 1536101 | 496 | 0.044735569 | 4.655430567 | 3.047010005 | 0.088926653 | 0.088679451 | 0 |
| AACCGCGTCCTACGAA-1 | 563392 | 7733 | 53940 | 48652 | 453067 | 146 | 0.0131859 | 1.413549933 | 3.029506986 | 0.154139539 | 0.147379952 | 0 |
| AACGTTGGTGGTAACG-1 | 1827248 | 62088 | 191353 | 157117 | 1416690 | 458 | 0.041250358 | 4.3122855 | 3.039901684 | 0.09630223 | 0.093155669 | 0 |
| AACTCAGAGTGGGTAC-1 | 2821904 | 73049 | 301236 | 250316 | 2197303 | 710 | 0.063968984 | 6.547577067 | 2.900968998 | 0.078979288 | 0.078016643 | 0 |
| AACTCAGAGTGGTAGC-1 | 2247774 | 30306 | 243754 | 206857 | 1766857 | 571 | 0.051485642 | 5.318281967 | 3.037099937 | 0.082656861 | 0.08161954 | 0 |
| AACTCAGCATGCAGTT-1 | 1867088 | 14187 | 208911 | 153233 | 1490757 | 482 | 0.043465605 | 4.5154008 | 3.11380768 | 0.088638568 | 0.087189797 | 0 |
| AACTCAGTCGAGTATC-1 | 3006250 | 21408 | 332188 | 260471 | 2392183 | 773 | 0.069816244 | 7.090951033 | 2.924756626 | 0.076415497 | 0.075939943 | 0 |
| AACTCAGTCTAACACG-1 | 1403818 | 90210 | 145694 | 121478 | 1046436 | 338 | 0.030443965 | 3.2138271 | 3.095987898 | 0.1051481 | 0.102782668 | 0 |
| AACTCCCTCAGTTCTT-1 | 2126942 | 49183 | 229009 | 191197 | 1657553 | 535 | 0.048225871 | 5.007724567 | 3.134622898 | 0.088129786 | 0.087240166 | 0 |
| AACTCCCTCCACGATA-1 | 1534614 | 15986 | 170798 | 126146 | 1221684 | 395 | 0.035660949 | 3.7311491 | 3.026584373 | 0.095971137 | 0.095608002 | 0 |
| AACTCCCTCGATAGGG-1 | 1475146 | 15241 | 162822 | 121556 | 1175527 | 380 | 0.034315791 | 3.600711433 | 2.985248265 | 0.097488988 | 0.096216596 | 0 |
| AACTCCCTCTAATGGC-1 | 2693360 | 65471 | 290321 | 232267 | 2105301 | 680 | 0.061300555 | 6.2800128 | 3.167214862 | 0.079541131 | 0.079541131 | 0 |
| AACTCTTAGAGTAGCC-1 | 2286696 | 49254 | 244128 | 204061 | 1789253 | 578 | 0.052149528 | 5.4014467 | 3.011585535 | 0.088088396 | 0.088574925 | 0 |
| AACTCTTGTCTTACCC-1 | 1792512 | 54661 | 191208 | 159484 | 1387159 | 448 | 0.040445744 | 4.219485567 | 3.046055576 | 0.091453925 | 0.089293918 | 0 |
| AACTCTTTCCTGCGTT-1 | 1948040 | 19303 | 214334 | 176285 | 1538118 | 497 | 0.044688642 | 4.625854433 | 3.105571181 | 0.086000033 | 0.086000033 | 1 |
| AACTGGTAGAAACCAT-1 | 6448042 | 48827 | 710064 | 589019 | 5100132 | 1648 | 0.148897051 | 14.4383907 | 3.081798283 | 0.050573647 | 0.050573647 | 0 |
| AACTGGTAGCTGAACG-1 | 1725966 | 92510 | 179330 | 150747 | 1303379 | 421 | 0.037917929 | 3.983259233 | 3.143699347 | 0.093700469 | 0.093335474 | 0 |
| AACTGGTAGGGCCATA-1 | 1855552 | 20621 | 201050 | 158042 | 1475839 | 477 | 0.043092167 | 4.488310433 | 3.035765023 | 0.089645377 | 0.089243381 | 0 |
| AACTGGTGTCGTGAAG-1 | 1255874 | 27883 | 123899 | 104209 | 999883 | 323 | 0.029186444 | 3.076570033 | 3.086339776 | 0.102153256 | 0.102153256 | 0 |
| AACTGGTGTTCAGTAC-1 | 2249426 | 50915 | 244846 | 197894 | 1755771 | 567 | 0.050985723 | 5.247257533 | 3.006791654 | 0.080135328 | 0.080135328 | 0 |
| AACTGGTTCGTACGCG-1 | 980516 | 138696 | 90446 | 80860 | 670514 | 217 | 0.019437477 | 2.075725667 | 3.054446165 | 0.120807581 | 0.124841242 | 0 |
| AACTTTCCACGTAGGA-1 | 1367584 | 18896 | 151288 | 112795 | 1084605 | 350 | 0.031604757 | 3.325611067 | 3.057852531 | 0.101656719 | 0.099247645 | 0 |
| AACTTTCTCTGCTGAA-1 | 1688648 | 22641 | 186709 | 151057 | 1328241 | 429 | 0.038749421 | 4.0454271 | 3.091688631 | 0.090553483 | 0.088523146 | 0 |
| AAGACCTCACAAGCGA-1 | 774862 | 40377 | 79832 | 71791 | 582862 | 188 | 0.017061097 | 1.828308833 | 3.010372811 | 0.139719946 | 0.13524537 | 0 |
| AAGACCTCACCAGTGC-1 | 1547280 | 122948 | 155960 | 131134 | 1137238 | 367 | 0.032934267 | 3.4613962 | 3.072078406 | 0.098307929 | 0.097219793 | 0 |
| AAGACCTCACTCCCTA-1 | 2081424 | 27512 | 225963 | 175333 | 1652616 | 534 | 0.048210729 | 4.991967 | 3.308575897 | 0.121725674 | 0.119637301 | 1 |
| AAGACCTCACTTGAGT-1 | 1968878 | 19080 | 216679 | 166042 | 1567077 | 506 | 0.045742909 | 4.751080033 | 3.08331972 | 0.086776176 | 0.085854819 | 0 |
| AAGACCTCATAGCACT-1 | 2356728 | 60362 | 251951 | 210185 | 1834230 | 593 | 0.053478813 | 5.5286958 | 3.072519355 | 0.084303431 | 0.083832883 | 0 |
| AAGACCTGTGGACGTA-1 | 1273004 | 15331 | 139501 | 106601 | 1011571 | 327 | 0.029516765 | 3.1151629 | 3.140806659 | 0.105485391 | 0.102580799 | 0 |
| AAGACCTTCACGGTAT-1 | 2233960 | 17432 | 248183 | 181187 | 1787158 | 577 | 0.05212545 | 5.3641339 | 3.068237517 | 0.085376316 | 0.080627504 | 0 |
| AAGACCTTCCGAGATT-1 | 1166124 | 26337 | 127734 | 96854 | 915199 | 296 | 0.02671377 | 2.816703467 | 3.018662254 | 0.109037378 | 0.105013852 | 0 |
| AAGCCGCCATCTCACC-1 | 2708982 | 74197 | 285899 | 237985 | 2110901 | 682 | 0.061513954 | 6.314070333 | 3.043628071 | 0.080022708 | 0.079190761 | 0 |
| AAGGAGCCAAGAATCA-1 | 1116436 | 41741 | 116288 | 100776 | 857631 | 277 | 0.025076927 | 2.661535433 | 3.018511549 | 0.111501177 | 0.113028341 | 0 |
| AAGGCAGAGAGCCCAA-1 | 2335452 | 14835 | 255573 | 210527 | 1854517 | 599 | 0.054147331 | 5.599737033 | 3.072701689 | 0.080106992 | 0.081562932 | 0 |
| AAGGCAGGTAAAGTCA-1 | 3459226 | 194178 | 350672 | 310763 | 2603613 | 841 | 0.075704884 | 7.6920089 | 3.047464443 | 0.068770368 | 0.067686204 | 0 |
| AAGGTTCAGCGTGTGA-1 | 1431840 | 19756 | 150647 | 121448 | 1139989 | 368 | 0.033251916 | 3.492838367 | 2.909760066 | 0.102286839 | 0.095630929 | 0 |
| AAGGTTCCAGATTGTC-1 | 2080386 | 47393 | 225218 | 179731 | 1628044 | 526 | 0.047436772 | 4.9170601 | 3.014072401 | 0.086751793 | 0.085054525 | 0 |
| AAGGTTCGTCCGAATT-1 | 1763662 | 42806 | 187132 | 148263 | 1385461 | 448 | 0.040337872 | 4.2054647 | 2.978691214 | 0.093311446 | 0.092194392 | 0 |
| AAGGTTCGTGTAATGA-1 | 874982 | 20862 | 91887 | 73200 | 689033 | 223 | 0.020055948 | 2.133062767 | 3.084092571 | 0.123041112 | 0.122001727 | 0 |
| AAGGTTCGTTGAGCAG-1 | 1748494 | 58248 | 181067 | 153257 | 1355922 | 438 | 0.039574736 | 4.143653467 | 3.034990164 | 0.091991493 | 0.090799195 | 0 |
| AAGGTTCGTTTGCCTC-1 | 2575348 | 38730 | 278037 | 214739 | 2043842 | 660 | 0.05953717 | 6.102514167 | 3.552102078 | 0.132374251 | 0.132374251 | 1 |
| AAGGTTCTCCGCGACA-1 | 2134632 | 69346 | 223136 | 187515 | 1654635 | 534 | 0.048231319 | 5.024670833 | 2.952552774 | 0.086350371 | 0.086350371 | 0 |
| AAGTCTGAGGCCCGTT-1 | 1663256 | 18066 | 181469 | 140776 | 1322945 | 427 | 0.038595661 | 4.025418733 | 3.108119625 | 0.093715987 | 0.093597908 | 0 |
| AAGTCTGAGTGCGATG-1 | 1709082 | 23876 | 188048 | 133689 | 1363469 | 440 | 0.039843722 | 4.152287333 | 2.989865057 | 0.090676158 | 0.090578468 | 0 |
| AAGTCTGTCACCGCGA-1 | 1405766 | 16157 | 149493 | 131967 | 1108149 | 358 | 0.032291929 | 3.415382333 | 3.002989951 | 0.100584012 | 0.100109823 | 0 |
| AATCCAGGTTGAACAA-1 | 1472704 | 24058 | 162103 | 129085 | 1157458 | 374 | 0.033768803 | 3.5431151 | 3.048945735 | 0.093381783 | 0.093381783 | 0 |
| AATCCAGTCGACTGCG-1 | 1501682 | 101547 | 153596 | 132549 | 1113990 | 360 | 0.032409317 | 3.4175184 | 2.981799841 | 0.101928189 | 0.100844939 | 0 |
| AATCGGTAGAAGGGTA-1 | 1738968 | 20720 | 187638 | 144910 | 1385700 | 448 | 0.040410101 | 4.222183 | 3.084431219 | 0.097108865 | 0.093875085 | 0 |
| AATCGGTCAGATGAAT-1 | 2083510 | 18324 | 232455 | 169505 | 1663226 | 537 | 0.048516826 | 5.0141978 | 3.15247242 | 0.089356775 | 0.083903648 | 0 |
| AATCGGTCATGAATGA-1 | 1141724 | 21695 | 124020 | 96014 | 899995 | 291 | 0.026150378 | 2.770480167 | 3.058156794 | 0.106586341 | 0.110073637 | 0 |
| AATCGGTGTGCGCTTG-1 | 1377036 | 36062 | 145279 | 120108 | 1075587 | 347 | 0.031342054 | 3.299241667 | 3.121105178 | 0.101415096 | 0.100277759 | 0 |
| ACACCAAGTGCAACCC-1 | 1564696 | 61763 | 164251 | 138949 | 1199733 | 388 | 0.034921638 | 3.6709974 | 3.035879663 | 0.094193068 | 0.094981267 | 0 |
| ACACCCTAGCGTAATA-1 | 1308692 | 19257 | 142891 | 105386 | 1041158 | 336 | 0.030332771 | 3.193255233 | 3.100212341 | 0.105521785 | 0.103687227 | 0 |
| ACACCCTCAACGCCGT-1 | 2677530 | 23668 | 284999 | 243202 | 2125661 | 687 | 0.061982831 | 6.390309767 | 3.07863356 | 0.072123215 | 0.072123215 | 0 |
| ACACCGGAGTGTGAAT-1 | 2298220 | 21337 | 260232 | 191802 | 1824849 | 589 | 0.053210932 | 5.493141067 | 3.03444356 | 0.082292507 | 0.077533505 | 0 |
| ACACCGGTCTGTGGCG-1 | 1500758 | 61695 | 156073 | 133208 | 1149782 | 371 | 0.03348087 | 3.534954067 | 3.080574814 | 0.102033577 | 0.099738367 | 0 |
| ACACTGAAGACAGGCT-1 | 2166830 | 41758 | 237185 | 190773 | 1697114 | 548 | 0.049312887 | 5.095376333 | 3.000113666 | 0.087629544 | 0.086901296 | 0 |
| ACACTGAAGCTCTCGG-1 | 1732842 | 34867 | 184174 | 165484 | 1348317 | 436 | 0.03936718 | 4.120503 | 3.046892093 | 0.098925254 | 0.094127327 | 0 |
| ACACTGACACAGACCC-1 | 1749536 | 61962 | 185899 | 157247 | 1344428 | 434 | 0.039159413 | 4.103213933 | 3.03137695 | 0.091223368 | 0.089724053 | 0 |
| ACACTGACAGAGGCTA-1 | 2149830 | 49924 | 231901 | 195011 | 1672994 | 540 | 0.048825964 | 5.073285233 | 3.092605744 | 0.084236344 | 0.08413419 | 0 |
| ACACTGACAGTTCCGG-1 | 2435104 | 54813 | 263137 | 222177 | 1894977 | 612 | 0.055279632 | 5.718920433 | 2.980186273 | 0.077411177 | 0.076989279 | 0 |
| ACAGCCGAGTCTCGTA-1 | 1325632 | 12262 | 144257 | 111403 | 1057710 | 342 | 0.030840806 | 3.249797767 | 3.035836287 | 0.104421375 | 0.101959922 | 0 |
| ACAGCCGGTGATTGTC-1 | 2070006 | 33065 | 224963 | 172111 | 1639867 | 530 | 0.047820785 | 4.9551575 | 3.059110151 | 0.098641944 | 0.093297281 | 1 |

| | | | | | | | | | | | | |
|---|---|---|---|---|---|---|---|---|---|---|---|---|
| ACAGCCGGTTTGGCGC-1 | 1300972 | 20708 | 138945 | 120580 | 1020739 | 330 | 0.029695062 | 3.144617367 | 3.056738848 | 0.105839368 | 0.105839368 | 0 |
| ACAGCTAAGACTCGGA-1 | 2329338 | 36147 | 255300 | 186817 | 1851074 | 598 | 0.05392884 | 5.563354867 | 3.730724366 | 0.142051914 | 0.142003883 | 1 |
| ACAGCTACAAGCTACT-1 | 2054712 | 25559 | 227526 | 190268 | 1611359 | 521 | 0.046946572 | 4.8795573 | 3.072039558 | 0.084071237 | 0.085329516 | 1 |
| ACAGCTACAGAACAGC-1 | 1390540 | 25218 | 147780 | 142095 | 1075447 | 347 | 0.031320565 | 3.270889167 | 3.046268073 | 0.123598583 | 0.106551431 | 0 |
| ACAGCTATCCGAAGCC-1 | 2047718 | 67040 | 219115 | 181967 | 1579596 | 510 | 0.045983917 | 4.788213533 | 3.038096524 | 0.086802322 | 0.086345479 | 0 |
| ACATACGAGCCGAACA-1 | 1933334 | 82909 | 202149 | 172557 | 1475719 | 477 | 0.042939418 | 4.4796397 | 3.039761022 | 0.089031637 | 0.087489465 | 0 |
| ACATACGAGGTAACGC-1 | 2155174 | 38268 | 230722 | 193333 | 1692851 | 547 | 0.049287004 | 5.131931267 | 3.046721736 | 0.085187709 | 0.085468485 | 0 |
| ACATACGAGTTAGGGC-1 | 1843254 | 56872 | 195811 | 159734 | 1430837 | 462 | 0.041699229 | 4.3516068 | 3.108740278 | 0.089285868 | 0.087070242 | 0 |
| ACATACGCACTCCGAG-1 | 1852398 | 56273 | 197785 | 154926 | 1443414 | 466 | 0.042020874 | 4.3845015 | 3.037744025 | 0.092099546 | 0.087989202 | 0 |
| ACATACGGTATAAGTG-1 | 1216200 | 16186 | 131761 | 97258 | 970995 | 314 | 0.028311585 | 2.986117567 | 3.141729879 | 0.108040973 | 0.106239597 | 0 |
| ACATACGGTCTCTCGT-1 | 1947230 | 17401 | 207461 | 180122 | 1542246 | 498 | 0.045008917 | 4.6810789 | 2.964606422 | 0.090716768 | 0.085540393 | 0 |
| ACATCAGAGTGTACGG-1 | 1548242 | 30265 | 165430 | 138008 | 1214539 | 392 | 0.035438602 | 3.729571833 | 3.141483965 | 0.094647312 | 0.093557268 | 0 |
| ACATCAGAGTTAGGGC-1 | 2162160 | 68138 | 230836 | 186021 | 1677165 | 542 | 0.048843643 | 5.0707868 | 3.044915965 | 0.085819087 | 0.083514394 | 0 |
| ACATGGTTCGAGCACC-1 | 2342724 | 42177 | 250391 | 208241 | 1841915 | 595 | 0.053685334 | 5.5509165 | 3.094818583 | 0.079690932 | 0.080236603 | 0 |
| ACCAGTAAGATTCACC-1 | 1976604 | 34859 | 211875 | 189856 | 1540014 | 497 | 0.04495541 | 4.6606386 | 2.915582109 | 0.09155485 | 0.08523766 | 0 |
| ACCAGTAGTGTTACGT-1 | 1900212 | 14933 | 209610 | 164185 | 1511484 | 488 | 0.044075688 | 4.5936624 | 3.03761348 | 0.082385431 | 0.083474961 | 0 |
| ACCAGTATCGTTCGAA-1 | 1747368 | 17087 | 189798 | 157215 | 1383268 | 447 | 0.040387415 | 4.213793267 | 3.130242669 | 0.089748891 | 0.088434455 | 0 |
| ACCCACTAGTTATCGC-1 | 1888022 | 33922 | 201901 | 173865 | 1478334 | 478 | 0.043060069 | 4.489605733 | 3.047354849 | 0.090463628 | 0.087961445 | 0 |
| ACCCACTGTAGGGTGT-1 | 2538200 | 14179 | 276220 | 219811 | 2027990 | 655 | 0.0591447 | 6.0762124 | 3.098019858 | 0.079529912 | 0.078203858 | 0 |
| ACCGTAACACGTAGGA-1 | 1380940 | 13625 | 148441 | 116163 | 1102711 | 356 | 0.032154616 | 3.387403767 | 3.012291852 | 0.100655279 | 0.099934378 | 0 |
| ACCGTAACATCTCGTC-1 | 1371276 | 27029 | 145865 | 119787 | 1078595 | 348 | 0.031406954 | 3.312304133 | 3.047964476 | 0.104143741 | 0.101846825 | 0 |
| ACCTTTAAGTGGGCTA-1 | 1984054 | 31521 | 215983 | 173725 | 1562825 | 505 | 0.045653576 | 4.752964833 | 3.073153056 | 0.087346541 | 0.087971547 | 0 |
| ACGAGCCGTGCGCTTG-1 | 480602 | 16054 | 50702 | 39264 | 374582 | 121 | 0.010972491 | 1.1839196 | 3.006909039 | 0.165813839 | 0.166136113 | 0 |
| ACGAGGACACCGGCAT-1 | 2202826 | 39079 | 235824 | 198605 | 1729318 | 559 | 0.050400606 | 5.1903268 | 2.949807498 | 0.082234569 | 0.081124728 | 0 |
| ACGAGGACAGTAGGGT-1 | 1970072 | 20694 | 211295 | 168772 | 1569311 | 507 | 0.045824131 | 4.755401733 | 2.900216673 | 0.086821051 | 0.088564303 | 0 |
| ACGAGGACATCTCGAA-1 | 1589852 | 13520 | 175320 | 138171 | 1262841 | 408 | 0.036813441 | 3.8619104 | 2.963469995 | 0.097026897 | 0.090469689 | 0 |
| ACGATACAGAAGATTC-1 | 2405454 | 45256 | 258237 | 206864 | 1895097 | 612 | 0.055211206 | 5.684149267 | 3.034979292 | 0.084124204 | 0.08313122 | 0 |
| ACGATACCAGCCAAAG-1 | 1565708 | 43644 | 170207 | 130158 | 1221699 | 395 | 0.035579155 | 3.730359167 | 3.096921274 | 0.096255677 | 0.097437977 | 0 |
| ACGATACCAGTCCCTT-1 | 2182200 | 51047 | 233881 | 196106 | 1701166 | 550 | 0.049620587 | 5.136295233 | 3.120036095 | 0.083921356 | 0.083654971 | 0 |
| ACGATACTCGAGGTGA-1 | 2134112 | 23342 | 233072 | 180790 | 1696908 | 548 | 0.049549346 | 5.133871267 | 3.095033425 | 0.085013719 | 0.08237047 | 0 |
| ACGATGTCAACATGGG-1 | 1533672 | 18629 | 167922 | 128048 | 1219073 | 394 | 0.035624279 | 3.727744667 | 3.072894475 | 0.093725177 | 0.095030389 | 0 |
| ACGATGTGTCATACCA-1 | 1729378 | 53666 | 184144 | 143803 | 1347765 | 435 | 0.039166738 | 4.0891198 | 3.036356662 | 0.098969759 | 0.096214583 | 0 |
| ACGATGTTCTTCACAT-1 | 3345290 | 27813 | 369751 | 280877 | 2666849 | 861 | 0.077807053 | 7.906098167 | 3.078657514 | 0.07609091 | 0.07609091 | 0 |
| ACGCAGCGTTACGGCC-1 | 1792344 | 57475 | 185968 | 168114 | 1380787 | 446 | 0.040184418 | 4.200522933 | 2.971514243 | 0.090680299 | 0.088751421 | 0 |
| ACGCAGCTCCGCTCTA-1 | 2087834 | 29942 | 224002 | 185826 | 1648064 | 532 | 0.048021425 | 4.992167267 | 3.108778436 | 0.085242302 | 0.086010237 | 0 |
| ACGCCAGCAAGAGAGA-1 | 1496968 | 26486 | 159756 | 127188 | 1183538 | 382 | 0.034569834 | 3.6316096 | 3.066274643 | 0.099224777 | 0.096272719 | 0 |
| ACGCCAGCAAGGGTGT-1 | 2271446 | 17733 | 244522 | 201180 | 1808011 | 584 | 0.052738271 | 5.446306333 | 2.874476928 | 0.084700755 | 0.08500079 | 0 |
| ACGCCAGCACAGCTGC-1 | 1625818 | 24915 | 175227 | 146652 | 1279024 | 413 | 0.037190945 | 3.894130367 | 2.405532311 | 0.093930142 | 0.095435505 | 0 |
| ACGCCAGTCCACCCAT-1 | 2425936 | 23263 | 261979 | 213298 | 1927396 | 623 | 0.056289291 | 5.790497933 | 3.031712879 | 0.081790805 | 0.079392069 | 0 |
| ACGCCAGTCTCGGAAT-1 | 2524986 | 35640 | 270869 | 220915 | 1997562 | 645 | 0.058245061 | 5.999015767 | 3.10649454 | 0.082614331 | 0.079362381 | 0 |
| ACGCCGAAGACAAGCC-1 | 2059208 | 82015 | 213546 | 184685 | 1578962 | 510 | 0.045918662 | 4.777867533 | 3.052397528 | 0.085790207 | 0.087685137 | 0 |
| ACGCCGAAGCAGCCTC-1 | 1879398 | 33050 | 194090 | 171141 | 1481117 | 478 | 0.043161277 | 4.507858033 | 3.03377188 | 0.085986667 | 0.086625096 | 0 |
| ACGCCGACATGACCGC-1 | 1274780 | 29596 | 133322 | 115760 | 996102 | 322 | 0.028963708 | 3.064372933 | 3.031834421 | 0.104529933 | 0.103030621 | 0 |
| ACGGAGAAGCCCTAAT-1 | 1510314 | 12939 | 167795 | 118186 | 1211394 | 391 | 0.035365851 | 3.697122133 | 3.044725235 | 0.102422677 | 0.102156751 | 1 |
| ACGGAGAAGCCGAACA-1 | 1870200 | 59658 | 200760 | 165518 | 1444264 | 467 | 0.04211348 | 4.395974267 | 3.060841769 | 0.092934158 | 0.090285877 | 0 |
| ACGGAGAGTTGCAGCC-1 | 1687642 | 66288 | 176812 | 155015 | 1289527 | 417 | 0.037574239 | 3.9443703 | 3.092616925 | 0.097058217 | 0.096416639 | 0 |
| ACGGAGATCGGATCCG-1 | 1698712 | 36263 | 180595 | 153861 | 1327993 | 429 | 0.038666782 | 4.045069267 | 3.090801242 | 0.093251565 | 0.092367631 | 0 |
| ACGGCCAAGAAGGTTT-1 | 2095810 | 58802 | 225630 | 170473 | 1640905 | 530 | 0.047833024 | 4.944299767 | 3.131815501 | 0.106846154 | 0.106290561 | 1 |
| ACGGCCAAGAATGTGT-1 | 2317226 | 15830 | 251569 | 195441 | 1854386 | 599 | 0.054170794 | 5.567949033 | 3.16001307 | 0.081579968 | 0.080154444 | 0 |
| ACGGCCACAACGAAAT-1 | 1818180 | 29968 | 197937 | 153539 | 1436736 | 464 | 0.041907997 | 4.348304167 | 2.989077325 | 0.08821607 | 0.088891438 | 0 |
| ACGGCCACACACACGC-1 | 1745414 | 33669 | 187703 | 154960 | 1369082 | 442 | 0.03983508 | 4.169640667 | 3.132814775 | 0.089239635 | 0.090296218 | 0 |
| ACGGGCTAGTCTAGCT-1 | 2027816 | 67519 | 211866 | 174659 | 1573772 | 508 | 0.045693949 | 4.739875233 | 3.120302301 | 0.089259435 | 0.087321671 | 0 |
| ACGGGCTCATCGGTCG-1 | 1082650 | 153573 | 97437 | 91438 | 740202 | 239 | 0.021415896 | 2.282546467 | 3.077201602 | 0.121474159 | 0.121474159 | 0 |
| ACGGGCTCATGCAGTT-1 | 1555772 | 19283 | 166411 | 132949 | 1237129 | 400 | 0.036089276 | 3.771228433 | 3.118545609 | 0.094622538 | 0.092835435 | 0 |
| ACGGGCTGTCGTCTTC-1 | 2180516 | 59532 | 224953 | 194276 | 1701755 | 550 | 0.049553681 | 5.1406903 | 3.046652366 | 0.084406043 | 0.082046355 | 0 |
| ACGGGTCCACGATGGA-1 | 1712888 | 19805 | 185174 | 139869 | 1368040 | 442 | 0.039957114 | 4.171670333 | 3.225529406 | 0.089381461 | 0.089452451 | 0 |
| ACGGGTCTCGTCCCTA-1 | 1148600 | 9419 | 123872 | 104283 | 911026 | 294 | 0.026611324 | 2.817042633 | 3.047678216 | 0.10857965 | 0.109505651 | 0 |
| ACGTCAAAGATGTGGC-1 | 2449318 | 41004 | 265303 | 216361 | 1926650 | 622 | 0.05619078 | 5.790395767 | 3.033245011 | 0.080239317 | 0.079049282 | 0 |
| ACGTCAAAGGAGCGAG-1 | 2217700 | 51849 | 231815 | 191802 | 1742234 | 563 | 0.050832196 | 5.277068233 | 3.017137011 | 0.084226197 | 0.083394547 | 0 |
| ACGTCAACATTTGTGG-1 | 1383764 | 24743 | 151157 | 118318 | 1089546 | 352 | 0.031795891 | 3.3498854 | 3.00124984 | 0.105654742 | 0.101124001 | 0 |
| ACTATCTAGAAGGCCT-1 | 2340500 | 45750 | 244962 | 206087 | 1843701 | 596 | 0.053762996 | 5.561933233 | 3.079507183 | 0.081917808 | 0.079889741 | 0 |
| ACTATCTAGTACAGTA-1 | 1373640 | 10603 | 151092 | 109642 | 1102303 | 356 | 0.032171528 | 3.3730243 | 2.936983713 | 0.101853901 | 0.102282196 | 0 |
| ACTATCTCATGGCTAT-1 | 2381058 | 24025 | 260827 | 207782 | 1888424 | 610 | 0.055100183 | 5.672152967 | 2.989273054 | 0.081251062 | 0.079224372 | 0 |
| ACTATCTTCGGAGAAC-1 | 2561748 | 31205 | 277288 | 226690 | 2026565 | 655 | 0.059167535 | 6.0588698 | 3.388715224 | 0.093515303 | 0.094152367 | 1 |
| ACTATCTTCTCGAACA-1 | 2087154 | 61595 | 221458 | 182462 | 1621639 | 524 | 0.047239503 | 4.9084405 | 3.082060499 | 0.084946899 | 0.085283738 | 0 |
| ACTATCTTCTGGTAGT-1 | 1379718 | 5169 | 151366 | 122603 | 1100580 | 356 | 0.032233351 | 3.374320867 | 3.073016392 | 0.098855663 | 0.099042487 | 0 |
| ACTGAACAGCTAAACA-1 | 1813600 | 50069 | 196045 | 157746 | 1409740 | 455 | 0.041107915 | 4.306804667 | 3.073844427 | 0.088706972 | 0.089138143 | 0 |
| ACTGAACAGTGCGATG-1 | 1410344 | 18306 | 154109 | 116718 | 1121211 | 362 | 0.032742614 | 3.438070667 | 3.046910406 | 0.104102781 | 0.103905052 | 0 |
| ACTGAACCACGCGATC-1 | 2013756 | 61007 | 212582 | 178930 | 1561237 | 504 | 0.04551106 | 4.726455133 | 3.058686063 | 0.092290925 | 0.09067528 | 0 |
| ACTGAGTCACGTTTAG-1 | 1102454 | 16367 | 120770 | 93160 | 872157 | 282 | 0.025507936 | 2.700768867 | 3.170097016 | 0.108813419 | 0.110268539 | 0 |
| ACTGAGTTCTTCTAAC-1 | 2190698 | 22791 | 242101 | 192580 | 1733226 | 560 | 0.050609432 | 5.2344077 | 3.032377218 | 0.081053923 | 0.080287096 | 0 |
| ACTGATGCACGACTAT-1 | 2184730 | 22637 | 242816 | 187936 | 1731341 | 559 | 0.050534506 | 5.223113867 | 3.080569091 | 0.088297215 | 0.083819471 | 0 |
| ACTGATGCATACCAGT-1 | 2430222 | 13424 | 269744 | 211919 | 1935135 | 625 | 0.056503135 | 5.818503667 | 3.096903218 | 0.080059274 | 0.078694419 | 0 |
| ACTGATGGTACGTCAT-1 | 2376510 | 23895 | 257856 | 208841 | 1885918 | 609 | 0.054981658 | 5.666858133 | 3.147523979 | 0.085689203 | 0.08075692 | 0 |
| ACTGATGTCGTTCGTC-1 | 1803874 | 31510 | 189485 | 160372 | 1422507 | 460 | 0.041483939 | 4.335570633 | 3.093830581 | 0.087089762 | 0.087962696 | 0 |
| ACTGATGTCTTATGGG-1 | 1536906 | 16390 | 167930 | 127867 | 1224719 | 396 | 0.035839708 | 3.7464362 | 3.043617225 | 0.09557357 | 0.093558139 | 0 |
| ACTGCTCAGATAGCAT-1 | 1975236 | 15691 | 218189 | 171038 | 1570318 | 507 | 0.045775088 | 4.755239333 | 3.206565667 | 0.085855248 | 0.085988091 | 0 |
| ACTGCTCAGCGCCTTG-1 | 1238026 | 14333 | 133668 | 107576 | 982449 | 317 | 0.028676488 | 3.029836333 | 3.047631899 | 0.108652568 | 0.105278423 | 0 |
| ACTGCTCCAATCGTCA-1 | 1772796 | 65297 | 186642 | 159768 | 1361089 | 440 | 0.039581459 | 4.154666133 | 3.106394919 | 0.090209465 | 0.089435262 | 0 |
| ACTGCTCGTGGAAAGA-1 | 1125862 | 20746 | 118026 | 100092 | 886998 | 287 | 0.025923185 | 2.744586533 | 2.949218639 | 0.110110618 | 0.110110618 | 0 |
| ACTTACTCACATAAAG-1 | 1145880 | 15642 | 127372 | 93137 | 909729 | 294 | 0.026486209 | 2.7934199 | 3.125732302 | 0.110359012 | 0.10959121 | 0 |
| ACTTACTTCCGAGATT-1 | 1161312 | 30222 | 126547 | 96522 | 908021 | 293 | 0.026498883 | 2.796564633 | 3.047326184 | 0.111954611 | 0.109939767 | 0 |
| ACTTACTTCTCATGTT-1 | 830544 | 5440 | 92263 | 66095 | 666746 | 215 | 0.019556487 | 2.063727167 | 3.029907551 | 0.126517812 | 0.130726055 | 0 |
| ACTTGTTAGGGTTTCT-1 | 2007414 | 67129 | 208535 | 180126 | 1551624 | 501 | 0.045093868 | 4.668288967 | 3.086407739 | 0.093588969 | 0.089122439 | 0 |
| ACTTGTTAGTTGTAGA-1 | 1407786 | 12403 | 153720 | 119717 | 1121946 | 362 | 0.032773897 | 3.447640233 | 3.156688115 | 0.099948837 | 0.098970935 | 0 |
| ACTTGTTGTCTAGGTT-1 | 2230772 | 26787 | 243670 | 185047 | 1775268 | 573 | 0.051795108 | 5.359951533 | 5.411388583 | 0.079391008 | 0.079391008 | 0 |
| ACTTGTTTCTCAGATG-1 | 1354672 | 15772 | 151326 | 109253 | 1078321 | 348 | 0.031508377 | 3.308018633 | 3.046968233 | 0.102145154 | 0.099140309 | 0 |
| ACTTGTTTCTGTGGCG-1 | 1437846 | 110970 | 144459 | 124653 | 1057764 | 342 | 0.030723092 | 3.238338333 | 3.17890827 | 0.102472444 | 0.100926321 | 0 |
| ACTTTCACAAACGAGC-1 | 2284376 | 30787 | 245830 | 212764 | 1794995 | 580 | 0.052342236 | 5.398161667 | 3.106089773 | 0.083718647 | 0.082206056 | 0 |
| ACTTTCACACATTCAG-1 | 1881816 | 25671 | 207654 | 163702 | 1484789 | 480 | 0.043340149 | 4.527164267 | 3.164901363 | 0.088559292 | 0.088963935 | 0 |
| ACTTTCACAGCGGTCT-1 | 1699466 | 77603 | 176749 | 150333 | 1294781 | 418 | 0.037617491 | 3.9313497 | 3.166321023 | 0.092560531 | 0.094528253 | 0 |
| ACTTTCAGTAGCAGTG-1 | 2130118 | 25382 | 235611 | 175558 | 1693567 | 547 | 0.049247292 | 5.1126856 | 2.854217732 | 0.079162952 | 0.078942267 | 0 |
| AGAATAGAGAAAGCTT-1 | 561428 | 4795 | 58768 | 49587 | 448278 | 145 | 0.01307636 | 1.401687 | 3.135153942 | 0.167327965 | 0.1532965 | 0 |
| AGAATAGAGTCAAGGC-1 | 1274676 | 41008 | 133170 | 110554 | 989944 | 320 | 0.028964719 | 3.063083833 | 3.027646365 | 0.107340015 | 0.105737256 | 0 |

| | | | | | | | | | | | | |
|---|---|---|---|---|---|---|---|---|---|---|---|---|
| AGAATAGCAAGCAAGC-1 | 2156808 | 44481 | 227759 | 190487 | 1694081 | 547 | 0.049406282 | 5.112295633 | 2.967717643 | 0.085479864 | 0.083294191 | 0 |
| AGAATAGGTGAAGCCA-1 | 1572888 | 52876 | 163326 | 143226 | 1213460 | 392 | 0.035362654 | 3.7213061 | 3.033423077 | 0.097017399 | 0.097017399 | 0 |
| AGACGTTTCCTCAGTC-1 | 1918948 | 26735 | 210538 | 162272 | 1519403 | 491 | 0.044235042 | 4.604272233 | 3.037907958 | 0.088403319 | 0.088556612 | 0 |
| AGAGCGAAGTCCATAC-1 | 2044638 | 35223 | 218603 | 168348 | 1622464 | 524 | 0.047287512 | 4.890652167 | 3.112902653 | 0.086209627 | 0.08564284 | 0 |
| AGAGCGACAGTGTGGA-1 | 1460510 | 22996 | 149562 | 128787 | 1159165 | 374 | 0.033863924 | 3.559892733 | 2.922134379 | 0.096894229 | 0.095514297 | 0 |
| AGAGCGAGTTTGTCAG-1 | 2044704 | 49614 | 214486 | 181666 | 1598938 | 517 | 0.046655888 | 4.850100467 | 2.958417389 | 0.086154332 | 0.084403797 | 0 |
| AGAGCGATCCCACGGA-1 | 1990708 | 49217 | 203124 | 171154 | 1567213 | 506 | 0.045640518 | 4.761051433 | 3.069606854 | 0.085818954 | 0.085188129 | 0 |
| AGAGCGATCGTGCCTT-1 | 528298 | 7433 | 45153 | 45646 | 430066 | 139 | 0.012560575 | 1.347902567 | 2.942494531 | 0.160100003 | 0.160100003 | 0 |
| AGAGCTTAGACGTCGA-1 | 1901952 | 23668 | 199568 | 176547 | 1502169 | 485 | 0.043834308 | 4.575904767 | 3.019696001 | 0.080964695 | 0.081844519 | 0 |
| AGAGCTTCATTCTTCA-1 | 1950338 | 65432 | 202472 | 175183 | 1507251 | 487 | 0.043876453 | 4.571817367 | 3.016754856 | 0.088704655 | 0.088931732 | 0 |
| AGAGTGGAGGCATGGT-1 | 1550684 | 17525 | 165018 | 134619 | 1233522 | 398 | 0.035976035 | 3.7670154 | 3.109204667 | 0.096247216 | 0.096247216 | 0 |
| AGAGTGGCACTACATG-1 | 848824 | 14877 | 91631 | 68597 | 673719 | 218 | 0.019592001 | 2.075590933 | 2.926195227 | 0.12482493 | 0.127679228 | 0 |
| AGAGTGGCATGAAAGT-1 | 1634482 | 33198 | 173472 | 148342 | 1279470 | 413 | 0.037343874 | 3.904139367 | 3.066837151 | 0.093745959 | 0.095522393 | 0 |
| AGAGTGGGTACAAGGC-1 | 1574718 | 62628 | 164436 | 135532 | 1212122 | 392 | 0.035291039 | 3.704511433 | 3.144035231 | 0.093973977 | 0.095234517 | 0 |
| AGAGTGGGTTCTTCGC-1 | 2160054 | 42859 | 225552 | 197871 | 1693772 | 547 | 0.049381983 | 5.1403705 | 3.09598579 | 0.085432457 | 0.083441706 | 0 |
| AGATCTGCACCAGTGC-1 | 1302742 | 111607 | 130852 | 112584 | 947699 | 306 | 0.027392843 | 2.904960333 | 3.003018637 | 0.110801733 | 0.10833319 | 0 |
| AGATCTGTCTTCTTGA-1 | 1027810 | 17927 | 111568 | 89262 | 809053 | 261 | 0.0236172 | 2.506383433 | 3.151715517 | 0.110572552 | 0.111794884 | 0 |
| AGATTGCAGCAAAGTT-1 | 1725934 | 45722 | 184586 | 150101 | 1345525 | 435 | 0.039155553 | 4.076863833 | 3.144242657 | 0.095546842 | 0.093409757 | 0 |
| AGATTGCCAGAATTCC-1 | 1775544 | 73194 | 183743 | 160486 | 1358121 | 439 | 0.03940685 | 4.109750467 | 3.141270382 | 0.091239103 | 0.092254147 | 0 |
| AGATTGCCATGATGCT-1 | 2359312 | 52078 | 251012 | 214016 | 1842206 | 595 | 0.053201518 | 5.4074756 | 3.018030178 | 0.0875062 | 0.08479324 | 0 |
| AGATTGCGTCACCGCA-1 | 1361118 | 25537 | 148711 | 122783 | 1064087 | 344 | 0.030926321 | 3.2684244 | 3.000106522 | 0.103068232 | 0.103068232 | 0 |
| AGATTGCTCGCGAGAA-1 | 1617266 | 19635 | 172416 | 137220 | 1287995 | 416 | 0.037576083 | 3.931550167 | 3.050215921 | 0.09602684 | 0.092561879 | 0 |
| AGCAGCCAGGGTCGAT-1 | 2197878 | 29517 | 233974 | 198460 | 1735927 | 561 | 0.050672239 | 5.250421533 | 3.283035353 | 0.081620525 | 0.081620525 | 0 |
| AGCAGCCAGTGTGATA-1 | 1620070 | 16473 | 177381 | 135473 | 1290743 | 417 | 0.037708206 | 3.941775667 | 3.110052196 | 0.093259462 | 0.090649856 | 0 |
| AGCAGCCGTTGACGGA-1 | 1443074 | 28203 | 155875 | 128520 | 1130476 | 365 | 0.032949853 | 3.478139333 | 3.14314121 | 0.101264371 | 0.098761295 | 0 |
| AGCAGCCTCTCATGTT-1 | 1247746 | 13976 | 136865 | 99636 | 997269 | 322 | 0.02910808 | 3.055169933 | 3.098491249 | 0.104614137 | 0.101126923 | 0 |
| AGCATACCAGGTACGA-1 | 1474976 | 23637 | 159689 | 130397 | 1161253 | 375 | 0.033815691 | 3.560159367 | 3.020477743 | 0.097256597 | 0.096098827 | 0 |
| AGCATACCATCATGGT-1 | 1791440 | 16376 | 195968 | 152142 | 1426954 | 461 | 0.041667075 | 4.327912133 | 2.832952156 | 0.090749601 | 0.090749601 | 0 |
| AGCCTAAAGTAAGTAC-1 | 2358660 | 59689 | 252197 | 209593 | 1837181 | 593 | 0.053447476 | 5.528049633 | 3.063391937 | 0.083021023 | 0.083341018 | 0 |
| AGCCTAACACGTCATA-1 | 1577272 | 22571 | 171513 | 138100 | 1245088 | 402 | 0.036342669 | 3.811247 | 2.990751264 | 0.098630928 | 0.096221741 | 0 |
| AGCCTAAGTCTCTCGT-1 | 2205756 | 33923 | 236537 | 196392 | 1738904 | 562 | 0.050758799 | 5.262497333 | 3.013723248 | 0.081779361 | 0.082401627 | 0 |
| AGCCTAAGTGTCGCTG-1 | 1209060 | 25888 | 129084 | 110236 | 943852 | 305 | 0.027478375 | 2.903478767 | 3.040305344 | 0.103093333 | 0.104352378 | 0 |
| AGCCTAATCCTCATCG-1 | 1593474 | 58089 | 167157 | 149951 | 1218277 | 394 | 0.035471316 | 3.736384133 | 3.063203525 | 0.100146994 | 0.098470001 | 0 |
| AGCGGTCAGCGTAGTG-1 | 1023310 | 17482 | 106164 | 80445 | 819219 | 265 | 0.02388401 | 2.519530667 | 3.039068215 | 0.114023001 | 0.111648753 | 0 |
| AGCGGTCCACAAGCGA-1 | 1365968 | 44907 | 141731 | 126755 | 1052575 | 340 | 0.030660726 | 3.246291167 | 3.108231502 | 0.097617216 | 0.099952199 | 0 |
| AGCGGTCCAGGCCGTT-1 | 1148618 | 11633 | 121041 | 105371 | 910573 | 294 | 0.026533706 | 2.8174782 | 2.988449654 | 0.106565146 | 0.107953076 | 0 |
| AGCGGTCGTAACTGTC-1 | 1374064 | 26408 | 138507 | 117266 | 1091883 | 353 | 0.031798974 | 3.347293433 | 3.063481127 | 0.109992841 | 0.106060754 | 1 |
| AGCGGTCGTCATCTGA-1 | 919006 | 18041 | 93317 | 82387 | 725261 | 234 | 0.021152135 | 2.257329233 | 2.922124949 | 0.124304729 | 0.123691695 | 0 |
| AGCGGTCTCATCGCCT-1 | 1720890 | 29976 | 181616 | 158431 | 1350867 | 436 | 0.039301563 | 4.121036333 | 2.861561308 | 0.094711954 | 0.092806781 | 0 |
| AGCGTATAGGCTATCT-1 | 2271176 | 38756 | 238477 | 205649 | 1788294 | 578 | 0.05174908 | 5.2981225 | 3.129033404 | 0.084654327 | 0.085006466 | 0 |
| AGCGTATCAGTGTGGA-1 | 1763302 | 21936 | 185483 | 160013 | 1395870 | 451 | 0.040754746 | 4.247786033 | 3.099565993 | 0.092988907 | 0.088487988 | 0 |
| AGCGTCGCACTCACCT-1 | 329072 | 21274 | 24516 | 27433 | 255849 | 83 | 0.007439767 | 0.808143467 | 3.091444192 | 0.202630239 | 0.191800209 | 0 |
| AGCTCCTAGACTTCCA-1 | 1971326 | 52313 | 208430 | 181948 | 1528635 | 494 | 0.044484478 | 4.6452097 | 3.045161703 | 0.088704676 | 0.085352278 | 0 |
| AGCTCCTAGGATGCGT-1 | 2353654 | 20983 | 253451 | 211655 | 1867565 | 603 | 0.054495186 | 5.618355233 | 2.999019629 | 0.080396012 | 0.080759701 | 0 |
| AGCTCCTCAGTACCTA-1 | 1787978 | 24588 | 194188 | 155884 | 1413318 | 457 | 0.041262715 | 4.305795233 | 3.067225936 | 0.085437463 | 0.086391948 | 0 |
| AGCTCCTGTAGTAGTA-1 | 654126 | 9848 | 62407 | 52142 | 529729 | 171 | 0.01547593 | 1.6500158 | 3.065543045 | 0.138996835 | 0.138320039 | 0 |
| AGCTCCTTCGAGTTGT-1 | 2480190 | 25318 | 273164 | 219795 | 1961913 | 634 | 0.057235343 | 5.895325367 | 3.110919332 | 0.077788927 | 0.077247803 | 0 |
| AGCTCTCAGCCGGTAA-1 | 1532874 | 15948 | 168798 | 128963 | 1219165 | 394 | 0.035608948 | 3.726568233 | 2.993757014 | 0.09378936 | 0.094398821 | 0 |
| AGCTCTCAGTTAGCGG-1 | 1500694 | 45535 | 158641 | 139343 | 1157175 | 374 | 0.033725253 | 3.550303967 | 3.056084258 | 0.102155779 | 0.098523681 | 0 |
| AGCTCTCGTACAACGG-1 | 1331684 | 33482 | 141517 | 117259 | 1039426 | 336 | 0.030322955 | 3.204334967 | 3.117052772 | 0.10090187 | 0.101664733 | 0 |
| AGCTCTCGTACTCCGG-1 | 1850330 | 51131 | 194030 | 166326 | 1438843 | 465 | 0.041962515 | 4.3804916 | 3.036522023 | 0.089909879 | 0.089106476 | 0 |
| AGCTCTCGTGATTACC-1 | 2812576 | 58022 | 295117 | 254261 | 2205176 | 712 | 0.063958728 | 6.489611833 | 3.075038361 | 0.081157984 | 0.079902206 | 0 |
| AGCTCTCTCGAGTTGT-1 | 3359056 | 31623 | 367746 | 277568 | 2682119 | 866 | 0.078285805 | 7.913010467 | 4.325234842 | 0.125119579 | 0.125119579 | 1 |
| AGCTTGAAGGCCCGTT-1 | 1720304 | 23476 | 186376 | 147660 | 1362792 | 440 | 0.039825946 | 4.1376289 | 3.016888605 | 0.095196976 | 0.089450383 | 0 |
| AGCTTGAAGTCGCGTC-1 | 1053408 | 52122 | 98986 | 88481 | 813819 | 263 | 0.023684823 | 2.5099128 | 4.699638435 | 0.152915307 | 0.152915307 | 1 |
| AGCTTGACACATCGAT-1 | 1968272 | 23166 | 216453 | 172744 | 1555909 | 503 | 0.045311812 | 4.6991641 | 2.914184009 | 0.09464809 | 0.092702729 | 1 |
| AGCTTGAGTCAACTGT-1 | 2299630 | 43004 | 250500 | 200548 | 1805578 | 583 | 0.052700057 | 5.4480065 | 3.108053195 | 0.0833082 | 0.082573039 | 0 |
| AGGCCACAGATCTGAA-1 | 1749286 | 27152 | 185499 | 151857 | 1384778 | 447 | 0.040337724 | 4.213841667 | 3.07027338 | 0.087132657 | 0.086964727 | 0 |
| AGGCCACAGTTTGCGT-1 | 2204010 | 30928 | 237658 | 197792 | 1737632 | 561 | 0.050733872 | 5.249468267 | 3.048412873 | 0.083782048 | 0.080772231 | 0 |
| AGGCCACGTTGGCGTC-1 | 1423172 | 37314 | 150379 | 125953 | 1109526 | 358 | 0.032314932 | 3.406177833 | 3.000369321 | 0.096717111 | 0.098356915 | 0 |
| AGGCCACTCAAGAGGC-1 | 1892180 | 38572 | 202662 | 163886 | 1487060 | 480 | 0.043325615 | 4.517488 | 2.966464216 | 0.090024812 | 0.08962511 | 0 |
| AGGCCACTCCACCGGA-1 | 1603934 | 32574 | 170544 | 144874 | 1255942 | 406 | 0.036584129 | 3.844998967 | 3.140054411 | 0.090084149 | 0.091609034 | 0 |
| AGGCCACTCTAGCTAG-1 | 409572 | 3201 | 43476 | 33921 | 328974 | 106 | 0.009612159 | 1.038991633 | 3.038397625 | 0.181373079 | 0.175496421 | 0 |
| AGGCCGTAGCACAGGT-1 | 1583174 | 15312 | 173080 | 133501 | 1261281 | 407 | 0.036874958 | 3.866255933 | 3.143004247 | 0.094611003 | 0.093961567 | 0 |
| AGGCCGTCATAGTGAA-1 | 1649578 | 19734 | 178405 | 136028 | 1315411 | 425 | 0.038364494 | 4.0086198 | 3.023141341 | 0.09412431 | 0.094374613 | 0 |
| AGGCCGTGTCTAGGTT-1 | 417406 | 4283 | 44720 | 32526 | 335877 | 108 | 0.009827153 | 1.0562403 | 3.034644328 | 0.182642423 | 0.182642423 | 0 |
| AGGCCGTGTCTCTATT-1 | 750368 | 22606 | 79750 | 59082 | 588930 | 190 | 0.017158877 | 1.8215811 | 3.034321417 | 0.128367768 | 0.132775678 | 0 |
| AGGCCGTGTGTGTATC-1 | 1148566 | 27913 | 119625 | 100513 | 900515 | 291 | 0.026291303 | 2.780606433 | 2.999424769 | 0.114266837 | 0.110833141 | 0 |
| AGGCCGTTCGTTAGGT-1 | 1801584 | 17503 | 193153 | 150139 | 1440789 | 465 | 0.042106187 | 4.387917567 | 3.024531947 | 0.091926877 | 0.091042759 | 0 |
| AGGCCGTTCTGCACAA-1 | 1537946 | 13667 | 167960 | 129639 | 1226680 | 396 | 0.035798326 | 3.746041633 | 2.999239981 | 0.09784165 | 0.096891759 | 0 |
| AGGGAGTTCGCTATGA-1 | 876860 | 12731 | 93741 | 73801 | 696587 | 225 | 0.020275435 | 2.152470567 | 3.099620069 | 0.118109483 | 0.119282838 | 0 |
| AGGGATGAGGCATTGG-1 | 1663006 | 31429 | 180490 | 138202 | 1312885 | 424 | 0.038318826 | 4.014689167 | 3.035017576 | 0.095834927 | 0.092147237 | 0 |
| AGGGATGCAAATCGCT-1 | 1401128 | 40333 | 150137 | 129017 | 1081641 | 349 | 0.031482126 | 3.324889367 | 3.169809847 | 0.099356491 | 0.101228004 | 0 |
| AGGGATGGTATAAGTG-1 | 1149906 | 19981 | 124271 | 90984 | 914670 | 295 | 0.026666984 | 2.814235133 | 2.972223011 | 0.107295848 | 0.10589313 | 0 |
| AGGGATGTCGTACGCG-1 | 893812 | 133266 | 82388 | 68572 | 609586 | 197 | 0.01774024 | 1.9008593 | 3.03236115 | 0.134096269 | 0.129364559 | 0 |
| AGGGATGTCTGGCCTT-1 | 1978200 | 52721 | 209735 | 175412 | 1540332 | 498 | 0.044828087 | 4.663919133 | 2.966789563 | 0.089551548 | 0.087588747 | 0 |
| AGGGTGAGTCCTACCT-1 | 1846048 | 21366 | 196755 | 167070 | 1460857 | 472 | 0.042498688 | 4.433703067 | 3.065491792 | 0.089080199 | 0.090014913 | 0 |
| AGGGTGAGTGACTGAG-1 | 1702810 | 28855 | 181835 | 143145 | 1348975 | 436 | 0.039390755 | 4.113775267 | 3.015918228 | 0.090255251 | 0.089502727 | 0 |
| AGGTCATAGCCATGCC-1 | 861790 | 4740 | 92524 | 76985 | 687541 | 222 | 0.020067168 | 2.141876333 | 3.130373643 | 0.128597655 | 0.124515747 | 0 |
| AGGTCATAGGCATTTC-1 | 2690112 | 90212 | 283915 | 239171 | 2076814 | 671 | 0.059996559 | 6.084890167 | 3.151966077 | 0.086005509 | 0.085088698 | 1 |
| AGGTCATAGTGATTGA-1 | 1689778 | 22184 | 183484 | 141027 | 1343083 | 434 | 0.039168435 | 4.0873525 | 3.010681591 | 0.093728168 | 0.092958044 | 0 |
| AGGTCATCAACATAAG-1 | 1174394 | 15627 | 129094 | 91346 | 938327 | 303 | 0.027418795 | 2.881191933 | 3.046460931 | 0.111822744 | 0.109433147 | 0 |
| AGGTCATTCGACAATC-1 | 2533004 | 37228 | 275819 | 226325 | 1993632 | 644 | 0.057993473 | 5.945801967 | 3.047537513 | 0.07929588 | 0.078320134 | 0 |
| AGGTCATTCGCGGCAT-1 | 2614406 | 47636 | 277343 | 223643 | 2065784 | 667 | 0.060215849 | 6.1721901 | 3.080780954 | 0.078036424 | 0.077922745 | 0 |
| AGGTCCGCAGTGTGGA-1 | 1863294 | 24403 | 202809 | 158068 | 1478014 | 477 | 0.043176622 | 4.501095 | 2.995651618 | 0.091251477 | 0.089331377 | 0 |
| AGGTCCGGTTGCCTCT-1 | 3346364 | 40672 | 357020 | 280321 | 2668351 | 862 | 0.077779102 | 7.9095452 | 2.842413903 | 0.098164342 | 0.098164342 | 1 |
| AGGTCCGTCTCGAGCG-1 | 1075000 | 94614 | 103951 | 94696 | 781739 | 253 | 0.022783577 | 2.428875767 | 3.061461004 | 0.116626259 | 0.114173925 | 0 |
| AGTAGTCCAATCGCGC-1 | 1347544 | 39366 | 138889 | 123782 | 1045507 | 338 | 0.030438488 | 3.2221532 | 3.070319713 | 0.105434103 | 0.10316717 | 0 |
| AGTCTTTAGTCTAGCT-1 | 1757466 | 24188 | 191071 | 160807 | 1381400 | 446 | 0.040149387 | 4.175805133 | 2.918222689 | 0.091972992 | 0.091972992 | 0 |
| AGTCTTTGTTCCACAA-1 | 1696910 | 14501 | 188090 | 144795 | 1349524 | 436 | 0.039333171 | 4.114171967 | 3.109465249 | 0.092673814 | 0.091358931 | 0 |
| AGTCTTTTCCGTCTAC-1 | 2442606 | 40672 | 265703 | 221502 | 1914729 | 619 | 0.055856444 | 5.7734663 | 3.106367861 | 0.081495182 | 0.080163436 | 0 |
| AGTGAGGAGGCATGGT-1 | 1666548 | 18462 | 178529 | 148797 | 1320760 | 427 | 0.038545755 | 4.034064233 | 2.973534448 | 0.091553745 | 0.091891211 | 0 |
| AGTGAGGCATTCCGAA-1 | 1648850 | 23461 | 178964 | 137282 | 1309143 | 423 | 0.038175487 | 3.987958 | 2.938579791 | 0.090520298 | 0.093213992 | 0 |
| AGTGAGGTCTGTATGG-1 | 1237886 | 21509 | 133156 | 99947 | 983274 | 318 | 0.028707147 | 3.022858633 | 2.985521756 | 0.107002386 | 0.104174017 | 0 |

| | | | | | | | | | | | | |
|---|---|---|---|---|---|---|---|---|---|---|---|---|
| AGTGGGAAGCTGAAAT-1 | 1490226 | 36794 | 157588 | 132316 | 1163528 | 376 | 0.033895017 | 3.557828 | 3.108770235 | 0.098696662 | 0.097026677 | 0 |
| AGTGGGAGTCACCAGC-1 | 1978564 | 42869 | 208403 | 180238 | 1547054 | 500 | 0.045106961 | 4.709262133 | 3.047462059 | 0.087613557 | 0.089622417 | 0 |
| AGTGGGATCTGACGCG-1 | 592084 | 177657 | 44217 | 38054 | 332156 | 107 | 0.009612948 | 1.042086633 | 3.121591014 | 0.161938344 | 0.169504918 | 0 |
| AGTGGGATCTGGGACC-1 | 1927970 | 53434 | 200149 | 174713 | 1499674 | 484 | 0.043682922 | 4.5523684 | 3.140047447 | 0.090020696 | 0.088305152 | 0 |
| AGTGTCACATGTAGCT-1 | 1892756 | 57650 | 197711 | 172374 | 1465021 | 473 | 0.042408426 | 4.405439567 | 3.118092166 | 0.093100892 | 0.091984156 | 0 |
| AGTGTCAGTACGTCAT-1 | 2299514 | 22851 | 245912 | 203154 | 1827597 | 590 | 0.053315924 | 5.4916656 | 3.174907131 | 0.08045447 | 0.080076041 | 0 |
| AGTGTCATCGAGCGTC-1 | 459060 | 45085 | 36731 | 36910 | 340334 | 110 | 0.009920517 | 1.069982567 | 3.075239806 | 0.169163897 | 0.167422671 | 0 |
| AGTTGGTCAAACGAGC-1 | 2297322 | 47934 | 242853 | 201074 | 1805461 | 583 | 0.052653154 | 5.448442233 | 3.065757591 | 0.081712516 | 0.081095118 | 0 |
| AGTTGGTCACTTCAGA-1 | 1380784 | 19083 | 148848 | 123652 | 1089201 | 352 | 0.031800664 | 3.348369933 | 3.119108321 | 0.100975167 | 0.09953487 | 0 |
| AGTTGGTGTGTTCGAT-1 | 2197724 | 28037 | 238073 | 185774 | 1745840 | 564 | 0.050898742 | 5.2592854 | 3.114321037 | 0.086512712 | 0.083922449 | 0 |
| ATAACGCCAACTGGAG-1 | 1633966 | 27716 | 178350 | 133748 | 1294152 | 418 | 0.037747828 | 3.9454821 | 2.992149814 | 0.093127091 | 0.093264072 | 0 |
| ATAACGCCACTTTCCG-1 | 1491974 | 125952 | 147673 | 126689 | 1091660 | 353 | 0.031778458 | 3.356884067 | 3.080169376 | 0.09975734 | 0.09975734 | 0 |
| ATAACGCGTGCCATTA-1 | 1066366 | 13213 | 111443 | 87408 | 854302 | 276 | 0.024879644 | 2.628302433 | 3.011752509 | 0.110566824 | 0.109181002 | 0 |
| ATAACGCGTGCTCGAC-1 | 1482980 | 60366 | 150134 | 136338 | 1136142 | 367 | 0.033069285 | 3.479349867 | 3.098668788 | 0.096776066 | 0.097489479 | 0 |
| ATAACGCTCCACTTCG-1 | 1359418 | 87900 | 137189 | 116187 | 1018142 | 329 | 0.029627614 | 3.135118967 | 3.033646028 | 0.103041904 | 0.104082418 | 0 |
| ATAAGAGAGCGCCTTG-1 | 962836 | 18972 | 96333 | 81371 | 766160 | 247 | 0.022368929 | 2.3683318 | 2.923421025 | 0.116149964 | 0.1138576 | 0 |
| ATAAGAGAGTAGCCGA-1 | 1766028 | 24052 | 184462 | 162312 | 1395202 | 451 | 0.040708318 | 4.254402967 | 3.021925244 | 0.091070329 | 0.08999183 | 0 |
| ATAAGAGCAGCGTCAC-1 | 1923086 | 40385 | 192026 | 172885 | 1517790 | 490 | 0.044271089 | 4.6048537 | 3.047690745 | 0.089157223 | 0.087402921 | 0 |
| ATAAGAGTCCGTGCTT-1 | 2169648 | 76523 | 223539 | 186154 | 1683432 | 544 | 0.049035212 | 5.074431067 | 3.177838879 | 0.083238835 | 0.084094245 | 0 |
| ATAGACCGTACCGCTG-1 | 1066608 | 32690 | 109798 | 93470 | 830650 | 268 | 0.024171501 | 2.5663864 | 3.046567775 | 0.114307193 | 0.109126515 | 0 |
| ATAGACCGTCCGAATT-1 | 1419948 | 40923 | 145886 | 123415 | 1109724 | 358 | 0.032328595 | 3.3939711 | 3.038751651 | 0.093175952 | 0.094536517 | 0 |
| ATCACGAGTACCTAGT-1 | 1363550 | 10720 | 151820 | 114518 | 1086492 | 351 | 0.031754131 | 3.327121167 | 3.093358329 | 0.102322145 | 0.100989869 | 0 |
| ATCATCTCATTACGCA-1 | 1111666 | 57838 | 114091 | 95954 | 843783 | 273 | 0.024518383 | 2.6064685 | 3.041226719 | 0.115094106 | 0.114188872 | 0 |
| ATCATCTGTGCGAAAC-1 | 2060480 | 64865 | 216164 | 190571 | 1588880 | 513 | 0.046243751 | 4.809747367 | 2.706719126 | 0.086284701 | 0.0867528 | 0 |
| ATCATGGAGTCCAGGA-1 | 1407318 | 16771 | 152163 | 124295 | 1114089 | 360 | 0.032485614 | 3.419469933 | 3.094611289 | 0.098210797 | 0.09773197 | 0 |
| ATCCGAACAGCCAAAG-1 | 1578844 | 38222 | 172314 | 134998 | 1233310 | 398 | 0.035927912 | 3.769150133 | 3.080589255 | 0.096002777 | 0.094600629 | 0 |
| ATCCGAACAGTAATCC-1 | 2136110 | 72536 | 226911 | 199460 | 1637203 | 529 | 0.047484221 | 4.8896513 | 3.091830502 | 0.09093289 | 0.088214773 | 0 |
| ATCCGAACATTCCGAA-1 | 1630784 | 15642 | 179768 | 141355 | 1294019 | 418 | 0.037714044 | 3.945072 | 3.031704198 | 0.09351833 | 0.092960022 | 0 |
| ATCGAGTGTCGGAAGT-1 | 2006760 | 37408 | 211820 | 176055 | 1581477 | 511 | 0.046189328 | 4.7856537 | 3.024615079 | 0.08520628 | 0.085615149 | 0 |
| ATCGAGTTCTGTGAAC-1 | 2395576 | 46901 | 260437 | 216339 | 1871899 | 605 | 0.054497131 | 5.6056252 | 3.097042278 | 0.08097007 | 0.079564362 | 0 |
| ATCTACTAGGGATGGG-1 | 2243542 | 25124 | 240907 | 194336 | 1783175 | 576 | 0.052102973 | 5.372276467 | 3.062564372 | 0.081146123 | 0.077335945 | 0 |
| ATCTACTAGTCACGCC-1 | 1218714 | 77336 | 123496 | 105423 | 912459 | 295 | 0.026521964 | 2.813396533 | 3.037725533 | 0.113991438 | 0.111485821 | 0 |
| ATCTACTAGTTTGCGT-1 | 2018732 | 24268 | 218536 | 174761 | 1601167 | 517 | 0.046717501 | 4.852593867 | 3.083351685 | 0.084552506 | 0.085514678 | 0 |
| ATCTACTCAGTGCTGC-1 | 1869846 | 61539 | 197307 | 169369 | 1441631 | 466 | 0.041918418 | 4.3723061 | 3.09101308 | 0.092589504 | 0.092595723 | 0 |
| ATCTGCCCATTCGTTT-1 | 1907868 | 52012 | 206181 | 157952 | 1491723 | 482 | 0.04346324 | 4.524551767 | 3.087888905 | 0.091098702 | 0.090487793 | 0 |
| ATCTGCCGTCATGCCG-1 | 1474172 | 58169 | 155039 | 125925 | 1135039 | 367 | 0.032977291 | 3.472968533 | 2.919518031 | 0.100677608 | 0.096420616 | 0 |
| ATCTGCCTCCCGCATT-1 | 2262360 | 27429 | 245848 | 205020 | 1784063 | 576 | 0.05181607 | 5.352986367 | 3.021787313 | 0.082319085 | 0.08157567 | 0 |
| ATGAGGGAGAGGTAGA-1 | 1567174 | 14967 | 165020 | 136004 | 1251183 | 404 | 0.036573547 | 3.8403556 | 3.070503168 | 0.098352273 | 0.095818013 | 0 |
| ATGAGGGAGTTTGCGT-1 | 1008106 | 17222 | 107020 | 90035 | 793829 | 256 | 0.023240011 | 2.469176367 | 3.032866383 | 0.116876157 | 0.116876157 | 0 |
| ATGAGGGTCCGACCAG-1 | 1942688 | 56312 | 203040 | 174595 | 1508741 | 487 | 0.043975265 | 4.5877036 | 3.049438059 | 0.088278537 | 0.087012533 | 0 |
| ATGCGATCATTACAGC-1 | 1909072 | 52262 | 201697 | 169549 | 1485564 | 480 | 0.043211991 | 4.500713433 | 3.075562975 | 0.0891428 | 0.087521738 | 0 |
| ATGCGATGTCAAGTTC-1 | 2058878 | 104904 | 211876 | 184501 | 1557597 | 503 | 0.045104044 | 4.6538191 | 3.179768897 | 0.090574869 | 0.088390852 | 0 |
| ATGCGATGTGATCCTA-1 | 2085366 | 33770 | 224296 | 184043 | 1643257 | 531 | 0.04794138 | 4.963607333 | 3.071358547 | 0.087753191 | 0.087266572 | 0 |
| ATGCGATGTTTAGGAA-1 | 1470398 | 18040 | 158690 | 124689 | 1168979 | 378 | 0.034072363 | 3.5788921 | 3.121255826 | 0.095290695 | 0.094863344 | 0 |
| ATGGGAGAGTCCCACG-1 | 1766016 | 34326 | 189818 | 158677 | 1383195 | 447 | 0.040208424 | 4.2134936 | 3.035173595 | 0.094134316 | 0.090766932 | 0 |
| ATGGGAGGTACCGTGC-1 | 1385244 | 60296 | 139029 | 121939 | 1063980 | 344 | 0.030932442 | 3.262052533 | 3.137004396 | 0.102294989 | 0.101197135 | 0 |
| ATGGGAGTCGGAGAGT-1 | 1975858 | 34805 | 209927 | 169141 | 1561985 | 505 | 0.045619693 | 4.735090867 | 2.959424902 | 0.088808111 | 0.086337311 | 0 |
| ATGGGAGTCTTAAGGC-1 | 1826300 | 67326 | 192514 | 156042 | 1410418 | 456 | 0.041097733 | 4.296669133 | 3.115217947 | 0.089298604 | 0.089962246 | 0 |
| ATGGGAGTCTTCTAAC-1 | 2084424 | 22514 | 225818 | 184041 | 1652051 | 534 | 0.048239738 | 5.0031236 | 3.106227575 | 0.083307849 | 0.081645377 | 0 |
| ATGTGTGCAAAGTAAC-1 | 2569986 | 39138 | 275783 | 228889 | 2026176 | 655 | 0.05914677 | 6.091579033 | 3.188429787 | 0.075207603 | 0.076189977 | 0 |
| ATGTGTGGTCTCGTCT-1 | 1995796 | 74139 | 203644 | 177756 | 1540257 | 498 | 0.044687175 | 4.617085167 | 3.158436945 | 0.090986539 | 0.089578389 | 0 |
| ATTACTCAGTAATCCC-1 | 1337548 | 26864 | 141917 | 116809 | 1051958 | 340 | 0.030764596 | 3.250707433 | 3.011244655 | 0.105755499 | 0.106521129 | 0 |
| ATTACTCCACACACGC-1 | 1548624 | 45305 | 163553 | 137239 | 1202527 | 388 | 0.035001052 | 3.676174233 | 3.047204051 | 0.098126055 | 0.099315894 | 0 |
| ATTACTCCATCCCAAA-1 | 1987428 | 16436 | 219320 | 154340 | 1597332 | 516 | 0.046582195 | 4.822011433 | 3.771541333 | 0.141879491 | 0.140008827 | 1 |
| ATTACTCGTAAAGTCA-1 | 1744250 | 52039 | 180308 | 156162 | 1355741 | 438 | 0.039433127 | 4.127583 | 2.955779366 | 0.094309345 | 0.093588856 | 0 |
| ATTACTCGTAGCGTCC-1 | 1670256 | 59486 | 165415 | 144106 | 1301249 | 420 | 0.037846569 | 3.949823767 | 3.001519412 | 0.100548567 | 0.09522938 | 0 |
| ATTATCCAGCGATGAC-1 | 2217690 | 39668 | 241927 | 193510 | 1742585 | 563 | 0.050813994 | 5.2627629 | 3.063954025 | 0.084168073 | 0.081576849 | 0 |
| ATTATCCAGGAGTACC-1 | 2707544 | 35067 | 289054 | 243790 | 2139633 | 691 | 0.062329197 | 6.382326533 | 3.136278382 | 0.078477025 | 0.076157035 | 0 |
| ATTATCCTCCGATACA-1 | 2139058 | 21981 | 229721 | 196304 | 1691052 | 546 | 0.049317571 | 5.108521867 | 2.777008261 | 0.086556331 | 0.085288966 | 0 |
| ATTCTACAGAGCCCAA-1 | 2060292 | 22490 | 219590 | 182448 | 1635764 | 528 | 0.0477225 | 4.954347167 | 2.944145706 | 0.093042607 | 0.093128964 | 1 |
| ATTCTACAGCTAAGAT-1 | 2064014 | 14793 | 228841 | 169328 | 1651052 | 533 | 0.048195085 | 4.985337667 | 2.963087677 | 0.085101675 | 0.084175553 | 0 |
| ATTCTACAGCTATGCT-1 | 2596256 | 36588 | 281287 | 229536 | 2048845 | 662 | 0.059551579 | 6.0883123 | 2.989998153 | 0.081210287 | 0.081210287 | 0 |
| ATTCTACCAGCGCTTG-1 | 1452584 | 56756 | 152366 | 126294 | 1117168 | 361 | 0.03255095 | 3.4311143 | 3.105762673 | 0.104128246 | 0.100776669 | 0 |
| ATTCTACGTGGAAAGA-1 | 1591534 | 24363 | 169634 | 141647 | 1255890 | 406 | 0.036730197 | 3.857382267 | 3.144478617 | 0.093540158 | 0.094047604 | 0 |
| ATTCTACTCTCCCTAG-1 | 1510114 | 16220 | 164234 | 126805 | 1202855 | 389 | 0.035136408 | 3.6800159 | 3.137884749 | 0.098618769 | 0.09447671 | 0 |
| ATTGGACTCGTAGCGC-1 | 1502062 | 34205 | 155188 | 141608 | 1171061 | 378 | 0.034091309 | 3.5875044 | 3.118677292 | 0.102657624 | 0.098518056 | 0 |
| ATTGGTGAGGGATACC-1 | 2898504 | 36297 | 307873 | 264693 | 2289641 | 740 | 0.066589811 | 6.730545967 | 3.134991334 | 0.080125052 | 0.076933884 | 0 |
| ATTGGTGCACTAAGGG-1 | 1321824 | 25484 | 140623 | 109817 | 1045900 | 338 | 0.030539829 | 3.217182233 | 3.104599301 | 0.103206742 | 0.101168889 | 0 |
| ATTGGTGTCGCCAACG-1 | 1482642 | 49269 | 154248 | 132413 | 1146712 | 370 | 0.033319327 | 3.504995833 | 3.10595829 | 0.095741267 | 0.096858061 | 0 |
| ATTTCTGAGTAGGTGC-1 | 1845118 | 93176 | 189058 | 165375 | 1397509 | 451 | 0.040489794 | 4.219122133 | 2.985819783 | 0.09149749 | 0.090020103 | 0 |
| ATTTCTGCACACACGC-1 | 1595764 | 45601 | 169772 | 140549 | 1239842 | 401 | 0.036164772 | 3.794070933 | 3.047636296 | 0.098452769 | 0.096132399 | 0 |
| ATTTCTGCACTAACTG-1 | 1008686 | 21032 | 110333 | 83713 | 793608 | 256 | 0.02312485 | 2.451968367 | 3.018147209 | 0.117774508 | 0.114419546 | 0 |
| ATTTCTGTCACACGGC-1 | 1453340 | 97174 | 150025 | 124396 | 1081745 | 349 | 0.031395006 | 3.310948267 | 3.014474946 | 0.100164721 | 0.098925344 | 0 |
| CAACCAAAGTACGCGA-1 | 1597864 | 21992 | 172280 | 146078 | 1257514 | 406 | 0.036706816 | 3.863218233 | 3.036903409 | 0.094499379 | 0.094278815 | 0 |
| CAACCAACAGGACCAA-1 | 2383248 | 31888 | 263675 | 204989 | 1882696 | 608 | 0.054895062 | 5.644899967 | 3.420929191 | 0.113811415 | 0.113811415 | 1 |
| CAACCAAGTTCGATAC-1 | 2184686 | 56982 | 238559 | 189257 | 1699888 | 549 | 0.049453485 | 5.122475567 | 3.084998757 | 0.084065414 | 0.082895283 | 0 |
| CAACCTCAGACTTCCA-1 | 2057964 | 63195 | 219232 | 188474 | 1587063 | 513 | 0.046241875 | 4.8158543 | 2.985362665 | 0.087438411 | 0.086138597 | 0 |
| CAACCTCCAAGAACTA-1 | 1797418 | 41635 | 193005 | 164085 | 1398693 | 452 | 0.040783497 | 4.2561898 | 3.238520118 | 0.089537275 | 0.090395684 | 0 |
| CAACCTCTCAACTGCA-1 | 1758068 | 75293 | 185033 | 154587 | 1343155 | 434 | 0.039096418 | 4.095538067 | 3.049996094 | 0.091796676 | 0.093617344 | 0 |
| CAACCTCTCTGAAGCT-1 | 1979090 | 84286 | 207340 | 176098 | 1511366 | 488 | 0.043935039 | 4.5664448 | 3.149577445 | 0.089649852 | 0.088202545 | 0 |
| CAACTAGGTCGTCTTC-1 | 1909782 | 48781 | 201426 | 171116 | 1488459 | 481 | 0.043383801 | 4.525891533 | 3.108283196 | 0.085867864 | 0.085867864 | 0 |
| CAACTAGGTTTAGGAA-1 | 1891012 | 20874 | 206809 | 154091 | 1509238 | 488 | 0.044026727 | 4.576249533 | 3.190401091 | 0.105697113 | 0.105438172 | 1 |
| CAACTAGTCGACCATA-1 | 1807742 | 21965 | 195938 | 154350 | 1435489 | 464 | 0.041938014 | 4.371614367 | 3.012053755 | 0.09330378 | 0.090452159 | 0 |
| CAAGAAAAGCAAAGCC-1 | 1752374 | 64436 | 183082 | 159416 | 1345440 | 435 | 0.039222606 | 4.114444833 | 3.005206669 | 0.092162654 | 0.094093578 | 0 |
| CAAGAAACATAGGACG-1 | 1418324 | 45284 | 144868 | 130069 | 1098103 | 355 | 0.031899746 | 3.362620567 | 2.854314465 | 0.101363577 | 0.101363577 | 0 |
| CAAGAAAGTTTGTCTT-1 | 2477684 | 54833 | 267058 | 218578 | 1937215 | 626 | 0.056480331 | 5.823839 | 3.046951826 | 0.081697062 | 0.081875163 | 0 |
| CAAGAAATCGAACAAA-1 | 1458782 | 30671 | 156567 | 127737 | 1143807 | 369 | 0.033305719 | 3.495096533 | 3.197160995 | 0.09447116 | 0.091437441 | 0 |
| CAAGATCAGAATCGTA-1 | 1818014 | 34506 | 194101 | 158414 | 1430993 | 462 | 0.041712241 | 4.3520173 | 3.219108588 | 0.111694782 | 0.111551102 | 1 |
| CAAGATCAGGTGTTAA-1 | 1190032 | 19321 | 130293 | 94865 | 945553 | 305 | 0.027586259 | 2.907830533 | 3.017025357 | 0.103580542 | 0.103429902 | 0 |
| CAAGATCCAATCCATG-1 | 1473704 | 10809 | 161272 | 121793 | 1179830 | 381 | 0.034425745 | 3.611339467 | 3.066071505 | 0.105198217 | 0.101471305 | 1 |
| CAAGATCCATTACGCA-1 | 1066406 | 53836 | 107686 | 93919 | 810965 | 262 | 0.023568025 | 2.509716 | 2.916521085 | 0.120624669 | 0.115225017 | 0 |
| CAAGATCGTGCTTAGT-1 | 930938 | 14282 | 100135 | 79489 | 737032 | 238 | 0.021535447 | 2.284486133 | 3.12105009 | 0.12361219 | 0.120276877 | 0 |
| CAAGATCGTGTTCGAT-1 | 1884784 | 23427 | 205586 | 158409 | 1497362 | 484 | 0.043685095 | 4.5423082 | 3.116891604 | 0.092643431 | 0.090361415 | 0 |
| CAAGATCGTTGAACAA-1 | 1925126 | 41205 | 209169 | 150760 | 1523992 | 492 | 0.044413612 | 4.6114049 | 3.780541555 | 0.137842668 | 0.136743016 | 1 |

| | | | | | | | | | | | | |
|---|---|---|---|---|---|---|---|---|---|---|---|---|
| CAAGATCTCACACGTA-1 | 991914 | 12484 | 107692 | 81580 | 790158 | 255 | 0.023013542 | 2.435500633 | 3.037202994 | 0.121628821 | 0.116786107 | 0 |
| CAAGGCCCAAACCCTA-1 | 1982342 | 34460 | 210586 | 177131 | 1560165 | 504 | 0.045499442 | 4.738578333 | 3.207876622 | 0.084360017 | 0.084360017 | 0 |
| CAAGGCCGTCACCTGG-1 | 1642794 | 16366 | 174084 | 150650 | 1301694 | 420 | 0.037984392 | 3.980356533 | 3.045971052 | 0.09541856 | 0.094411553 | 0 |
| CAAGTTGTCGCGGCAT-1 | 2345074 | 44713 | 249073 | 205616 | 1845672 | 596 | 0.053863238 | 5.560302867 | 3.143348322 | 0.082821352 | 0.083367702 | 0 |
| CACAAACCAAGAATCA-1 | 1729018 | 67888 | 182399 | 160697 | 1318034 | 426 | 0.038366752 | 4.009094767 | 3.032866651 | 0.09653591 | 0.091907152 | 0 |
| CACAAACCAGGCCTTG-1 | 1351118 | 17069 | 145960 | 120904 | 1067185 | 345 | 0.031105434 | 3.278082667 | 2.986091577 | 0.096941961 | 0.097064539 | 0 |
| CACAAACCATCAAGAA-1 | 1165236 | 17462 | 128755 | 96458 | 922561 | 298 | 0.026889665 | 2.838878167 | 3.047493928 | 0.11209445 | 0.109224276 | 0 |
| CACAAACTCGAGAGGT-1 | 2443022 | 24979 | 267409 | 206032 | 1944602 | 628 | 0.056820198 | 5.843351667 | 3.049721166 | 0.082299467 | 0.079579961 | 0 |
| CACACAAGTACGTTTG-1 | 928390 | 20228 | 101387 | 77988 | 728787 | 235 | 0.021232293 | 2.263496533 | 3.01765625 | 0.117243272 | 0.116471876 | 0 |
| CACACAAGTTGCACTA-1 | 1933074 | 34106 | 212271 | 170927 | 1515770 | 490 | 0.044184454 | 4.600446 | 3.137858782 | 0.084475217 | 0.084982044 | 0 |
| CACACCTGTACGTGCC-1 | 2062168 | 35032 | 220476 | 187637 | 1619023 | 523 | 0.047089689 | 4.889799167 | 3.141059062 | 0.0865587 | 0.086558862 | 0 |
| CACACTCAGACGTCGA-1 | 2053630 | 22949 | 217545 | 186061 | 1627075 | 526 | 0.047478526 | 4.942961233 | 3.141051047 | 0.082146815 | 0.083602286 | 0 |
| CACACTCAGCGTAGTG-1 | 1192988 | 6904 | 129618 | 95532 | 960934 | 310 | 0.028099984 | 2.955771133 | 3.122481149 | 0.106505245 | 0.103953171 | 0 |
| CACACTCGTTAAAGGT-1 | 2103860 | 23152 | 230295 | 182487 | 1667926 | 539 | 0.048760026 | 5.048418333 | 3.009809841 | 0.08461972 | 0.08477514 | 0 |
| CACACTCTCTACAGTG-1 | 1359016 | 9662 | 152476 | 103334 | 1093544 | 353 | 0.031936724 | 3.341532667 | 3.001150617 | 0.098627228 | 0.096331187 | 0 |
| CACACTCTCTTCTTGA-1 | 2613982 | 33908 | 289054 | 199878 | 2091142 | 676 | 0.06108842 | 6.2823981 | 2.712066405 | 0.114845132 | 0.114845132 | 1 |
| CACAGGCCATAGTGTC-1 | 2029164 | 38403 | 212992 | 170413 | 1607356 | 519 | 0.046819279 | 4.855185333 | 2.998415799 | 0.092037576 | 0.086417265 | 0 |
| CACAGGCGTAGTCTGT-1 | 2309698 | 28174 | 250334 | 204225 | 1826965 | 590 | 0.053342574 | 5.4959476 | 3.164826655 | 0.078036549 | 0.078651936 | 0 |
| CACAGGCTCCGTCTAC-1 | 2688526 | 49236 | 285156 | 249771 | 2104363 | 680 | 0.061367495 | 6.295448267 | 2.876395234 | 0.071789887 | 0.07251362 | 0 |
| CACAGTAAGTCGTACT-1 | 2200514 | 39005 | 237214 | 193563 | 1730732 | 559 | 0.050397763 | 5.219707967 | 3.081170377 | 0.086395515 | 0.085912878 | 0 |
| CACATAGAGGTATCGG-1 | 1183238 | 10630 | 126100 | 108176 | 938332 | 303 | 0.027391 | 2.893459 | 2.914334212 | 0.114091348 | 0.110658795 | 0 |
| CACATAGAGTCGTCCG-1 | 1515588 | 57818 | 155019 | 134938 | 1167813 | 377 | 0.033978205 | 3.580842433 | 3.169035695 | 0.096275507 | 0.095912789 | 0 |
| CACATAGCACAACTTG-1 | 1050146 | 17569 | 113673 | 88780 | 830124 | 268 | 0.024182653 | 2.565279033 | 3.093081386 | 0.110535305 | 0.112199818 | 0 |
| CACATAGGTACAATAG-1 | 1199238 | 12084 | 131995 | 93874 | 961285 | 311 | 0.028054842 | 2.950229667 | 3.188603531 | 0.109001291 | 0.108953086 | 0 |
| CACATAGGTTGGTGAG-1 | 2356402 | 20803 | 252947 | 197040 | 1885612 | 609 | 0.05507309 | 5.6918504 | 3.06920639 | 0.080163426 | 0.080163426 | 0 |
| CACATAGTCTCAGTTT-1 | 1270040 | 47238 | 135468 | 104184 | 983150 | 318 | 0.028613145 | 3.015816267 | 3.049484492 | 0.106395602 | 0.103756911 | 0 |
| CACATTTAGCAGCGTA-1 | 1491390 | 19800 | 161357 | 131303 | 1178930 | 381 | 0.034392231 | 3.6129607 | 2.948396352 | 0.098328031 | 0.098052822 | 0 |
| CACATTTAGGAATCGC-1 | 1788962 | 34177 | 185234 | 172467 | 1397084 | 451 | 0.040719766 | 4.253678067 | 3.046801074 | 0.094168534 | 0.092212784 | 0 |
| CACATTTCATGGAGAC-1 | 1666454 | 92810 | 172264 | 143753 | 1257627 | 406 | 0.036591425 | 3.8443176 | 3.035002735 | 0.096454132 | 0.094607793 | 0 |
| CACATTTGTCGCTTCT-1 | 1757246 | 135783 | 174807 | 147486 | 1299170 | 420 | 0.03781055 | 3.9602074 | 3.001122191 | 0.095752837 | 0.095752837 | 0 |
| CACATTTGTTTCCTGC-1 | 2148104 | 51278 | 231960 | 185871 | 1678995 | 542 | 0.048864179 | 5.043830467 | 3.042189087 | 0.08653235 | 0.08637982 | 0 |
| CACCACTCATCTAAGC-1 | 1668586 | 59918 | 176146 | 144962 | 1287560 | 416 | 0.037440282 | 3.926901567 | 3.048625072 | 0.097970564 | 0.093402462 | 0 |
| CACCACTGTCATGTTG-1 | 1316502 | 9603 | 145111 | 104483 | 1057305 | 342 | 0.030818023 | 3.234075733 | 3.200086642 | 0.105972239 | 0.103616163 | 0 |
| CACCAGGAGACAGTTA-1 | 1559160 | 19763 | 171614 | 129296 | 1238487 | 400 | 0.03612783 | 3.785085533 | 3.129568602 | 0.095698047 | 0.096346331 | 0 |
| CACCAGGAGGAGTAGA-1 | 1392524 | 17119 | 150017 | 121675 | 1103713 | 357 | 0.032289982 | 3.404810733 | 3.049786902 | 0.098582256 | 0.09949105 | 0 |
| CACCAGGCAATCTCTT-1 | 1906360 | 104015 | 196994 | 170916 | 1434435 | 463 | 0.041766678 | 4.3389478 | 3.070338194 | 0.091310537 | 0.087684687 | 0 |
| CACCAGGCACGGCTGT-1 | 1937944 | 30805 | 208909 | 176708 | 1521522 | 491 | 0.044396381 | 4.6231115 | 3.067488367 | 0.090932528 | 0.088326177 | 0 |
| CACCAGGCATTACTTC-1 | 1964936 | 116751 | 198865 | 179890 | 1469430 | 475 | 0.042561497 | 4.423673 | 3.049172291 | 0.091000425 | 0.089485077 | 0 |
| CACCTTGAGGCTCTCG-1 | 1682028 | 30158 | 179114 | 152624 | 1320132 | 426 | 0.038484609 | 4.046814633 | 3.077061301 | 0.096785777 | 0.091425229 | 0 |
| CACCTTGCAGACACAG-1 | 1506556 | 13544 | 165383 | 126112 | 1201517 | 388 | 0.035047324 | 3.675612133 | 2.999730075 | 0.097557947 | 0.096757213 | 0 |
| CACCTTGCAGACAGTG-1 | 1167746 | 7345 | 129372 | 96153 | 934876 | 302 | 0.027236624 | 2.875836167 | 2.769659649 | 0.112960963 | 0.108018225 | 0 |
| CACCTTGTCCGACGTG-1 | 1201956 | 10831 | 130159 | 101537 | 959429 | 310 | 0.027912864 | 2.948543467 | 3.114056773 | 0.109595445 | 0.10604133 | 0 |
| CACTCCAAGCAGACTG-1 | 1537726 | 14797 | 169410 | 132680 | 1220839 | 394 | 0.035390546 | 3.7114328 | 3.002130863 | 0.099970291 | 0.097099322 | 0 |
| CACTCCAAGTGTCCAT-1 | 3946044 | 36235 | 431076 | 350045 | 3128688 | 1011 | 0.091421551 | 9.2212151 | 3.141047874 | 0.062160242 | 0.063227203 | 0 |
| CAGAATCAGGGATGGG-1 | 2840352 | 24671 | 308520 | 243478 | 2263683 | 731 | 0.066114011 | 6.7573483 | 3.108206436 | 0.081856135 | 0.081856135 | 1 |
| CAGAATCTCGAAAGGC-1 | 1891002 | 60973 | 196657 | 170970 | 1462402 | 472 | 0.042655192 | 4.458532267 | 3.02222317 | 0.087376953 | 0.087376953 | 0 |
| CAGAATCTCGGAATGG-1 | 1225462 | 11899 | 130654 | 104135 | 978774 | 316 | 0.028550801 | 3.011946667 | 3.04646852 | 0.112917241 | 0.111593102 | 0 |
| CAGAATCTCTACGGGC-1 | 2253986 | 71928 | 235339 | 199186 | 1747533 | 565 | 0.050911135 | 5.281735667 | 3.146310803 | 0.07902844 | 0.079290348 | 0 |
| CAGAGAGAGTCGTACT-1 | 2352722 | 50537 | 245415 | 206056 | 1850714 | 598 | 0.053917879 | 5.5658189 | 3.001221634 | 0.08711482 | 0.084943367 | 0 |
| CAGAGAGAGTGAAGAG-1 | 1478856 | 20427 | 156862 | 124942 | 1176625 | 380 | 0.034324704 | 3.608175967 | 3.10594051 | 0.101274798 | 0.100235971 | 0 |
| CAGAGAGCACGAGCTC-1 | 1774744 | 52820 | 186143 | 162178 | 1373603 | 444 | 0.040001568 | 4.185607533 | 3.13860261 | 0.090185855 | 0.090544604 | 0 |
| CAGAGAGTCCGAAGCC-1 | 1705046 | 63664 | 176436 | 154110 | 1310836 | 423 | 0.038233998 | 4.019616833 | 3.100105105 | 0.09371606 | 0.092919834 | 0 |
| CAGAGAGTCCGAGATT-1 | 1155904 | 42167 | 121931 | 94700 | 897106 | 290 | 0.026197245 | 2.763708733 | 3.234893514 | 0.109084548 | 0.109084548 | 0 |
| CAGATCAAGAAGGGTA-1 | 1734182 | 17335 | 186685 | 147320 | 1382842 | 447 | 0.040384936 | 4.216310267 | 2.981912861 | 0.09170747 | 0.091885929 | 0 |
| CAGATCACATGCCGCA-1 | 1223384 | 68330 | 125705 | 102848 | 926501 | 299 | 0.026925257 | 2.851030033 | 3.038350931 | 0.116217107 | 0.114460693 | 1 |
| CAGATCAGTTCTGTCC-1 | 2148434 | 83796 | 226658 | 188404 | 1649576 | 533 | 0.047881561 | 4.9450431 | 3.033485854 | 0.08925158 | 0.09032128 | 0 |
| CAGCAGCGTTCAAGGG-1 | 1627262 | 23578 | 173280 | 140863 | 1289541 | 417 | 0.037623303 | 3.9476944 | 3.110804976 | 0.098387713 | 0.096416835 | 0 |
| CAGCAGCTCCCGGACA-1 | 1666458 | 67209 | 172420 | 150578 | 1276251 | 412 | 0.037159283 | 3.8980215 | 3.257905883 | 0.124362349 | 0.123446837 | 1 |
| CAGCAGCTCGCAACGC-1 | 1471326 | 29835 | 153922 | 127792 | 1159777 | 375 | 0.033759072 | 3.5558087 | 2.99610812 | 0.1005587 | 0.098032183 | 0 |
| CAGCATAAGCGTTGCC-1 | 1967240 | 32020 | 211599 | 175080 | 1548541 | 500 | 0.045222155 | 4.707643133 | 3.020674515 | 0.089996748 | 0.088435738 | 0 |
| CAGCATACAAACGGGT-1 | 2404860 | 50224 | 257668 | 217096 | 1879872 | 607 | 0.054887542 | 5.659009167 | 3.021835643 | 0.080833971 | 0.079576122 | 0 |
| CAGCATACACGGAATG-1 | 1409624 | 14844 | 153066 | 117269 | 1124445 | 363 | 0.032796582 | 3.444442667 | 3.09794953 | 0.104063051 | 0.101565969 | 0 |
| CAGCATAGTCCGCAGT-1 | 2033658 | 30808 | 217829 | 183230 | 1601791 | 517 | 0.046681942 | 4.846023567 | 3.122813748 | 0.089004641 | 0.087618766 | 0 |
| CAGCCGAAGCGTCAAG-1 | 1362142 | 12211 | 142308 | 121258 | 1086365 | 351 | 0.031720829 | 3.341885167 | 3.045185301 | 0.103428574 | 0.103428574 | 0 |
| CAGCCGACAAACCGAG-1 | 1934040 | 42893 | 208502 | 169865 | 1512780 | 489 | 0.044060794 | 4.592530967 | 3.354003441 | 0.114962202 | 0.114627548 | 1 |
| CAGCGACAGCGCCTCA-1 | 1779958 | 31732 | 188649 | 162297 | 1397280 | 451 | 0.040733308 | 4.2494217 | 3.096851647 | 0.089900021 | 0.089784253 | 0 |
| CAGCGACCAAGTCGGA-1 | 1834776 | 26808 | 197830 | 162518 | 1447620 | 468 | 0.042207866 | 4.410499733 | 3.038100854 | 0.092498097 | 0.091444546 | 0 |
| CAGCTAATCCACTTCG-1 | 1364382 | 34342 | 146702 | 127806 | 1055532 | 341 | 0.030690493 | 3.2488078 | 3.144119569 | 0.097905374 | 0.098244086 | 0 |
| CAGCTAATCGAAAGGC-1 | 2131100 | 53270 | 225661 | 195028 | 1657141 | 535 | 0.048249805 | 5.022174367 | 3.036382367 | 0.086622451 | 0.085172597 | 0 |
| CAGCTGGAGCCACCTG-1 | 1344510 | 9808 | 144109 | 119732 | 1070861 | 346 | 0.031148232 | 3.2741524 | 2.991271555 | 0.103786316 | 0.101445495 | 0 |
| CAGCTGGGTCCTTGGG-1 | 1598564 | 17058 | 172500 | 141194 | 1267812 | 410 | 0.037042855 | 3.8858174 | 3.022776503 | 0.0903093 | 0.092059113 | 0 |
| CAGCTGGTCACGGTAT-1 | 2317042 | 32578 | 249417 | 191528 | 1843519 | 596 | 0.053706398 | 5.5315445 | 3.001349566 | 0.083494453 | 0.080823439 | 0 |
| CAGGTGCCAGCATGTT-1 | 1211912 | 13658 | 131366 | 94811 | 972077 | 314 | 0.028363309 | 2.9803552 | 3.144247505 | 0.104912773 | 0.105203312 | 0 |
| CAGGTGCTCGAGGTGA-1 | 1542410 | 19206 | 161765 | 132391 | 1229048 | 397 | 0.035856848 | 3.763961 | 3.13177353 | 0.098896097 | 0.096888131 | 0 |
| CAGTAACAGACCACGA-1 | 3172984 | 34101 | 344305 | 283953 | 2510625 | 811 | 0.07325867 | 7.483114167 | 3.086413789 | 0.067127382 | 0.067339226 | 0 |
| CAGTAACAGCACCGCT-1 | 1872596 | 97273 | 192526 | 164935 | 1417862 | 458 | 0.041293209 | 4.325315767 | 3.109573569 | 0.091550959 | 0.091788048 | 0 |
| CAGTAACCACACCTAA-1 | 1163364 | 10172 | 129244 | 95135 | 928813 | 300 | 0.027140454 | 2.860819867 | 3.009871598 | 0.107057769 | 0.106233733 | 0 |
| CAGTAACCATACGTTG-1 | 960866 | 8542 | 105429 | 74022 | 772873 | 250 | 0.022603045 | 2.391350833 | 3.144620148 | 0.123669736 | 0.119851878 | 0 |
| CAGTCCTAGTCAAGGC-1 | 1663460 | 54138 | 176876 | 145983 | 1286463 | 416 | 0.037426059 | 3.921003433 | 3.120669082 | 0.0943415 | 0.09327306 | 0 |
| CAGTCCTGTCTTCACC-1 | 2654606 | 53363 | 285662 | 236403 | 2079178 | 672 | 0.060637447 | 6.2349062 | 3.022116443 | 0.075799411 | 0.075529548 | 0 |
| CATATGGTCTAGACAC-1 | 2190552 | 49579 | 227484 | 210433 | 1703056 | 550 | 0.049611835 | 5.1519616 | 3.099322414 | 0.082441828 | 0.082654153 | 0 |
| CATATTCCAGTAGAAT-1 | 944598 | 5651 | 102968 | 78281 | 757698 | 245 | 0.022194856 | 2.344679733 | 3.035925592 | 0.12605078 | 0.121101664 | 0 |
| CATATTCGTCGCGCTA-1 | 1456806 | 45022 | 145073 | 133927 | 1132784 | 366 | 0.033009108 | 3.4713631 | 3.062532403 | 0.099785576 | 0.099723478 | 0 |
| CATATTCGTGACCTGC-1 | 2194712 | 48274 | 231466 | 194156 | 1720816 | 556 | 0.05014238 | 5.1916818 | 3.142990116 | 0.081806639 | 0.083317721 | 0 |
| CATATTCTCTTGCTTA-1 | 1745252 | 19774 | 186370 | 152115 | 1386993 | 448 | 0.04049298 | 4.230865933 | 3.119178345 | 0.094213673 | 0.09163831 | 0 |
| CATCAAGTCCTACGAA-1 | 1292354 | 14362 | 142175 | 112046 | 1023771 | 331 | 0.029808468 | 3.144663133 | 3.103958076 | 0.098270695 | 0.100233643 | 0 |
| CATCAAGTCTAGCCTC-1 | 1936188 | 42093 | 203728 | 176775 | 1513592 | 489 | 0.044182151 | 4.612622067 | 3.067859223 | 0.090567345 | 0.088854853 | 0 |
| CATCAAGTCTCCATGC-1 | 861250 | 19557 | 93483 | 67811 | 680399 | 220 | 0.019937511 | 2.120938 | 3.072394624 | 0.119737478 | 0.120953151 | 0 |
| CATCAGAGTTAGTGAA-1 | 1848376 | 16636 | 202758 | 158478 | 1470504 | 475 | 0.042867983 | 4.4630392 | 3.022221907 | 0.087816962 | 0.087834853 | 0 |
| CATCCACCATTACAGC-1 | 2145348 | 64069 | 229629 | 180207 | 1671443 | 540 | 0.048604205 | 5.0351886 | 2.897610897 | 0.086157784 | 0.084686221 | 0 |
| CATCCACTCCTGCGTT-1 | 1880724 | 46527 | 201187 | 161541 | 1471469 | 475 | 0.042860826 | 4.4599898 | 3.15650086 | 0.088925049 | 0.0895555 | 0 |
| CATCCACTCCTTGAAG-1 | 1150670 | 9958 | 128273 | 97456 | 914983 | 296 | 0.02677273 | 2.824968 | 3.037549548 | 0.107877789 | 0.107370924 | 0 |
| CATCGAACAATGGGAC-1 | 2130288 | 106028 | 222168 | 182676 | 1619416 | 523 | 0.04709756 | 4.904087 | 3.060867218 | 0.088242545 | 0.084986262 | 0 |
| CATCGAACACGACGCT-1 | 1490770 | 89436 | 151836 | 126513 | 1122985 | 363 | 0.032646802 | 3.4358403 | 3.105751747 | 0.12825662 | 0.129381225 | 1 |

| | | | | | | | | | | | | |
|---|---|---|---|---|---|---|---|---|---|---|---|---|
| CATCGAACATTTCGTT-1 | 1699224 | 27548 | 184226 | 144439 | 1343011 | 434 | 0.039102581 | 4.086608867 | 3.104212246 | 0.095348887 | 0.091467305 | 0 |
| CATCGAAGTGCTCGAC-1 | 1639564 | 43619 | 172083 | 152945 | 1270917 | 411 | 0.036959962 | 3.8742628 | 3.095286114 | 0.094336494 | 0.094012531 | 0 |
| CATCGAAGTTTGGCGC-1 | 1335302 | 48508 | 137663 | 119909 | 1029222 | 332 | 0.029960412 | 3.1659727 | 3.045423551 | 0.106472668 | 0.105051972 | 0 |
| CATCGGGCACCCAAAT-1 | 2032452 | 27036 | 223609 | 166025 | 1615782 | 522 | 0.047116435 | 4.8829961 | 3.097529434 | 0.089316046 | 0.085285175 | 0 |
| CATCGGGTCTCAGTTT-1 | 1473728 | 69926 | 156649 | 119774 | 1127379 | 364 | 0.032778375 | 3.441406367 | 3.044830182 | 0.101884961 | 0.100650497 | 0 |
| CATGACACAACTCGCG-1 | 777788 | 71451 | 75117 | 65515 | 565705 | 183 | 0.01640252 | 1.762613333 | 3.046929691 | 0.137557003 | 0.132870605 | 0 |
| CATGACAGTTTACTCT-1 | 2172104 | 95810 | 225968 | 191806 | 1658520 | 536 | 0.048025304 | 4.939264933 | 3.189062078 | 0.097212671 | 0.095394562 | 1 |
| CATGACATCTCCGGAG-1 | 1568590 | 20516 | 166384 | 138106 | 1243584 | 402 | 0.03631234 | 3.811180633 | 3.099857205 | 0.099159135 | 0.097825721 | 0 |
| CATGCCTGTCCGCAAC-1 | 2225014 | 34727 | 236967 | 202839 | 1750481 | 565 | 0.050945972 | 5.2705876 | 3.026614943 | 0.084344354 | 0.084453862 | 0 |
| CATGCCTGTTACGGCC-1 | 1722280 | 35215 | 182601 | 164982 | 1339482 | 433 | 0.038976587 | 4.074889233 | 3.060549664 | 0.096649873 | 0.093045758 | 0 |
| CATGCCTTCATGTGAC-1 | 2072428 | 46502 | 219769 | 185529 | 1620628 | 524 | 0.047210641 | 4.9043615 | 3.130275138 | 0.083843643 | 0.084605473 | 0 |
| CATGGCGAGATATGGT-1 | 1958860 | 19573 | 211885 | 158726 | 1568676 | 507 | 0.045845386 | 4.7574801 | 3.045589552 | 0.089383324 | 0.086786014 | 0 |
| CATGGCGCAGTCGGAA-1 | 797928 | 12642 | 84291 | 66710 | 634285 | 205 | 0.018594529 | 1.9882921 | 3.108719114 | 0.131005622 | 0.129136956 | 0 |
| CATGGCGCATTTGCGA-1 | 555382 | 6119 | 57978 | 50679 | 440606 | 142 | 0.012879764 | 1.389392667 | 2.910910847 | 0.155277681 | 0.153079916 | 0 |
| CATGGCGTCGAGCACC-1 | 832524 | 22845 | 71074 | 75381 | 663224 | 214 | 0.019352581 | 2.070670133 | 3.134123828 | 0.124376726 | 0.127485053 | 0 |
| CATTATCAGCGTGAAC-1 | 2578918 | 49940 | 272708 | 225105 | 2031165 | 656 | 0.059238997 | 6.088024067 | 3.047621754 | 0.079983391 | 0.079364727 | 0 |
| CATTATCCAACAGCCC-1 | 1041272 | 39354 | 107291 | 94142 | 800485 | 259 | 0.023389264 | 2.490006533 | 3.135348385 | 0.120549121 | 0.120193261 | 0 |
| CATTATCCATAGGTCT-1 | 4119828 | 157262 | 412876 | 362343 | 3187347 | 1030 | 0.091834417 | 9.090414267 | 3.177268076 | 0.085530783 | 0.085530783 | 1 |
| CATTATCGTAAGAAGG-1 | 1289738 | 8519 | 134863 | 111724 | 1034632 | 334 | 0.030267281 | 3.1898554 | 3.061892721 | 0.103716114 | 0.103745886 | 0 |
| CATTCGCAGCAAATCA-1 | 2069318 | 94509 | 214843 | 179313 | 1580653 | 511 | 0.046019035 | 4.7839259 | 3.009198488 | 0.08562156 | 0.08617256 | 0 |
| CATTCGCCACGCACGT-1 | 1499436 | 27556 | 158099 | 136232 | 1177549 | 380 | 0.034345838 | 3.5979809 | 3.004603417 | 0.104500002 | 0.099901386 | 0 |
| CCAATCCGTCAAGCCC-1 | 1518854 | 83271 | 154447 | 131380 | 1149756 | 371 | 0.033475339 | 3.527414267 | 2.930275177 | 0.100931798 | 0.099227766 | 0 |
| CCAATCCTCCGACGTG-1 | 1038500 | 16441 | 111213 | 86067 | 824779 | 266 | 0.023961547 | 2.5396019 | 2.987573053 | 0.121559224 | 0.11735385 | 1 |
| CCACCTAAGCTCCTTC-1 | 2333238 | 54287 | 247855 | 213557 | 1817539 | 587 | 0.05291587 | 5.4486446 | 3.109640711 | 0.084620198 | 0.081204889 | 0 |
| CCACCTAAGTGGTAGC-1 | 2819020 | 23751 | 314679 | 248923 | 2231667 | 721 | 0.065052307 | 6.648738 | 3.021405077 | 0.079658054 | 0.078581152 | 0 |
| CCACCTAAGTTAGGTA-1 | 1533368 | 11662 | 168247 | 123262 | 1230197 | 397 | 0.03587655 | 3.761328533 | 3.04702682 | 0.102283183 | 0.100528589 | 1 |
| CCACCTAGTTCGAGCC-1 | 1443238 | 84463 | 146910 | 129206 | 1082659 | 350 | 0.031516396 | 3.3330021 | 2.907310733 | 0.104482803 | 0.104008508 | 0 |
| CCACGGAAGACTGTAA-1 | 1362894 | 15869 | 145102 | 108168 | 1093755 | 353 | 0.031715842 | 3.3277473 | 3.081269399 | 0.106512975 | 0.102926828 | 0 |
| CCACGGACAAAGTGGC-1 | 2764182 | 33997 | 303195 | 222935 | 2204055 | 712 | 0.064280665 | 6.584963233 | 4.319796485 | 0.147060799 | 0.147060799 | 1 |
| CCACGGACAGAAGACC-1 | 2244778 | 60109 | 234007 | 200669 | 1749993 | 565 | 0.050920427 | 5.2669944 | 3.11915335 | 0.082582582 | 0.083585775 | 0 |
| CCACTACAGACAAAGG-1 | 1303838 | 10658 | 142717 | 111096 | 1039367 | 336 | 0.030349435 | 3.203572967 | 3.079971586 | 0.102440786 | 0.10300518 | 0 |
| CCACTACAGAGACTAT-1 | 1957250 | 22063 | 213289 | 166056 | 1555842 | 503 | 0.045391394 | 4.7124118 | 3.027965271 | 0.089037625 | 0.087139292 | 1 |
| CCACTACTCTACGGCG-1 | 1005072 | 64878 | 101137 | 87973 | 751084 | 243 | 0.021789337 | 2.323451367 | 3.056400133 | 0.116816241 | 0.118947753 | 0 |
| CCACTACTCTCGGAAT-1 | 2590632 | 36409 | 280649 | 223299 | 2050275 | 662 | 0.05977173 | 6.143101433 | 3.118657072 | 0.080655487 | 0.077971244 | 0 |
| CCAGCGATCTACAGTG-1 | 1080628 | 14437 | 117828 | 85861 | 862502 | 279 | 0.025019448 | 2.639526633 | 2.983874518 | 0.1131981 | 0.112845636 | 0 |
| CCATGTCTCGAGTACT-1 | 2195948 | 60394 | 232538 | 189247 | 1713769 | 554 | 0.049952154 | 5.1725539 | 3.030984075 | 0.091623391 | 0.088923945 | 0 |
| CCATGTCTCGCTGAAT-1 | 1423912 | 19745 | 152685 | 125435 | 1126047 | 364 | 0.032807593 | 3.432056633 | 3.130147884 | 0.10012276 | 0.100492304 | 0 |
| CCATTCGAGTGCTGCC-1 | 2016674 | 54727 | 212415 | 177452 | 1572080 | 508 | 0.045870466 | 4.762353567 | 3.02468706 | 0.091874845 | 0.088054897 | 0 |
| CCATTCGCACATAAAG-1 | 1134864 | 12700 | 125307 | 91455 | 905402 | 292 | 0.026394447 | 2.785054533 | 3.106716476 | 0.111821729 | 0.111062171 | 0 |
| CCCAATCGTCGACCAC-1 | 2185784 | 51776 | 222876 | 200972 | 1710160 | 552 | 0.049845776 | 5.167617067 | 2.955684555 | 0.084922902 | 0.084186368 | 0 |
| CCCAGTTAGCACGCCT-1 | 2071046 | 60665 | 216533 | 190897 | 1602951 | 518 | 0.046680255 | 4.856996933 | 3.174450159 | 0.089537421 | 0.086243707 | 0 |
| CCCATACAGCCACGTC-1 | 1188740 | 30847 | 126489 | 102521 | 928883 | 300 | 0.027002391 | 2.856022367 | 2.916234532 | 0.111025193 | 0.107253706 | 0 |
| CCCATACCAACGAAAT-1 | 1926286 | 33499 | 206859 | 168495 | 1517433 | 490 | 0.044252309 | 4.596507033 | 2.917374627 | 0.089063067 | 0.087354741 | 0 |
| CCCATACCACAAGCTT-1 | 1789668 | 116127 | 182162 | 151371 | 1340008 | 433 | 0.038954023 | 4.079265767 | 3.043598375 | 0.094062886 | 0.092869364 | 0 |
| CCCATACCATCTTTCA-1 | 1726126 | 78372 | 175896 | 155637 | 1316221 | 425 | 0.038309048 | 4.018839167 | 2.989254109 | 0.09121458 | 0.09198234 | 0 |
| CCCTCCTAGCAATCTC-1 | 2956678 | 50191 | 315387 | 251316 | 2339784 | 756 | 0.068251678 | 6.977899467 | 2.769408519 | 0.101589542 | 0.101589542 | 1 |
| CCCTCCTCAGAATTCC-1 | 1609118 | 78850 | 162984 | 143542 | 1223742 | 395 | 0.035537397 | 3.718200133 | 3.017356845 | 0.102890529 | 0.098055735 | 0 |
| CCCTCCTGTATTCGCA-1 | 1118800 | 39483 | 114055 | 103616 | 861646 | 278 | 0.025080974 | 2.654751067 | 3.034976604 | 0.117007329 | 0.113952885 | 0 |
| CCGGGATCACCAATGT-1 | 1278264 | 13265 | 137914 | 105949 | 1021136 | 330 | 0.02978967 | 3.124733633 | 3.036200381 | 0.100532822 | 0.102965265 | 0 |
| CCGGTAGAGCCAGTTT-1 | 1334556 | 58485 | 132447 | 109832 | 1033792 | 334 | 0.030072848 | 3.1621244 | 2.989823094 | 0.109399071 | 0.107345104 | 0 |
| CCGGTAGAGCCTATGT-1 | 416750 | 5912 | 42314 | 32470 | 336054 | 109 | 0.009772588 | 1.048926633 | 3.00515398 | 0.177187606 | 0.172792232 | 0 |
| CCGGTAGGTTCAGTAC-1 | 2180036 | 58397 | 225555 | 187195 | 1708889 | 552 | 0.049744978 | 5.1313449 | 3.002486431 | 0.087660932 | 0.086101791 | 0 |
| CCGGTAGTCGAGTTGT-1 | 2292410 | 35433 | 239561 | 196162 | 1821254 | 588 | 0.053260601 | 5.503595867 | 3.003019174 | 0.082671635 | 0.081534192 | 0 |
| CCGGTAGTCGCGATGC-1 | 1351426 | 44332 | 137523 | 113547 | 1056024 | 341 | 0.030735482 | 3.237693267 | 3.109921951 | 0.103296274 | 0.101006173 | 0 |
| CCGTACTAGGCTCATT-1 | 1942124 | 36088 | 199129 | 166966 | 1539941 | 497 | 0.044926167 | 4.666113567 | 3.205143868 | 0.086423123 | 0.086425008 | 0 |
| CCGTACTTCGGAGAGT-1 | 1747252 | 19932 | 185489 | 155321 | 1386510 | 448 | 0.040537741 | 4.227180567 | 2.896149474 | 0.09232074 | 0.091112967 | 0 |
| CCGTGGAAGTAGATGT-1 | 1260710 | 26687 | 128976 | 101156 | 1003891 | 324 | 0.029157273 | 3.071701167 | 3.109937085 | 0.107421282 | 0.10655571 | 0 |
| CCGTGGACACCAAAGG-1 | 946662 | 9488 | 97137 | 81474 | 758563 | 245 | 0.022118144 | 2.355058067 | 3.13896721 | 0.116664631 | 0.114339434 | 0 |
| CCGTGGACACTGGCCA-1 | 1445510 | 46353 | 144686 | 126682 | 1127789 | 364 | 0.032824053 | 3.464837833 | 2.988734515 | 0.101564215 | 0.09994302 | 0 |
| CCGTGGAGTCAACCGC-1 | 640124 | 15586 | 60323 | 57542 | 506673 | 164 | 0.014741518 | 1.580894833 | 3.042224621 | 0.145569656 | 0.147044507 | 0 |
| CCGTTCAAGTCGCCGT-1 | 2035446 | 23552 | 209026 | 181368 | 1621500 | 524 | 0.047267563 | 4.917614 | 3.142883532 | 0.087005164 | 0.085787862 | 0 |
| CCTAAAGGTACCGTGC-1 | 1243776 | 46411 | 128052 | 110346 | 958967 | 310 | 0.027888401 | 2.9471118 | 3.107868137 | 0.10849734 | 0.106254396 | 0 |
| CCTAAAGTCAAGCCGC-1 | 1537612 | 16542 | 162673 | 142891 | 1215506 | 393 | 0.035457668 | 3.734648633 | 3.037934047 | 0.097823625 | 0.096879615 | 0 |
| CCTACACAGGGATCTG-1 | 484660 | 3565 | 53439 | 41539 | 386117 | 125 | 0.011300511 | 1.215420433 | 3.029604493 | 0.164619597 | 0.165390755 | 0 |
| CCTACACCAACCGCTG-1 | 975884 | 18856 | 103875 | 86228 | 766925 | 248 | 0.022305086 | 2.3680561 | 2.987677294 | 0.123047018 | 0.119256821 | 0 |
| CCTACACGTGGAAGGG-1 | 1835666 | 14078 | 194920 | 156800 | 1469868 | 475 | 0.042932256 | 4.477480433 | 3.0814264 | 0.089731758 | 0.088296032 | 0 |
| CCTACACTCGCGATGC-1 | 1559590 | 39518 | 162674 | 146887 | 1210511 | 391 | 0.035201655 | 3.693471067 | 3.130901912 | 0.098758539 | 0.096346185 | 0 |
| CCTACACTCGTTATTC-1 | 2175604 | 42149 | 231522 | 188271 | 1713662 | 554 | 0.049824044 | 5.146113633 | 3.22477641 | 0.084354647 | 0.084077862 | 0 |
| CCTACACTCTCAGATG-1 | 963422 | 11643 | 105567 | 78705 | 767507 | 248 | 0.0223925 | 2.379014467 | 2.984660616 | 0.11636765 | 0.115234527 | 0 |
| CCTACCATCGGAGAGT-1 | 1835762 | 16302 | 197301 | 161268 | 1460891 | 472 | 0.042682893 | 4.4362076 | 2.952258825 | 0.088619568 | 0.087058738 | 0 |
| CCTAGCTCAATCCGTA-1 | 1056978 | 11035 | 113506 | 90980 | 841457 | 272 | 0.024568881 | 2.6031397 | 3.025838014 | 0.113415215 | 0.111208005 | 0 |
| CCTAGCTTCATGTCAG-1 | 1898290 | 17077 | 204875 | 163866 | 1512472 | 489 | 0.044214768 | 4.597760067 | 3.046934716 | 0.087189949 | 0.086869516 | 0 |
| CCTAGCTTCTACATGG-1 | 1041660 | 7911 | 113352 | 86255 | 834142 | 269 | 0.024321711 | 2.5681664 | 3.054034669 | 0.110805327 | 0.112811552 | 0 |
| CCTATTAAGCCAACAG-1 | 1267868 | 15635 | 136738 | 109598 | 1005897 | 325 | 0.029329422 | 3.094562467 | 3.046630358 | 0.103215933 | 0.102076079 | 0 |
| CCTATTACAACAGCCC-1 | 1252004 | 60568 | 127948 | 115999 | 947489 | 306 | 0.027562091 | 2.9239806 | 3.021051286 | 0.106883923 | 0.106883923 | 0 |
| CCTATTACACGAGTTT-1 | 1175852 | 61566 | 121358 | 96311 | 896617 | 290 | 0.026114357 | 2.762864233 | 3.08807591 | 0.108026642 | 0.110122632 | 0 |
| CCTATTAGTATTCGTG-1 | 958152 | 9371 | 101614 | 80355 | 766812 | 248 | 0.022349945 | 2.371164967 | 3.04552782 | 0.118831577 | 0.116626095 | 0 |
| CCTATTAGTGACCCAC-1 | 2067548 | 28843 | 219818 | 182638 | 1636249 | 529 | 0.04774126 | 4.954334967 | 3.019546821 | 0.088762312 | 0.088748546 | 0 |
| CCTATTATCGAACCCG-1 | 1295192 | 42853 | 131573 | 120488 | 1000278 | 323 | 0.029089953 | 3.083704033 | 2.991798739 | 0.10645367 | 0.103417402 | 0 |
| CCTCTGACATGGGCCT-1 | 1626814 | 77879 | 168080 | 152149 | 1228706 | 397 | 0.035779979 | 3.753770633 | 2.941390406 | 0.102659578 | 0.09603784 | 0 |
| CCTTACGAGTGACTCT-1 | 1754386 | 75780 | 179257 | 170786 | 1328563 | 429 | 0.038483818 | 3.9767946 | 2.934507832 | 0.104435114 | 0.098624255 | 0 |
| CCTTACGCAAGGTATA-1 | 1468810 | 11290 | 159381 | 120769 | 1177370 | 380 | 0.034396962 | 3.6057068 | 3.01705716 | 0.099685019 | 0.096888761 | 0 |
| CCTTACGCATACGTTG-1 | 1006224 | 10050 | 108505 | 80568 | 807101 | 261 | 0.023543257 | 2.486580033 | 2.897869047 | 0.116461993 | 0.114317207 | 0 |
| CCTTACGTCATCAGCA-1 | 1504350 | 42862 | 156605 | 127678 | 1177205 | 380 | 0.034321714 | 3.610882133 | 3.170590826 | 0.095350166 | 0.094845554 | 0 |
| CCTTCCCCAACCCTGG-1 | 1575962 | 12697 | 169033 | 138471 | 1255761 | 406 | 0.036654866 | 3.841571133 | 3.033778649 | 0.096032997 | 0.094423511 | 0 |
| CCTTCCCCAACGTCTA-1 | 1817908 | 43010 | 192616 | 164028 | 1418254 | 458 | 0.04134362 | 4.3208427 | 3.059488374 | 0.090809598 | 0.089121934 | 0 |
| CCTTCCCGTTCAGGTT-1 | 1694660 | 24383 | 182617 | 138091 | 1349569 | 436 | 0.039354856 | 4.112665633 | 3.071710685 | 0.092256499 | 0.090459843 | 0 |
| CCTTCCCTCGCGGACT-1 | 2050046 | 46035 | 214846 | 184208 | 1604957 | 518 | 0.046728603 | 4.845035 | 2.985304145 | 0.089090817 | 0.085770045 | 0 |
| CCTTCGAAGGTGTTAA-1 | 1401088 | 16519 | 153925 | 113399 | 1117245 | 361 | 0.032599099 | 3.415647233 | 3.127166316 | 0.112885397 | 0.109581471 | 1 |
| CCTTCGAAGTCGTCCG-1 | 1693476 | 45337 | 175468 | 152401 | 1320270 | 426 | 0.038408118 | 4.035253067 | 3.206790737 | 0.088881931 | 0.091309034 | 0 |
| CCTTCGATCAGCAGCC-1 | 1722542 | 51913 | 181552 | 156484 | 1332593 | 430 | 0.038817053 | 4.0649955 | 2.754725962 | 0.089872244 | 0.090579615 | 0 |
| CCTTCGATCAGTACCA-1 | 2142354 | 59744 | 227802 | 183099 | 1671709 | 540 | 0.048629465 | 5.0442593 | 3.082047237 | 0.084993575 | 0.083795302 | 0 |
| CGAACATAGGCTAGTG-1 | 1480506 | 13051 | 161909 | 116417 | 1189129 | 384 | 0.03471488 | 3.632922867 | 2.983017139 | 0.099890169 | 0.09806791 | 0 |
| CGAACATGTGACTGAG-1 | 1382770 | 15416 | 149996 | 134251 | 1083107 | 350 | 0.031641299 | 3.317707267 | 2.058132845 | 0.104618959 | 0.104174908 | 0 |

| | | | | | | | | | | | | |
|---|---|---|---|---|---|---|---|---|---|---|---|---|
| CGAACATTCAGTACCA-1 | 2148394 | 48823 | 228924 | 188575 | 1682072 | 543 | 0.048941104 | 5.080563433 | 3.009293535 | 0.080530416 | 0.080787058 | 0 |
| CGAACATTCTGGGATT-1 | 1519344 | 41931 | 162564 | 126803 | 1188046 | 384 | 0.0346916 | 3.635237333 | 3.143560173 | 0.097641207 | 0.095362412 | 0 |
| CGAATGTAGTACGACG-1 | 1426342 | 32493 | 148871 | 127547 | 1117431 | 361 | 0.032433703 | 3.4169298 | 2.977258841 | 0.101805633 | 0.101805633 | 0 |
| CGAATGTCAGACAGTG-1 | 1338032 | 12233 | 146804 | 104602 | 1074393 | 347 | 0.031369086 | 3.295534567 | 3.140815167 | 0.106761605 | 0.103310249 | 0 |
| CGAATGTCAGCCCATG-1 | 1143972 | 10785 | 124248 | 96891 | 912048 | 295 | 0.026593352 | 2.814320367 | 3.116965376 | 0.109629412 | 0.107721666 | 0 |
| CGAATGTGTGTCAACT-1 | 2357762 | 88055 | 245848 | 218626 | 1805233 | 583 | 0.052561526 | 5.426710667 | 3.076750382 | 0.086988275 | 0.082486644 | 0 |
| CGACCTTAGGGTAACC-1 | 1258438 | 20218 | 133529 | 112249 | 992442 | 321 | 0.028918691 | 3.051503733 | 3.143517598 | 0.108110338 | 0.105988674 | 0 |
| CGACCTTAGTGTTTGC-1 | 2188530 | 74852 | 227238 | 195990 | 1690450 | 546 | 0.04922356 | 5.1166358 | 3.045921811 | 0.083718198 | 0.084270308 | 0 |
| CGACCTTCAGTCGGAA-1 | 1778448 | 13729 | 195386 | 155289 | 1414044 | 457 | 0.041206559 | 4.304715 | 2.959141053 | 0.089443342 | 0.088539087 | 0 |
| CGACTTCGTGGTGCAA-1 | 1744136 | 17817 | 185550 | 154936 | 1385833 | 448 | 0.040444265 | 4.228205233 | 3.052149331 | 0.089513632 | 0.088980994 | 0 |
| CGAGAAGCAAACTCAC-1 | 2477078 | 40756 | 262920 | 229109 | 1944293 | 628 | 0.056719544 | 5.852171933 | 3.143746226 | 0.077793572 | 0.078285362 | 0 |
| CGAGAAGTCCGAGATT-1 | 1016972 | 34507 | 108194 | 82513 | 791758 | 256 | 0.023128642 | 2.4440609 | 3.118636725 | 0.124355243 | 0.127883459 | 1 |
| CGAGCACAGGCGATAC-1 | 2158970 | 51247 | 229803 | 179207 | 1698713 | 549 | 0.049497493 | 5.122792333 | 3.01069173 | 0.086084918 | 0.085664782 | 0 |
| CGAGCACCACGATGGA-1 | 1678868 | 17816 | 179208 | 135181 | 1346663 | 435 | 0.03930458 | 4.1065903 | 3.308262396 | 0.120854351 | 0.121043797 | 1 |
| CGAGCACCAGATATCC-1 | 1889582 | 75681 | 196412 | 161726 | 1455763 | 470 | 0.042251788 | 4.371760367 | 2.914028371 | 0.094835681 | 0.092273852 | 0 |
| CGAGCACGTCGAGTAG-1 | 343464 | 9168 | 31598 | 26752 | 275946 | 89 | 0.008033105 | 0.8666299 | 2.968632407 | 0.189519313 | 0.191629913 | 0 |
| CGAGCACTCAAGTTGC-1 | 1805596 | 38366 | 189155 | 167254 | 1410821 | 456 | 0.041085969 | 4.302524667 | 3.108889015 | 0.084895407 | 0.084895407 | 0 |
| CGAGCACTCGGAAGAC-1 | 1926266 | 72299 | 193499 | 165750 | 1494718 | 483 | 0.043576107 | 4.544577533 | 3.043908121 | 0.087129844 | 0.089049136 | 0 |
| CGAGCACTCTGATCAG-1 | 1820306 | 21160 | 193213 | 161955 | 1443978 | 466 | 0.042170325 | 4.401477767 | 2.883137912 | 0.089079906 | 0.089377631 | 0 |
| CGAGCCAAGCTGGTCC-1 | 1951710 | 61554 | 201483 | 171413 | 1517260 | 490 | 0.044033118 | 4.5694158 | 2.991084495 | 0.095186119 | 0.094264226 | 0 |
| CGAGCCAAGGCATTTC-1 | 2237266 | 65543 | 230560 | 195846 | 1745317 | 564 | 0.050321366 | 5.130891067 | 3.03029808 | 0.08997202 | 0.089707251 | 1 |
| CGAGCCACAAGAACTA-1 | 1589222 | 41654 | 165668 | 145203 | 1236697 | 399 | 0.036052537 | 3.7864152 | 3.021389956 | 0.092931724 | 0.093825847 | 0 |
| CGAGCCACAGACGAGT-1 | 2070508 | 37207 | 220476 | 183209 | 1629616 | 526 | 0.047577055 | 4.931601733 | 3.191411967 | 0.083701966 | 0.083345154 | 0 |
| CGAGCCACAGGACCAA-1 | 1804880 | 20873 | 189680 | 164570 | 1429757 | 462 | 0.041699948 | 4.358821633 | 2.986801165 | 0.088339018 | 0.08767696 | 0 |
| CGAGCCACATAGTGAA-1 | 1358980 | 14795 | 145889 | 113244 | 1085052 | 351 | 0.031617177 | 3.327787633 | 3.060361173 | 0.103306732 | 0.101395228 | 0 |
| CGATCGGGTACCGTGC-1 | 1496770 | 64643 | 153610 | 128283 | 1150234 | 372 | 0.033419882 | 3.516942167 | 3.074631953 | 0.099323335 | 0.098132113 | 0 |
| CGATGGCTCATCGCGG-1 | 2162090 | 41153 | 225029 | 179754 | 1716154 | 554 | 0.050008102 | 5.191420967 | 4.479600759 | 0.117620208 | 0.117620208 | 1 |
| CGATGGCTCGACAATC-1 | 2559804 | 49842 | 267733 | 212367 | 2029862 | 656 | 0.05906969 | 6.0417001 | 3.518475332 | 0.128309497 | 0.125134692 | 1 |
| CGATGTAAGATCGGGT-1 | 2377726 | 29888 | 252704 | 204255 | 1890879 | 611 | 0.055216544 | 5.696744833 | 3.045996859 | 0.077632274 | 0.079135459 | 0 |
| CGATGTACAAGTACAA-1 | 1296938 | 16032 | 140285 | 106126 | 1034495 | 334 | 0.030060611 | 3.162617933 | 2.997529023 | 0.108042332 | 0.103919237 | 0 |
| CGATGTAGTCGCCAAC-1 | 367326 | 3105 | 35119 | 31141 | 297961 | 96 | 0.008709902 | 0.943983233 | 3.049572577 | 0.180590162 | 0.18201443 | 0 |
| CGATGTATCTCCAGCT-1 | 561236 | 11227 | 59593 | 50085 | 440331 | 142 | 0.012860527 | 1.382100167 | 2.915619275 | 0.155885834 | 0.15310397 | 0 |
| CGATTGAAGCTAGCCC-1 | 1698570 | 83075 | 172577 | 150416 | 1292502 | 418 | 0.037606604 | 3.953399167 | 3.141614166 | 0.094070955 | 0.094066148 | 0 |
| CGATTGACACCTCCAT-1 | 1961760 | 20872 | 211484 | 172691 | 1556713 | 503 | 0.045503996 | 4.7313784 | 2.918223668 | 0.087403403 | 0.088695461 | 0 |
| CGATTGAGTAAGAAGG-1 | 1363116 | 12149 | 146058 | 113640 | 1091269 | 353 | 0.031916699 | 3.360823 | 3.003032584 | 0.095656472 | 0.095656472 | 0 |
| CGATTGAGTACTCCGG-1 | 1845060 | 24862 | 195973 | 168821 | 1455404 | 470 | 0.042477243 | 4.441960733 | 3.048004602 | 0.086656822 | 0.085607148 | 0 |
| CGATTGAGTATTCGCA-1 | 1750174 | 56348 | 183494 | 151761 | 1358571 | 439 | 0.039559769 | 4.1530741 | 3.04420019 | 0.093576335 | 0.09260972 | 0 |
| CGATTGAGTCAACTCA-1 | 1480450 | 43704 | 157932 | 136270 | 1142544 | 369 | 0.033225904 | 3.4861054 | 3.03279783 | 0.105482754 | 0.098161774 | 0 |
| CGATTGAGTGCATCTA-1 | 1754316 | 29993 | 186564 | 151898 | 1385861 | 448 | 0.04039558 | 4.220101733 | 3.034126178 | 0.092569803 | 0.090407596 | 0 |
| CGATTGAGTTGGCGTC-1 | 1516144 | 83879 | 150376 | 128771 | 1153118 | 372 | 0.033579 | 3.529186233 | 3.037356159 | 0.118217191 | 0.118217191 | 1 |
| CGATTGATCGTGCTAA-1 | 1771556 | 13391 | 193330 | 145494 | 1419341 | 458 | 0.041400225 | 4.305628633 | 3.253779398 | 0.120424673 | 0.1180887 | 1 |
| CGCCAAGAGGCTCAGA-1 | 1693462 | 13969 | 184756 | 146293 | 1348444 | 436 | 0.03937136 | 4.124616633 | 3.073836276 | 0.09362089 | 0.091428262 | 0 |
| CGCCAAGCATAGGTCT-1 | 2155770 | 72872 | 214828 | 193507 | 1674563 | 541 | 0.048089467 | 4.870143233 | 2.981287962 | 0.093425868 | 0.093165367 | 1 |
| CGCGGTAGTCGGAAGT-1 | 2052592 | 42452 | 214872 | 177329 | 1617939 | 523 | 0.04725868 | 4.9047119 | 3.109414632 | 0.085024731 | 0.083492845 | 0 |
| CGCGTTTCACAAGCTT-1 | 2137846 | 36807 | 223571 | 194298 | 1683170 | 544 | 0.049007556 | 5.1064282 | 3.139172622 | 0.080896741 | 0.080896741 | 0 |
| CGCGTTTGTTGAACTC-1 | 1216418 | 16242 | 126888 | 105461 | 967827 | 313 | 0.028258384 | 2.9822901 | 2.94057901 | 0.120874878 | 0.120874878 | 1 |
| CGCGTTTTCTGGATCA-1 | 2059390 | 67399 | 212019 | 181103 | 1598869 | 516 | 0.046534442 | 4.8442183 | 3.108150537 | 0.084128606 | 0.085803001 | 0 |
| CGCTATCAGGGATTTA-1 | 1965940 | 30304 | 209092 | 166645 | 1559899 | 504 | 0.045550926 | 4.734718067 | 3.180456095 | 0.099664876 | 0.098100239 | 1 |
| CGCTATCGTGATTGTC-1 | 2033294 | 41042 | 213930 | 169523 | 1608799 | 520 | 0.046852392 | 4.854877167 | 2.999396174 | 0.08684798 | 0.084919248 | 0 |
| CGCTATCTCCGGCTAG-1 | 1670860 | 18484 | 179356 | 148689 | 1324331 | 428 | 0.038681771 | 4.048924167 | 3.169886913 | 0.094033051 | 0.092995347 | 0 |
| CGCTATCTCGACTTCC-1 | 2075530 | 65721 | 215389 | 186045 | 1608375 | 520 | 0.046779397 | 4.850945267 | 3.153732478 | 0.087208548 | 0.087311307 | 0 |
| CGCTTCACAAGCACGA-1 | 1475008 | 15596 | 156589 | 133370 | 1169453 | 378 | 0.034099746 | 3.5946963 | 3.045859438 | 0.09753253 | 0.095879095 | 0 |
| CGCTTCACACGAAGGT-1 | 2160344 | 21364 | 232354 | 187257 | 1719369 | 555 | 0.050203306 | 5.1996286 | 2.926952803 | 0.082835828 | 0.081670113 | 0 |
| CGCTTCAGTTCTAGTG-1 | 1125858 | 10442 | 123382 | 86312 | 905722 | 293 | 0.026464776 | 2.786333567 | 3.039223483 | 0.112076048 | 0.10882621 | 0 |
| CGGACACAGACGCTTT-1 | 2279682 | 90829 | 237807 | 201293 | 1749753 | 565 | 0.05092207 | 5.282295633 | 3.181534002 | 0.084169152 | 0.08215828 | 0 |
| CGGACACGTAGTAGTA-1 | 1179800 | 13142 | 130791 | 94287 | 941580 | 304 | 0.027461848 | 2.898907 | 2.984392973 | 0.104294226 | 0.106273974 | 0 |
| CGGACACGTCTAGGTT-1 | 1425554 | 23793 | 154782 | 116280 | 1130699 | 365 | 0.03299077 | 3.464803133 | 3.107460969 | 0.097939358 | 0.098166887 | 0 |
| CGGACACTCGAGTTGT-1 | 2539928 | 39105 | 276136 | 222285 | 2002402 | 647 | 0.058415071 | 6.0041277 | 2.996910489 | 0.076561324 | 0.077960211 | 0 |
| CGGACGTCACTTGCTC-1 | 318906 | 2638 | 34223 | 27581 | 254464 | 82 | 0.007429967 | 0.806293267 | 3.066369922 | 0.200467593 | 0.20363059 | 0 |
| CGGACGTTCCTGATTT-1 | 2189420 | 72218 | 235230 | 176236 | 1705736 | 551 | 0.04955984 | 5.117037433 | 3.055690124 | 0.093311264 | 0.093171871 | 1 |
| CGGAGCTAGGCTACGA-1 | 1415898 | 9461 | 162235 | 127161 | 1117041 | 361 | 0.032548156 | 3.440116033 | 3.071341831 | 0.101466785 | 0.098245839 | 0 |
| CGGAGCTGTCCGAAGA-1 | 1349038 | 21643 | 151981 | 118411 | 1057003 | 341 | 0.030842733 | 3.255923333 | 3.104127467 | 0.101141302 | 0.100333567 | 0 |
| CGGAGCTTCTAGACAC-1 | 2244246 | 62228 | 249449 | 197465 | 1735104 | 560 | 0.050526125 | 5.2387393 | 3.108122505 | 0.081918221 | 0.079921158 | 0 |
| CGGAGTCAGAAACCTA-1 | 1116420 | 24811 | 121079 | 103571 | 866959 | 280 | 0.025275752 | 2.6820006 | 3.065763034 | 0.115097845 | 0.111812554 | 0 |
| CGGAGTCAGCATGGCA-1 | 2001692 | 53690 | 218702 | 172657 | 1556643 | 503 | 0.045329141 | 4.724172233 | 3.083985138 | 0.084180152 | 0.084710122 | 0 |
| CGGAGTCAGCCAACAG-1 | 1462388 | 23969 | 161942 | 126379 | 1150098 | 372 | 0.033492163 | 3.523158067 | 3.06412551 | 0.095297027 | 0.095686603 | 0 |
| CGGAGTCCAGAGGCAT-1 | 2891232 | 44068 | 312975 | 254950 | 2279239 | 736 | 0.066509804 | 6.800387833 | 3.184105443 | 0.074475843 | 0.074475843 | 0 |
| CGGAGTCCAGCTTGAT-1 | 3160210 | 32843 | 348492 | 266420 | 2512455 | 812 | 0.073218564 | 7.462112733 | 3.037301452 | 0.067936598 | 0.066710324 | 0 |
| CGGAGTCGTATCAGTC-1 | 1544512 | 25173 | 174847 | 132727 | 1211765 | 391 | 0.03530745 | 3.7008477 | 3.089177631 | 0.093254624 | 0.092795813 | 0 |
| CGGAGTCGTCTTCACC-1 | 1830202 | 34642 | 192569 | 169211 | 1433780 | 463 | 0.041811763 | 4.370971733 | 3.110430755 | 0.088543555 | 0.088543555 | 0 |
| CGGCTAGAGACGCACA-1 | 1764988 | 23547 | 186281 | 162522 | 1392638 | 450 | 0.040598309 | 4.249622967 | 3.010355776 | 0.088675467 | 0.08776305 | 0 |
| CGGCTAGAGTAGCCGA-1 | 1733798 | 6645 | 186940 | 161369 | 1378844 | 445 | 0.040242073 | 4.217474667 | 3.073471446 | 0.093730481 | 0.091590753 | 0 |
| CGGCTAGAGTAGTATG-1 | 1272664 | 12240 | 138067 | 97388 | 1024969 | 331 | 0.02992082 | 3.145150067 | 2.997776625 | 0.105874965 | 0.105795625 | 0 |
| CGGCTAGGTGTCAACT-1 | 2996998 | 67147 | 318703 | 266036 | 2345112 | 758 | 0.068394622 | 7.007945833 | 2.93081618 | 0.072745607 | 0.073044926 | 0 |
| CGGCTAGGTTCTTCGC-1 | 1976848 | 33868 | 211107 | 178245 | 1553628 | 502 | 0.045324104 | 4.7249509 | 2.978735951 | 0.0914803 | 0.091413088 | 0 |
| CGGCTAGTCATGTGAC-1 | 2053004 | 56203 | 215644 | 178289 | 1602868 | 518 | 0.046691605 | 4.856228767 | 3.069134996 | 0.084791659 | 0.083939045 | 0 |
| CGGGTCAGTCATTACG-1 | 1552808 | 31640 | 163699 | 136234 | 1221235 | 394 | 0.035482946 | 3.724395533 | 3.005420381 | 0.097765404 | 0.096501275 | 0 |
| CGGTTAACACCACATA-1 | 1270688 | 10741 | 141264 | 104608 | 1014075 | 328 | 0.029588975 | 3.1140416 | 3.13241539 | 0.105146632 | 0.103621209 | 0 |
| CGGTTAACATGGAGAC-1 | 1498302 | 104708 | 151215 | 128010 | 1114369 | 360 | 0.032419558 | 3.4171267 | 2.997954705 | 0.098773103 | 0.09959121 | 0 |
| CGTAGCGAGGACAGAA-1 | 1180076 | 15168 | 121516 | 98800 | 944592 | 305 | 0.027531339 | 2.905745867 | 3.033501218 | 0.105040508 | 0.106026699 | 0 |
| CGTAGCGAGTACGATA-1 | 1236012 | 12302 | 131506 | 102018 | 990186 | 320 | 0.028847102 | 3.027583 | 3.193678051 | 0.09882375 | 0.09882375 | 0 |
| CGTAGCGCAACGGGAT-1 | 1587150 | 17055 | 160982 | 133511 | 1275602 | 412 | 0.037296799 | 3.8849361 | 3.003189487 | 0.100662729 | 0.097948111 | 0 |
| CGTAGGCAGGCGTTGA-1 | 1084734 | 16901 | 104742 | 95325 | 867766 | 280 | 0.025317433 | 2.689115967 | 3.04192229 | 0.115174428 | 0.109683775 | 0 |
| CGTAGGCCACTTGTTT-1 | 1328928 | 34264 | 121929 | 110305 | 1062430 | 343 | 0.030967144 | 3.246652733 | 3.009145844 | 0.103872194 | 0.102256919 | 0 |
| CGTAGGCCATGAGGGT-1 | 1709230 | 30056 | 159809 | 148095 | 1371270 | 443 | 0.040037392 | 4.1726686 | 3.041909064 | 0.092343454 | 0.091042365 | 0 |
| CGTAGGCGTTGAACAA-1 | 1527450 | 27372 | 153095 | 128221 | 1218762 | 394 | 0.035540104 | 3.723282767 | 3.560688425 | 0.137336532 | 0.137336532 | 1 |
| CGTCACTAGGCGCTCT-1 | 1903772 | 53819 | 189760 | 164639 | 1495554 | 483 | 0.043463025 | 4.534249733 | 3.074450025 | 0.091774458 | 0.090285576 | 0 |
| CGTCACTAGTCCCACG-1 | 1526544 | 20560 | 158092 | 137905 | 1209987 | 391 | 0.035189623 | 3.7037983 | 3.059660786 | 0.096177632 | 0.095894489 | 0 |
| CGTCACTCATTCCGAA-1 | 1515310 | 15582 | 161060 | 128295 | 1210373 | 391 | 0.035277863 | 3.7071591 | 3.047565947 | 0.095296598 | 0.092106725 | 0 |
| CGTCACTTCGTTACCC-1 | 2000396 | 22121 | 212880 | 176879 | 1588516 | 513 | 0.046236181 | 4.810863767 | 3.1446212 | 0.082039367 | 0.081452631 | 0 |
| CGTCACTTCTTAGCTT-1 | 1458070 | 84081 | 147266 | 121206 | 1105517 | 357 | 0.032158885 | 3.3604032 | 3.015565612 | 0.140675295 | 0.140675295 | 1 |
| CGTCAGGAGCAGGCTA-1 | 1527678 | 32382 | 154354 | 135539 | 1205403 | 389 | 0.03518498 | 3.698296533 | 2.940615435 | 0.096113874 | 0.092637486 | 0 |
| CGTCAGGAGGACAGCT-1 | 1527512 | 42237 | 153792 | 140363 | 1191120 | 385 | 0.034703085 | 3.657480367 | 2.998110702 | 0.095648262 | 0.095308049 | 0 |
| CGTCAGGCACGAGCTC-1 | 1277054 | 45275 | 130755 | 114934 | 986090 | 319 | 0.028722727 | 3.038112133 | 3.046093095 | 0.104993677 | 0.103600498 | 0 |

| | | | | | | | | | | | | |
|---|---|---|---|---|---|---|---|---|---|---|---|---|
| CGTCCATGTAGATGTA-1 | 856004 | 11200 | 86610 | 68326 | 689868 | 223 | 0.020070448 | 2.129878933 | 2.998870859 | 0.11940271 | 0.121479744 | 0 |
| CGTCCATTCTCCACAC-1 | 2261368 | 58891 | 236159 | 191907 | 1774411 | 573 | 0.051684896 | 5.3540041 | 3.011403602 | 0.085501897 | 0.081899965 | 0 |
| CGTCTACCAACGACTT-1 | 1810266 | 37553 | 186456 | 160975 | 1425282 | 460 | 0.041445643 | 4.328347767 | 3.098777186 | 0.091330796 | 0.091093904 | 0 |
| CGTCTACCAAGACGAC-1 | 1287782 | 20438 | 133208 | 108240 | 1025896 | 331 | 0.029991754 | 3.1611919 | 2.641412132 | 0.13897844 | 0.137894809 | 1 |
| CGTCTACCACAGCTGC-1 | 1566836 | 50419 | 153256 | 137943 | 1225218 | 396 | 0.035689298 | 3.7490021 | 3.265588931 | 0.0959431 | 0.0959431 | 0 |
| CGTCTACGTGGAGCTA-1 | 1612578 | 48519 | 163870 | 141223 | 1258966 | 407 | 0.036718456 | 3.857782 | 3.099096219 | 0.09586259 | 0.094173526 | 0 |
| CGTCTACTCTTAAGGC-1 | 1624182 | 57473 | 166136 | 142202 | 1258371 | 406 | 0.036640768 | 3.842845833 | 3.249960938 | 0.095563647 | 0.095563647 | 0 |
| CGTGAGCTCGAATGTC-1 | 1474586 | 28227 | 147809 | 128549 | 1170001 | 378 | 0.034089169 | 3.5751218 | 3.02306425 | 0.099664807 | 0.099664807 | 0 |
| CGTGTAAAGTTATCGC-1 | 1296870 | 32280 | 116722 | 106930 | 1040938 | 336 | 0.030340915 | 3.188387233 | 4.040878962 | 0.150169026 | 0.150169026 | 1 |
| CGTGTAAGTAGGTTTC-1 | 1200112 | 49514 | 104926 | 107762 | 937910 | 303 | 0.027271992 | 2.882236067 | 3.153406428 | 0.106159608 | 0.106159608 | 0 |
| CGTGTAAGTTTCCGAA-1 | 1069042 | 18652 | 103870 | 90994 | 855526 | 276 | 0.024903713 | 2.633148633 | 3.001398929 | 0.112026035 | 0.11223839 | 0 |
| CGTGTCTCAACCGCTG-1 | 561062 | 14089 | 50039 | 49135 | 447799 | 145 | 0.013013182 | 1.398346867 | 3.007562041 | 0.151918899 | 0.146535128 | 0 |
| CGTGTCTCACAGCTAT-1 | 1019540 | 18698 | 94079 | 88603 | 818160 | 264 | 0.023803447 | 2.520560033 | 3.089156233 | 0.114906707 | 0.111612645 | 0 |
| CGTGTCTCATCCATCC-1 | 1065860 | 58156 | 92685 | 94804 | 820215 | 265 | 0.023782541 | 2.526501767 | 3.107425869 | 0.113917663 | 0.114738243 | 0 |
| CGTGTCTTCCAACAAC-1 | 1304530 | 21459 | 122330 | 112239 | 1048502 | 339 | 0.030528301 | 3.212134233 | 2.888343654 | 0.103957743 | 0.102364398 | 0 |
| CGTGTCTTCCGAGTCG-1 | 758556 | 22010 | 67223 | 69933 | 599390 | 194 | 0.017359875 | 1.862627067 | 3.125389712 | 0.130734206 | 0.125173532 | 0 |
| CGTGTCTTCGTTGCAA-1 | 1085152 | 11904 | 101242 | 93296 | 878710 | 284 | 0.025598769 | 2.711610467 | 3.099141329 | 0.113507578 | 0.111478881 | 0 |
| CGTGTCTTCTGGGACC-1 | 1249184 | 32063 | 114553 | 112388 | 990180 | 320 | 0.028818333 | 3.051273033 | 2.971897704 | 0.113199887 | 0.112806338 | 1 |
| CGTTAGAAGGGTAACC-1 | 1962970 | 34043 | 201856 | 179821 | 1547250 | 500 | 0.04502531 | 4.668406767 | 3.110819067 | 0.089901162 | 0.088631057 | 0 |
| CGTTAGAAGTGGCACA-1 | 1747090 | 32014 | 183794 | 161003 | 1370279 | 443 | 0.0399684 | 4.187541567 | 3.047082892 | 0.088000853 | 0.085504199 | 0 |
| CGTTAGAGTCATCCAA-1 | 1819246 | 20179 | 191152 | 158861 | 1449054 | 468 | 0.042312736 | 4.419032333 | 3.140867082 | 0.089852713 | 0.087021225 | 0 |
| CGTTAGATCAGCCTGG-1 | 1890494 | 22916 | 198496 | 167954 | 1501128 | 485 | 0.043797465 | 4.569540367 | 3.108940929 | 0.087506704 | 0.088208283 | 0 |
| CGTTCTGAGCTGCGAA-1 | 1713214 | 22873 | 179651 | 147780 | 1362910 | 440 | 0.039729403 | 4.155219667 | 3.038428754 | 0.093552412 | 0.090640457 | 0 |
| CGTTCTGAGGCTAGCA-1 | 1593886 | 57906 | 161829 | 139414 | 1234737 | 399 | 0.035925359 | 3.7787763 | 3.060055261 | 0.096239458 | 0.095846518 | 0 |
| CGTTCTGGTTGAACAA-1 | 1553610 | 15878 | 165343 | 135072 | 1237317 | 400 | 0.036092728 | 3.7966688 | 3.109311396 | 0.09686683 | 0.094260737 | 0 |
| CGTTCTGTCTCTAGTT-1 | 1345264 | 23410 | 143652 | 108747 | 1069455 | 345 | 0.031158203 | 3.276483967 | 3.14158695 | 0.106424549 | 0.099919684 | 0 |
| CGTTGGGAGGCGCTCT-1 | 1660516 | 89504 | 159156 | 143097 | 1268759 | 410 | 0.036880549 | 3.866992733 | 3.143487185 | 0.095359577 | 0.095326369 | 0 |
| CGTTGGGAGTCCCACG-1 | 1886736 | 12994 | 198184 | 170367 | 1505191 | 486 | 0.043781336 | 4.577657933 | 3.025501325 | 0.089755476 | 0.086078881 | 0 |
| CTAACTTAGGGATGGG-1 | 2390956 | 16866 | 253506 | 215612 | 1904972 | 615 | 0.055700603 | 5.7358368 | 3.156351468 | 0.077399607 | 0.076847069 | 0 |
| CTAACTTGTTGACGGA-1 | 1475372 | 36077 | 157947 | 128908 | 1152440 | 372 | 0.033596678 | 3.536945633 | 3.144316721 | 0.101394701 | 0.098694754 | 0 |
| CTAACTTTCTTGGCCT-1 | 2119528 | 91541 | 218779 | 190476 | 1618732 | 523 | 0.047067813 | 4.898149867 | 3.08417695 | 0.092322576 | 0.090279252 | 0 |
| CTAAGACGTGATTGTC-1 | 2081728 | 32821 | 224345 | 175754 | 1648808 | 533 | 0.048068205 | 4.977690467 | 3.109839362 | 0.086901226 | 0.086963097 | 0 |
| CTAATGGCAACCTGAT-1 | 1718314 | 17066 | 186825 | 142959 | 1371464 | 443 | 0.04003356 | 4.170205033 | 3.042408574 | 0.091785458 | 0.092584103 | 0 |
| CTACACCAGTTACCCA-1 | 1958588 | 60403 | 206718 | 167964 | 1523503 | 492 | 0.044354111 | 4.629236667 | 3.016740651 | 0.088999901 | 0.085584082 | 0 |
| CTACACCGTACCGTGC-1 | 1555884 | 42826 | 162026 | 136786 | 1214246 | 392 | 0.035303895 | 3.707090933 | 3.119404151 | 0.097042807 | 0.096598519 | 0 |
| CTACATTCACGAGATG-1 | 1003378 | 9074 | 109793 | 83999 | 800512 | 259 | 0.023417138 | 2.4778293 | 3.167192048 | 0.118303211 | 0.116576924 | 0 |
| CTACATTTCAAAGTGA-1 | 1559394 | 18679 | 168518 | 138887 | 1233310 | 398 | 0.035985596 | 3.7766569 | 3.044849165 | 0.09820599 | 0.096442306 | 0 |
| CTACATTTCACGTTAG-1 | 1528406 | 9107 | 168006 | 126221 | 1225072 | 396 | 0.0357958 | 3.746199633 | 3.088261197 | 0.096814348 | 0.09699749 | 0 |
| CTACCCACAGTAGGGT-1 | 2157756 | 16509 | 234716 | 185599 | 1720932 | 556 | 0.050253513 | 5.1932489 | 3.052444621 | 0.085181507 | 0.083930582 | 0 |
| CTACCCAGTAGAACAT-1 | 2353070 | 36677 | 254140 | 209280 | 1852973 | 599 | 0.054068063 | 5.583048233 | 3.107911932 | 0.079946022 | 0.07912283 | 0 |
| CTACCCAGTATATGAG-1 | 1089474 | 9226 | 119248 | 86667 | 874333 | 282 | 0.025510375 | 2.687537233 | 3.073386936 | 0.110560528 | 0.110309541 | 0 |
| CTACGTCGTAGTGGCA-1 | 1545156 | 55036 | 148526 | 137100 | 1204494 | 389 | 0.03509039 | 3.695667833 | 3.009249853 | 0.095477954 | 0.096567373 | 0 |
| CTAGAGTTCGACAATC-1 | 2030534 | 43385 | 213719 | 176129 | 1597301 | 516 | 0.046504562 | 4.818978933 | 3.095566873 | 0.090475805 | 0.087801758 | 0 |
| CTAGCCTAGGGATACC-1 | 3113612 | 45134 | 329328 | 274740 | 2464410 | 796 | 0.071582578 | 7.2100198 | 3.064339202 | 0.076610104 | 0.075563579 | 0 |
| CTAGCCTTCCAACGAT-1 | 2121236 | 32899 | 230296 | 186094 | 1671947 | 540 | 0.04879306 | 5.057247533 | 2.921123859 | 0.084012584 | 0.081305627 | 0 |
| CTAGCCTTCTCCGGAG-1 | 1832240 | 19858 | 196106 | 159233 | 1457043 | 471 | 0.042484065 | 4.438456967 | 3.054795007 | 0.095222995 | 0.091999596 | 0 |
| CTAGCCTTCTTGGATG-1 | 940192 | 12741 | 102392 | 78371 | 746688 | 241 | 0.02176956 | 2.317985367 | 2.969414204 | 0.124466028 | 0.120055896 | 0 |
| CTAGCCTTCTTGTCCG-1 | 1383720 | 25384 | 146577 | 130856 | 1080903 | 349 | 0.031498144 | 3.3335749 | 3.145155676 | 0.098591508 | 0.101380616 | 0 |
| CTAGTGACAGTTTGCA-1 | 1906118 | 55236 | 199053 | 179562 | 1472267 | 476 | 0.042893913 | 4.493857567 | 3.01199838 | 0.086045215 | 0.086229583 | 0 |
| CTAGTGAGTAGGGTGT-1 | 1960660 | 29613 | 206076 | 170498 | 1554473 | 502 | 0.045458609 | 4.7179248 | 3.10617319 | 0.090680571 | 0.088180024 | 0 |
| CTAGTGATCTTAAGGC-1 | 2560254 | 62047 | 268051 | 211049 | 2019107 | 652 | 0.058867817 | 6.0466642 | 2.697471764 | 0.116611617 | 0.116611617 | 1 |
| CTCACACCATAGGTCT-1 | 327056 | 2826 | 34877 | 28125 | 261228 | 84 | 0.007624999 | 0.826155533 | 3.06905389 | 0.195228871 | 0.199457096 | 0 |
| CTCACACGTCTCTCTG-1 | 1495918 | 20166 | 160517 | 128269 | 1186966 | 383 | 0.034656715 | 3.646884433 | 3.036616897 | 0.094412411 | 0.094875573 | 0 |
| CTCACACTCAAGAGGC-1 | 2071306 | 54424 | 219228 | 182541 | 1615113 | 522 | 0.04706732 | 4.891227633 | 3.034239034 | 0.083363121 | 0.084712562 | 0 |
| CTCAGAACATGGAGAC-1 | 1688592 | 65014 | 176089 | 153020 | 1294469 | 418 | 0.03767659 | 3.956617167 | 3.096021597 | 0.091829391 | 0.093110915 | 0 |
| CTCAGAATCATGACCA-1 | 2000786 | 51382 | 212940 | 176225 | 1560239 | 504 | 0.04536542 | 4.7227471 | 3.011010478 | 0.086531912 | 0.086037822 | 0 |
| CTCATTAGTGTTCCGG-1 | 2253188 | 20534 | 241162 | 208554 | 1782938 | 576 | 0.052034523 | 5.4071209 | 3.2069284 | 0.083426564 | 0.081867434 | 0 |
| CTCATTATCATTGGAC-1 | 1463202 | 35485 | 145228 | 131485 | 1151004 | 372 | 0.033488103 | 3.5243579 | 3.019333348 | 0.104325116 | 0.103806784 | 1 |
| CTCATTATCTGATCAG-1 | 1501058 | 11591 | 160293 | 133403 | 1195771 | 386 | 0.0349011 | 3.653897233 | 2.758148853 | 0.093801825 | 0.093801825 | 0 |
| CTCCTAGAGGAGTACC-1 | 2763282 | 28714 | 299137 | 250751 | 2184680 | 706 | 0.063662812 | 6.5240663 | 3.035823885 | 0.078652034 | 0.076497298 | 0 |
| CTCCTAGCAAACCGAG-1 | 1563740 | 15633 | 166675 | 136542 | 1244890 | 402 | 0.036289342 | 3.811587767 | 3.06562662 | 0.097550723 | 0.095756039 | 0 |
| CTCGAAAAGCAGGTCA-1 | 2285172 | 54679 | 236644 | 201940 | 1791909 | 579 | 0.052162523 | 5.3902435 | 3.064757859 | 0.085205805 | 0.085943593 | 0 |
| CTCGAAACAAGCTACT-1 | 1975938 | 47327 | 211690 | 179274 | 1537647 | 497 | 0.044849458 | 4.675464833 | 3.035554482 | 0.085418231 | 0.085903952 | 0 |
| CTCGAAACATTTGTGG-1 | 1583110 | 20931 | 171427 | 141419 | 1249333 | 404 | 0.03638018 | 3.817449667 | 3.175836429 | 0.095546106 | 0.096686208 | 0 |
| CTCGAAATCTACAAGC-1 | 2286928 | 55356 | 244776 | 201887 | 1784909 | 577 | 0.051973004 | 5.382899333 | 3.104763055 | 0.078236923 | 0.079604333 | 0 |
| CTCGAGGAGCGATATA-1 | 1127428 | 11930 | 120783 | 92320 | 902395 | 292 | 0.026336249 | 2.7811031 | 3.168086677 | 0.110343678 | 0.107689797 | 0 |
| CTCGAGGAGTGACATA-1 | 1450604 | 15001 | 155857 | 123205 | 1156541 | 374 | 0.033745701 | 3.546709667 | 3.089834096 | 0.101246607 | 0.097694343 | 0 |
| CTCGAGGCAACTAGTC-1 | 1256054 | 17485 | 137605 | 103090 | 997874 | 322 | 0.029113872 | 3.065941367 | 3.007549019 | 0.103258312 | 0.099868202 | 0 |
| CTCGAGGCAAGAATCA-1 | 1917166 | 73487 | 197937 | 170916 | 1474826 | 476 | 0.042934146 | 4.477838733 | 3.036442165 | 0.088850377 | 0.088620471 | 0 |
| CTCGAGGCAAGTGTTC-1 | 2530396 | 83816 | 263584 | 222531 | 1960465 | 633 | 0.056914471 | 5.841298967 | 3.110339046 | 0.080418918 | 0.079721384 | 0 |
| CTCGAGGGTCATGTTG-1 | 1208110 | 12446 | 128280 | 94304 | 973080 | 314 | 0.028419392 | 2.9924117 | 3.052217393 | 0.108598181 | 0.107906959 | 0 |
| CTCGGAGAGAGCAATT-1 | 1380234 | 42952 | 146410 | 113540 | 1077332 | 348 | 0.031410026 | 3.3020091 | 3.079942893 | 0.101833668 | 0.102480522 | 0 |
| CTCGGAGAGCTAACAA-1 | 1148382 | 16213 | 123609 | 97643 | 910917 | 294 | 0.026510858 | 2.8047236 | 2.976884072 | 0.107621346 | 0.105815703 | 0 |
| CTCGGAGTCCACTAAG-1 | 1127786 | 10427 | 124394 | 91174 | 901791 | 291 | 0.026230658 | 2.763890367 | 3.028395938 | 0.108325242 | 0.109467182 | 0 |
| CTCGGGAAGTGTTAGA-1 | 1086558 | 14814 | 114747 | 91561 | 865436 | 280 | 0.025334325 | 2.682456533 | 3.099958105 | 0.119023147 | 0.11721102 | 0 |
| CTCGGGACAAGGGTGT-1 | 1852084 | 29013 | 183354 | 167900 | 1471817 | 475 | 0.042941982 | 4.454483367 | 2.970665739 | 0.094505531 | 0.090080585 | 0 |
| CTCGGGATCATATCTC-1 | 2347498 | 43649 | 241693 | 215304 | 1846852 | 597 | 0.053714975 | 5.537878133 | 3.079866439 | 0.081804424 | 0.081587434 | 0 |
| CTCGTACAGAGCTTCT-1 | 1707842 | 61590 | 168877 | 151166 | 1326209 | 428 | 0.038592984 | 4.0413505 | 3.106658429 | 0.093188539 | 0.093408881 | 0 |
| CTCGTACAGCTGCGAA-1 | 1506866 | 13044 | 159950 | 132667 | 1201205 | 388 | 0.035013864 | 3.672032133 | 3.017811488 | 0.096144591 | 0.092721417 | 0 |
| CTCGTACCACGTACAT-1 | 1392466 | 11753 | 149129 | 120273 | 1111311 | 359 | 0.032390161 | 3.4060444 | 3.011575882 | 0.100579748 | 0.098989968 | 0 |
| CTCGTACCACGTTTAG-1 | 1017728 | 7727 | 108943 | 86717 | 814341 | 263 | 0.023805404 | 2.528387833 | 3.020280922 | 0.11094554 | 0.109822986 | 0 |
| CTCGTACCATGCCATA-1 | 1401664 | 13665 | 147200 | 116738 | 1124061 | 363 | 0.03280752 | 3.451266767 | 2.98432918 | 0.102711589 | 0.101284571 | 0 |
| CTCGTACGTTGCCTCT-1 | 1926940 | 41840 | 200068 | 170757 | 1514275 | 489 | 0.044026277 | 4.5806802 | 3.119848538 | 0.08879369 | 0.089327653 | 0 |
| CTCGTCAAGCGGCTTC-1 | 1937546 | 45395 | 197517 | 180330 | 1514304 | 489 | 0.044134374 | 4.6051607 | 3.014394623 | 0.084633441 | 0.084402212 | 0 |
| CTCGTCACAACGACGA-1 | 1169706 | 7971 | 126257 | 105027 | 930451 | 301 | 0.027140094 | 2.8762414 | 2.997755921 | 0.109017193 | 0.108069849 | 0 |
| CTCGTCACAAGGTGAC-1 | 2379404 | 27065 | 261445 | 219291 | 1871603 | 605 | 0.054468836 | 5.641332567 | 3.059064685 | 0.083036265 | 0.082664522 | 0 |
| CTCGTCAGTGTCCCTT-1 | 2062898 | 49037 | 214362 | 183490 | 1616009 | 522 | 0.047117519 | 4.887136767 | 3.046675192 | 0.086456425 | 0.086336405 | 0 |
| CTCGTCATCACTGCGG-1 | 1832716 | 17204 | 193852 | 164337 | 1457323 | 471 | 0.042481094 | 4.437792867 | 3.007473024 | 0.086396807 | 0.086858796 | 0 |
| CTCTAATGTATCGCCG-1 | 900322 | 38672 | 89970 | 80783 | 690897 | 223 | 0.020044247 | 2.138781 | 3.032719372 | 0.124292118 | 0.123826826 | 0 |
| CTCTAATTCATATCTC-1 | 2380144 | 30615 | 247426 | 215230 | 1886873 | 610 | 0.054812533 | 5.6182659 | 3.166246368 | 0.084554504 | 0.084554504 | 0 |
| CTCTACGAGAAAGTGG-1 | 1638324 | 13021 | 174957 | 142413 | 1307933 | 423 | 0.038199491 | 4.002699933 | 3.084457924 | 0.093192981 | 0.093673781 | 0 |
| CTCTACGGTGATTGGG-1 | 2120548 | 22138 | 224005 | 186674 | 1687731 | 545 | 0.04928551 | 5.125126633 | 3.089007765 | 0.082912574 | 0.082912574 | 0 |
| CTGAAGTAGTGATTGA-1 | 443610 | 2809 | 47634 | 37859 | 355308 | 115 | 0.010414924 | 1.123269433 | 3.084361132 | 0.171023474 | 0.170009204 | 0 |
| CTGAAGTCAGACTTGT-1 | 596420 | 4356 | 64716 | 51746 | 475602 | 154 | 0.013910614 | 1.4905837 | 3.1104633 | 0.160810591 | 0.158074567 | 0 |

| | | | | | | | | | | | | |
|---|---|---|---|---|---|---|---|---|---|---|---|---|
| CTGAAGTGTAACGGCA-1 | 1902978 | 72754 | 199621 | 165034 | 1465569 | 473 | 0.042662696 | 4.457422533 | 3.081537379 | 0.088519466 | 0.089200565 | 0 |
| CTGAAGTGTCAACCGC-1 | 1320288 | 38113 | 138178 | 120671 | 1023326 | 331 | 0.029790664 | 3.146162267 | 2.984300666 | 0.106719634 | 0.105070744 | 0 |
| CTGAAGTGTCCCACAG-1 | 1428234 | 13240 | 156141 | 122560 | 1136293 | 367 | 0.033062068 | 3.473943067 | 3.17162239 | 0.101202051 | 0.099015825 | 0 |
| CTGAAGTTCGTAGCGC-1 | 1367116 | 24192 | 141288 | 126217 | 1075419 | 347 | 0.031325224 | 3.317167833 | 3.088183083 | 0.102826412 | 0.102826412 | 0 |
| CTGATAGAGCAGCGTA-1 | 1181584 | 14862 | 124325 | 103446 | 938951 | 303 | 0.027436522 | 2.9010493 | 3.134874869 | 0.106246545 | 0.104990891 | 0 |
| CTGATAGCAGCTCATA-1 | 1553248 | 14719 | 167225 | 135488 | 1235816 | 399 | 0.036078473 | 3.7733072 | 2.995929241 | 0.092266373 | 0.092674369 | 0 |
| CTGATAGTCAGAAACA-1 | 1537562 | 38305 | 161344 | 145742 | 1192171 | 385 | 0.034774663 | 3.652282733 | 3.058908771 | 0.101834517 | 0.095929179 | 0 |
| CTGATCCGTTCGTCAA-1 | 1696836 | 15592 | 181047 | 153512 | 1346685 | 435 | 0.039304368 | 4.1021824 | 3.02087521 | 0.095379332 | 0.092068477 | 0 |
| CTGCCTAGTGGTCAGA-1 | 1938248 | 12978 | 206739 | 181199 | 1537332 | 497 | 0.044861043 | 4.680603867 | 3.11474676 | 0.089646995 | 0.088526282 | 0 |
| CTGCCTAGTTCCACAA-1 | 1323522 | 10719 | 145921 | 113899 | 1052983 | 340 | 0.030751861 | 3.241486 | 2.771150322 | 0.101382131 | 0.101653732 | 0 |
| CTGCCTATCGTAGCGC-1 | 1402058 | 17933 | 145451 | 131879 | 1106795 | 358 | 0.032211239 | 3.391182033 | 3.045507493 | 0.104149384 | 0.101045308 | 0 |
| CTGCGGAAGAGTGACC-1 | 2253888 | 39985 | 228552 | 206359 | 1778992 | 575 | 0.051873029 | 5.364512233 | 3.195462491 | 0.082609261 | 0.081208457 | 0 |
| CTGCGGAAGATGTCTC-1 | 2097292 | 43793 | 213585 | 188166 | 1651748 | 534 | 0.04812277 | 4.988983433 | 3.143177814 | 0.082486877 | 0.083263919 | 0 |
| CTGCGGAAGTTGTCGT-1 | 1437632 | 35351 | 140076 | 127987 | 1134218 | 366 | 0.033089927 | 3.482450233 | 2.971983325 | 0.102500698 | 0.100634521 | 0 |
| CTGCGGAGTCTCCACT-1 | 2302124 | 43820 | 242140 | 213695 | 1802469 | 582 | 0.052483142 | 5.415455567 | 3.144684962 | 0.085789383 | 0.084892361 | 0 |
| CTGCGGATCAGAAACA-1 | 1794346 | 50036 | 187028 | 164039 | 1393243 | 450 | 0.040634764 | 4.257108267 | 2.913233999 | 0.090116101 | 0.090116101 | 0 |
| CTGCGGATCCAAAGAA-1 | 1294600 | 23917 | 137384 | 112916 | 1020383 | 330 | 0.029718838 | 3.1313968 | 3.083663612 | 0.109444621 | 0.104363896 | 0 |
| CTGCGGATCCCAGAAT-1 | 2037080 | 31364 | 217101 | 176835 | 1611780 | 521 | 0.047007967 | 4.860889233 | 3.050676612 | 0.084439999 | 0.084115103 | 0 |
| CTGCTGTAGTAGGCCA-1 | 2904584 | 46632 | 307361 | 257150 | 2293441 | 741 | 0.06690233 | 6.878235 | 3.04479817 | 0.070958185 | 0.070842514 | 0 |
| CTGCTGTAGTCTCGGC-1 | 1587634 | 46644 | 166037 | 143424 | 1231529 | 398 | 0.035867426 | 3.781468433 | 3.016763269 | 0.095915563 | 0.096778388 | 0 |
| CTGCTGTCAACATAAG-1 | 999008 | 8249 | 108803 | 78317 | 803639 | 260 | 0.023518168 | 2.480268167 | 3.01775837 | 0.115024183 | 0.113822096 | 0 |
| CTGGTCTAGCGATATA-1 | 1369272 | 12013 | 147479 | 111928 | 1097852 | 355 | 0.03203925 | 3.360740733 | 2.985688107 | 0.104414385 | 0.103164377 | 0 |
| CTGGTCTAGGTCGGAT-1 | 2144268 | 25502 | 228582 | 180198 | 1709986 | 552 | 0.049937113 | 5.168721 | 2.936170364 | 0.085501076 | 0.083118416 | 0 |
| CTGGTCTTCCTGATTT-1 | 2043226 | 73879 | 214536 | 164843 | 1589968 | 514 | 0.046226042 | 4.782915867 | 3.22314678 | 0.118126368 | 0.120136739 | 1 |
| CTGGTCTTCTCCCTAG-1 | 1760714 | 10573 | 189650 | 145661 | 1414830 | 457 | 0.041297216 | 4.306031767 | 2.827139368 | 0.090564736 | 0.088004571 | 0 |
| CTGTGCTCAATGGGAC-1 | 1792334 | 85480 | 181386 | 157408 | 1368060 | 442 | 0.039808665 | 4.174099333 | 3.045392853 | 0.092743723 | 0.093281284 | 0 |
| CTGTGCTCAGACACGA-1 | 1560768 | 12728 | 167697 | 138184 | 1242159 | 401 | 0.03619035 | 3.810207633 | 3.000028408 | 0.096595105 | 0.09340183 | 0 |
| CTGTGCTCATGGCTCG-1 | 1031146 | 31848 | 103607 | 95599 | 800092 | 258 | 0.02321614 | 2.4718329 | 3.014880213 | 0.115128148 | 0.113494355 | 0 |
| CTGTGCTGTGGGCCTT-1 | 1782228 | 35880 | 185357 | 163305 | 1397686 | 451 | 0.040753745 | 4.260388433 | 3.005888955 | 0.092581503 | 0.089405956 | 0 |
| CTGTTTAAGAGTCTGG-1 | 2009608 | 17905 | 213162 | 177429 | 1601112 | 517 | 0.046778974 | 4.8758432 | 3.088770973 | 0.082092232 | 0.08255897 | 0 |
| CTGTTTAAGCTGCCCA-1 | 1733476 | 40875 | 181142 | 158901 | 1352558 | 437 | 0.039452659 | 4.1385167 | 3.141778428 | 0.093901957 | 0.093999334 | 0 |
| CTGTTTAAGGCTATCT-1 | 2211484 | 53041 | 233677 | 191273 | 1733493 | 560 | 0.050345876 | 5.187961733 | 3.109609866 | 0.091706718 | 0.087791857 | 0 |
| CTGTTTACACGCCTGA-1 | 1466792 | 16859 | 157859 | 129248 | 1162826 | 376 | 0.033941439 | 3.567886333 | 3.018909473 | 0.099649224 | 0.097774395 | 0 |
| CTGTTTAGTCTCTTTA-1 | 1054114 | 14082 | 112038 | 92653 | 835341 | 270 | 0.024443221 | 2.586924333 | 2.917447086 | 0.110020654 | 0.110018126 | 0 |
| CTGTTTATCACCAGCG-1 | 600228 | 27944 | 61771 | 54027 | 456486 | 147 | 0.01319896 | 1.413432567 | 2.944189613 | 0.147719573 | 0.143954214 | 0 |
| CTTAACTAGCATCTTG-1 | 797650 | 8617 | 86789 | 67259 | 634985 | 205 | 0.018538829 | 1.971474667 | 2.983313688 | 0.125362199 | 0.125749205 | 0 |
| CTTAACTGTGGAAAGA-1 | 1335630 | 21708 | 141310 | 117150 | 1055462 | 341 | 0.030852233 | 3.258244967 | 3.072816361 | 0.100825335 | 0.101093223 | 0 |
| CTTAACTTCCTTGCAC-1 | 2206996 | 52103 | 237378 | 194151 | 1723364 | 557 | 0.050213775 | 5.2020779 | 2.9041598 | 0.089730691 | 0.086152171 | 0 |
| CTTACCGAGGGCTTCC-1 | 2357294 | 65991 | 245239 | 214025 | 1832039 | 592 | 0.053331553 | 5.514133267 | 2.895529067 | 0.083993979 | 0.083993979 | 0 |
| CTTACCGGTAAGTAAC-1 | 2149550 | 18786 | 229482 | 194908 | 1706374 | 551 | 0.049816621 | 5.164468867 | 3.119499893 | 0.081608867 | 0.081173409 | 0 |
| CTTACCGGTCACCTGG-1 | 1495446 | 16363 | 158509 | 131671 | 1188903 | 384 | 0.034709698 | 3.647437767 | 3.163036683 | 0.096466298 | 0.094218776 | 0 |
| CTTACCGGTGCTATTG-1 | 1344662 | 12942 | 145731 | 106768 | 1079221 | 349 | 0.031485573 | 3.3107972 | 3.100494374 | 0.100402233 | 0.098392658 | 0 |
| CTTACCGTCACACGTA-1 | 1179048 | 9844 | 127635 | 97632 | 943937 | 305 | 0.027515661 | 2.897565533 | 3.085791989 | 0.108736155 | 0.109693089 | 0 |
| CTTACCGTCCGCGACA-1 | 1663906 | 45922 | 172190 | 152250 | 1293544 | 418 | 0.03765714 | 3.9596711 | 3.097094068 | 0.09157913 | 0.093586337 | 0 |
| CTTAGGACACACGCAC-1 | 2162164 | 39880 | 227760 | 191897 | 1702627 | 550 | 0.049644444 | 5.156974133 | 3.046675523 | 0.082369825 | 0.084887742 | 0 |
| CTTAGGATCTGGGCTG-1 | 1481830 | 34948 | 153830 | 130188 | 1162864 | 376 | 0.033888596 | 3.569604567 | 3.119705053 | 0.099946647 | 0.10084129 | 0 |
| CTTCTCTAGCTGCCCA-1 | 2289280 | 53969 | 241452 | 209231 | 1784628 | 576 | 0.052005103 | 5.3963407 | 3.107273787 | 0.079286156 | 0.079918073 | 0 |
| CTTCTCTAGGTCGGAT-1 | 2191864 | 22693 | 235133 | 188443 | 1745595 | 564 | 0.050982455 | 5.2703951 | 3.105562619 | 0.086059049 | 0.082589925 | 0 |
| CTTCTCTCAATGTAGA-1 | 976130 | 8538 | 105744 | 82690 | 779158 | 252 | 0.022754777 | 2.415797333 | 3.001966771 | 0.118342639 | 0.115189679 | 0 |
| CTTCTCTCAGCTATGT-1 | 1263638 | 15403 | 138880 | 100588 | 1008767 | 326 | 0.02944899 | 3.0968291 | 3.109206751 | 0.108412539 | 0.105914628 | 0 |
| CTTCTCTCATCGCATA-1 | 1557172 | 17041 | 166138 | 131029 | 1242964 | 402 | 0.036271866 | 3.783456533 | 2.967222546 | 0.09790768 | 0.095658259 | 0 |
| CTTCTCTTCCTATGAG-1 | 1250368 | 9979 | 136683 | 100188 | 1003518 | 324 | 0.029300541 | 3.075587533 | 3.0032322 | 0.106112775 | 0.10521838 | 0 |
| CTTGGCTAGACAGAGA-1 | 1600858 | 15526 | 172364 | 135920 | 1277048 | 413 | 0.037331676 | 3.913358767 | 3.060517606 | 0.09430851 | 0.0937657 | 0 |
| CTTGGCTAGCGCCTTG-1 | 1253604 | 6889 | 142183 | 107036 | 997496 | 322 | 0.029108119 | 3.0783841 | 3.071516348 | 0.102039166 | 0.100918306 | 0 |
| CTTGGCTAGTCTCGTA-1 | 1046986 | 11947 | 110730 | 96485 | 827824 | 267 | 0.024145741 | 2.5531081 | 3.00358703 | 0.113264749 | 0.111579134 | 0 |
| CTTGGCTGTTCTAGTG-1 | 1180350 | 12674 | 129704 | 93240 | 944732 | 305 | 0.027542008 | 2.896158567 | 3.095774068 | 0.108621867 | 0.106120356 | 0 |
| CTTTGCGGTCAAAGCG-1 | 1068926 | 35869 | 110974 | 96164 | 825919 | 267 | 0.023993883 | 2.557689433 | 3.114620182 | 0.112343159 | 0.113216065 | 0 |
| CTTTGCGTCCTCAGTC-1 | 1471636 | 29145 | 152586 | 124223 | 1165682 | 377 | 0.033927155 | 3.551343533 | 3.018356376 | 0.100917226 | 0.095813637 | 0 |
| GAAACTCGTGCGAACA-1 | 1571848 | 82219 | 162120 | 137717 | 1189792 | 384 | 0.034709358 | 3.6423319 | 3.094240726 | 0.097142868 | 0.098054525 | 0 |
| GAAACTCGTTCGTCAA-1 | 2135930 | 27021 | 230601 | 190929 | 1687379 | 545 | 0.049259709 | 5.110496567 | 2.979772072 | 0.084960479 | 0.083211653 | 0 |
| GAAACTCTCACGCTCT-1 | 2130534 | 106771 | 218451 | 192550 | 1612762 | 521 | 0.046673393 | 4.811697433 | 3.000518599 | 0.08935311 | 0.087046346 | 0 |
| GAAACTCTCACGTAGT-1 | 1542580 | 15913 | 170467 | 129785 | 1226415 | 396 | 0.03581356 | 3.741242833 | 3.042750401 | 0.094557746 | 0.093273384 | 0 |
| GAAACTCTCTGGTGAT-1 | 2495682 | 28178 | 275288 | 218563 | 1973653 | 638 | 0.05765117 | 5.926617433 | 2.976117579 | 0.081069753 | 0.077453221 | 0 |
| GAAATGAAGGCTCTTA-1 | 1624258 | 36501 | 176006 | 140795 | 1270956 | 411 | 0.037095872 | 3.8867127 | 3.072867677 | 0.097114455 | 0.094399575 | 0 |
| GAAATGACACCCTCTA-1 | 3053208 | 50683 | 331744 | 263350 | 2407431 | 778 | 0.070268292 | 7.194647133 | 3.071199204 | 0.071206171 | 0.070016325 | 0 |
| GAACATCGTACTTGCA-1 | 1716378 | 80359 | 177834 | 155279 | 1302906 | 421 | 0.03790936 | 3.972400433 | 3.050301511 | 0.09048902 | 0.0917392 | 0 |
| GAACATCTCATGCGCG-1 | 913584 | 74014 | 89652 | 81012 | 668906 | 216 | 0.019455263 | 2.085101733 | 3.001604977 | 0.126304485 | 0.125028462 | 0 |
| GAACCTACAAAGGTTA-1 | 1874472 | 15467 | 206866 | 163088 | 1489051 | 481 | 0.043479318 | 4.5191123 | 3.029858487 | 0.090336444 | 0.08918915 | 0 |
| GAACCTACAATCGCGC-1 | 1155310 | 23774 | 119606 | 115712 | 896218 | 290 | 0.026104245 | 2.764222367 | 3.111349462 | 0.111685547 | 0.108421981 | 0 |
| GAACCTAGTCACATGT-1 | 1320126 | 16512 | 148517 | 101515 | 1053582 | 340 | 0.03074443 | 3.228522067 | 3.032847182 | 0.100351012 | 0.099074915 | 0 |
| GAACCTAGTCGAGTAG-1 | 1478760 | 18729 | 156095 | 123110 | 1180826 | 381 | 0.034519334 | 3.624958133 | 3.023052013 | 0.096022323 | 0.096465549 | 0 |
| GAACCTAGTTCGAGCC-1 | 1327568 | 54281 | 138490 | 117847 | 1016950 | 329 | 0.029541528 | 3.1145524 | 3.011322792 | 0.10361213 | 0.101656405 | 0 |
| GAACGGACACTTGCAA-1 | 1806992 | 20850 | 198891 | 154785 | 1432466 | 463 | 0.041755683 | 4.350981333 | 3.072373375 | 0.087650834 | 0.087327631 | 0 |
| GAACGGACAGGTCTGC-1 | 2242342 | 43803 | 243718 | 180991 | 1773830 | 573 | 0.051648559 | 5.335640633 | 3.324063411 | 0.125915174 | 0.12395808 | 1 |
| GAACGGAGTACGTTTG-1 | 1641982 | 30817 | 178954 | 142092 | 1290119 | 417 | 0.037602925 | 3.932228267 | 3.083304916 | 0.093672218 | 0.093226763 | 0 |
| GAACGGATCTTCACGC-1 | 2074354 | 30954 | 224088 | 186351 | 1632961 | 527 | 0.047557803 | 4.9460844 | 3.14422544 | 0.086541769 | 0.086801784 | 0 |
| GAAGCAGAGGGAGTAA-1 | 1747690 | 20773 | 193324 | 145277 | 1388316 | 448 | 0.040430343 | 4.2181149 | 3.164989454 | 0.091741223 | 0.091516455 | 0 |
| GAAGCAGCACACGCAC-1 | 2517470 | 37601 | 273868 | 222433 | 1983568 | 641 | 0.05782491 | 5.955686333 | 3.025062336 | 0.079392679 | 0.07923693 | 0 |
| GAATAAGAGGCTACGA-1 | 1310700 | 15219 | 142569 | 114065 | 1038847 | 336 | 0.03031693 | 3.202001333 | 3.008851569 | 0.107183957 | 0.103695582 | 0 |
| GAATAAGCAAATACGA-1 | 1444902 | 28523 | 156080 | 126632 | 1133667 | 366 | 0.033020363 | 3.477850333 | 3.142372111 | 0.098606309 | 0.096146788 | 0 |
| GAATAAGCAGAGATCG-1 | 1405216 | 90974 | 142076 | 122745 | 1049421 | 339 | 0.030502127 | 3.223729967 | 2.790193697 | 0.106124835 | 0.102260099 | 0 |
| GAATGAACAACCTGCG-1 | 1353282 | 114117 | 135817 | 113857 | 989491 | 320 | 0.028754164 | 3.044604833 | 3.032668134 | 0.107976624 | 0.106225734 | 0 |
| GAATGAATCTTCACAT-1 | 1419246 | 14180 | 155458 | 128195 | 1121413 | 362 | 0.032700565 | 3.424381333 | 3.031093308 | 0.101512537 | 0.099594309 | 0 |
| GACACGCCAAATGATG-1 | 1339860 | 17337 | 148031 | 106291 | 1068201 | 345 | 0.031238517 | 3.272402133 | 3.138061583 | 0.102200963 | 0.102213792 | 0 |
| GACAGAGAGCCTAACT-1 | 2165482 | 63675 | 231520 | 191947 | 1678340 | 542 | 0.048705597 | 5.0075142 | 3.142702879 | 0.08445602 | 0.08445602 | 0 |
| GACAGAGCACAGACCC-1 | 1853144 | 35133 | 196862 | 171059 | 1450090 | 468 | 0.042228588 | 4.427032433 | 3.111223199 | 0.08924619 | 0.089764452 | 0 |
| GACAGAGCAGTAATCC-1 | 2410034 | 63379 | 256031 | 208503 | 1882121 | 608 | 0.054553541 | 5.603322767 | 3.101909878 | 0.086467359 | 0.086467359 | 0 |
| GACAGAGCATCGGGCT-1 | 1206666 | 97913 | 108314 | 100954 | 899485 | 291 | 0.026140103 | 2.7692769 | 3.000127856 | 0.108934316 | 0.111346305 | 0 |
| GACAGAGTCAGCCCGA-1 | 1977856 | 26529 | 213053 | 179911 | 1558363 | 503 | 0.045458385 | 4.7368325 | 3.035723163 | 0.085453581 | 0.0858874 | 0 |
| GACAGAGTCCTACGAA-1 | 1480510 | 16990 | 165520 | 122338 | 1175662 | 380 | 0.034257549 | 3.598007567 | 3.097890475 | 0.096885677 | 0.094605732 | 0 |
| GACAGAGTCCTGCGTT-1 | 2125662 | 50676 | 234646 | 180134 | 1660206 | 536 | 0.048283671 | 4.997184467 | 3.019277759 | 0.088629931 | 0.088218794 | 0 |
| GACCAATAGACTTTCG-1 | 1551048 | 75972 | 162503 | 139935 | 1172638 | 379 | 0.03414249 | 3.5937185 | 3.020096611 | 0.098492683 | 0.096527375 | 0 |
| GACCAATAGGACAGAA-1 | 1526494 | 16098 | 168112 | 125003 | 1217281 | 393 | 0.035524867 | 3.725419 | 3.077322005 | 0.098513191 | 0.09294981 | 0 |
| GACCAATTCCGGCTAG-1 | 2078448 | 47228 | 227554 | 184269 | 1619397 | 523 | 0.047290496 | 4.918062067 | 3.012175025 | 0.086372042 | 0.084574453 | 0 |

| | | | | | | | | | | | | |
|---|---|---|---|---|---|---|---|---|---|---|---|---|
| GACCTGGGTAGCCCTG-1 | 1507576 | 30056 | 160550 | 130904 | 1186066 | 383 | 0.03461075 | 3.6470282 | 3.064097919 | 0.095999814 | 0.097439951 | 0 |
| GACGCGTAGTAACCCT-1 | 1314248 | 56235 | 124120 | 116042 | 1017851 | 329 | 0.029634674 | 3.1338713 | 3.066245721 | 0.102695246 | 0.102695246 | 0 |
| GACGCGTCAGGTCAAG-1 | 1014420 | 31407 | 91650 | 87949 | 803414 | 260 | 0.023429188 | 2.491088867 | 3.093265831 | 0.119603439 | 0.115098895 | 0 |
| GACGGCTGTGTAATGA-1 | 1000748 | 18661 | 109270 | 79280 | 793537 | 256 | 0.023098136 | 2.444537467 | 2.966153212 | 0.117263836 | 0.114776129 | 0 |
| GACGGCTGTTGAGCAG-1 | 1885472 | 49625 | 202240 | 166556 | 1467051 | 474 | 0.042808146 | 4.4797851 | 3.084720841 | 0.090263068 | 0.089157872 | 0 |
| GACGTGCGTGCGTCAC-1 | 1450280 | 16227 | 143118 | 132494 | 1158441 | 374 | 0.033737767 | 3.545438667 | 3.047979941 | 0.097451603 | 0.097451603 | 0 |
| GACTAACCACCCGTGA-1 | 1395092 | 23880 | 152241 | 122273 | 1096698 | 354 | 0.031977537 | 3.371270333 | 2.984218302 | 0.099639901 | 0.099127411 | 0 |
| GACTAACCAGACACAG-1 | 1446066 | 22168 | 160403 | 123529 | 1139966 | 368 | 0.033261087 | 3.490100967 | 2.878661326 | 0.100443411 | 0.099138281 | 0 |
| GACTACACACAATCAC-1 | 2660244 | 39481 | 289705 | 248051 | 2083007 | 673 | 0.060716634 | 6.2424244 | 3.18591201 | 0.076658997 | 0.076522248 | 0 |
| GACTGCGCAGTAGGGT-1 | 2170000 | 36102 | 235259 | 185900 | 1712739 | 553 | 0.050026979 | 5.181878 | 3.050870452 | 0.083284338 | 0.083284338 | 0 |
| GACTGCGGTGTCTGTA-1 | 1635618 | 16369 | 178566 | 139524 | 1301159 | 420 | 0.037953245 | 3.9728971 | 3.051660464 | 0.093585147 | 0.091834371 | 0 |
| GACTGCGTCATATCTC-1 | 2264826 | 25519 | 242691 | 199311 | 1797305 | 581 | 0.052105801 | 5.362400933 | 3.070489777 | 0.083592645 | 0.083280216 | 0 |
| GACTGCGTCGTAGCGC-1 | 670938 | 9237 | 60215 | 60752 | 540734 | 175 | 0.01573814 | 1.691431467 | 2.993369303 | 0.143123183 | 0.140546077 | 0 |
| GAGCAGAAGAATCGTA-1 | 2216978 | 48046 | 239225 | 188015 | 1741692 | 563 | 0.050811488 | 5.266044067 | 3.084447507 | 0.085787459 | 0.0840052 | 0 |
| GAGCAGAAGTTATCGC-1 | 2176558 | 36693 | 235012 | 197154 | 1707699 | 552 | 0.049729883 | 5.155134933 | 3.055013675 | 0.087779692 | 0.086653753 | 0 |
| GAGCAGATCCGCCTAT-1 | 1711856 | 22022 | 188377 | 151855 | 1349602 | 436 | 0.039351329 | 4.091451767 | 3.068046202 | 0.093719704 | 0.09199494 | 0 |
| GAGGTGAAGCCGAACA-1 | 2121278 | 69618 | 223005 | 190890 | 1637765 | 529 | 0.04768697 | 4.964064667 | 3.031619992 | 0.084417538 | 0.08385759 | 0 |
| GAGGTGACACTAACTG-1 | 1059550 | 30858 | 113973 | 87748 | 826971 | 267 | 0.024081367 | 2.558490233 | 2.96361519 | 0.110460944 | 0.112755599 | 0 |
| GAGGTGACAGCTATGT-1 | 1249364 | 17608 | 137760 | 97935 | 996061 | 322 | 0.029071743 | 3.054865433 | 3.034710039 | 0.103394281 | 0.100634976 | 0 |
| GAGGTGACAGTGAATA-1 | 2726140 | 35698 | 297126 | 226754 | 2166562 | 700 | 0.063187278 | 6.491346267 | 3.044153686 | 0.072366932 | 0.071952718 | 0 |
| GAGGTGATCACACGGC-1 | 1401554 | 48905 | 148657 | 117683 | 1086309 | 351 | 0.031545617 | 3.324517333 | 3.125536328 | 0.117556725 | 0.118438842 | 1 |
| GAGTCCGAGGGTTCGA-1 | 2131810 | 33054 | 228821 | 194589 | 1675346 | 541 | 0.04887616 | 5.0922854 | 3.027165844 | 0.083107021 | 0.081583331 | 0 |
| GAGTCCGCATACGGTT-1 | 1850328 | 24091 | 202755 | 148461 | 1475021 | 476 | 0.043016021 | 4.4760191 | 3.093631709 | 0.087752162 | 0.085159131 | 0 |
| GAGTCCGGTACTGCTA-1 | 1311384 | 16339 | 140964 | 117972 | 1036109 | 335 | 0.030239065 | 3.181647967 | 3.037336855 | 0.106884186 | 0.103576913 | 0 |
| GAGTCCGGTCCTAGTA-1 | 1067466 | 15559 | 117361 | 84528 | 850018 | 275 | 0.024753602 | 2.6171019 | 3.098224558 | 0.114169704 | 0.114045212 | 0 |
| GAGTCCGTCAGACACT-1 | 2342610 | 41049 | 255765 | 207407 | 1838389 | 594 | 0.053505592 | 5.522209033 | 3.031324154 | 0.081281604 | 0.080735943 | 0 |
| GATCAGTCAAGCACGA-1 | 1438234 | 25436 | 155044 | 129042 | 1128712 | 365 | 0.032928964 | 3.475754633 | 2.957936824 | 0.096280823 | 0.09809861 | 0 |
| GATCAGTTCCACCCAT-1 | 2476684 | 27238 | 273097 | 211762 | 1964587 | 635 | 0.057389122 | 5.915584967 | 3.036962036 | 0.076831969 | 0.077662481 | 0 |
| GATCAGTTCCGCTCTA-1 | 2021266 | 55838 | 215003 | 186603 | 1563822 | 505 | 0.045595863 | 4.748382033 | 3.126734249 | 0.089424265 | 0.085164429 | 0 |
| GATCGATGTCCTACCT-1 | 692496 | 9082 | 75107 | 56286 | 552021 | 178 | 0.01611553 | 1.727162233 | 3.106981598 | 0.148158038 | 0.145026898 | 0 |
| GATCGCGAGTCCATAC-1 | 1973362 | 22739 | 213469 | 165444 | 1571710 | 508 | 0.045862498 | 4.7475962 | 3.0168857 | 0.089034164 | 0.087312138 | 0 |
| GATCGTAAGTAGGCCA-1 | 1791346 | 59937 | 183888 | 158114 | 1389407 | 449 | 0.040489652 | 4.2361245 | 3.012865231 | 0.088322654 | 0.087281407 | 0 |
| GATCGTACAACCGGAA-1 | 1640214 | 24329 | 175127 | 142220 | 1298538 | 419 | 0.037899615 | 3.9782238 | 3.067719802 | 0.098139582 | 0.094615624 | 0 |
| GATCGTACATACCAGT-1 | 2118092 | 33293 | 228755 | 188282 | 1667762 | 539 | 0.048694703 | 5.0415323 | 3.138066963 | 0.081609779 | 0.082855695 | 0 |
| GATCGTACATGATGCT-1 | 2710516 | 64379 | 290815 | 238436 | 2116886 | 684 | 0.061344933 | 6.2422825 | 3.163201523 | 0.085760418 | 0.084237103 | 1 |
| GATCTAGCAAGTGTTC-1 | 2368390 | 31654 | 257348 | 211351 | 1868037 | 603 | 0.054285403 | 5.588488167 | 3.108596192 | 0.085412275 | 0.082878267 | 0 |
| GATCTAGTCACGCTCT-1 | 1557342 | 50925 | 165106 | 139675 | 1201636 | 388 | 0.034868083 | 3.621311033 | 3.005199861 | 0.106458133 | 0.10034133 | 0 |
| GATGAAAAGTAATCCC-1 | 2292618 | 29727 | 246614 | 202032 | 1814245 | 586 | 0.052832474 | 5.453982867 | 3.12287395 | 0.083950471 | 0.083883712 | 0 |
| GATGAAACAAGTGCAG-1 | 1735164 | 31412 | 188885 | 154798 | 1360069 | 439 | 0.039705169 | 4.1515519 | 2.87047707 | 0.090783918 | 0.088652508 | 1 |
| GATGAAACATGCAGTT-1 | 1756572 | 28915 | 194158 | 144272 | 1389227 | 449 | 0.040497631 | 4.221143733 | 2.971020568 | 0.093727602 | 0.092974895 | 0 |
| GATGAAAGTAACGGCA-1 | 1916592 | 84951 | 201128 | 169503 | 1461010 | 472 | 0.042573285 | 4.4446107 | 3.016580826 | 0.087529974 | 0.086045265 | 0 |
| GATGAAAGTTCAGTAC-1 | 2110778 | 33609 | 230028 | 191638 | 1655503 | 535 | 0.048093077 | 4.952792467 | 2.903130017 | 0.086869133 | 0.086923523 | 0 |
| GATGAGGCAATGGCTT-1 | 2266998 | 108079 | 238016 | 196308 | 1724595 | 557 | 0.050182023 | 5.187408 | 3.003829811 | 0.091524376 | 0.091145284 | 1 |
| GATGCTAGTAGGCTGA-1 | 1521416 | 19389 | 165432 | 142624 | 1193971 | 386 | 0.034869701 | 3.676908867 | 3.068785737 | 0.095548724 | 0.095519433 | 0 |
| GATGCTAGTCTCTCGT-1 | 2148888 | 41912 | 229747 | 186792 | 1690437 | 546 | 0.04933529 | 5.113525 | 2.992848926 | 0.086053893 | 0.085153952 | 0 |
| GATGCTAGTTCCCTAC-1 | 2744570 | 28944 | 302181 | 233627 | 2179818 | 704 | 0.06346832 | 6.4779754 | 3.079342977 | 0.080868573 | 0.078277904 | 0 |
| GATGCTAGTTGGAACG-1 | 1665998 | 87995 | 173298 | 144934 | 1259771 | 407 | 0.036658219 | 3.8463398 | 2.960847267 | 0.090797073 | 0.092004266 | 0 |
| GATGCTATCACTATGG-1 | 1110078 | 16949 | 123829 | 88599 | 880701 | 284 | 0.025714216 | 2.7090558 | 3.108377008 | 0.115354161 | 0.112007878 | 0 |
| GATGCTATCATCAGCA-1 | 1808598 | 39890 | 193938 | 160610 | 1414160 | 457 | 0.041136882 | 4.2957505 | 3.092979639 | 0.090124196 | 0.088761988 | 0 |
| GATTCAGAGATGTGGC-1 | 2881214 | 31849 | 317688 | 251040 | 2280637 | 737 | 0.066539417 | 6.8103709 | 3.097838364 | 0.074389532 | 0.073366297 | 0 |
| GATTCAGGTTCAGTGT-1 | 2196964 | 35687 | 239077 | 188028 | 1734172 | 560 | 0.050545926 | 5.216228833 | 3.142120551 | 0.087716322 | 0.086008679 | 0 |
| GATTCAGTCGACAACT-1 | 2363820 | 66860 | 255198 | 214580 | 1827182 | 590 | 0.053148991 | 5.466758933 | 3.092181906 | 0.085699998 | 0.083781515 | 0 |
| GCAAACTAGCCACGTC-1 | 1280606 | 22599 | 138617 | 111399 | 1007991 | 326 | 0.029314657 | 3.090762933 | 3.044396787 | 0.105751905 | 0.106634817 | 0 |
| GCAAACTCAAGCCAAG-1 | 1970636 | 32455 | 213362 | 173018 | 1551801 | 501 | 0.045278175 | 4.7175169 | 3.142023056 | 0.084096916 | 0.083661195 | 0 |
| GCAAACTCACATTGGT-1 | 2304558 | 30833 | 253175 | 192942 | 1827608 | 590 | 0.053358368 | 5.503516067 | 3.081819796 | 0.080413701 | 0.079181238 | 0 |
| GCAAACTTCCTACGAA-1 | 1493416 | 19440 | 165142 | 127175 | 1181659 | 382 | 0.034400279 | 3.610446133 | 3.182539344 | 0.100484936 | 0.097690555 | 0 |
| GCAATCAAGGCAACAC-1 | 2040878 | 38276 | 220581 | 183989 | 1598032 | 516 | 0.04654745 | 4.8263927 | 2.973894327 | 0.086502293 | 0.083612672 | 0 |
| GCAATCAGTTAGTGAA-1 | 2844122 | 27445 | 313809 | 238324 | 2264544 | 732 | 0.066151562 | 6.7817772 | 3.122862603 | 0.072597467 | 0.070689066 | 0 |
| GCAATCAGTTGCCTCT-1 | 2284926 | 51110 | 247231 | 197286 | 1789299 | 578 | 0.052132401 | 5.3734342 | 3.004440592 | 0.085478066 | 0.084612502 | 0 |
| GCAATCATCCCTGAGG-1 | 1955304 | 29829 | 212996 | 167315 | 1545164 | 499 | 0.045125148 | 4.702526067 | 2.996079323 | 0.086022886 | 0.084392016 | 0 |
| GCAATCATCCGTTCAC-1 | 3329206 | 24212 | 362848 | 305603 | 2636543 | 852 | 0.077159929 | 7.851600267 | 3.009597215 | 0.063775236 | 0.063232964 | 0 |
| GCACATAGTGAGCGAT-1 | 2628698 | 42475 | 283979 | 237963 | 2064281 | 667 | 0.060225533 | 6.192304033 | 2.985358576 | 0.079038119 | 0.078528773 | 0 |
| GCACTCTAGACTTTCG-1 | 1519538 | 90881 | 155091 | 131986 | 1141580 | 369 | 0.033226163 | 3.5074068 | 3.1464047 | 0.095345632 | 0.09710856 | 0 |
| GCACTCTAGCATCTTG-1 | 1195050 | 21377 | 128951 | 98457 | 946265 | 306 | 0.027651376 | 2.9226492 | 3.128532167 | 0.107455851 | 0.105965383 | 0 |
| GCACTCTAGTCCAGGA-1 | 1568648 | 20518 | 170669 | 136877 | 1240584 | 401 | 0.036212069 | 3.808155433 | 3.033669316 | 0.096348778 | 0.096319297 | 0 |
| GCACTCTCAAGCATGG-1 | 1192130 | 15695 | 130380 | 100735 | 945320 | 305 | 0.027650322 | 2.9171426 | 3.081614974 | 0.107415174 | 0.107108926 | 0 |
| GCACTCTTCCACCTTG-1 | 1427692 | 28006 | 154927 | 119921 | 1124838 | 363 | 0.032830372 | 3.4552007 | 3.077858051 | 0.097327179 | 0.09730119 | 0 |
| GCAGCCACAGACACGA-1 | 1704214 | 28208 | 184453 | 146714 | 1344839 | 434 | 0.039198108 | 4.0858639 | 2.154069664 | 0.083564732 | 0.081002764 | 1 |
| GCAGCCACATTGCATG-1 | 1239600 | 14768 | 136645 | 102956 | 985231 | 318 | 0.02867376 | 3.021949333 | 2.982728009 | 0.107036511 | 0.10653768 | 0 |
| GCAGCCAGTCGTCTTC-1 | 2385760 | 22432 | 254762 | 222033 | 1886533 | 609 | 0.054962719 | 5.682672333 | 3.089569942 | 0.093613077 | 0.093613077 | 1 |
| GCAGCCATCGCTCTAC-1 | 2487958 | 25187 | 269366 | 214977 | 1978428 | 639 | 0.057625639 | 5.922392267 | 3.0587458 | 0.080318827 | 0.078455008 | 0 |
| GCAGTTAAGGCCCTTG-1 | 1509446 | 20626 | 164916 | 129629 | 1194275 | 386 | 0.034862012 | 3.656346067 | 3.106214232 | 0.094801539 | 0.096856946 | 0 |
| GCAGTTAGTAGCAGTG-1 | 1449734 | 17626 | 158807 | 117095 | 1156206 | 373 | 0.033794401 | 3.545397167 | 3.108552842 | 0.100839918 | 0.10051095 | 0 |
| GCAGTTAGTGGAAGGG-1 | 1980468 | 42189 | 211206 | 173171 | 1553902 | 502 | 0.045370147 | 4.722401833 | 2.993713417 | 0.084592204 | 0.085698546 | 0 |
| GCAGTTATCGCCCAGA-1 | 1966408 | 19823 | 211963 | 174201 | 1560421 | 504 | 0.045549685 | 4.743428867 | 2.987555139 | 0.088305559 | 0.088914554 | 0 |
| GCATACAAGATCCTGT-1 | 2392730 | 38323 | 263406 | 207929 | 1883072 | 608 | 0.054957698 | 5.675742867 | 3.01899458 | 0.081698688 | 0.081159666 | 0 |
| GCATACAAGGACAGAA-1 | 1489600 | 19846 | 163454 | 123558 | 1182742 | 382 | 0.034509225 | 3.623158433 | 2.947626016 | 0.0973647 | 0.095312642 | 0 |
| GCATACACACGTGCGT-1 | 1984236 | 40353 | 212795 | 178758 | 1552330 | 501 | 0.045267937 | 4.714747567 | 3.111939026 | 0.085088597 | 0.085727211 | 0 |
| GCATACATCAGATGTC-1 | 1673148 | 14625 | 183577 | 140737 | 1334209 | 431 | 0.038897689 | 4.0566664 | 3.00917555 | 0.094238436 | 0.092993825 | 0 |
| GCATACATCCGCTCTA-1 | 1703458 | 39949 | 183826 | 146943 | 1332740 | 431 | 0.03890419 | 4.0640419 | 3.031236236 | 0.089878028 | 0.091156646 | 0 |
| GCATGATCACGATAGG-1 | 1351994 | 19141 | 145826 | 113885 | 1073142 | 347 | 0.031370797 | 3.3006619 | 3.019103893 | 0.100141611 | 0.098599609 | 0 |
| GCATGATCACTCACCT-1 | 2106780 | 64803 | 221376 | 192370 | 1628231 | 526 | 0.047385202 | 4.933765667 | 3.168762522 | 0.085287861 | 0.085287861 | 0 |
| GCATGATCATGACCGC-1 | 1244540 | 15452 | 130972 | 115272 | 982844 | 317 | 0.028584445 | 3.022511833 | 3.082922568 | 0.107686824 | 0.104312839 | 0 |
| GCATGATGTCGCTCAG-1 | 1842078 | 42872 | 195426 | 167368 | 1436412 | 464 | 0.041909257 | 4.3852703 | 3.048477487 | 0.088146414 | 0.089335217 | 0 |
| GCATGATGTTCGCGTG-1 | 1396812 | 41917 | 147413 | 116344 | 1091138 | 352 | 0.031751923 | 3.3469385 | 2.987036787 | 0.105147542 | 0.100789056 | 0 |
| GCATGATTCCGTTAAG-1 | 1254200 | 21583 | 136786 | 104439 | 991392 | 320 | 0.028959035 | 3.058290833 | 3.038747523 | 0.102176359 | 0.101292125 | 0 |
| GCATGCGAGTAACCCT-1 | 2227086 | 91553 | 231739 | 191040 | 1712754 | 553 | 0.049878367 | 5.179085 | 3.0184376 | 0.085546861 | 0.08398285 | 0 |
| GCATGTATCCACAGCG-1 | 1077684 | 47690 | 112482 | 92792 | 824720 | 266 | 0.023897812 | 2.545190933 | 3.07210323 | 0.11729829 | 0.111778321 | 0 |
| GCCAAATAGTAAGTAC-1 | 1949582 | 33242 | 208713 | 181667 | 1525960 | 493 | 0.04436075 | 4.5977005 | 3.062986644 | 0.087507252 | 0.084651737 | 0 |
| GCCAAATCATCGGCGT-1 | 1585204 | 35825 | 166744 | 142314 | 1240321 | 401 | 0.036100626 | 3.791823667 | 3.102674481 | 0.092157279 | 0.092157279 | 1 |
| GCCAAATCATTTGTGG-1 | 1337694 | 25725 | 138141 | 115251 | 1058577 | 342 | 0.030846839 | 3.248924433 | 3.037419291 | 0.102947705 | 0.100829551 | 0 |
| GCCAAATTCCGCGGAT-1 | 2700238 | 43842 | 292444 | 232946 | 2131006 | 688 | 0.062189074 | 6.372317367 | 3.172082351 | 0.076799486 | 0.073196974 | 0 |
| GCCAAATTCTACAGTG-1 | 1237550 | 16284 | 137427 | 98354 | 985485 | 318 | 0.02874898 | 3.0219803 | 3.035727728 | 0.10841836 | 0.103366789 | 0 |
| GCCTCTAGTACGGTGA-1 | 1531882 | 21114 | 166086 | 128778 | 1215904 | 393 | 0.03546554 | 3.7245339 | 3.049646599 | 0.096928501 | 0.097903428 | 0 |

| | | | | | | | | | | | | |
|---|---|---|---|---|---|---|---|---|---|---|---|---|
| GCCTCTATCACGTAGT-1 | 1419696 | 14326 | 156326 | 122346 | 1126698 | 364 | 0.032922555 | 3.4554272 | 3.048362123 | 0.102377502 | 0.101427486 | 0 |
| GCGACCACAACTCATG-1 | 1136444 | 18209 | 123847 | 93056 | 901332 | 291 | 0.026248254 | 2.769514733 | 3.039311481 | 0.103652108 | 0.104602808 | 0 |
| GCGACCAGTCCTACCT-1 | 1891298 | 71540 | 199578 | 169206 | 1450974 | 469 | 0.042196593 | 4.400750267 | 2.884008041 | 0.091970732 | 0.089665349 | 0 |
| GCGAGAAAGCTTCAGT-1 | 1119460 | 7521 | 121890 | 94084 | 895965 | 289 | 0.026171881 | 2.760192933 | 3.089326166 | 0.114496316 | 0.112441357 | 0 |
| GCGCAGTAGCGATGAC-1 | 2212136 | 31338 | 244800 | 192987 | 1743011 | 563 | 0.050791746 | 5.266660233 | 2.999587986 | 0.084795099 | 0.082467346 | 0 |
| GCGCAGTGTACTCGTA-1 | 1001110 | 14102 | 105180 | 84161 | 797667 | 258 | 0.023252843 | 2.470162667 | 3.018899953 | 0.114932278 | 0.117597535 | 0 |
| GCGCAGTGTGACACGA-1 | 1393070 | 14829 | 149232 | 126156 | 1102853 | 356 | 0.032173614 | 3.3965511 | 3.106374213 | 0.099100041 | 0.100014594 | 0 |
| GCGCAGTGTGACCAAG-1 | 1699622 | 34643 | 179720 | 144847 | 1340412 | 433 | 0.039222512 | 4.097738267 | 3.117900209 | 0.09530551 | 0.093869249 | 0 |
| GCGCCAAAGCGAGAAA-1 | 1154256 | 21181 | 130667 | 95651 | 906757 | 293 | 0.026420885 | 2.798021633 | 3.033798036 | 0.112476028 | 0.108814456 | 0 |
| GCGCCAAGTTGCCAAT-1 | 2001368 | 19288 | 214930 | 169184 | 1597966 | 516 | 0.046684363 | 4.8431654 | 3.043044288 | 0.087918427 | 0.084132242 | 0 |
| GCGCCAATCCTCATCG-1 | 1388218 | 60090 | 142934 | 121297 | 1063897 | 344 | 0.030949828 | 3.2647979 | 3.171219353 | 0.099593979 | 0.099396853 | 0 |
| GCGCGATAGGACTGGT-1 | 2082546 | 30088 | 221435 | 183834 | 1647189 | 532 | 0.048114145 | 4.980123167 | 3.106563432 | 0.082741137 | 0.084254232 | 0 |
| GCGCGATAGGATATAC-1 | 1783566 | 21381 | 192003 | 150078 | 1420104 | 459 | 0.041428227 | 4.311192767 | 3.116289972 | 0.093549216 | 0.090382229 | 0 |
| GCGCGATGTCATCTAG-1 | 786220 | 11571 | 84865 | 67995 | 621789 | 201 | 0.018169034 | 1.932812467 | 2.992012723 | 0.14570691 | 0.143186783 | 1 |
| GCGGGTTAGGAGTTTA-1 | 3468752 | 87806 | 357613 | 307766 | 2715567 | 877 | 0.079353661 | 8.056599667 | 2.977217169 | 0.065007975 | 0.065007975 | 0 |
| GCGGGTTAGTGTATCC-1 | 2134686 | 57681 | 219032 | 188563 | 1669410 | 539 | 0.048313446 | 4.965499167 | 3.041017151 | 0.083847749 | 0.085234433 | 0 |
| GCGGGTTTCTATGCCC-1 | 1786222 | 32765 | 187875 | 171146 | 1394436 | 450 | 0.040545356 | 4.2169994 | 3.036351366 | 0.100398457 | 0.093364927 | 0 |
| GCTCCTAAGATCCCGC-1 | 2217126 | 12386 | 239320 | 203833 | 1761587 | 569 | 0.051396997 | 5.329333533 | 3.113858678 | 0.080990471 | 0.080220724 | 0 |
| GCTCCTAAGCCCGAAA-1 | 1569384 | 24000 | 170193 | 135645 | 1239546 | 400 | 0.03614803 | 3.7945494 | 2.892923352 | 0.096614095 | 0.093717989 | 0 |
| GCTCCTATCCAGAAT-1 | 1973500 | 30021 | 213755 | 180520 | 1549204 | 500 | 0.045136819 | 4.6742327 | 2.980773875 | 0.084712928 | 0.084712928 | 0 |
| GCTCTGTAGCATGGCA-1 | 1889490 | 29841 | 203833 | 168684 | 1487132 | 480 | 0.04331561 | 4.527392933 | 3.04766235 | 0.089311733 | 0.087208433 | 0 |
| GCTCTGTAGTTAGCGG-1 | 1473400 | 43378 | 155035 | 134302 | 1140685 | 368 | 0.033261167 | 3.502342533 | 2.972685241 | 0.10085759 | 0.10131738 | 0 |
| GCTCTGTCAAACTCAC-1 | 2338690 | 22205 | 254668 | 210923 | 1850894 | 598 | 0.053951013 | 5.582111967 | 2.970937171 | 0.080174493 | 0.07920302 | 0 |
| GCTCTGTTCACTCATT-1 | 2039068 | 26192 | 224314 | 172988 | 1615574 | 522 | 0.047124909 | 4.882893467 | 2.990539034 | 0.085816903 | 0.086585158 | 0 |
| GCTGCAGAGGAGTTGC-1 | 2145342 | 24600 | 230800 | 191143 | 1698799 | 549 | 0.049404521 | 5.125430267 | 3.0742614 | 0.087614577 | 0.086601345 | 0 |
| GCTGCAGGTAACGTGG-1 | 1601104 | 28418 | 170676 | 136656 | 1265354 | 409 | 0.03697472 | 3.8746192 | 3.040706141 | 0.090306945 | 0.090306945 | 0 |
| GCTGCAGGTCGCGAAA-1 | 1184218 | 29060 | 122966 | 107769 | 924423 | 299 | 0.026970945 | 2.858519567 | 3.113698027 | 0.106361241 | 0.104903508 | 0 |
| GCTGCAGGTTGAACAA-1 | 1378032 | 26260 | 148603 | 120869 | 1082300 | 350 | 0.031538656 | 3.3124057 | 2.92472049 | 0.101450336 | 0.100685233 | 0 |
| GCTGCGACACCTCCAT-1 | 2180056 | 24785 | 237846 | 190074 | 1727351 | 558 | 0.050421845 | 5.208405 | 3.081480534 | 0.083748554 | 0.082533339 | 0 |
| GCTGCGATCACTACAG-1 | 1196642 | 16891 | 130169 | 98628 | 950954 | 307 | 0.027671594 | 2.926947267 | 3.110108226 | 0.110095855 | 0.104244951 | 0 |
| GCTGCTTCACGATCTA-1 | 2009778 | 46929 | 218270 | 178555 | 1566024 | 506 | 0.045646137 | 4.7583443 | 3.111742438 | 0.089364508 | 0.087736537 | 0 |
| GCTGCTTGTCATCTAG-1 | 1271376 | 13723 | 139301 | 108471 | 1009881 | 326 | 0.02952753 | 3.1090056 | 3.091755565 | 0.103493786 | 0.103493786 | 0 |
| GCTGCTTTCAGTACAC-1 | 2594706 | 28978 | 282947 | 241106 | 2041675 | 660 | 0.059459654 | 6.117805267 | 3.113147371 | 0.076547431 | 0.075181457 | 0 |
| GCTGGGTAGGCATATA-1 | 1019942 | 13332 | 109971 | 78207 | 818432 | 264 | 0.023928221 | 2.526921433 | 3.080781902 | 0.113967818 | 0.115128256 | 0 |
| GCTGGGTCAACTCCGG-1 | 2003818 | 61327 | 208539 | 179873 | 1554079 | 502 | 0.045299218 | 4.7369394 | 2.988943367 | 0.083961261 | 0.083162585 | 0 |
| GCTGGGTGTGTAGTTC-1 | 2150790 | 78467 | 221476 | 187721 | 1663126 | 537 | 0.048009844 | 4.917046367 | 2.971244894 | 0.088401051 | 0.089070305 | 0 |
| GCTGGGTGTGTCGCTG-1 | 1083488 | 33420 | 111155 | 94914 | 843999 | 273 | 0.02462948 | 2.615526967 | 3.077092965 | 0.112552134 | 0.112552134 | 0 |
| GCTGGGTTCTATGGTG-1 | 1179430 | 18997 | 127464 | 93994 | 938975 | 303 | 0.027392034 | 2.8785921 | 3.024250472 | 0.10586632 | 0.10647569 | 0 |
| GCTTCCAAGGGTTTCT-1 | 2114276 | 63489 | 226483 | 188756 | 1635548 | 528 | 0.047591927 | 4.928545767 | 3.104959188 | 0.083995841 | 0.085598993 | 0 |
| GCTTCCATCCTACGAA-1 | 1383004 | 16058 | 152214 | 118858 | 1095874 | 354 | 0.031944853 | 3.360852333 | 2.949855818 | 0.101092708 | 0.098330771 | 0 |
| GCTTCCATCTGGCCTT-1 | 2235898 | 66248 | 236536 | 202863 | 1730251 | 559 | 0.050402382 | 5.228590933 | 3.108784619 | 0.082721347 | 0.081934021 | 0 |
| GCTTCCATCTTCCGAC-1 | 1562214 | 37524 | 166789 | 147342 | 1210559 | 391 | 0.035232653 | 3.692380167 | 3.0966491 | 0.101968317 | 0.096086885 | 0 |
| GCTTGAACAGTGCTTA-1 | 1907858 | 27898 | 207981 | 166443 | 1505536 | 486 | 0.043967841 | 4.575037067 | 3.019939903 | 0.08666868 | 0.085951881 | 0 |
| GCTTGAAGTGAGCGAT-1 | 2641892 | 44237 | 285824 | 239980 | 2071851 | 669 | 0.060461778 | 6.195382633 | 2.991228694 | 0.095440983 | 0.095296133 | 1 |
| GGAAAGCAGCCAGAAC-1 | 2147390 | 48220 | 227431 | 193417 | 1678322 | 542 | 0.048820358 | 5.043704733 | 2.992315614 | 0.08826692 | 0.087228323 | 0 |
| GGAAAGCCATGACTCA-1 | 1805330 | 87916 | 187988 | 154409 | 1375017 | 444 | 0.039960363 | 4.177748967 | 3.095924658 | 0.090422541 | 0.088148011 | 0 |
| GGAAAGCTCCACCTTG-1 | 1024530 | 20904 | 108410 | 90219 | 804997 | 260 | 0.023486247 | 2.487053367 | 3.019974863 | 0.117876784 | 0.114799225 | 0 |
| GGAAAGCTCGTGTACA-1 | 1224228 | 29047 | 114052 | 107820 | 973309 | 314 | 0.028435231 | 3.0005882 | 3.144325211 | 0.106542738 | 0.106503394 | 0 |
| GGAACTTAGGGTCTCC-1 | 2034862 | 13750 | 218315 | 193345 | 1609452 | 520 | 0.047032897 | 4.883649933 | 3.034424949 | 0.086651973 | 0.085101635 | 0 |
| GGAACTTCACAAGCTT-1 | 384970 | 6873 | 41438 | 30273 | 306386 | 99 | 0.008935834 | 0.965921567 | 3.250678026 | 0.224546104 | 0.224546104 | 1 |
| GGAACTTGTCGGGAAT-1 | 2125044 | 30224 | 230842 | 192963 | 1671015 | 540 | 0.048787707 | 5.050710967 | 2.963545699 | 0.08397347 | 0.082723821 | 0 |
| GGAACTTGTTTCCATT-1 | 2006678 | 30783 | 219528 | 177324 | 1579043 | 510 | 0.046050611 | 4.7822081 | 3.024420042 | 0.086849674 | 0.086205717 | 0 |
| GGAACTTTCCGATACA-1 | 1884300 | 49928 | 202080 | 166954 | 1465338 | 473 | 0.042715764 | 4.4421336 | 2.949379497 | 0.088699314 | 0.088212985 | 0 |
| GGAATAATCTAAGACC-1 | 1922192 | 16715 | 205603 | 169403 | 1530471 | 494 | 0.044509785 | 4.618114667 | 3.121633912 | 0.1085547 | 0.1085547 | 1 |
| GGACAAGAGGGATTTA-1 | 1730070 | 33024 | 187609 | 149545 | 1359892 | 439 | 0.03966722 | 4.146693567 | 3.0381591 | 0.092199968 | 0.089806455 | 0 |
| GGACAAGGTGTAAAGT-1 | 584380 | 6228 | 63939 | 49791 | 464422 | 150 | 0.013594107 | 1.453776067 | 3.045282858 | 0.14658493 | 0.149554478 | 0 |
| GGACAAGGTTCAGTAC-1 | 1868202 | 40044 | 200839 | 168189 | 1459130 | 471 | 0.042351675 | 4.369242867 | 2.932403731 | 0.093069566 | 0.092551832 | 0 |
| GGACAAGTCGACGAAG-1 | 1799638 | 34909 | 192998 | 151754 | 1419977 | 459 | 0.041449417 | 4.3244544 | 2.934117831 | 0.09306896 | 0.090186927 | 0 |
| GGACAAGTCGCGCACA-1 | 1676210 | 52543 | 175573 | 148622 | 1299472 | 420 | 0.037823346 | 3.970268967 | 3.106661363 | 0.093188151 | 0.091578687 | 0 |
| GGACAGAAGGTATCGG-1 | 1786330 | 43952 | 191537 | 158790 | 1392051 | 450 | 0.04063412 | 4.2553263 | 2.890626665 | 0.092998267 | 0.092998267 | 0 |
| GGACAGACAAACCCTA-1 | 2020568 | 50815 | 217363 | 180771 | 1571619 | 508 | 0.045860992 | 4.771284633 | 3.064808303 | 0.083490179 | 0.082223704 | 0 |
| GGACAGACAGCCAGTC-1 | 1672180 | 24472 | 182937 | 141415 | 1323356 | 427 | 0.038509462 | 4.027986867 | 3.039098567 | 0.094487666 | 0.091095923 | 0 |
| GGACATTAGTCAAGCG-1 | 1105060 | 110959 | 109411 | 93064 | 791626 | 256 | 0.022994249 | 2.446703267 | 2.772892524 | 0.115093295 | 0.11677698 | 0 |
| GGACGTCAGAACTTCC-1 | 2248020 | 19705 | 242742 | 204418 | 1781155 | 575 | 0.05185345 | 5.3748853 | 3.017651655 | 0.081179634 | 0.080278905 | 0 |
| GGACGTCCACACGTGC-1 | 1778986 | 34586 | 193117 | 157425 | 1393858 | 450 | 0.040520955 | 4.2343649 | 3.057730831 | 0.090386625 | 0.089840611 | 0 |
| GGAGCAAAGGTGCTTT-1 | 2379744 | 56090 | 255102 | 210141 | 1858411 | 600 | 0.054038866 | 5.555326967 | 3.000241494 | 0.084029062 | 0.08230655 | 0 |
| GGAGCAACACGAGCTC-1 | 2606732 | 27342 | 282881 | 222454 | 2074055 | 670 | 0.060514957 | 6.244203067 | 3.493896287 | 0.097029229 | 0.097029229 | 1 |
| GGATGTTAGCCAGGAT-1 | 2517512 | 38568 | 272946 | 207528 | 1998470 | 646 | 0.058273923 | 5.982906333 | 3.008021599 | 0.075374659 | 0.076221388 | 0 |
| GGATGTTAGTGTACCT-1 | 2090596 | 63638 | 220121 | 186411 | 1620426 | 523 | 0.047151012 | 4.8946039 | 3.023555578 | 0.086974661 | 0.086974661 | 0 |
| GGATTACAGCCACCTG-1 | 1378584 | 37754 | 148784 | 116319 | 1075727 | 347 | 0.031337621 | 3.302494733 | 3.001143295 | 0.099802176 | 0.10032042 | 0 |
| GGATTACGTACTGTTG-1 | 1255810 | 23756 | 136323 | 101847 | 993884 | 321 | 0.028969399 | 3.0563242 | 3.073144399 | 0.10839763 | 0.10659828 | 0 |
| GGATTACTCCAACAAC-1 | 2254992 | 26544 | 243061 | 203461 | 1781926 | 576 | 0.05191863 | 5.3652253 | 3.067935922 | 0.086126394 | 0.085369169 | 0 |
| GGATTACTCTCGGTTC-1 | 387132 | 2855 | 41428 | 33730 | 309119 | 100 | 0.009041154 | 0.975458633 | 3.024146598 | 0.191868686 | 0.187923039 | 0 |
| GGCAATTGTACAACGG-1 | 1403666 | 58903 | 146902 | 118508 | 1079353 | 349 | 0.031476651 | 3.319677767 | 2.991177724 | 0.098373906 | 0.098373906 | 0 |
| GGCAATTGTGCAACCC-1 | 1687972 | 30092 | 181641 | 152512 | 1323727 | 428 | 0.03847926 | 4.025946 | 3.016164674 | 0.094471381 | 0.093736223 | 0 |
| GGCAATTTCTTGGATG-1 | 1460768 | 28610 | 159964 | 121103 | 1151091 | 372 | 0.033585528 | 3.5322566 | 3.109663263 | 0.105848369 | 0.100069599 | 0 |
| GGCCGATTCTGACGCG-1 | 784478 | 126886 | 70759 | 60443 | 526390 | 170 | 0.015264412 | 1.640160667 | 3.151735694 | 0.135468921 | 0.137234815 | 0 |
| GGCGACTGTTACTGAC-1 | 1920364 | 31606 | 208282 | 171279 | 1509197 | 488 | 0.043994761 | 4.571335667 | 3.014604036 | 0.088309739 | 0.084742618 | 0 |
| GGCGACTGTTCCGTCT-1 | 2109792 | 43688 | 222805 | 191908 | 1651391 | 533 | 0.047836834 | 4.920229467 | 2.981569733 | 0.08965188 | 0.090865277 | 0 |
| GGCGACTGTTGGAACG-1 | 1468714 | 93733 | 147908 | 131536 | 1095537 | 354 | 0.031853542 | 3.355657167 | 3.1061781 | 0.104906329 | 0.100966602 | 0 |
| GGCGACTTCCTTATTG-1 | 1070734 | 19102 | 116899 | 83166 | 851567 | 275 | 0.024846022 | 2.621863 | 3.110307273 | 0.117908791 | 0.112281924 | 0 |
| GGCTCGACAACTACTG-1 | 1156426 | 18761 | 127850 | 89158 | 920657 | 297 | 0.026797514 | 2.821837833 | 3.13171896 | 0.131819228 | 0.133369181 | 1 |
| GGCTCGACAATGGCTT-1 | 2281420 | 94077 | 241632 | 198034 | 1747677 | 565 | 0.05086476 | 5.263622633 | 2.826766227 | 0.083810565 | 0.084136408 | 0 |
| GGCTCGACATACTGCA-1 | 1705694 | 59064 | 180713 | 152049 | 1313868 | 424 | 0.038208023 | 4.000563467 | 3.131593713 | 0.094236485 | 0.094116841 | 0 |
| GGCTCGACATCATGGT-1 | 1755774 | 18913 | 190235 | 139595 | 1407031 | 455 | 0.04113467 | 4.277713067 | 3.213947277 | 0.12173873 | 0.12173873 | 1 |
| GGCTGGTCAACCTGCG-1 | 1140684 | 62288 | 116943 | 100005 | 861448 | 278 | 0.025031366 | 2.661605433 | 3.059652836 | 0.10871609 | 0.108700018 | 0 |
| GGCTGGTTCCTACGAA-1 | 1276010 | 18884 | 139108 | 107622 | 1010396 | 326 | 0.029384382 | 3.104276467 | 2.975456986 | 0.100221739 | 0.101691167 | 0 |
| GGCTGGTTCTGTGGCG-1 | 1433216 | 95065 | 145589 | 124125 | 1068437 | 345 | 0.031060575 | 3.280702133 | 3.058879939 | 0.100027742 | 0.099960763 | 0 |
| GGGAATGCAGATTGTC-1 | 1897772 | 31866 | 204762 | 164907 | 1496237 | 483 | 0.043589667 | 4.5239738 | 3.112996959 | 0.085446017 | 0.086981508 | 0 |
| GGGAATGCATCATGGT-1 | 1890202 | 21823 | 204921 | 152512 | 1510946 | 488 | 0.044145652 | 4.5846845 | 3.097117243 | 0.087561752 | 0.087069945 | 0 |
| GGGAATGGTGGTACAG-1 | 1391714 | 27533 | 148012 | 119988 | 1096181 | 354 | 0.031970575 | 3.370131367 | 3.014215181 | 0.097950606 | 0.09816503 | 0 |
| GGGACCTCATCCATCC-1 | 1929474 | 37118 | 204642 | 174871 | 1512843 | 489 | 0.043898288 | 4.561367867 | 3.12774254 | 0.085555745 | 0.083796839 | 0 |
| GGGACCTGTCCGGGTT-1 | 1934026 | 50292 | 207266 | 171814 | 1504654 | 486 | 0.043806415 | 4.5520625 | 3.04489451 | 0.089193446 | 0.088621007 | 0 |
| GGGACCTTCATCGACA-1 | 1810676 | 59627 | 190027 | 162308 | 1398714 | 452 | 0.040749255 | 4.270926733 | 2.986481687 | 0.093322513 | 0.092294866 | 0 |

| | | | | | | | | | | | | |
|---|---|---|---|---|---|---|---|---|---|---|---|---|
| GGGAGATAGGCTAGCA-1 | 1464372 | 81476 | 148641 | 124443 | 1109812 | 359 | 0.032276755 | 3.396830567 | 3.011594038 | 0.103625357 | 0.101485172 | 0 |
| GGGAGATCACAGACCC-1 | 1445274 | 16720 | 153282 | 137037 | 1138235 | 368 | 0.033112131 | 3.487409533 | 3.094999645 | 0.097551081 | 0.098875391 | 0 |
| GGGAGATCAGCGCTTG-1 | 1322744 | 61694 | 134038 | 109678 | 1017334 | 329 | 0.029634641 | 3.1273177 | 3.082320364 | 0.104040786 | 0.102598551 | 0 |
| GGGAGATCATGCAGTT-1 | 1610696 | 32403 | 172213 | 131210 | 1274870 | 412 | 0.037112186 | 3.884437167 | 3.056647465 | 0.099132984 | 0.094966948 | 0 |
| GGGCACTGTGCAACCC-1 | 1673214 | 15677 | 179753 | 158546 | 1319238 | 426 | 0.038371518 | 4.0164547 | 3.144706843 | 0.093349688 | 0.09247291 | 0 |
| GGGCACTGTTCAAGGG-1 | 1698298 | 43519 | 181239 | 149795 | 1323745 | 428 | 0.038650285 | 4.055688 | 3.167953737 | 0.09228736 | 0.089680329 | 0 |
| GGGCATCCACGCTGAC-1 | 2290296 | 40688 | 245640 | 205613 | 1798355 | 581 | 0.052392253 | 5.420374567 | 2.986430419 | 0.082881467 | 0.079505152 | 0 |
| GGGCATCGTCTCTATT-1 | 827990 | 29393 | 87000 | 65324 | 646273 | 209 | 0.018833598 | 1.9986004 | 3.031525806 | 0.127816069 | 0.125584586 | 0 |
| GGGCATCTCGAACCCG-1 | 1123326 | 56474 | 115030 | 99332 | 852490 | 275 | 0.024788732 | 2.640591633 | 2.976532489 | 0.110860214 | 0.110860214 | 0 |
| GGGTCTGCATCGGTCG-1 | 1079282 | 189727 | 93001 | 85205 | 711349 | 230 | 0.02057078 | 2.1924806 | 3.00572305 | 0.122371709 | 0.119515638 | 0 |
| GGGTCTGTCCACGATA-1 | 1569498 | 23022 | 169627 | 136331 | 1240518 | 401 | 0.036185353 | 3.796488767 | 3.130163983 | 0.091140917 | 0.09221101 | 0 |
| GGGTTGCGTGTCGCTG-1 | 1373334 | 39325 | 144117 | 121052 | 1068840 | 345 | 0.031163606 | 3.288426133 | 3.04180665 | 0.100745967 | 0.101794964 | 0 |
| GGTATTGAGGGTTCGA-1 | 2073240 | 38561 | 217346 | 185050 | 1632283 | 527 | 0.04768345 | 4.960465867 | 2.938348963 | 0.086203725 | 0.083793186 | 0 |
| GGTATTGAGGTTCCTA-1 | 1882342 | 39405 | 199714 | 170079 | 1473144 | 476 | 0.04297718 | 4.484453533 | 3.151106375 | 0.089310983 | 0.088926564 | 0 |
| GGTGAAGGTATTCGTG-1 | 1246628 | 34043 | 131859 | 101387 | 979339 | 316 | 0.028520823 | 3.0128921 | 3.044196698 | 0.108376062 | 0.10543602 | 0 |
| GGTGCGTCACTAACTG-1 | 738116 | 18923 | 73369 | 64911 | 580913 | 188 | 0.016917567 | 1.8065352 | 2.91571741 | 0.135975037 | 0.134444291 | 0 |
| GGTGCGTGTGTTTCAG-1 | 1834224 | 44300 | 190541 | 159274 | 1440109 | 465 | 0.042074139 | 4.405351667 | 3.03859213 | 0.090032357 | 0.089373655 | 0 |
| GGTGTTAGTCAGTGGA-1 | 1852090 | 28677 | 197814 | 167148 | 1458451 | 471 | 0.042593949 | 4.453754367 | 2.955483903 | 0.087557538 | 0.087974715 | 0 |
| GTAACGTAGGCAGTCA-1 | 2001772 | 72998 | 210064 | 171982 | 1546728 | 500 | 0.044934947 | 4.654238867 | 3.037518202 | 0.088666133 | 0.089078704 | 0 |
| GTAACGTAGTCCGTAT-1 | 1844214 | 14606 | 204022 | 149438 | 1476148 | 477 | 0.043127049 | 4.472399033 | 3.068077852 | 0.095639637 | 0.094890625 | 1 |
| GTAACGTTCCCACAAA-1 | 1116246 | 10907 | 122530 | 92843 | 889966 | 287 | 0.025963503 | 2.729980533 | 3.003813406 | 0.102803297 | 0.104546331 | 0 |
| GTAACTGAGCCGTCGT-1 | 1257814 | 30327 | 120442 | 113606 | 993439 | 321 | 0.028961561 | 3.066166533 | 3.025107752 | 0.108256055 | 0.105861299 | 0 |
| GTAACTGCATAGGGCA-1 | 1642580 | 59903 | 170003 | 151810 | 1260864 | 407 | 0.036683539 | 3.843016233 | 2.901381245 | 0.095753029 | 0.095336948 | 0 |
| GTAACTGGTAGATGCG-1 | 1634170 | 97216 | 167123 | 139436 | 1230395 | 397 | 0.035725585 | 3.743003933 | 3.124192013 | 0.095018641 | 0.096035094 | 0 |
| GTAACTGGTTGAACTC-1 | 2035178 | 34486 | 221429 | 183040 | 1596223 | 516 | 0.046497263 | 4.831227333 | 3.080909252 | 0.087264088 | 0.085633263 | 0 |
| GTAACTGTCGAATGTC-1 | 1770892 | 18470 | 194034 | 154829 | 1403559 | 453 | 0.04091448 | 4.2600948 | 3.144631621 | 0.091701272 | 0.090223346 | 0 |
| GTACGTACACTGGCCA-1 | 1606312 | 40276 | 168200 | 146002 | 1251834 | 404 | 0.036486075 | 3.842740067 | 2.968607732 | 0.097376631 | 0.09504945 | 0 |
| GTACGTAGTGCGCTTG-1 | 1263726 | 43782 | 132324 | 109365 | 978255 | 316 | 0.028513093 | 3.016493 | 3.091430114 | 0.103142521 | 0.102085807 | 0 |
| GTACTCCGTCAGTTTG-1 | 1334892 | 36216 | 142369 | 114123 | 1042184 | 337 | 0.030393146 | 3.194262633 | 2.956359014 | 0.104837605 | 0.102125765 | 0 |
| GTACTCCGTTGGTATC-1 | 2846454 | 21926 | 308439 | 252168 | 2263921 | 731 | 0.065921134 | 6.727206767 | 3.011988686 | 0.075101852 | 0.075586041 | 0 |
| GTACTTTAGAGTAATC-1 | 2239942 | 24738 | 240418 | 204154 | 1770632 | 572 | 0.051473221 | 5.326178367 | 2.985659187 | 0.089545725 | 0.087273998 | 0 |
| GTACTTTCAGGCCGTT-1 | 1991260 | 43421 | 215430 | 169752 | 1562657 | 505 | 0.045575805 | 4.7349636 | 3.089619358 | 0.084394794 | 0.084875202 | 0 |
| GTACTTTTCAGTACAC-1 | 2374966 | 20315 | 259198 | 218719 | 1876734 | 606 | 0.054634413 | 5.645589533 | 3.143260063 | 0.080331976 | 0.080864509 | 0 |
| GTAGGCCCACGTGGCT-1 | 2141392 | 62480 | 223524 | 190381 | 1665007 | 538 | 0.048486815 | 5.035866067 | 3.060673647 | 0.088463094 | 0.088252045 | 0 |
| GTAGGCCGTGTAATGA-1 | 1065974 | 20763 | 111345 | 85220 | 848646 | 274 | 0.024685543 | 2.614044367 | 3.001604692 | 0.109364782 | 0.110188677 | 0 |
| GTAGGCCTCCTTGCAC-1 | 2451642 | 27215 | 259484 | 221394 | 1943549 | 628 | 0.056639835 | 5.840827867 | 3.074964127 | 0.078540254 | 0.07719403 | 0 |
| GTAGGCCTCGTGCTAA-1 | 389230 | 2101 | 38896 | 32986 | 315247 | 102 | 0.009232303 | 1.000579067 | 3.077161765 | 0.188946594 | 0.188946594 | 0 |
| GTAGGCCTCTCGAGCG-1 | 1020332 | 90127 | 95815 | 88987 | 745403 | 241 | 0.021690516 | 2.3167042 | 2.968234359 | 0.118037919 | 0.115256473 | 0 |
| GTAGTCAGTCATTACG-1 | 1645666 | 40285 | 174907 | 144186 | 1286288 | 416 | 0.037559748 | 3.928586733 | 3.002614546 | 0.093161192 | 0.095377007 | 0 |
| GTAGTCATCGAACCCG-1 | 1257550 | 118777 | 123058 | 104485 | 911230 | 294 | 0.026486752 | 2.8071866 | 3.039002635 | 0.105174573 | 0.103161903 | 0 |
| GTATCTTAGGGAGTAA-1 | 1635648 | 14206 | 176172 | 137134 | 1308136 | 423 | 0.03822822 | 3.9960803 | 2.958546647 | 0.091547556 | 0.092472624 | 0 |
| GTATCTTAGGGATTTA-1 | 1875084 | 24057 | 201154 | 160999 | 1488874 | 481 | 0.043457073 | 4.5264354 | 3.073600455 | 0.090817347 | 0.090647063 | 0 |
| GTATCTTCATTTGTGG-1 | 1351348 | 31057 | 146734 | 115280 | 1058277 | 342 | 0.030847717 | 3.253622933 | 2.987731629 | 0.105623456 | 0.103280723 | 0 |
| GTATCTTGTCGCTTCT-1 | 1770066 | 88836 | 174652 | 154685 | 1351893 | 437 | 0.039283815 | 4.1061602 | 3.193944388 | 0.097796014 | 0.096691525 | 0 |
| GTATCTTGTCTCGGTG-1 | 774170 | 20818 | 83043 | 60777 | 609532 | 197 | 0.017853147 | 1.902446067 | 3.033195374 | 0.131273389 | 0.127993029 | 0 |
| GTATTCTCATAGGTCT-1 | 2330920 | 67938 | 241992 | 203999 | 1816991 | 587 | 0.052491338 | 5.367131367 | 2.960373672 | 0.087367284 | 0.08780864 | 0 |
| GTATTCTCATGCAGTT-1 | 1598848 | 25942 | 173238 | 132186 | 1267482 | 409 | 0.036961082 | 3.8502267 | 2.985837565 | 0.098873686 | 0.097289455 | 0 |
| GTCAAGTAGGACCAGT-1 | 1476490 | 32005 | 145279 | 133466 | 1165740 | 377 | 0.034050153 | 3.573816867 | 2.949946729 | 0.096644917 | 0.095370895 | 0 |
| GTCACAAAGTGAACAT-1 | 2453136 | 24743 | 269655 | 213325 | 1945413 | 628 | 0.056755193 | 5.8373346 | 3.096696052 | 0.077679055 | 0.076751249 | 0 |
| GTCACAAGTCAGTGGA-1 | 2014432 | 21212 | 221349 | 180384 | 1591487 | 514 | 0.046483077 | 4.835501367 | 3.22683554 | 0.085288719 | 0.085502515 | 0 |
| GTCACGGGTATTGAAG-1 | 1433324 | 26861 | 152281 | 122714 | 1131468 | 365 | 0.032958049 | 3.464893667 | 3.094852758 | 0.095394308 | 0.09775826 | 0 |
| GTCACGGTCCACAGAT-1 | 1775292 | 22176 | 192278 | 144423 | 1416415 | 458 | 0.041329321 | 4.301526767 | 3.031854372 | 0.092895443 | 0.089721 | 0 |
| GTCATTTAGCAAAGCC-1 | 1511770 | 18383 | 158710 | 143595 | 1191082 | 385 | 0.03458358 | 3.6389753 | 3.115326974 | 0.09546827 | 0.094547089 | 0 |
| GTCATTTGTCGCTTCT-1 | 1483466 | 35535 | 154085 | 131660 | 1162186 | 375 | 0.033861231 | 3.5679104 | 3.072048075 | 0.103044878 | 0.103044878 | 0 |
| GTCATTTTCCCTTATA-1 | 899814 | 14154 | 98075 | 73006 | 714579 | 231 | 0.020837688 | 2.211096267 | 2.9220865 | 0.12011275 | 0.120782399 | 0 |
| GTCCTCAGTTTCCTGC-1 | 3228202 | 30552 | 352233 | 282062 | 2563355 | 828 | 0.074799936 | 7.619784767 | 2.970329803 | 0.07108829 | 0.070112591 | 0 |
| GTCCTCATCTAAGACC-1 | 1913362 | 23407 | 203198 | 174860 | 1511897 | 488 | 0.044006035 | 4.565550367 | 3.142264467 | 0.090899084 | 0.089991499 | 0 |
| GTCGGGTCACAGCTGC-1 | 885556 | 25783 | 82545 | 79272 | 697956 | 225 | 0.020338418 | 2.1634484 | 3.060128649 | 0.124289448 | 0.124156093 | 0 |
| GTCTCGTCAGGTTAAA-1 | 696708 | 32079 | 72905 | 54023 | 537701 | 174 | 0.015603735 | 1.662614067 | 3.037659073 | 0.139063466 | 0.138186057 | 0 |
| GTCTCGTCATTCCGAA-1 | 1540622 | 18202 | 165333 | 134074 | 1223013 | 395 | 0.03564067 | 3.7383638 | 3.138160044 | 0.091928443 | 0.093000884 | 0 |
| GTCTCGTGTACGTGCC-1 | 1861646 | 14481 | 188363 | 165302 | 1493500 | 482 | 0.043445378 | 4.522677467 | 3.132970561 | 0.089542069 | 0.088852261 | 0 |
| GTCTCGTTCACACGGC-1 | 1198602 | 24020 | 129472 | 103753 | 941357 | 304 | 0.027268499 | 2.8834418 | 3.034634618 | 0.10723361 | 0.10700517 | 0 |
| GTCTCGTTCATTGCTT-1 | 2116490 | 48303 | 226083 | 178114 | 1663990 | 538 | 0.048463044 | 5.0273037 | 3.078546007 | 0.086222639 | 0.084601785 | 0 |
| GTCTTCGAGGTGCTAG-1 | 905402 | 8112 | 98515 | 77063 | 721712 | 233 | 0.021129132 | 2.238332867 | 3.04192025 | 0.122883327 | 0.122402448 | 0 |
| GTCTTCGAGTGTTGAA-1 | 1694516 | 22335 | 182912 | 144561 | 1344708 | 434 | 0.039234418 | 4.103284367 | 3.037509058 | 0.094878938 | 0.092133989 | 0 |
| GTCTTCGTCTGGTAGT-1 | 1632334 | 19253 | 176747 | 139880 | 1296454 | 419 | 0.037934373 | 3.9564849 | 3.104458318 | 0.094023561 | 0.094758434 | 0 |
| GTGAAGGGTCCTCGGA-1 | 1633264 | 28194 | 173116 | 146827 | 1285127 | 415 | 0.037477955 | 3.924708267 | 3.022338393 | 0.090832477 | 0.091343888 | 0 |
| GTGAAGGGTCTCGTCT-1 | 2115862 | 107611 | 214629 | 182046 | 1611576 | 521 | 0.046828531 | 4.8593122 | 3.112649541 | 0.08871914 | 0.088659693 | 0 |
| GTGCAGCAGCTGAAAT-1 | 1792186 | 39896 | 194953 | 152301 | 1405036 | 454 | 0.040969284 | 4.273322033 | 2.976950178 | 0.091532119 | 0.088308962 | 0 |
| GTGCAGCCAAAGTTAG-1 | 1039452 | 30172 | 110392 | 89865 | 809023 | 261 | 0.023610648 | 2.507332733 | 2.998707616 | 0.114358081 | 0.112205642 | 0 |
| GTGCAGCGTCCTACCT-1 | 1299694 | 58541 | 118854 | 111451 | 1010848 | 327 | 0.029395232 | 3.098637767 | 3.094415357 | 0.103374494 | 0.103597224 | 0 |
| GTGCATACACTCTTAT-1 | 1093226 | 21456 | 120444 | 89034 | 862292 | 279 | 0.025094953 | 2.653809033 | 2.975702435 | 0.11274585 | 0.108489259 | 0 |
| GTGCGGTCAATCGTCA-1 | 1711268 | 52186 | 177364 | 151508 | 1330210 | 430 | 0.03874931 | 4.0702481 | 2.939159096 | 0.093034038 | 0.094529476 | 0 |
| GTGCGGTCAGATCGAG-1 | 980174 | 38188 | 101130 | 84929 | 755927 | 244 | 0.0220431 | 2.3396341 | 3.136760633 | 0.128055952 | 0.124571636 | 0 |
| GTGCGGTGTACTCACA-1 | 1841358 | 25615 | 196108 | 167314 | 1452321 | 469 | 0.0423378 | 4.427631033 | 3.128866126 | 0.086874439 | 0.086366293 | 0 |
| GTGCTTCGTGATACCT-1 | 2252352 | 69269 | 230266 | 199143 | 1753674 | 566 | 0.051013328 | 5.282086033 | 3.224049617 | 0.101951109 | 0.100893579 | 1 |
| GTGGGTCCACAGAGCA-1 | 1481810 | 74488 | 150214 | 130563 | 1126545 | 364 | 0.032759416 | 3.452152367 | 3.016680988 | 0.099946927 | 0.098410379 | 0 |
| GTGGGTCGTACCACTA-1 | 1341734 | 12203 | 142424 | 114857 | 1072250 | 346 | 0.03121292 | 3.2835632 | 3.063260392 | 0.100444938 | 0.101593609 | 0 |
| GTGGGTCGTTGGAACG-1 | 1683602 | 94124 | 168932 | 144731 | 1275815 | 412 | 0.037139408 | 3.908941167 | 3.097200721 | 0.092640204 | 0.092640204 | 0 |
| GTGGGTCTCCGCTTGT-1 | 1747864 | 54716 | 181506 | 151849 | 1359793 | 439 | 0.039680098 | 4.145982733 | 3.073307616 | 0.091852387 | 0.093497703 | 0 |
| GTGGGTCTCCTAGACA-1 | 1730908 | 51546 | 181447 | 145675 | 1352240 | 437 | 0.039379205 | 4.118041333 | 3.044293862 | 0.09322998 | 0.092465077 | 0 |
| GTGTGCGAGAAGGTTT-1 | 1837266 | 71453 | 191840 | 152393 | 1421580 | 459 | 0.041434519 | 4.317740167 | 3.005703773 | 0.093993735 | 0.091291511 | 0 |
| GTGTGCGAGCGCGATA-1 | 1549748 | 26625 | 160136 | 136678 | 1226309 | 396 | 0.035800036 | 3.750629867 | 3.056336527 | 0.100554073 | 0.098854116 | 0 |
| GTGTGCGAGGCTCTCG-1 | 1552632 | 61860 | 160532 | 138087 | 1192153 | 385 | 0.034708134 | 3.645746167 | 3.119507934 | 0.093540976 | 0.093540976 | 0 |
| GTGTGCGGTCGCTTCT-1 | 607086 | 20047 | 46048 | 55563 | 485428 | 157 | 0.014076292 | 1.516978167 | 3.024836518 | 0.141466861 | 0.142390868 | 0 |
| GTTAAGCAGTGTTTGC-1 | 1961858 | 41027 | 210453 | 177675 | 1532703 | 495 | 0.044622561 | 4.6521443 | 3.048411154 | 0.085016591 | 0.084837747 | 0 |
| GTTAAGCCAGATATCC-1 | 1880652 | 53068 | 198457 | 171915 | 1457212 | 471 | 0.042061461 | 4.352649667 | 3.09974995 | 0.090632679 | 0.091308253 | 0 |
| GTTAAGCTCCACTTAT-1 | 1114700 | 13744 | 123878 | 92192 | 884886 | 286 | 0.025766818 | 2.715397 | 3.112393794 | 0.111342831 | 0.110824099 | 0 |
| GTTAAGCTCTGTCGAA-1 | 1699638 | 25331 | 184573 | 151630 | 1338104 | 432 | 0.039040787 | 4.083055567 | 3.080596018 | 0.096071151 | 0.09373875 | 0 |
| GTTACAGAGTACACCT-1 | 2109174 | 43594 | 227138 | 192571 | 1645871 | 532 | 0.047926819 | 4.9692936 | 3.036850743 | 0.086193198 | 0.084861352 | 0 |
| GTTACAGTCAGGTGAG-1 | 2256030 | 32381 | 243186 | 196696 | 1783767 | 576 | 0.052117528 | 5.403341767 | 3.081913397 | 0.081668922 | 0.082023423 | 0 |
| GTTCATTGTCAAGTTC-1 | 1998780 | 50666 | 213268 | 181905 | 1552941 | 502 | 0.044866572 | 4.620334567 | 3.113578563 | 0.09389273 | 0.09389273 | 1 |
| GTTCATTTCGAGTATC-1 | 3383148 | 24227 | 369239 | 295786 | 2693896 | 870 | 0.078524122 | 7.930237867 | 3.032839894 | 0.072799619 | 0.072260333 | 0 |
| GTTCATTTCGTAGGGA-1 | 1758616 | 17675 | 188914 | 151523 | 1400504 | 452 | 0.040978257 | 4.2880114 | 3.024010457 | 0.093978705 | 0.092574819 | 0 |

| | | | | | | | | | | | | |
|---|---|---|---|---|---|---|---|---|---|---|---|---|
| GTTCATTTCTCTTCCT-1 | 2500448 | 86148 | 264900 | 224229 | 1925171 | 622 | 0.056041184 | 5.783205433 | 3.054840062 | 0.079472687 | 0.079929142 | 0 |
| GTTCGGGTCAACAGTC-1 | 1619528 | 15864 | 177436 | 135603 | 1290625 | 417 | 0.037657843 | 3.933121167 | 3.114955674 | 0.090089994 | 0.088387205 | 0 |
| GTTCTCGCAACCCACG-1 | 1844272 | 82054 | 191909 | 161299 | 1409010 | 455 | 0.041003255 | 4.291125533 | 3.035637164 | 0.087873578 | 0.086646599 | 0 |
| GTTCTCGCACGCACCA-1 | 2383140 | 63509 | 249735 | 213140 | 1856756 | 600 | 0.05410296 | 5.606783233 | 3.044880357 | 0.080431506 | 0.077218934 | 0 |
| GTTCTCGCAGTGCTCG-1 | 1445760 | 43722 | 151910 | 129037 | 1121091 | 362 | 0.032537244 | 3.4317832 | 3.038235607 | 0.102041796 | 0.102041796 | 0 |
| GTTCTCGGTCATGTTG-1 | 1426946 | 21829 | 154751 | 113578 | 1136788 | 367 | 0.033140846 | 3.475283267 | 3.140984305 | 0.100905367 | 0.100045999 | 0 |
| GTTCTCGTCATGACCA-1 | 1926700 | 34403 | 204839 | 176308 | 1511150 | 488 | 0.044017541 | 4.585549333 | 3.066047787 | 0.08643448 | 0.087836924 | 0 |
| GTTTCTAAGGTGATTA-1 | 1474894 | 14792 | 160824 | 126010 | 1173268 | 379 | 0.03424198 | 3.591324633 | 3.137968256 | 0.102475823 | 0.101351876 | 0 |
| GTTTCTAAGTGGGATC-1 | 2129706 | 31555 | 228021 | 196455 | 1673675 | 541 | 0.048740485 | 5.046120867 | 3.139892582 | 0.086724879 | 0.085393309 | 0 |
| GTTTCTAGTCCGGTGT-1 | 2294738 | 38806 | 248070 | 208924 | 1798938 | 581 | 0.052448355 | 5.407919867 | 3.030339625 | 0.082791774 | 0.080337898 | 0 |
| GTTTCTATCAGATTCG-1 | 1549146 | 105927 | 156771 | 131347 | 1155101 | 373 | 0.033524796 | 3.5201264 | 3.133513245 | 0.100205931 | 0.10195152 | 0 |
| TAAACCGAGGCTAGAC-1 | 1606784 | 59255 | 172177 | 138724 | 1236628 | 399 | 0.03594455 | 3.772815167 | 2.786122119 | 0.100407888 | 0.095219047 | 0 |
| TAAACCGAGGTAGCCA-1 | 1883616 | 49185 | 201627 | 168819 | 1463985 | 473 | 0.042638881 | 4.459989833 | 3.038975707 | 0.088064812 | 0.089010031 | 0 |
| TAAACCGTCCAAGGGA-1 | 2052240 | 21689 | 222509 | 180116 | 1627926 | 526 | 0.047570188 | 4.944835367 | 2.973779552 | 0.085991536 | 0.082768288 | 0 |
| TAAGAGACAGCGTCAC-1 | 2021510 | 30838 | 207135 | 185908 | 1597629 | 516 | 0.046612458 | 4.8464496 | 3.114361211 | 0.087556081 | 0.085752913 | 0 |
| TAAGAGACATCTCGTC-1 | 1438314 | 27253 | 154199 | 123417 | 1133445 | 366 | 0.033001567 | 3.471796067 | 3.038906609 | 0.102228963 | 0.098373614 | 0 |
| TAAGAGAGTGTGTTCA-1 | 1709262 | 48093 | 177023 | 152680 | 1331466 | 430 | 0.03885601 | 4.071347267 | 3.046895533 | 0.096601904 | 0.093608064 | 0 |
| TAAGAGATCCAGGTTA-1 | 1870170 | 18035 | 202464 | 158301 | 1491370 | 482 | 0.043481416 | 4.532251567 | 3.013277543 | 0.092614958 | 0.088445214 | 0 |
| TAAGCGTTCATTGGAC-1 | 2005238 | 54013 | 213631 | 176753 | 1560841 | 504 | 0.045468 | 4.7339861 | 3.055968055 | 0.084665875 | 0.083192005 | 0 |
| TAAGTGCAGGGATTCG-1 | 1838360 | 112233 | 183584 | 161008 | 1381535 | 446 | 0.040140794 | 4.197646933 | 3.026109067 | 0.091156818 | 0.089988679 | 0 |
| TAAGTGCAGGTAAGTT-1 | 1846748 | 54539 | 194761 | 153129 | 1444319 | 467 | 0.042002948 | 4.360207933 | 3.067755798 | 0.089554369 | 0.091194077 | 0 |
| TAAGTGCTCGGGTCAC-1 | 2548824 | 33163 | 271000 | 227126 | 2017535 | 652 | 0.058840149 | 6.065913733 | 3.097083102 | 0.08068748 | 0.08098319 | 0 |
| TACACGACAAGCAAGC-1 | 2457252 | 51836 | 265080 | 215911 | 1924425 | 622 | 0.056110923 | 5.791251167 | 3.153943406 | 0.07861523 | 0.0795881 | 0 |
| TACACGACACTAGGCC-1 | 1840308 | 87270 | 191647 | 170053 | 1391338 | 449 | 0.040438959 | 4.237664967 | 3.072875241 | 0.089145616 | 0.089145616 | 0 |
| TACACGATCCGACCAG-1 | 1994292 | 59666 | 212915 | 177214 | 1544497 | 499 | 0.045021029 | 4.688506733 | 3.046838258 | 0.081193008 | 0.082048943 | 0 |
| TACACGATCCGTTCAC-1 | 2617154 | 29936 | 283810 | 235621 | 2067787 | 668 | 0.060311595 | 6.189951833 | 2.953940264 | 0.075792921 | 0.075739094 | 0 |
| TACAGTGAGTCCCGAC-1 | 1573904 | 59124 | 165585 | 137046 | 1212149 | 392 | 0.035265247 | 3.698016633 | 2.903111926 | 0.095695339 | 0.093762183 | 0 |
| TACAGTGGTCAATTCG-1 | 1377138 | 127317 | 133646 | 118511 | 997664 | 322 | 0.028914494 | 3.0519058 | 3.119232818 | 0.102008863 | 0.101485807 | 0 |
| TACAGTGGTCTAACGT-1 | 1992352 | 36011 | 210619 | 173128 | 1572594 | 508 | 0.045860036 | 4.769692033 | 3.036585886 | 0.084667341 | 0.08483458 | 0 |
| TACCTATAGTAGGTGC-1 | 1772926 | 43195 | 185705 | 156623 | 1387403 | 448 | 0.040257508 | 4.212792533 | 3.141638492 | 0.093183879 | 0.090590427 | 0 |
| TACCTATGTAAACAGT-1 | 1884048 | 16806 | 205766 | 172576 | 1488900 | 481 | 0.043443777 | 4.5236024 | 3.086331804 | 0.093790943 | 0.092250193 | 0 |
| TACCTATGTCAGTTTG-1 | 1471908 | 29892 | 159045 | 125828 | 1157143 | 374 | 0.033721213 | 3.546665267 | 3.021670111 | 0.095727009 | 0.095727009 | 0 |
| TACCTATGTTGAACAA-1 | 1627134 | 21265 | 177394 | 140625 | 1287850 | 416 | 0.037556918 | 3.933166733 | 3.011619731 | 0.097503679 | 0.09541239 | 0 |
| TACCTTAAGACTGGCA-1 | 1965718 | 20795 | 212826 | 180607 | 1551490 | 501 | 0.045212203 | 4.710638433 | 2.91538877 | 0.086503626 | 0.086790213 | 0 |
| TACCTTAAGGAGCGAG-1 | 1813566 | 45494 | 188916 | 169000 | 1410156 | 456 | 0.04113946 | 4.285842167 | 3.034252024 | 0.090067254 | 0.088637981 | 0 |
| TACCTTAAGTGAATTG-1 | 1447110 | 19203 | 157720 | 119833 | 1150354 | 372 | 0.033577584 | 3.525355633 | 3.130051501 | 0.102142266 | 0.10000367 | 0 |
| TACCTTACATTCCGAA-1 | 1798922 | 17481 | 198853 | 154623 | 1427965 | 461 | 0.041605488 | 4.3375444 | 2.957831197 | 0.093500743 | 0.08766323 | 0 |
| TACCTTATCACTCATT-1 | 1841270 | 18579 | 202226 | 155135 | 1465330 | 473 | 0.042781928 | 4.432972867 | 2.957391953 | 0.088339479 | 0.08474174 | 0 |
| TACGGATAGCTGAACG-1 | 1543264 | 89785 | 156329 | 134002 | 1163148 | 376 | 0.033824508 | 3.561573533 | 2.972797523 | 0.102887576 | 0.100850955 | 0 |
| TACGGATAGTGTTAGA-1 | 1138328 | 12417 | 121299 | 99383 | 905229 | 292 | 0.026508456 | 2.806755833 | 2.960549677 | 0.109238983 | 0.110520069 | 0 |
| TACGGATCAACACCGC-1 | 1280980 | 23555 | 132893 | 119139 | 1005393 | 325 | 0.02924133 | 3.092972833 | 2.979622856 | 0.104494799 | 0.104209962 | 0 |
| TACGGATCAGAGGCAT-1 | 2658116 | 37968 | 284234 | 235997 | 2099917 | 678 | 0.06131188 | 6.292786933 | 3.037294613 | 0.076025963 | 0.075843835 | 0 |
| TACGGATCATCCGTTC-1 | 2062042 | 34193 | 214906 | 180673 | 1632270 | 527 | 0.047324493 | 4.8944453 | 3.076870047 | 0.10770914 | 0.103634728 | 1 |
| TACGGATGTGATGTTC-1 | 2629786 | 57253 | 279746 | 229188 | 2063599 | 667 | 0.05976055 | 6.059645633 | 3.129416864 | 0.078813164 | 0.080272963 | 0 |
| TACGGATTCAAGCCGC-1 | 1825626 | 16398 | 192748 | 168876 | 1447604 | 468 | 0.042191896 | 4.408129233 | 3.106567684 | 0.090289182 | 0.092089561 | 0 |
| TACGGATTCACACGGC-1 | 1225208 | 76669 | 124422 | 106882 | 917235 | 296 | 0.026598068 | 2.8251283 | 2.98828752 | 0.112535635 | 0.10789928 | 0 |
| TACGGGCAGGTAAGTT-1 | 1748176 | 40338 | 184325 | 150730 | 1372783 | 443 | 0.039992115 | 4.161671633 | 3.14508556 | 0.091692189 | 0.090903422 | 0 |
| TACGGGCGTATCTCTT-1 | 1818566 | 91090 | 186033 | 156396 | 1385047 | 447 | 0.040334224 | 4.194349767 | 3.059606506 | 0.093338711 | 0.092352899 | 0 |
| TACGGTAAGAACAATC-1 | 2395126 | 40184 | 253628 | 216884 | 1884430 | 609 | 0.054874113 | 5.6413592 | 3.076647727 | 0.085404633 | 0.084261454 | 0 |
| TACGGTAAGTTTCCTT-1 | 2008894 | 34502 | 214339 | 184723 | 1575330 | 509 | 0.045894132 | 4.765208333 | 2.91291624 | 0.090336247 | 0.089275258 | 0 |
| TACGGTAGTGCTTTCC-1 | 2188622 | 90951 | 223078 | 199169 | 1675424 | 541 | 0.048630725 | 5.019006167 | 3.115155648 | 0.087836053 | 0.085980937 | 0 |
| TACGGTATCTCGTGGG-1 | 2254158 | 32351 | 238433 | 200306 | 1783068 | 576 | 0.052069543 | 5.391689933 | 3.046808964 | 0.080859965 | 0.081305074 | 0 |
| TACTCATAGAACAATC-1 | 1908910 | 21588 | 203637 | 174814 | 1508871 | 487 | 0.043997999 | 4.566169267 | 3.111281581 | 0.08930226 | 0.089589834 | 0 |
| TACTCATAGAAGGTGA-1 | 2744076 | 18788 | 297741 | 219775 | 2207772 | 713 | 0.064471205 | 6.616008167 | 2.725081905 | 0.110486014 | 0.111082072 | 1 |
| TACTCATAGGCATGGT-1 | 1899894 | 12973 | 207588 | 164357 | 1514976 | 489 | 0.044270956 | 4.593671667 | 3.061652351 | 0.09136594 | 0.086192342 | 0 |
| TACTCATCAACGGCTC-1 | 1529160 | 40457 | 161249 | 147802 | 1179652 | 381 | 0.034354423 | 3.613087533 | 3.018038235 | 0.09930974 | 0.097187797 | 0 |
| TACTCGCAGAAGGTGA-1 | 1807448 | 15870 | 198431 | 158835 | 1434312 | 463 | 0.04188882 | 4.381434533 | 3.055011261 | 0.092710832 | 0.090939921 | 0 |
| TACTCGCAGGACGAAA-1 | 1696788 | 11323 | 187001 | 142709 | 1355755 | 438 | 0.039641256 | 4.1196104 | 3.253370192 | 0.130680165 | 0.130680165 | 1 |
| TACTCGCCATGATGCT-1 | 2694362 | 67691 | 286408 | 237647 | 2102616 | 679 | 0.060945835 | 6.1905062 | 2.927095619 | 0.082892106 | 0.082452461 | 0 |
| TACTCGCTCCACGATA-1 | 1660108 | 9588 | 184221 | 138339 | 1327960 | 429 | 0.038806552 | 4.053344067 | 2.973555234 | 0.096613646 | 0.091560813 | 0 |
| TACTTGTAGTGGGTAC-1 | 2342126 | 50090 | 242355 | 206085 | 1843596 | 596 | 0.053615955 | 5.5212529 | 3.072042957 | 0.084584144 | 0.082268447 | 0 |
| TACTTGTCAGACACAG-1 | 1152354 | 19407 | 124664 | 100750 | 907533 | 293 | 0.026414836 | 2.7858613 | 3.098120389 | 0.110917178 | 0.109883986 | 0 |
| TACTTGTGTAGGTTTC-1 | 2237696 | 60061 | 225643 | 193912 | 1758080 | 568 | 0.0510299 | 5.245183733 | 3.0954151 | 0.093755519 | 0.092766627 | 1 |
| TACTTGTGTTCGGTGC-1 | 1586448 | 103650 | 159654 | 138337 | 1184807 | 383 | 0.034314209 | 3.6003808 | 3.020198577 | 0.102745888 | 0.101117904 | 0 |
| TACTTGTTCACCGGTG-1 | 1472560 | 24510 | 160367 | 122798 | 1164885 | 376 | 0.033907372 | 3.5579543 | 3.18585045 | 0.096535558 | 0.095476795 | 0 |
| TAGACCAAGCGGCTTC-1 | 2639720 | 51525 | 286484 | 223094 | 2078617 | 671 | 0.060585722 | 6.2218663 | 3.964392388 | 0.132129298 | 0.132129298 | 1 |
| TAGACCAAGTCGATAA-1 | 1448302 | 12233 | 159938 | 118882 | 1157249 | 374 | 0.03375581 | 3.535116533 | 3.123943522 | 0.101020814 | 0.098925996 | 0 |
| TAGACCACATCCACGG-1 | 1388392 | 40812 | 149503 | 120884 | 1077193 | 348 | 0.031385537 | 3.308094767 | 3.035295538 | 0.100293418 | 0.09941708 | 0 |
| TAGACCACATGATGCT-1 | 2362116 | 63280 | 254571 | 209409 | 1834856 | 593 | 0.053045659 | 5.3957752 | 3.142151778 | 0.08738112 | 0.086219694 | 0 |
| TAGACCATCATCAGCA-1 | 1735454 | 35589 | 186189 | 151578 | 1362098 | 440 | 0.039643078 | 4.1472061 | 3.017394084 | 0.094260041 | 0.091597755 | 0 |
| TAGAGCTAGCTTTCAG-1 | 1671676 | 23320 | 178206 | 154165 | 1315985 | 425 | 0.038448005 | 4.016525633 | 3.133997557 | 0.092198778 | 0.090791697 | 0 |
| TAGAGCTAGGCTAGGT-1 | 1522574 | 17114 | 164059 | 126546 | 1214855 | 392 | 0.0354429 | 3.709278967 | 3.075013676 | 0.091171922 | 0.091415476 | 0 |
| TAGAGCTCACGTCGGT-1 | 1873838 | 53519 | 190033 | 167370 | 1462916 | 473 | 0.042637764 | 4.452796133 | 3.025882336 | 0.09663332 | 0.09663332 | 1 |
| TAGAGCTCATTGGTCA-1 | 1798664 | 52476 | 186267 | 160402 | 1399519 | 452 | 0.040591388 | 4.227031267 | 3.04577002 | 0.091616179 | 0.090952221 | 0 |
| TAGAGCTTCTCAGATG-1 | 1066822 | 16807 | 115506 | 85641 | 848868 | 274 | 0.024754702 | 2.615024067 | 3.084252986 | 0.118236248 | 0.111015305 | 0 |
| TAGCCGGAGTCGTGTT-1 | 1719226 | 35853 | 165496 | 144869 | 1373008 | 444 | 0.039965106 | 4.166606 | 3.086648676 | 0.093591441 | 0.092444112 | 0 |
| TAGCCGGCAACCTCAA-1 | 1699768 | 14337 | 185035 | 151693 | 1348703 | 436 | 0.039355208 | 4.099660567 | 2.997223603 | 0.094315456 | 0.09076989 | 0 |
| TAGCCGGCACCACACG-1 | 1613736 | 87616 | 166716 | 138992 | 1220412 | 394 | 0.035466549 | 3.7289898 | 3.14193695 | 0.097713836 | 0.097675164 | 0 |
| TAGCCGGCACGCTGAC-1 | 2149056 | 31516 | 233674 | 198499 | 1685367 | 544 | 0.049138322 | 5.098764367 | 3.077626447 | 0.080284416 | 0.080484712 | 0 |
| TAGCCGGGTTCCACAA-1 | 1720700 | 14396 | 190206 | 144618 | 1371480 | 443 | 0.039981265 | 4.171733933 | 3.121113534 | 0.092645728 | 0.092968793 | 0 |
| TAGCCGGTCAGCGGCT-1 | 1652736 | 16042 | 180987 | 153479 | 1302228 | 421 | 0.03794329 | 3.977796667 | 3.038023057 | 0.093708298 | 0.092433383 | 0 |
| TAGGCATAGCTAAACA-1 | 1880108 | 43011 | 201242 | 168380 | 1467475 | 474 | 0.042739127 | 4.472179133 | 3.08244354 | 0.091694569 | 0.090843795 | 0 |
| TAGGCATAGTAGGTGC-1 | 1831152 | 92209 | 187097 | 160619 | 1391227 | 449 | 0.040388379 | 4.221929433 | 2.938904674 | 0.092317314 | 0.093281819 | 0 |
| TAGGCATAGTGAACAT-1 | 2297936 | 34347 | 247848 | 205435 | 1810306 | 585 | 0.052880005 | 5.464162867 | 3.01296395 | 0.081594676 | 0.081895536 | 0 |
| TAGGCATCAATCGTCA-1 | 1677626 | 46443 | 176817 | 150514 | 1303852 | 421 | 0.037974718 | 3.980201733 | 3.142045172 | 0.090782586 | 0.09034173 | 0 |
| TAGGCATGTCGGCATC-1 | 2017848 | 37585 | 213233 | 186154 | 1580876 | 511 | 0.046125569 | 4.8110013 | 3.015569619 | 0.08840677 | 0.088232286 | 0 |
| TAGGCATGTTCAGTAC-1 | 2361030 | 67512 | 248871 | 209321 | 1835326 | 593 | 0.053313917 | 5.4729078 | 3.049120938 | 0.083683044 | 0.0838148 | 0 |
| TAGTGGTCAAGTGTTC-1 | 2377520 | 66659 | 247470 | 209144 | 1854247 | 599 | 0.053917522 | 5.554775167 | 3.076293465 | 0.082902497 | 0.081753669 | 0 |
| TAGTGGTCACACACGC-1 | 1573220 | 23809 | 169762 | 143012 | 1236637 | 399 | 0.036001704 | 3.7807721 | 3.010343921 | 0.097535314 | 0.092951225 | 0 |
| TAGTGGTGTGTGACAG-1 | 1471092 | 26684 | 157461 | 125412 | 1161535 | 375 | 0.033869839 | 3.560299633 | 3.002350635 | 0.100299748 | 0.098285711 | 0 |
| TAGTTGGAGAATGGGA-1 | 2231870 | 17582 | 242619 | 184770 | 1786899 | 577 | 0.052227332 | 5.388131367 | 3.448329553 | 0.12445789 | 0.12445789 | 1 |
| TAGTTGGCATGAGAAT-1 | 2274300 | 27607 | 247737 | 194937 | 1804019 | 583 | 0.052701432 | 5.4342 | 2.961995823 | 0.087861884 | 0.083238694 | 0 |
| TAGTTGGCATGGCACC-1 | 2029112 | 37042 | 217042 | 184565 | 1590463 | 514 | 0.046349106 | 4.8279328 | 3.038451156 | 0.085264114 | 0.086804847 | 0 |
| TAGTTGGTCCTGATTT-1 | 2128642 | 77062 | 224142 | 176803 | 1650635 | 533 | 0.048034296 | 4.967066 | 2.841439765 | 0.08779661 | 0.08779661 | 0 |

| | | | | | | | | | | | | |
|---|---|---|---|---|---|---|---|---|---|---|---|---|
| TAGTTGGTCGAACCCG-1 | 978608 | 58920 | 97952 | 87306 | 734430 | 237 | 0.021372209 | 2.285116233 | 3.120054812 | 0.123919155 | 0.123904924 | 0 |
| TATCAGGAGTCCGTAT-1 | 1895928 | 20292 | 204965 | 156337 | 1514334 | 489 | 0.044208812 | 4.579943567 | 3.068984375 | 0.088408242 | 0.085964452 | 0 |
| TATCAGGGTGACCTGC-1 | 2035436 | 24126 | 229256 | 178225 | 1603829 | 518 | 0.046758622 | 4.853525367 | 3.12536936 | 0.102896559 | 0.100725232 | 1 |
| TATCAGGGTTCGAGCC-1 | 1194712 | 78532 | 117765 | 107548 | 890867 | 288 | 0.02588351 | 2.742308967 | 3.020123125 | 0.111574242 | 0.108808232 | 0 |
| TATCAGGGTTCGTCAA-1 | 2132816 | 24665 | 229122 | 188592 | 1690437 | 546 | 0.049316887 | 5.120336267 | 2.989728655 | 0.087162189 | 0.08686263 | 0 |
| TATCAGGTCTGGTAGT-1 | 1484660 | 20849 | 159410 | 129624 | 1174777 | 379 | 0.034359066 | 3.600746067 | 3.061210141 | 0.0978681 | 0.098157159 | 0 |
| TATCAGGTCTTCTTGA-1 | 1070870 | 12394 | 115449 | 94067 | 848960 | 274 | 0.0247734 | 2.628853433 | 3.114960898 | 0.112447556 | 0.112189801 | 0 |
| TATCTCAAGACTGGGT-1 | 1943192 | 18108 | 211125 | 166154 | 1547805 | 500 | 0.045228185 | 4.700175433 | 3.026383837 | 0.088079902 | 0.088749325 | 0 |
| TATCTCAGTAGTGGCA-1 | 1987226 | 47552 | 210026 | 176405 | 1553243 | 502 | 0.045205922 | 4.706531667 | 3.03515428 | 0.086128461 | 0.085422059 | 0 |
| TATCTCAGTCCGAAGA-1 | 1332600 | 16196 | 144456 | 119894 | 1052054 | 340 | 0.03068444 | 3.2394976 | 3.208025283 | 0.102858496 | 0.100544042 | 0 |
| TATCTCAGTGACCCGT-1 | 2399688 | 35833 | 255230 | 220159 | 1888466 | 610 | 0.055107937 | 5.696481633 | 3.106650944 | 0.079152825 | 0.079616981 | 0 |
| TATGCCCAGGGCTTGA-1 | 1193432 | 8955 | 128593 | 108072 | 947812 | 306 | 0.027686248 | 2.9351697 | 3.145415936 | 0.104360195 | 0.103989722 | 0 |
| TATGCCCTCTAGAACC-1 | 2119420 | 57842 | 221612 | 193340 | 1646626 | 532 | 0.047852412 | 4.951597333 | 3.033173504 | 0.087337155 | 0.087736099 | 0 |
| TATTACCAGCCAGGAT-1 | 2357222 | 31599 | 254160 | 201168 | 1870295 | 604 | 0.054578677 | 5.602899433 | 3.215561034 | 0.07885708 | 0.07885708 | 0 |
| TATTACCGTGCTATTG-1 | 1068362 | 14236 | 116007 | 81020 | 857099 | 277 | 0.025011776 | 2.639855933 | 3.036052349 | 0.113729029 | 0.114407409 | 0 |
| TCAACGAAGAGCAATT-1 | 1514202 | 40925 | 163230 | 126395 | 1183652 | 382 | 0.03452396 | 3.6203681 | 3.006697443 | 0.09608982 | 0.098813516 | 0 |
| TCAACGAAGCTGCACC-1 | 2731640 | 55908 | 295631 | 239785 | 2140316 | 691 | 0.062323655 | 6.381016233 | 3.261517586 | 0.101217011 | 0.101217011 | 1 |
| TCAACGAAGGCTGCTC-1 | 2375976 | 35402 | 259367 | 212945 | 1868262 | 604 | 0.054500182 | 5.639611933 | 3.142790149 | 0.078940415 | 0.077403796 | 0 |
| TCAACGATCTTTCGCG-1 | 1006886 | 87701 | 99167 | 86887 | 733131 | 237 | 0.021293752 | 2.2659808 | 3.000205948 | 0.124325758 | 0.122997971 | 0 |
| TCACAAGCATTTCCAT-1 | 2428758 | 33245 | 268139 | 218792 | 1908582 | 617 | 0.055739015 | 5.735632467 | 3.100528048 | 0.079620771 | 0.079616424 | 0 |
| TCACGAAGTCAAACTC-1 | 1883808 | 59643 | 200888 | 170401 | 1452876 | 469 | 0.042276497 | 4.414824933 | 3.03380994 | 0.088892474 | 0.090075376 | 0 |
| TCACGAAGTGTCTAAC-1 | 2385080 | 20024 | 267184 | 204313 | 1893559 | 612 | 0.05527542 | 5.6887297 | 3.027020308 | 0.080188933 | 0.079646573 | 0 |
| TCAGATGAGAGCCCAA-1 | 1782712 | 12819 | 193261 | 160107 | 1416525 | 458 | 0.041356747 | 4.325691467 | 3.110492292 | 0.090396302 | 0.089239886 | 0 |
| TCAGATGTCCGATACA-1 | 2234366 | 32645 | 242764 | 200009 | 1758948 | 568 | 0.051348646 | 5.3113818 | 3.060627371 | 0.083400134 | 0.082244677 | 0 |
| TCAGCAAGTAGATGCG-1 | 1717858 | 69067 | 180703 | 149643 | 1318445 | 426 | 0.038298375 | 4.013314867 | 3.17336828 | 0.09204946 | 0.09200351 | 0 |
| TCAGCAAGTGTTTCAG-1 | 2142000 | 33022 | 234487 | 188049 | 1686442 | 545 | 0.049281704 | 5.1203846 | 3.008899274 | 0.080966075 | 0.082010651 | 0 |
| TCAGCAAGTTGGTGAG-1 | 1805496 | 23764 | 196511 | 154542 | 1430679 | 462 | 0.041852428 | 4.365999 | 3.049341498 | 0.084281508 | 0.08582369 | 0 |
| TCAGCTCAGTTGTCGT-1 | 2409122 | 39697 | 259176 | 217743 | 1892506 | 611 | 0.055219606 | 5.701687033 | 3.022730663 | 0.078378422 | 0.078378422 | 0 |
| TCAGGATTCAGCAGCC-1 | 1890788 | 73172 | 198775 | 163177 | 1455664 | 470 | 0.042325748 | 4.4177091 | 3.23887997 | 0.112427382 | 0.111311027 | 1 |
| TCAGGATTCGGGTCAC-1 | 2536316 | 35436 | 267269 | 229468 | 2004143 | 647 | 0.058426383 | 6.009323133 | 2.941340703 | 0.077788918 | 0.077788918 | 0 |
| TCAGGTACATAGACAA-1 | 1369112 | 11427 | 151520 | 116338 | 1089827 | 352 | 0.03180681 | 3.335566733 | 3.095461292 | 0.104877998 | 0.101074052 | 0 |
| TCAGGTACATGTAGCT-1 | 1784354 | 87745 | 185048 | 153143 | 1358418 | 439 | 0.039373614 | 4.099629967 | 3.119308615 | 0.091695668 | 0.092316008 | 0 |
| TCAGGTATCAAGTGCT-1 | 2482604 | 74659 | 261254 | 221828 | 1924863 | 622 | 0.056058553 | 5.784886067 | 3.167909918 | 0.079709086 | 0.079301662 | 0 |
| TCATTACAGCCAGTTT-1 | 1449460 | 59268 | 154061 | 119913 | 1116218 | 361 | 0.032456195 | 3.400645067 | 3.072161068 | 0.099878446 | 0.100094104 | 0 |
| TCATTACCACAGACCC-1 | 1703842 | 60908 | 177942 | 154422 | 1310570 | 423 | 0.038107208 | 3.9986053 | 3.01047801 | 0.093693184 | 0.094433549 | 0 |
| TCATTACCATTCAGTG-1 | 1093652 | 11949 | 121086 | 85014 | 875603 | 283 | 0.025496159 | 2.674225433 | 2.993075825 | 0.115836704 | 0.110648578 | 0 |
| TCATTACTCTCCACAC-1 | 2153296 | 53154 | 229852 | 189234 | 1681056 | 543 | 0.048962721 | 5.075311633 | 3.059659616 | 0.082729944 | 0.083647912 | 0 |
| TCATTTGAGTAGCCGA-1 | 1970502 | 13022 | 212363 | 182168 | 1562949 | 505 | 0.045572831 | 4.751489233 | 3.107867371 | 0.089203524 | 0.089203524 | 0 |
| TCATTTGAGTCTCGTA-1 | 1454064 | 13237 | 158614 | 119732 | 1162481 | 376 | 0.033913925 | 3.557729567 | 3.079127496 | 0.104366262 | 0.099789953 | 0 |
| TCCACACCAAGCAAGC-1 | 1987106 | 41680 | 213393 | 181299 | 1550734 | 501 | 0.045166804 | 4.695992267 | 2.93807359 | 0.086718991 | 0.08713187 | 0 |
| TCCACACGTGCTCGAC-1 | 1279132 | 49309 | 133570 | 120258 | 975995 | 315 | 0.028389809 | 3.0005559 | 3.06282187 | 0.109443716 | 0.103326904 | 0 |
| TCCACACTCGCAACGC-1 | 1622502 | 27952 | 174283 | 146153 | 1274114 | 412 | 0.037075865 | 3.886508867 | 3.168780058 | 0.089282531 | 0.091521986 | 0 |
| TCCCGATAGAATCGTA-1 | 1714450 | 11802 | 186425 | 147506 | 1368717 | 442 | 0.039944936 | 4.179425267 | 3.309900273 | 0.088813272 | 0.088697814 | 0 |
| TCCCGATAGTGCGATG-1 | 1560746 | 16818 | 170267 | 129241 | 1244420 | 402 | 0.036354768 | 3.811432933 | 3.140919811 | 0.092670414 | 0.093716336 | 0 |
| TCCCGATGTTAAGTAG-1 | 1467288 | 14425 | 161276 | 117597 | 1173990 | 379 | 0.034368288 | 3.601870367 | 3.107809781 | 0.10321635 | 0.097594892 | 0 |
| TCGAGGCAGATCCTGT-1 | 2022518 | 26861 | 214784 | 175609 | 1605264 | 519 | 0.046851795 | 4.868547367 | 3.020345967 | 0.08244857 | 0.082337718 | 0 |
| TCGAGGCCACCGAACC-1 | 1251800 | 57789 | 119127 | 110570 | 964314 | 312 | 0.028023378 | 2.963238567 | 2.847185729 | 0.107184538 | 0.106051786 | 0 |
| TCGAGGCCAGTGAAAT-1 | 1564870 | 31340 | 161170 | 130508 | 1241852 | 401 | 0.036270595 | 3.787356767 | 2.995887346 | 0.094687357 | 0.092600434 | 0 |
| TCGAGGCGTAAGAAGG-1 | 831570 | 14085 | 84042 | 69121 | 664322 | 215 | 0.019427947 | 2.073578667 | 2.511854971 | 0.120572043 | 0.121557928 | 1 |
| TCGCGAGAGAGTTGGC-1 | 1757016 | 40278 | 170395 | 156789 | 1389554 | 449 | 0.040552641 | 4.247035667 | 2.960811473 | 0.092739676 | 0.091218985 | 0 |
| TCGCGAGGTGATCCTA-1 | 1704922 | 29476 | 169892 | 151277 | 1354277 | 437 | 0.039528013 | 4.1386832 | 3.033847268 | 0.093346152 | 0.093369258 | 0 |
| TCGCGAGGTTCAAGAA-1 | 918630 | 13371 | 91959 | 78718 | 734582 | 237 | 0.021331567 | 2.2744739 | 3.031315685 | 0.117110944 | 0.116227317 | 0 |
| TCGCGAGTCCGCTTGT-1 | 1499842 | 21251 | 157210 | 130860 | 1190521 | 385 | 0.034740065 | 3.646671833 | 3.048385379 | 0.095924028 | 0.096760376 | 0 |
| TCGCGAGTCTACGGCG-1 | 869502 | 33313 | 83893 | 76510 | 675786 | 218 | 0.019629586 | 2.0963251 | 2.958857029 | 0.123615223 | 0.125576355 | 0 |
| TCGCGTTAGACTGGGT-1 | 1524144 | 18854 | 150819 | 126856 | 1227615 | 397 | 0.03574705 | 3.74515 | 3.123850865 | 0.092588765 | 0.094887985 | 0 |
| TCGCGTTAGGACGAAA-1 | 1245744 | 13566 | 120686 | 108300 | 1003192 | 324 | 0.029176966 | 3.078576567 | 3.110922484 | 0.102602023 | 0.104752288 | 0 |
| TCGCGTTCAACAACTC-1 | 1240408 | 21638 | 118648 | 111291 | 988831 | 319 | 0.028765721 | 3.0411341 | 3.041434659 | 0.103263449 | 0.106729534 | 0 |
| TCGCGTTGTCGCGAAA-1 | 1112142 | 9546 | 123767 | 98794 | 880035 | 284 | 0.025655342 | 2.7211949 | 3.114356942 | 0.114966438 | 0.110097885 | 0 |
| TCGCGTTGTCTAGCGC-1 | 779442 | 11718 | 73885 | 71662 | 622177 | 201 | 0.018085483 | 1.938495933 | 2.96523937 | 0.127377853 | 0.128619531 | 0 |
| TCGCGTTTCCGTATTT-1 | 809016 | 33189 | 78846 | 59682 | 637299 | 206 | 0.018570187 | 1.966083267 | 3.564289214 | 0.168044553 | 0.170914663 | 1 |
| TCGGGACAGTCAAGGC-1 | 1443188 | 68931 | 141784 | 125941 | 1106532 | 357 | 0.032192986 | 3.3986331 | 3.106600283 | 0.100921843 | 0.100921843 | 0 |
| TCGGGACGTTTCCATT-1 | 1709586 | 32267 | 177054 | 148644 | 1351621 | 437 | 0.039403376 | 4.115725867 | 3.094161771 | 0.090570369 | 0.090576515 | 0 |
| TCGGGACTCGTAAGGG-1 | 815546 | 15420 | 81958 | 68581 | 649587 | 210 | 0.019017113 | 2.020077967 | 3.038223118 | 0.129631172 | 0.124338115 | 0 |
| TCGGTAAGTATACACC-1 | 1902158 | 33803 | 196881 | 169280 | 1502194 | 485 | 0.043679214 | 4.553322867 | 3.015438063 | 0.091583938 | 0.090325061 | 0 |
| TCGGTAATCAGAAACA-1 | 1452842 | 30640 | 148145 | 121939 | 1152118 | 372 | 0.03362408 | 3.535642933 | 3.662500089 | 0.100260321 | 0.100260321 | 0 |
| TCGTACCAATCCATG-1 | 703032 | 15761 | 65858 | 59422 | 561991 | 182 | 0.016336549 | 1.751279967 | 3.024922428 | 0.134192991 | 0.13902432 | 0 |
| TCGTACCCACGTTTAG-1 | 467408 | 10349 | 43615 | 39923 | 373521 | 121 | 0.010898856 | 1.1791157 | 3.022875399 | 0.165467634 | 0.158921586 | 0 |
| TCGTACCGTGTGACAG-1 | 713804 | 12504 | 67684 | 61364 | 572252 | 185 | 0.01671725 | 1.787778 | 3.106399898 | 0.135132332 | 0.134120751 | 0 |
| TCGTAGACAATCCGTA-1 | 757554 | 7956 | 74918 | 67610 | 607070 | 196 | 0.017674093 | 1.8922723 | 3.091573895 | 0.132244439 | 0.130803199 | 0 |
| TCGTAGAGTGACTGAG-1 | 1249064 | 18577 | 124440 | 102268 | 1003779 | 324 | 0.02925285 | 3.0885475 | 3.00949837 | 0.103444751 | 0.101652038 | 0 |
| TCTATTGCAACGAAAT-1 | 1550156 | 27309 | 166373 | 136574 | 1219900 | 394 | 0.035554697 | 3.725597367 | 3.170744178 | 0.092663719 | 0.094544823 | 0 |
| TCTATTGCAGAGATCG-1 | 1491700 | 63832 | 152949 | 131109 | 1143810 | 369 | 0.033250082 | 3.506239733 | 3.08438432 | 0.100770089 | 0.099524503 | 0 |
| TCTATTGCATGCGTGC-1 | 1463972 | 46563 | 152919 | 133660 | 1130830 | 365 | 0.032846115 | 3.448698733 | 3.09257181 | 0.099286243 | 0.097966923 | 0 |
| TCTATTGGTATTCGCA-1 | 1485748 | 41330 | 154350 | 130428 | 1159640 | 375 | 0.03373445 | 3.5473501 | 3.050445947 | 0.09892392 | 0.099207901 | 0 |
| TCTATTGGTTGGCGTC-1 | 1309892 | 56782 | 129539 | 118581 | 1004990 | 325 | 0.029286836 | 3.1010971 | 3.02418341 | 0.102263945 | 0.101759004 | 0 |
| TCTATTGTCAGCCCGA-1 | 1435714 | 8394 | 154295 | 127456 | 1145569 | 370 | 0.033464417 | 3.518868467 | 3.081762257 | 0.095578972 | 0.097135823 | 0 |
| TCTATTGTCATCGACA-1 | 1691866 | 32828 | 180721 | 151629 | 1326688 | 429 | 0.038648951 | 4.040234567 | 2.977955172 | 0.097095003 | 0.093814055 | 0 |
| TCTCATAAGCAGCCCT-1 | 1914610 | 45999 | 201821 | 172317 | 1494473 | 483 | 0.04365011 | 4.560044267 | 3.102080389 | 0.090590517 | 0.090590517 | 0 |
| TCTCATAAGGGATCTG-1 | 1826846 | 32333 | 193085 | 155081 | 1446347 | 467 | 0.042219191 | 4.401677933 | 3.057160925 | 0.091793597 | 0.091866872 | 0 |
| TCTCATAGTGCACACC-1 | 1922910 | 44922 | 202785 | 170730 | 1504473 | 486 | 0.043782983 | 4.5588568 | 3.094669764 | 0.090290402 | 0.08994826 | 0 |
| TCTCATATCTAAGACC-1 | 1919434 | 36747 | 201858 | 179158 | 1501671 | 485 | 0.04354388 | 4.488891867 | 2.964530795 | 0.095792709 | 0.090121616 | 0 |
| TCTCATATCTGTTGTT-1 | 1915492 | 21120 | 210213 | 158209 | 1525950 | 493 | 0.044457448 | 4.609705933 | 2.967072547 | 0.091889034 | 0.091232638 | 0 |
| TCTCTAAAGTAAGTAC-1 | 3199414 | 26317 | 349989 | 267759 | 2555349 | 825 | 0.07459488 | 7.625192267 | 3.142392339 | 0.065123412 | 0.064612121 | 0 |
| TCTCTAAGTAGGTTTC-1 | 2811158 | 35105 | 299375 | 259192 | 2217486 | 716 | 0.064631517 | 6.631572033 | 3.058591613 | 0.074405561 | 0.073218337 | 0 |
| TCTGAGAAGATCTGCT-1 | 1614660 | 46963 | 170188 | 147851 | 1249658 | 404 | 0.036439563 | 3.806809933 | 3.045866803 | 0.101752925 | 0.097130137 | 0 |
| TCTGGAACACGTGGCT-1 | 2179820 | 90735 | 225571 | 191672 | 1671842 | 540 | 0.048656105 | 5.0534868 | 3.045488129 | 0.088527348 | 0.086383276 | 0 |
| TCTGGAATCAGCGGCT-1 | 2129686 | 79186 | 222249 | 189823 | 1638428 | 529 | 0.047715771 | 4.969048533 | 3.110234878 | 0.088620685 | 0.087910695 | 0 |
| TCTTCGGAGAGGGCTT-1 | 2299512 | 70552 | 243777 | 203395 | 1781788 | 576 | 0.051949805 | 5.374983233 | 3.019576077 | 0.087564576 | 0.084500925 | 0 |
| TCTTCGGAGCCAGGAT-1 | 2313476 | 33011 | 252534 | 193239 | 1834692 | 593 | 0.053450381 | 5.507792533 | 3.000838247 | 0.083395649 | 0.081630146 | 0 |
| TCTTCGGAGCTATGCT-1 | 2461126 | 54716 | 265431 | 219865 | 1921114 | 621 | 0.055784619 | 5.716179633 | 3.167027907 | 0.079664064 | 0.079664064 | 0 |
| TCTTCGGGTCGGATGA-1 | 2103426 | 14088 | 229373 | 174624 | 1685341 | 544 | 0.04918171 | 5.1139456 | 3.048181902 | 0.084325521 | 0.084325521 | 0 |
| TCTTTCCCACAGCTAT-1 | 1908746 | 21087 | 211015 | 162260 | 1514384 | 489 | 0.044236023 | 4.601227967 | 3.108020198 | 0.088522896 | 0.08551334 | 0 |
| TCTTTCCCACTAGGCC-1 | 1827330 | 75098 | 192428 | 160970 | 1398834 | 452 | 0.04064839 | 4.237020033 | 3.012111439 | 0.088941913 | 0.089117618 | 0 |
| TCTTTCCGTGGTGAGT-1 | 2129564 | 28226 | 231447 | 187834 | 1682057 | 543 | 0.049141497 | 5.082704567 | 2.974179204 | 0.086624963 | 0.085923215 | 0 |

| | | | | | | | | | | | | |
|---|---|---|---|---|---|---|---|---|---|---|---|---|
| TGAAAGAAGCATGAGC-1 | 2855072 | 53304 | 309220 | 240430 | 2252118 | 728 | 0.06547785 | 6.639474967 | 1.844582972 | 0.123332143 | 0.123332143 | 1 |
| TGAAAGAAGGTACTCT-1 | 2405202 | 78936 | 253646 | 214812 | 1857808 | 600 | 0.053822853 | 5.519801067 | 3.147028739 | 0.083862208 | 0.083807081 | 0 |
| TGAAAGACACCGTCGA-1 | 1794500 | 29796 | 189658 | 155050 | 1419996 | 459 | 0.041365309 | 4.3137469 | 3.530497381 | 0.129582722 | 0.128212542 | 1 |
| TGACAACAGCGAACTG-1 | 1445430 | 37622 | 155968 | 124444 | 1127396 | 364 | 0.0328747 | 3.4672824 | 3.037330341 | 0.097228975 | 0.09649809 | 0 |
| TGACAACAGCTCCTTC-1 | 2416094 | 54856 | 260468 | 226412 | 1874358 | 605 | 0.054573947 | 5.614417167 | 3.071969024 | 0.083393345 | 0.079977711 | 0 |
| TGACGGCAGTTACGGG-1 | 1633552 | 54441 | 175684 | 139920 | 1263507 | 408 | 0.036806702 | 3.857627367 | 3.060383334 | 0.094447921 | 0.094447921 | 0 |
| TGACGGCCAAGCACGA-1 | 1558486 | 14215 | 170157 | 141599 | 1232515 | 398 | 0.035936443 | 3.7842658 | 3.109346365 | 0.09632907 | 0.097112701 | 1 |
| TGACGGCGTCCGATGC-1 | 1356338 | 51445 | 137956 | 116926 | 1050011 | 339 | 0.030513016 | 3.218863867 | 3.033563385 | 0.100819747 | 0.101379759 | 0 |
| TGACGGCGTGTCAACT-1 | 2170004 | 62230 | 225873 | 200723 | 1681178 | 543 | 0.048974778 | 5.073674667 | 3.038201896 | 0.085646725 | 0.085174194 | 0 |
| TGACTAGGTAGGGTGT-1 | 2310132 | 32885 | 246646 | 197514 | 1833087 | 592 | 0.05351069 | 5.5239914 | 3.073518022 | 0.0800951 | 0.081072366 | 0 |
| TGACTAGTCCTCGCTA-1 | 1821144 | 25080 | 198781 | 158841 | 1438442 | 465 | 0.041785297 | 4.328979867 | 2.986659658 | 0.0928327 | 0.093590972 | 0 |
| TGACTAGTCGGGCTTG-1 | 1650928 | 55814 | 175416 | 143120 | 1276578 | 412 | 0.037234729 | 3.906748567 | 3.037652239 | 0.095529789 | 0.095529789 | 0 |
| TGACTAGTCTCATGTT-1 | 1780014 | 20443 | 198948 | 130902 | 1429721 | 462 | 0.041704168 | 4.330304567 | 3.442600699 | 0.124649058 | 0.124840574 | 1 |
| TGACTTTAGATTCACC-1 | 2301090 | 33712 | 250247 | 210277 | 1806854 | 584 | 0.05266166 | 5.4429393 | 3.035469758 | 0.078268887 | 0.079929734 | 0 |
| TGACTTTAGCAACGGT-1 | 2163340 | 50546 | 234468 | 194196 | 1684130 | 544 | 0.049124199 | 5.0935564 | 3.114092434 | 0.081999572 | 0.081999572 | 0 |
| TGACTTTAGGTGTGCA-1 | 1787378 | 15000 | 195578 | 163894 | 1412906 | 456 | 0.041235461 | 4.2998279 | 2.948745996 | 0.088117964 | 0.088117964 | 0 |
| TGACTTTAGTACAGTA-1 | 1181854 | 8987 | 131323 | 96081 | 945463 | 305 | 0.027611339 | 2.910682933 | 3.073652661 | 0.108493702 | 0.107849118 | 0 |
| TGAGAGGAGTGTCCAT-1 | 2246190 | 30857 | 239147 | 200185 | 1776001 | 574 | 0.051879822 | 5.3431709 | 3.071426034 | 0.082182112 | 0.079518386 | 0 |
| TGAGAGGGTCCCACAG-1 | 1274998 | 30185 | 135619 | 105503 | 1003691 | 324 | 0.029205907 | 3.081825967 | 3.042186901 | 0.102022481 | 0.099510792 | 0 |
| TGAGAGGGTCGCGTAC-1 | 2287218 | 47869 | 232259 | 206250 | 1800840 | 582 | 0.052507691 | 5.420204133 | 3.049986857 | 0.080797919 | 0.080797919 | 0 |
| TGAGAGGTCTACGGCG-1 | 1028696 | 109705 | 97781 | 84543 | 736667 | 238 | 0.021444762 | 2.286978967 | 3.098434004 | 0.121248648 | 0.11877367 | 0 |
| TGAGCATCATAGGTCT-1 | 513730 | 4927 | 54398 | 42745 | 411660 | 133 | 0.012021247 | 1.2935369 | 3.096401539 | 0.165979854 | 0.165362042 | 0 |
| TGAGCATCATCGGGCT-1 | 1674960 | 73389 | 173226 | 147939 | 1280406 | 414 | 0.037205671 | 3.8943667 | 3.106671874 | 0.09599866 | 0.096358153 | 0 |
| TGAGCATTCTGTATGG-1 | 1263814 | 18346 | 137911 | 101915 | 1005642 | 325 | 0.029381168 | 3.091855867 | 2.927811399 | 0.102974575 | 0.105867354 | 0 |
| TGAGCCGAGGCATGGT-1 | 2230514 | 18235 | 245149 | 182230 | 1784900 | 577 | 0.052152634 | 5.3759433 | 3.127876002 | 0.081809335 | 0.081618828 | 0 |
| TGAGCCGGTGCTGTAT-1 | 1483950 | 18989 | 162847 | 124426 | 1177688 | 380 | 0.034299336 | 3.604278533 | 3.058877011 | 0.098434442 | 0.097729371 | 0 |
| TGAGCCGGTTCCCTAC-1 | 2850160 | 32697 | 311529 | 246758 | 2259176 | 730 | 0.06574085 | 6.701109767 | 3.142385235 | 0.080585868 | 0.076401185 | 0 |
| TGAGGGACACCAATGT-1 | 1587742 | 19326 | 173797 | 131538 | 1263081 | 408 | 0.036805808 | 3.849979733 | 3.148286002 | 0.092765422 | 0.093065131 | 0 |
| TGATTTCTCACGATAC-1 | 2264160 | 37396 | 247189 | 190991 | 1788584 | 578 | 0.052051663 | 5.3831451 | 3.044897524 | 0.088143788 | 0.083797329 | 0 |
| TGATTTCTCCGACCAG-1 | 2242330 | 63221 | 240928 | 194338 | 1743843 | 563 | 0.050864673 | 5.269453367 | 3.077897534 | 0.080692727 | 0.081399508 | 0 |
| TGCACCTAGTACGCGA-1 | 1570974 | 11805 | 168678 | 152865 | 1237626 | 400 | 0.036107073 | 3.799542033 | 3.071863434 | 0.09129586 | 0.093694244 | 0 |
| TGCACCTTCAGCCCGA-1 | 2126416 | 17097 | 230974 | 196023 | 1682322 | 543 | 0.049042675 | 5.087482 | 3.101381408 | 0.089714739 | 0.089540122 | 0 |
| TGCCAAAAGACAGGCT-1 | 2205220 | 33594 | 240866 | 195316 | 1735444 | 561 | 0.050362034 | 5.1986704 | 3.104849996 | 0.084475637 | 0.085388156 | 0 |
| TGCCAAACAAGTGTTC-1 | 2318874 | 48619 | 250360 | 209154 | 1810741 | 585 | 0.05258351 | 5.408093033 | 3.048254496 | 0.082998839 | 0.08339146 | 0 |
| TGCCAAACACTACATG-1 | 1159192 | 15428 | 128451 | 92876 | 922437 | 298 | 0.026889479 | 2.8334616 | 3.09073673 | 0.113927365 | 0.113192954 | 0 |
| TGCCCATAGGATGCGT-1 | 2322016 | 26031 | 249902 | 210296 | 1835787 | 593 | 0.053553383 | 5.5353585 | 2.989264654 | 0.082938338 | 0.082355028 | 0 |
| TGCCCATAGGCCATAG-1 | 1179806 | 11927 | 132453 | 95052 | 940374 | 304 | 0.027405508 | 2.877125467 | 3.090991087 | 0.110113703 | 0.107881722 | 0 |
| TGCCCATGTACCAGCC-1 | 1480952 | 34141 | 157916 | 138183 | 1150712 | 372 | 0.033434594 | 3.527727133 | 3.175128761 | 0.09815268 | 0.099764682 | 0 |
| TGCCCATTCCAACAAC-1 | 2280048 | 42613 | 249404 | 199731 | 1788300 | 578 | 0.052184137 | 5.3928318 | 3.139472263 | 0.082650292 | 0.082580424 | 0 |
| TGCCCTAAGCCAAACG-1 | 1356656 | 74585 | 140587 | 114172 | 1027312 | 332 | 0.029860053 | 3.147235967 | 3.114372772 | 0.098688163 | 0.098688163 | 0 |
| TGCCCTACAAATTCCG-1 | 777146 | 95937 | 74855 | 63385 | 542969 | 175 | 0.015791569 | 1.6980025 | 3.173872294 | 0.161906754 | 0.158341636 | 1 |
| TGCCCTAGTCGTAATC-1 | 1975040 | 51457 | 209701 | 171162 | 1542720 | 498 | 0.044757802 | 4.642507733 | 3.070620666 | 0.092433183 | 0.088201743 | 0 |
| TGCCCTAGTGTAGTTC-1 | 1911972 | 53415 | 197436 | 186818 | 1474303 | 476 | 0.042523953 | 4.372696133 | 3.12513853 | 0.098267159 | 0.093047783 | 0 |
| TGCCCTAGTTCTAGAC-1 | 1332094 | 35052 | 145172 | 116857 | 1035013 | 334 | 0.030070314 | 3.175142967 | 2.938426025 | 0.110306522 | 0.104334152 | 0 |
| TGCGCAGAGAGGTACC-1 | 2591744 | 33170 | 276489 | 235836 | 2046249 | 661 | 0.059659311 | 6.130635167 | 2.971792668 | 0.082219126 | 0.080051685 | 0 |
| TGCGCAGAGTACTTGC-1 | 2644076 | 52139 | 288877 | 217996 | 2085064 | 674 | 0.060636132 | 6.220085633 | 4.213582425 | 0.137010351 | 0.137010351 | 1 |
| TGCGGGTCAGACAGTG-1 | 2046096 | 22868 | 217821 | 161263 | 1644144 | 531 | 0.047909237 | 4.982054333 | 3.073867551 | 0.082229454 | 0.083707994 | 0 |
| TGCGGGTCATACGTCA-1 | 1722598 | 26392 | 181826 | 156019 | 1358361 | 439 | 0.039451685 | 4.109787033 | 3.096341385 | 0.093814912 | 0.092925259 | 0 |
| TGCGGGTCATCTGTAG-1 | 1201460 | 16588 | 127037 | 98139 | 959696 | 310 | 0.027976549 | 2.950293333 | 2.883502936 | 0.110425252 | 0.105812401 | 0 |
| TGCGGGTTCTATGGTG-1 | 1058004 | 18855 | 112879 | 86075 | 840195 | 271 | 0.024440142 | 2.583938733 | 3.142219035 | 0.116078293 | 0.113178676 | 0 |
| TGCGTGGAGTAGTGCG-1 | 1740654 | 61939 | 177294 | 157398 | 1344023 | 434 | 0.039124318 | 4.112230867 | 3.003047221 | 0.091864 | 0.092656977 | 0 |
| TGCTACCGTACGTCAT-1 | 2338412 | 32231 | 251948 | 207502 | 1846731 | 597 | 0.053848683 | 5.557345533 | 3.019380368 | 0.08522223 | 0.081802217 | 0 |
| TGCTACCTCCTGAAAC-1 | 2328798 | 57185 | 251701 | 197415 | 1822497 | 589 | 0.053076687 | 5.472549767 | 3.78013005 | 0.133720725 | 0.133720725 | 1 |
| TGCTGCTTCGGAGAGT-1 | 1893980 | 30603 | 204229 | 168862 | 1490286 | 481 | 0.043508521 | 4.5235827 | 3.110624907 | 0.090581256 | 0.087734681 | 0 |
| TGGACGCAGACGCTTT-1 | 2202638 | 52602 | 236071 | 198834 | 1715131 | 554 | 0.049821263 | 5.1413468 | 3.108028046 | 0.089715809 | 0.087198547 | 0 |
| TGGACGCCATCGGCGT-1 | 1888278 | 24496 | 208867 | 176236 | 1478679 | 478 | 0.043051265 | 4.489986367 | 3.10999936 | 0.088229532 | 0.086696381 | 0 |
| TGGACGCGTTACTGAC-1 | 2579716 | 48930 | 276725 | 215949 | 2038112 | 658 | 0.059413806 | 6.102296433 | 3.923585536 | 0.132668797 | 0.132668797 | 1 |
| TGGACGCTCGAACCCG-1 | 1099690 | 94206 | 108603 | 92807 | 804074 | 260 | 0.023385061 | 2.491230233 | 3.097826319 | 0.118451133 | 0.114550716 | 0 |
| TGGCCAGGTCCTAGTA-1 | 1081622 | 10923 | 119159 | 92642 | 858898 | 277 | 0.025005342 | 2.6334069 | 3.313451173 | 0.100811049 | 0.103749212 | 0 |
| TGGCCAGTCAAGCCAT-1 | 2429658 | 18020 | 268174 | 215629 | 1927835 | 623 | 0.056254529 | 5.797977733 | 3.037953455 | 0.078442906 | 0.077700931 | 0 |
| TGGCGCAGTCGACCAC-1 | 2510174 | 34666 | 269607 | 225037 | 1980864 | 640 | 0.0577571 | 5.954932467 | 3.012374972 | 0.077358669 | 0.079038133 | 0 |
| TGGCGCAGTGTCCGAC-1 | 2057088 | 51484 | 217459 | 192990 | 1595155 | 515 | 0.046428273 | 4.8419747 | 3.074563148 | 0.085518767 | 0.086336798 | 0 |
| TGGCGCATCCAACCGG-1 | 1653480 | 79616 | 170058 | 143265 | 1260541 | 407 | 0.036719662 | 3.854260367 | 3.037506483 | 0.094652758 | 0.095303736 | 0 |
| TGGCGCATCCGGTTCT-1 | 2697504 | 82272 | 283406 | 233043 | 2098783 | 678 | 0.061026998 | 6.2583796 | 3.107118495 | 0.079939894 | 0.080148024 | 0 |
| TGGCTGGAGTAAGTAC-1 | 2488488 | 69708 | 265930 | 216674 | 1936176 | 625 | 0.056346496 | 5.814364333 | 3.074736206 | 0.082311399 | 0.078424091 | 0 |
| TGGCTGGCAACGCGCT-1 | 1279616 | 48076 | 131238 | 113366 | 986936 | 319 | 0.028682965 | 3.025945233 | 2.999573806 | 0.104871499 | 0.103700837 | 0 |
| TGGCTGGGTCGAGTAG-1 | 1106740 | 17546 | 111502 | 88090 | 889602 | 287 | 0.026013373 | 2.7488078 | 3.058615082 | 0.120210438 | 0.120811857 | 1 |
| TGGGAAGAGGAATGGA-1 | 1715148 | 19481 | 180949 | 150992 | 1363726 | 441 | 0.039728546 | 4.1606326 | 3.072323984 | 0.092723306 | 0.091157984 | 0 |
| TGGGAAGCAAGGTGAC-1 | 2099758 | 49322 | 218925 | 192744 | 1638767 | 529 | 0.047776744 | 4.958985833 | 2.918677416 | 0.085996629 | 0.085231535 | 0 |
| TGGGAAGGTGGTAACG-1 | 1753774 | 60578 | 180999 | 151271 | 1360926 | 440 | 0.0396101 | 4.1445208 | 3.016784578 | 0.088769045 | 0.089751218 | 0 |
| TGGGAAGTCACTTTAC-1 | 2087938 | 60473 | 221803 | 173378 | 1632284 | 527 | 0.047544771 | 4.9363335 | 3.052657005 | 0.086052078 | 0.084871897 | 0 |
| TGGGCGTAGGCTCAGA-1 | 1803992 | 18247 | 197036 | 161316 | 1427393 | 461 | 0.041678408 | 4.360065833 | 3.051034179 | 0.088222086 | 0.088273144 | 0 |
| TGGGCGTGTTTCCTGC-1 | 2065052 | 30114 | 220357 | 196594 | 1617987 | 523 | 0.047237485 | 4.904689333 | 3.067431535 | 0.086477404 | 0.084336553 | 0 |
| TGGGCGTTCACTGTTT-1 | 1628738 | 45453 | 173929 | 136418 | 1272938 | 411 | 0.037030956 | 3.8643664 | 3.064156197 | 0.093996453 | 0.095283801 | 0 |
| TGGTTAGAGGGATGGG-1 | 2136016 | 27446 | 227248 | 191032 | 1690290 | 546 | 0.049415262 | 5.109077333 | 2.762450992 | 0.077885845 | 0.080686337 | 0 |
| TGGTTAGAGTTAGCGG-1 | 1509668 | 56410 | 157554 | 133924 | 1161780 | 375 | 0.033887053 | 3.571696733 | 3.020323962 | 0.101588912 | 0.098339965 | 0 |
| TGGTTAGTCTTATGGG-1 | 1720824 | 21747 | 186238 | 143678 | 1369161 | 442 | 0.040019666 | 4.182032267 | 3.048977127 | 0.093677668 | 0.091973687 | 0 |
| TGGTTCCAGACAAAGG-1 | 1279028 | 21754 | 138799 | 110897 | 1007578 | 325 | 0.029427004 | 3.1106308 | 3.164177196 | 0.105412974 | 0.102548187 | 0 |
| TGGTTCCAGAGTAAGG-1 | 958374 | 17717 | 103707 | 80834 | 756116 | 244 | 0.022100321 | 2.35485 | 3.062393454 | 0.120235971 | 0.119427301 | 0 |
| TGGTTCCAGGTGCACA-1 | 2036712 | 23394 | 219339 | 187701 | 1606278 | 519 | 0.046816131 | 4.867484833 | 3.022094816 | 0.085064895 | 0.085325529 | 0 |
| TGGTTCCTCACACCTC-1 | 1978410 | 30220 | 210729 | 189525 | 1547936 | 500 | 0.045095221 | 4.6917585 | 3.023925552 | 0.087341338 | 0.08666308 | 0 |
| TGTATTCCAACATAAG-1 | 1119632 | 13749 | 121920 | 86867 | 897096 | 290 | 0.026196871 | 2.756805967 | 3.043910787 | 0.111815906 | 0.108764804 | 0 |
| TGTATTCCAATCGCGC-1 | 1437018 | 40625 | 149085 | 130389 | 1116919 | 361 | 0.032512509 | 3.4302753 | 3.036806636 | 0.10028618 | 0.101031764 | 0 |
| TGTCCCACAGTCGGAA-1 | 1277180 | 12412 | 137860 | 117169 | 1009739 | 326 | 0.029450601 | 3.1107462 | 3.054171904 | 0.106362034 | 0.103478361 | 0 |
| TGTCCCAGTAGATAGT-1 | 1775788 | 13563 | 195489 | 145980 | 1420756 | 459 | 0.041532861 | 4.319871567 | 3.038462631 | 0.08922246 | 0.089671268 | 0 |
| TGTGGTACATTATCGG-1 | 2011992 | 46673 | 213888 | 185191 | 1566240 | 506 | 0.045669386 | 4.764361367 | 2.873889043 | 0.090011291 | 0.090011291 | 0 |
| TGTGGTACATTGAAGA-1 | 1353890 | 15942 | 147028 | 116939 | 1073981 | 347 | 0.03132305 | 3.2990804 | 3.040106534 | 0.098954405 | 0.099557534 | 0 |
| TGTGGTAGTTTGGGCC-1 | 1778828 | 89157 | 182485 | 158408 | 1348778 | 436 | 0.039210661 | 4.090710833 | 3.531602236 | 0.092120743 | 0.090091769 | 0 |
| TGTGGTATCAAGCCGC-1 | 2098862 | 24944 | 221309 | 191189 | 1661420 | 537 | 0.048396289 | 5.032009967 | 3.141153237 | 0.086389544 | 0.085456139 | 0 |
| TGTGTTTGTAAACCTC-1 | 1961044 | 35769 | 206872 | 183266 | 1535137 | 496 | 0.044711685 | 4.668730267 | 2.91513636 | 0.088265971 | 0.087412959 | 1 |
| TGTTCCGCATAGAGCG-1 | 1049952 | 32599 | 108495 | 98293 | 810565 | 262 | 0.023596366 | 2.5237046 | 3.03280784 | 0.118320551 | 0.115384736 | 0 |
| TTAACTCCAGCTCATA-1 | 1529170 | 13449 | 168292 | 129522 | 1217907 | 393 | 0.035558147 | 3.7234074 | 3.06188379 | 0.099634408 | 0.095964961 | 0 |
| TTAACTCCATGGCTAT-1 | 2028180 | 22758 | 224155 | 177117 | 1604150 | 518 | 0.046836777 | 4.8517156 | 2.882402245 | 0.08920444 | 0.086001427 | 0 |
| TTAACTCTCAGTACAC-1 | 2105048 | 34679 | 229456 | 185738 | 1655175 | 535 | 0.048248106 | 5.0062402 | 3.14321135 | 0.086020146 | 0.081571967 | 0 |

| | | | | | | | | | | | | |
|---|---|---|---|---|---|---|---|---|---|---|---|---|
| TTAACTCTCGATAGTC-1 | 1474440 | 17808 | 160921 | 120129 | 1175582 | 380 | 0.034241005 | 3.577867233 | 3.147446914 | 0.100643886 | 0.095227836 | 0 |
| TTAGGACAGAGCTGCA-1 | 1595386 | 32558 | 164107 | 147049 | 1251672 | 404 | 0.036456041 | 3.8217652 | 3.094770337 | 0.093285795 | 0.093285795 | 0 |
| TTAGGACAGTTACCCA-1 | 1550908 | 27499 | 166178 | 136792 | 1220439 | 394 | 0.03558407 | 3.7387442 | 2.949396581 | 0.093831592 | 0.093504739 | 0 |
| TTAGGACTCTGAAGTC-1 | 1979764 | 52024 | 210974 | 170108 | 1546658 | 500 | 0.044987534 | 4.677815067 | 3.243281833 | 0.087948562 | 0.087308485 | 0 |
| TTAGGCAAGGTAAGTT-1 | 1838286 | 41598 | 193838 | 158775 | 1444075 | 466 | 0.041963435 | 4.3667543 | 3.143374572 | 0.094747524 | 0.091035258 | 0 |
| TTAGGCACACCAATGT-1 | 1659096 | 18028 | 182100 | 132665 | 1326303 | 428 | 0.038691834 | 4.0332062 | 2.989678345 | 0.095402005 | 0.092476229 | 0 |
| TTAGGCAGTAACGACG-1 | 1420648 | 83201 | 143746 | 126740 | 1066961 | 345 | 0.031034127 | 3.280928533 | 2.704059093 | 0.096312458 | 0.099365881 | 0 |
| TTAGTTCAGAGCCACA-1 | 2118102 | 24440 | 227912 | 192425 | 1673325 | 541 | 0.048824728 | 5.066397933 | 3.039220143 | 0.086366542 | 0.084610171 | 0 |
| TTATGCTAGTGCCATT-1 | 2088700 | 24895 | 224114 | 181109 | 1658582 | 536 | 0.048367567 | 5.005697033 | 3.068885146 | 0.086917604 | 0.083263099 | 0 |
| TTATGCTCAGAGACGT-1 | 1789210 | 22113 | 188081 | 172028 | 1406988 | 455 | 0.041107791 | 4.2985327 | 3.123509854 | 0.090509279 | 0.090509279 | 0 |
| TTCCCAGAGGAGCGTT-1 | 3410494 | 50339 | 371398 | 297977 | 2690780 | 869 | 0.078588861 | 7.989324733 | 3.025966737 | 0.064819593 | 0.063545584 | 0 |
| TTCCCAGCACGATCTA-1 | 2170762 | 31556 | 238421 | 187736 | 1713049 | 553 | 0.049962953 | 5.174475 | 3.080588214 | 0.082286245 | 0.083176806 | 0 |
| TTCCCAGCAGATTGTC-1 | 2269404 | 32642 | 252997 | 181872 | 1801893 | 582 | 0.052483089 | 5.405535033 | 3.61962767 | 0.128563063 | 0.128563063 | 1 |
| TTCCCAGGTTCTCTCG-1 | 1518232 | 59726 | 160765 | 133601 | 1164140 | 376 | 0.033947829 | 3.568523767 | 3.110572039 | 0.096483092 | 0.096893851 | 0 |
| TTCCCAGTCCGCCTTA-1 | 1558652 | 26935 | 167826 | 136977 | 1226914 | 396 | 0.035807866 | 3.757928467 | 3.019722212 | 0.100214809 | 0.095940773 | 0 |
| TTCGAAGGTACTCGCG-1 | 1097584 | 78473 | 109463 | 93208 | 816440 | 264 | 0.023725873 | 2.529146867 | 3.142569026 | 0.115683945 | 0.113221435 | 0 |
| TTCGGTCAGCGATGAC-1 | 2026556 | 32170 | 219239 | 180082 | 1595065 | 515 | 0.046512354 | 4.828305467 | 2.922493945 | 0.084211565 | 0.084915579 | 0 |
| TTCGGTCAGTGAATTG-1 | 1058934 | 20657 | 114151 | 87909 | 836217 | 270 | 0.024396762 | 2.5830278 | 3.032521942 | 0.112638425 | 0.112835725 | 0 |
| TTCGGTCCACTTGAGT-1 | 1611112 | 16542 | 178030 | 132313 | 1284227 | 415 | 0.037518591 | 3.914983 | 3.012017899 | 0.09532526 | 0.094131156 | 0 |
| TTCGGTCCATAGCACT-1 | 2238462 | 72314 | 236292 | 199512 | 1730344 | 559 | 0.050349712 | 5.222405633 | 3.144026198 | 0.084785477 | 0.083641909 | 0 |
| TTCTACAAGAGGTAGA-1 | 1694298 | 8195 | 183875 | 153921 | 1348307 | 436 | 0.039395826 | 4.124490433 | 2.99416189 | 0.090071562 | 0.088822567 | 0 |
| TTCTACAAGCAGAGAA-1 | 1574894 | 10153 | 173968 | 135173 | 1255600 | 406 | 0.03661971 | 3.831094067 | 3.135648122 | 0.099439762 | 0.096059036 | 0 |
| TTCTACAAGTGACTCT-1 | 2012298 | 55001 | 214907 | 181782 | 1560608 | 504 | 0.045334068 | 4.6965471 | 2.984026194 | 0.092837694 | 0.090940493 | 0 |
| TTCTACAGTCGCTTCT-1 | 888010 | 46345 | 60770 | 84420 | 696475 | 225 | 0.020109962 | 2.154067733 | 3.029495509 | 0.123385744 | 0.125334078 | 0 |
| TTCTACAGTCTAACGT-1 | 1707524 | 26049 | 182750 | 157019 | 1341706 | 433 | 0.039116385 | 4.080855133 | 3.027665681 | 0.096551442 | 0.093348527 | 0 |
| TTCTACATCTTAGCTT-1 | 1910458 | 67259 | 201417 | 166940 | 1474842 | 476 | 0.042920241 | 4.477946733 | 3.107831993 | 0.093060523 | 0.09022728 | 0 |
| TTCTCAAAGGCACATG-1 | 1193874 | 13237 | 131242 | 97648 | 951747 | 307 | 0.027800827 | 2.9285391 | 3.006988289 | 0.105918162 | 0.10452414 | 0 |
| TTCTCAACATACGGAG-1 | 1597084 | 22752 | 174803 | 132223 | 1267306 | 409 | 0.036932853 | 3.868071333 | 3.001512935 | 0.100648237 | 0.099907754 | 1 |
| TTCTCAAGTAGGGTGT-1 | 2393238 | 30384 | 256764 | 204592 | 1901498 | 614 | 0.055474223 | 5.708763667 | 3.135010515 | 0.08464697 | 0.083351221 | 0 |
| TTCTCAAGTGGCCACT-1 | 1674236 | 43591 | 179674 | 142926 | 1308045 | 423 | 0.038256929 | 4.012296967 | 3.000099454 | 0.090301191 | 0.089594098 | 0 |
| TTCTCCTAGACAGACC-1 | 2387694 | 39929 | 253713 | 214503 | 1879549 | 607 | 0.054630064 | 5.605624233 | 3.007757335 | 0.084625881 | 0.082675197 | 0 |
| TTCTCCTAGCAGCCCT-1 | 1833014 | 42327 | 195156 | 165598 | 1429933 | 462 | 0.041720331 | 4.361396533 | 3.113643628 | 0.090803904 | 0.088958474 | 0 |
| TTCTCCTCATTCGTTT-1 | 2042690 | 52324 | 219670 | 173413 | 1597283 | 516 | 0.046449137 | 4.8151224 | 3.036506358 | 0.088653452 | 0.087666527 | 0 |
| TTCTCCTTCATCGCGG-1 | 1724246 | 41249 | 182606 | 152687 | 1347704 | 435 | 0.039307074 | 4.1129284 | 2.936999595 | 0.093491467 | 0.092637133 | 0 |
| TTCTTAGAGTACTTGC-1 | 1816392 | 45713 | 192679 | 159270 | 1418730 | 458 | 0.041290324 | 4.312421867 | 3.046341481 | 0.094283308 | 0.091475731 | 0 |
| TTGAACGAGAATTGTG-1 | 1261492 | 19055 | 139620 | 107277 | 995540 | 322 | 0.029050556 | 3.062350367 | 2.991368961 | 0.103737354 | 0.104663631 | 0 |
| TTGAACGAGCACGCCT-1 | 1932494 | 58414 | 203161 | 178296 | 1492623 | 482 | 0.043510522 | 4.547870567 | 2.984749362 | 0.089137176 | 0.087115229 | 0 |
| TTGAACGAGTGTCCAT-1 | 2427774 | 19803 | 263911 | 221176 | 1922884 | 621 | 0.05613529 | 5.782304967 | 3.110133191 | 0.082190797 | 0.083340915 | 0 |
| TTGACTTAGGGCTTCC-1 | 2166276 | 55418 | 230056 | 199682 | 1681120 | 543 | 0.04895468 | 5.067576667 | 3.099070108 | 0.087104504 | 0.084332644 | 0 |
| TTGACTTAGTCCGTAT-1 | 2157434 | 17831 | 241457 | 177881 | 1720265 | 556 | 0.050215204 | 5.185823533 | 3.115134851 | 0.091242507 | 0.083895761 | 0 |
| TTGCCGTAGAGTAATC-1 | 2024858 | 43498 | 216751 | 186412 | 1578197 | 510 | 0.04585224 | 4.744569633 | 3.022962789 | 0.087024323 | 0.087380481 | 0 |
| TTGCCGTAGTCGTGTT-1 | 1775716 | 24061 | 188529 | 151571 | 1411555 | 456 | 0.04116602 | 4.278098033 | 3.108176253 | 0.089027017 | 0.089741527 | 0 |
| TTGCCGTGTAAGCGCA-1 | 1195024 | 21708 | 124798 | 109967 | 938551 | 303 | 0.027275758 | 2.887878967 | 3.046110677 | 0.105504833 | 0.10744099 | 0 |
| TTGCCGTGTGCACTTA-1 | 1392326 | 20664 | 151122 | 119506 | 1101034 | 356 | 0.032064819 | 3.377338067 | 3.011639302 | 0.105595861 | 0.100182464 | 0 |
| TTGCGTCAGGATGTAT-1 | 1799076 | 18768 | 191061 | 152468 | 1436779 | 464 | 0.041846457 | 4.347007633 | 3.120615188 | 0.090280685 | 0.088327953 | 0 |
| TTGCGTCTCATTTGCT-1 | 1728188 | 56945 | 174456 | 153017 | 1343770 | 434 | 0.039125135 | 4.0976749 | 2.886227587 | 0.092236296 | 0.090847506 | 0 |
| TTGGAACGTAAGGTGC-1 | 2068648 | 76586 | 213524 | 181729 | 1596809 | 516 | 0.046451737 | 4.8303182 | 3.097526321 | 0.087404416 | 0.085682804 | 0 |
| TTGGCAATCATGGTGT-1 | 2502470 | 31371 | 274371 | 205370 | 1991358 | 643 | 0.058012858 | 5.947461133 | 3.014485238 | 0.081714822 | 0.080489835 | 0 |
| TTGTAGGAGAGTACAT-1 | 1853772 | 16296 | 198373 | 158421 | 1480682 | 478 | 0.043220294 | 4.495245933 | 3.121473683 | 0.090192633 | 0.086523665 | 0 |
| TTGTAGGTCTTCTTGA-1 | 846372 | 8967 | 90600 | 73754 | 673051 | 217 | 0.019675513 | 2.095048733 | 2.967917628 | 0.123608574 | 0.127052684 | 0 |
| TTTACTGAGAAACGCC-1 | 1748232 | 57762 | 182000 | 157647 | 1350823 | 436 | 0.039343693 | 4.117528833 | 3.042934852 | 0.092619265 | 0.091344543 | 0 |
| TTTACTGAGACAGACC-1 | 2507462 | 41263 | 269344 | 227167 | 1969688 | 636 | 0.057368802 | 5.902468533 | 3.143568471 | 0.080711903 | 0.081412665 | 0 |
| TTTACTGAGCCCAATT-1 | 791808 | 23315 | 85173 | 59406 | 623914 | 202 | 0.018290932 | 1.942565367 | 3.034131898 | 0.1329755 | 0.12963281 | 0 |
| TTTACTGCACCACATA-1 | 1627156 | 12186 | 181064 | 123935 | 1309971 | 423 | 0.038184502 | 3.972668633 | 4.341722325 | 0.142063155 | 0.142288013 | 1 |
| TTTACTGGTAACAATG-1 | 1265020 | 14039 | 138758 | 102365 | 1009858 | 326 | 0.029491219 | 3.0987657 | 3.090294544 | 0.105374723 | 0.103842487 | 0 |
| TTTACTGTCCACGATA-1 | 1610830 | 14013 | 177652 | 133902 | 1285263 | 415 | 0.037522447 | 3.9263845 | 3.047494584 | 0.097954365 | 0.096434892 | 0 |
| TTTATGCAGTAGTATG-1 | 1110412 | 10943 | 120717 | 89115 | 889637 | 287 | 0.026015486 | 2.744050233 | 3.046512123 | 0.111317607 | 0.110367296 | 0 |
| TTTATGCTCTGAAGGG-1 | 1640902 | 26590 | 172497 | 144502 | 1297313 | 419 | 0.037880672 | 3.9750998 | 2.951717721 | 0.094170306 | 0.095011192 | 0 |
| TTTCCTCAGCTCCTTC-1 | 2516606 | 47918 | 272596 | 232277 | 1963815 | 634 | 0.057211154 | 5.9116326 | 3.107555909 | 0.08033103 | 0.080579297 | 0 |
| TTTCCTCCAACAGCCC-1 | 1528114 | 78079 | 158612 | 136226 | 1155197 | 373 | 0.033614186 | 3.543141167 | 2.939779998 | 0.098787269 | 0.098787269 | 0 |
| TTTCCTCCATTCAGCA-1 | 1630466 | 39279 | 175684 | 142551 | 1272952 | 411 | 0.037002093 | 3.8811907 | 3.081360001 | 0.101998451 | 0.098920324 | 1 |
| TTTCCTCGTCAAGTCT-1 | 2199828 | 81390 | 230726 | 200244 | 1687468 | 545 | 0.048730802 | 4.9934283 | 2.895542324 | 0.093373999 | 0.093373999 | 1 |
| TTTGCGCAGTGCCATT-1 | 2064206 | 28000 | 217583 | 182083 | 1636540 | 529 | 0.047712934 | 4.935847433 | 3.14365775 | 0.087003648 | 0.086134641 | 0 |
| TTTGCGCCAACTCATG-1 | 986010 | 13517 | 106183 | 81906 | 784404 | 253 | 0.022856781 | 2.428664367 | 2.908530435 | 0.118646476 | 0.118646476 | 0 |
| TTTGCGCTCGATAAGA-1 | 1067886 | 8933 | 115441 | 87614 | 855898 | 276 | 0.024961824 | 2.634196033 | 3.123445744 | 0.115750924 | 0.111189326 | 0 |
| TTTGCGCTCTCATCCG-1 | 1469370 | 67617 | 152246 | 129953 | 1119554 | 362 | 0.032602735 | 3.431065867 | 3.005660712 | 0.099645637 | 0.099645637 | 0 |
| TTTGGTTCAAGACGAC-1 | 1678288 | 58039 | 178003 | 149809 | 1292437 | 417 | 0.037571985 | 3.946982167 | 3.077345398 | 0.093458163 | 0.091135887 | 0 |
| TTTGGTTGTGATGTTC-1 | 2501734 | 57366 | 264979 | 225337 | 1954052 | 631 | 0.056596783 | 5.740793067 | 3.098451375 | 0.084508055 | 0.082261345 | 0 |
| TTTGGTTTCAGATTCG-1 | 1517258 | 145646 | 149613 | 127044 | 1094955 | 354 | 0.031717703 | 3.341627867 | 3.133985188 | 0.104922885 | 0.101959238 | 0 |
| TTTGTCAGTGTGTTCA-1 | 2632962 | 41823 | 284940 | 228845 | 2077354 | 671 | 0.060549859 | 6.2191403 | 3.061144741 | 0.079651328 | 0.07953071 | 0 |
| TTTGTCATCACTGCGG-1 | 1759810 | 30953 | 190198 | 159556 | 1379103 | 445 | 0.040243654 | 4.208228967 | 2.885560867 | 0.093087254 | 0.091058968 | 0 |

**Supplementary Table 3 | DAPC classification of single cells.** Assignment of each barcode to DAPC defined cluster and group.

| cell_id | barcode | cluster | group |
|---|---|---|---|
| 0 | AAACCTGAGGCTACGA-1 | 4 | A |
| 1 | AAACCTGGTCCTTGGG-1 | 4 | A |
| 2 | AAACCTGTCACACGGC-1 | 9 | D |
| 3 | AAACCTGTCGATACAC-1 | 11 | D |
| 4 | AAACGGGAGCAGATCG-1 | 2 | A |
| 5 | AAACGGGCAAAGTAAC-1 | 1 | A |
| 6 | AAACGGGCAACCTCAA-1 | 4 | A |
| 7 | AAACGGGGTCTTCGTC-1 | 8 | D |
| 8 | AAACGGGTCACGCTCT-1 | 10 | D |
| 9 | AAACGGGTCAGCGGCT-1 | 11 | D |
| 10 | AAAGATGAGAATTGTG-1 | 3 | A |
| 12 | AAAGCAAGTGACACGA-1 | 1 | A |
| 13 | AAAGCAATCGCGGACT-1 | 5 | A |
| 14 | AAAGCAATCTCACCTG-1 | 8 | D |
| 15 | AAAGTAGTCTCAGCGG-1 | 11 | D |
| 16 | AAAGTAGTCTTGCGAA-1 | 10 | D |
| 17 | AAATGCCAGTGGAGAA-1 | 11 | D |
| 18 | AAATGCCGTCGCGCTA-1 | 8 | D |
| 19 | AACACGTAGGACAGAA-1 | 1 | A |
| 20 | AACCATGAGGAATCGC-1 | 3 | A |
| 22 | AACCATGCATTCATCT-1 | 10 | D |
| 23 | AACCATGTCGACGAAG-1 | 8 | D |
| 24 | AACCGCGGTAACAGCG-1 | 2 | A |
| 25 | AACCGCGGTTTACTTC-1 | 3 | A |
| 26 | AACCGCGTCCTACGAA-1 | 2 | A |
| 27 | AACGTTGGTGGTAACG-1 | 5 | A |
| 28 | AACTCAGAGTGGGTAC-1 | 7 | C |
| 29 | AACTCAGAGTGGTAGC-1 | 4 | A |
| 30 | AACTCAGCATGCAGTT-1 | 10 | D |
| 31 | AACTCAGTCGAGTATC-1 | 5 | A |
| 32 | AACTCAGTCTAACACG-1 | 8 | D |
| 33 | AACTCCCTCAGTTCTT-1 | 10 | D |
| 34 | AACTCCCTCCACGATA-1 | 11 | D |
| 35 | AACTCCCTCGATAGGG-1 | 2 | A |
| 36 | AACTCCCTCTAATGGC-1 | 10 | D |
| 37 | AACTCTTAGAGTAGCC-1 | 5 | A |
| 38 | AACTCTTGTCTTACCC-1 | 3 | A |
| 40 | AACTGGTAGAAACCAT-1 | 5 | A |
| 41 | AACTGGTAGCTGAACG-1 | 11 | D |

| | | | |
|---|---|---|---|
| 42 | AACTGGTAGGGCCATA-1 | 1 | A |
| 43 | AACTGGTGTCGTGAAG-1 | 9 | D |
| 44 | AACTGGTGTTCAGTAC-1 | 5 | A |
| 45 | AACTGGTTCGTACGCG-1 | 11 | D |
| 46 | AACTTTCCACGTAGGA-1 | 8 | D |
| 47 | AACTTTCTCTGCTGAA-1 | 11 | D |
| 48 | AAGACCTCACAAGCGA-1 | 2 | A |
| 49 | AAGACCTCACCAGTGC-1 | 8 | D |
| 51 | AAGACCTCACTTGAGT-1 | 3 | A |
| 52 | AAGACCTCATAGCACT-1 | 8 | D |
| 53 | AAGACCTGTGGACGTA-1 | 11 | D |
| 54 | AAGACCTTCACGGTAT-1 | 5 | A |
| 55 | AAGACCTTCCGAGATT-1 | 6 | B |
| 56 | AAGCCGCCATCTCACC-1 | 5 | A |
| 57 | AAGGAGCCAAGAATCA-1 | 6 | B |
| 58 | AAGGCAGAGAGCCCAA-1 | 3 | A |
| 59 | AAGGCAGGTAAAGTCA-1 | 5 | A |
| 60 | AAGGTTCAGCGTGTGA-1 | 6 | B |
| 61 | AAGGTTCCAGATTGTC-1 | 5 | A |
| 62 | AAGGTTCGTCCGAATT-1 | 2 | A |
| 63 | AAGGTTCGTGTAATGA-1 | 9 | D |
| 64 | AAGGTTCGTTGAGCAG-1 | 8 | D |
| 66 | AAGGTTCTCCGCGACA-1 | 1 | A |
| 67 | AAGTCTGAGGCCCGTT-1 | 8 | D |
| 68 | AAGTCTGAGTGCGATG-1 | 1 | A |
| 69 | AAGTCTGTCACCGCGA-1 | 1 | A |
| 70 | AATCCAGGTTGAACAA-1 | 3 | A |
| 71 | AATCCAGTCGACTGCG-1 | 5 | A |
| 72 | AATCGGTAGAAGGGTA-1 | 4 | A |
| 73 | AATCGGTCAGATGAAT-1 | 11 | D |
| 74 | AATCGGTCATGAATGA-1 | 4 | A |
| 75 | AATCGGTGTGCGCTTG-1 | 4 | A |
| 76 | ACACCAAGTGCAACCC-1 | 1 | A |
| 77 | ACACCCTAGCGTAATA-1 | 9 | D |
| 78 | ACACCCTCAACGCCGT-1 | 8 | D |
| 79 | ACACCGGAGTGTGAAT-1 | 3 | A |
| 80 | ACACCGGTCTGTGGCG-1 | 7 | C |
| 81 | ACACTGAAGACAGGCT-1 | 5 | A |
| 82 | ACACTGAAGCTCTCGG-1 | 10 | D |
| 83 | ACACTGACACAGACCC-1 | 10 | D |
| 84 | ACACTGACAGAGGCTA-1 | 4 | A |
| 85 | ACACTGACAGTTCCGG-1 | 5 | A |

| | | | |
|---|---|---|---|
| 86 | ACAGCCGAGTCTCGTA-1 | 1 | A |
| 88 | ACAGCCGGTTTGGCGC-1 | 4 | A |
| 91 | ACAGCTACAGAACAGC-1 | 8 | D |
| 92 | ACAGCTATCCGAAGCC-1 | 3 | A |
| 93 | ACATACGAGCCGAACA-1 | 8 | D |
| 94 | ACATACGAGGTAACGC-1 | 11 | D |
| 95 | ACATACGAGTTAGGGC-1 | 8 | D |
| 96 | ACATACGCACTCCGAG-1 | 1 | A |
| 97 | ACATACGGTATAAGTG-1 | 9 | D |
| 98 | ACATACGGTCTCTCGT-1 | 6 | B |
| 99 | ACATCAGAGTGTACGG-1 | 11 | D |
| 100 | ACATCAGAGTTAGGGC-1 | 3 | A |
| 101 | ACATGGTTCGAGCACC-1 | 10 | D |
| 102 | ACCAGTAAGATTCACC-1 | 6 | B |
| 103 | ACCAGTAGTGTTACGT-1 | 1 | A |
| 104 | ACCAGTATCGTTCGAA-1 | 8 | D |
| 105 | ACCCACTAGTTATCGC-1 | 3 | A |
| 106 | ACCCACTGTAGGGTGT-1 | 10 | D |
| 107 | ACCGTAACACGTAGGA-1 | 1 | A |
| 108 | ACCGTAACATCTCGTC-1 | 3 | A |
| 109 | ACCTTTAAGTGGGCTA-1 | 3 | A |
| 110 | ACGAGCCGTGCGCTTG-1 | 2 | A |
| 111 | ACGAGGACACCGGCAT-1 | 1 | A |
| 112 | ACGAGGACAGTAGGGT-1 | 5 | A |
| 113 | ACGAGGACATCTCGAA-1 | 4 | A |
| 114 | ACGATACAGAAGATTC-1 | 5 | A |
| 115 | ACGATACCAGCCAAAG-1 | 8 | D |
| 116 | ACGATACCAGTCCCTT-1 | 10 | D |
| 117 | ACGATACTCGAGGTGA-1 | 8 | D |
| 118 | ACGATGTCAACATGGG-1 | 1 | A |
| 119 | ACGATGTGTCATACCA-1 | 1 | A |
| 120 | ACGATGTTCTTCACAT-1 | 5 | A |
| 121 | ACGCAGCGTTACGGCC-1 | 3 | A |
| 122 | ACGCAGCTCCGCTCTA-1 | 11 | D |
| 123 | ACGCCAGCAAGAGAGA-1 | 11 | D |
| 124 | ACGCCAGCAAGGGTGT-1 | 6 | B |
| 125 | ACGCCAGCACAGCTGC-1 | 5 | A |
| 126 | ACGCCAGTCCACCCAT-1 | 5 | A |
| 127 | ACGCCAGTCTCGGAAT-1 | 10 | D |
| 128 | ACGCCGAAGACAAGCC-1 | 7 | C |
| 129 | ACGCCGAAGCAGCCTC-1 | 3 | A |
| 130 | ACGCCGACATGACCGC-1 | 1 | A |

| | | | |
|---|---|---|---|
| 132 | ACGGAGAAGCCGAACA-1 | 8 | D |
| 133 | ACGGAGAGTTGCAGCC-1 | 4 | A |
| 134 | ACGGAGATCGGATCCG-1 | 11 | D |
| 136 | ACGGCCAAGAATGTGT-1 | 10 | D |
| 137 | ACGGCCACAACGAAAT-1 | 1 | A |
| 138 | ACGGCCACACACACGC-1 | 10 | D |
| 139 | ACGGGCTAGTCTAGCT-1 | 10 | D |
| 140 | ACGGGCTCATCGGTCG-1 | 8 | D |
| 141 | ACGGGCTCATGCAGTT-1 | 11 | D |
| 142 | ACGGGCTGTCGTCTTC-1 | 7 | C |
| 143 | ACGGGTCCACGATGGA-1 | 11 | D |
| 144 | ACGGGTCTCGTCCCTA-1 | 2 | A |
| 145 | ACGTCAAAGATGTGGC-1 | 5 | A |
| 146 | ACGTCAAAGGAGCGAG-1 | 5 | A |
| 147 | ACGTCAACATTTGTGG-1 | 1 | A |
| 148 | ACTATCTAGAAGGCCT-1 | 11 | D |
| 149 | ACTATCTAGTACAGTA-1 | 3 | A |
| 150 | ACTATCTCATGGCTAT-1 | 5 | A |
| 152 | ACTATCTTCTCGAACA-1 | 5 | A |
| 153 | ACTATCTTCTGGTAGT-1 | 11 | D |
| 154 | ACTGAACAGCTAAACA-1 | 11 | D |
| 155 | ACTGAACAGTGCGATG-1 | 11 | D |
| 156 | ACTGAACCACGCGATC-1 | 5 | A |
| 157 | ACTGAGTCACGTTTAG-1 | 11 | D |
| 158 | ACTGAGTTCTTCTAAC-1 | 6 | B |
| 159 | ACTGATGCACGACTAT-1 | 10 | D |
| 160 | ACTGATGCATACCAGT-1 | 10 | D |
| 161 | ACTGATGGTACGTCAT-1 | 10 | D |
| 162 | ACTGATGTCGTTCGTC-1 | 4 | A |
| 163 | ACTGATGTCTTATGGG-1 | 2 | A |
| 164 | ACTGCTCAGATAGCAT-1 | 11 | D |
| 165 | ACTGCTCAGCGCCTTG-1 | 3 | A |
| 166 | ACTGCTCCAATCGTCA-1 | 8 | D |
| 167 | ACTGCTCGTGGAAAGA-1 | 3 | A |
| 168 | ACTTACTCACATAAAG-1 | 11 | D |
| 169 | ACTTACTTCCGAGATT-1 | 3 | A |
| 170 | ACTTACTTCTCATGTT-1 | 4 | A |
| 171 | ACTTGTTAGGGTTTCT-1 | 11 | D |
| 172 | ACTTGTTAGTTGTAGA-1 | 11 | D |
| 173 | ACTTGTTGTCTAGGTT-1 | 10 | D |
| 174 | ACTTGTTTCTCAGATG-1 | 3 | A |
| 175 | ACTTGTTTCTGTGGCG-1 | 11 | D |

| | | | |
|---|---|---|---|
| 176 | ACTTTCACAAACGAGC-1 | 1 | A |
| 177 | ACTTTCACACATTCAG-1 | 11 | D |
| 178 | ACTTTCACAGCGGTCT-1 | 10 | D |
| 179 | ACTTTCAGTAGCAGTG-1 | 3 | A |
| 180 | AGAATAGAGAAAGCTT-1 | 9 | D |
| 181 | AGAATAGAGTCAAGGC-1 | 3 | A |
| 182 | AGAATAGCAAGCAAGC-1 | 1 | A |
| 183 | AGAATAGGTGAAGCCA-1 | 3 | A |
| 184 | AGACGTTTCCTCAGTC-1 | 5 | A |
| 185 | AGAGCGAAGTCCATAC-1 | 3 | A |
| 186 | AGAGCGACAGTGTGGA-1 | 6 | B |
| 187 | AGAGCGAGTTTGTCAG-1 | 3 | A |
| 188 | AGAGCGATCCCACGGA-1 | 3 | A |
| 189 | AGAGCGATCGTGCCTT-1 | 4 | A |
| 190 | AGAGCTTAGACGTCGA-1 | 6 | B |
| 191 | AGAGCTTCATTCTTCA-1 | 3 | A |
| 192 | AGAGTGGAGGCATGGT-1 | 8 | D |
| 193 | AGAGTGGCACTACATG-1 | 6 | B |
| 194 | AGAGTGGCATGAAAGT-1 | 8 | D |
| 195 | AGAGTGGGTACAAGGC-1 | 11 | D |
| 196 | AGAGTGGGTTCTTCGC-1 | 11 | D |
| 197 | AGATCTGCACCAGTGC-1 | 1 | A |
| 198 | AGATCTGTCTTCTTGA-1 | 11 | D |
| 199 | AGATTGCAGCAAAGTT-1 | 4 | A |
| 200 | AGATTGCCAGAATTCC-1 | 11 | D |
| 201 | AGATTGCCATGATGCT-1 | 10 | D |
| 202 | AGATTGCGTCACCGCA-1 | 2 | A |
| 203 | AGATTGCTCGCGAGAA-1 | 3 | A |
| 204 | AGCAGCCAGGGTCGAT-1 | 7 | C |
| 205 | AGCAGCCAGTGTGATA-1 | 11 | D |
| 206 | AGCAGCCGTTGACGGA-1 | 11 | D |
| 207 | AGCAGCCTCTCATGTT-1 | 8 | D |
| 208 | AGCATACCAGGTACGA-1 | 3 | A |
| 209 | AGCATACCATCATGGT-1 | 3 | A |
| 210 | AGCCTAAAGTAAGTAC-1 | 10 | D |
| 211 | AGCCTAACACGTCATA-1 | 3 | A |
| 212 | AGCCTAAGTCTCTCGT-1 | 3 | A |
| 213 | AGCCTAAGTGTCGCTG-1 | 1 | A |
| 214 | AGCCTAATCCTCATCG-1 | 4 | A |
| 215 | AGCGGTCAGCGTAGTG-1 | 8 | D |
| 216 | AGCGGTCCACAAGCGA-1 | 11 | D |
| 217 | AGCGGTCCAGGCCGTT-1 | 3 | A |

| | | | |
|---|---|---|---|
| 219 | AGCGGTCGTCATCTGA-1 | 6 | B |
| 220 | AGCGGTCTCATCGCCT-1 | 7 | C |
| 221 | AGCGTATAGGCTATCT-1 | 5 | A |
| 222 | AGCGTATCAGTGTGGA-1 | 8 | D |
| 223 | AGCGTCGCACTCACCT-1 | 9 | D |
| 224 | AGCTCCTAGACTTCCA-1 | 3 | A |
| 225 | AGCTCCTAGGATGCGT-1 | 5 | A |
| 226 | AGCTCCTCAGTACCTA-1 | 3 | A |
| 227 | AGCTCCTGTAGTAGTA-1 | 2 | A |
| 228 | AGCTCCTTCGAGTTGT-1 | 8 | D |
| 229 | AGCTCTCAGCCGGTAA-1 | 8 | D |
| 230 | AGCTCTCAGTTAGCGG-1 | 4 | A |
| 231 | AGCTCTCGTACAACGG-1 | 9 | D |
| 232 | AGCTCTCGTACTCCGG-1 | 1 | A |
| 233 | AGCTCTCGTGATTACC-1 | 10 | D |
| 235 | AGCTTGAAGGCCCGTT-1 | 10 | D |
| 238 | AGCTTGAGTCAACTGT-1 | 11 | D |
| 239 | AGGCCACAGATCTGAA-1 | 11 | D |
| 240 | AGGCCACAGTTTGCGT-1 | 3 | A |
| 241 | AGGCCACGTTGGCGTC-1 | 1 | A |
| 242 | AGGCCACTCAAGAGGC-1 | 1 | A |
| 243 | AGGCCACTCCACCGGA-1 | 11 | D |
| 244 | AGGCCACTCTAGCTAG-1 | 2 | A |
| 245 | AGGCCGTAGCACAGGT-1 | 9 | D |
| 246 | AGGCCGTCATAGTGAA-1 | 3 | A |
| 247 | AGGCCGTGTCTAGGTT-1 | 2 | A |
| 248 | AGGCCGTGTCTCTATT-1 | 2 | A |
| 249 | AGGCCGTGTGTGTATC-1 | 1 | A |
| 250 | AGGCCGTTCGTTAGGT-1 | 5 | A |
| 251 | AGGCCGTTCTGCACAA-1 | 1 | A |
| 252 | AGGGAGTTCGCTATGA-1 | 9 | D |
| 253 | AGGGATGAGGCATTGG-1 | 3 | A |
| 254 | AGGGATGCAAATCGCT-1 | 11 | D |
| 255 | AGGGATGGTATAAGTG-1 | 4 | A |
| 256 | AGGGATGTCGTACGCG-1 | 3 | A |
| 257 | AGGGATGTCTGGCCTT-1 | 4 | A |
| 258 | AGGGTGAGTCCTACCT-1 | 3 | A |
| 259 | AGGGTGAGTGACTGAG-1 | 5 | A |
| 260 | AGGTCATAGCCATGCC-1 | 2 | A |
| 262 | AGGTCATAGTGATTGA-1 | 1 | A |
| 263 | AGGTCATCAACATAAG-1 | 1 | A |
| 264 | AGGTCATTCGACAATC-1 | 3 | A |

| | | | |
|---|---|---|---|
| 265 | AGGTCATTCGCGGCAT-1 | 8 | D |
| 266 | AGGTCCGCAGTGTGGA-1 | 2 | A |
| 268 | AGGTCCGTCTCGAGCG-1 | 2 | A |
| 269 | AGTAGTCCAATCGCGC-1 | 11 | D |
| 270 | AGTCTTTAGTCTAGCT-1 | 6 | B |
| 271 | AGTCTTTGTTCCACAA-1 | 11 | D |
| 272 | AGTCTTTTCCGTCTAC-1 | 3 | A |
| 273 | AGTGAGGAGGCATGGT-1 | 6 | B |
| 274 | AGTGAGGCATTCCGAA-1 | 4 | A |
| 275 | AGTGAGGTCTGTATGG-1 | 2 | A |
| 276 | AGTGGGAAGCTGAAAT-1 | 9 | D |
| 277 | AGTGGGAGTCACCAGC-1 | 5 | A |
| 278 | AGTGGGATCTGACGCG-1 | 9 | D |
| 279 | AGTGGGATCTGGGACC-1 | 11 | D |
| 280 | AGTGTCACATGTAGCT-1 | 11 | D |
| 281 | AGTGTCAGTACGTCAT-1 | 10 | D |
| 282 | AGTGTCATCGAGCGTC-1 | 9 | D |
| 283 | AGTTGGTCAAACGAGC-1 | 4 | A |
| 284 | AGTTGGTCACTTCAGA-1 | 4 | A |
| 285 | AGTTGGTGTGTTCGAT-1 | 10 | D |
| 286 | ATAACGCCAACTGGAG-1 | 6 | B |
| 287 | ATAACGCCACTTTCCG-1 | 11 | D |
| 288 | ATAACGCGTGCCATTA-1 | 2 | A |
| 289 | ATAACGCGTGCTCGAC-1 | 8 | D |
| 290 | ATAACGCTCCACTTCG-1 | 1 | A |
| 291 | ATAAGAGAGCGCCTTG-1 | 1 | A |
| 292 | ATAAGAGAGTAGCCGA-1 | 5 | A |
| 293 | ATAAGAGCAGCGTCAC-1 | 1 | A |
| 294 | ATAAGAGTCCGTGCTT-1 | 11 | D |
| 295 | ATAGACCGTACCGCTG-1 | 2 | A |
| 296 | ATAGACCGTCCGAATT-1 | 1 | A |
| 297 | ATCACGAGTACCTAGT-1 | 8 | D |
| 298 | ATCATCTCATTACGCA-1 | 3 | A |
| 299 | ATCATCTGTGCGAAAC-1 | 7 | C |
| 300 | ATCATGGAGTCCAGGA-1 | 8 | D |
| 301 | ATCCGAACAGCCAAAG-1 | 7 | C |
| 302 | ATCCGAACAGTAATCC-1 | 4 | A |
| 303 | ATCCGAACATTCCGAA-1 | 3 | A |
| 304 | ATCGAGTGTCGGAAGT-1 | 1 | A |
| 305 | ATCGAGTTCTGTGAAC-1 | 10 | D |
| 306 | ATCTACTAGGGATGGG-1 | 10 | D |
| 307 | ATCTACTAGTCACGCC-1 | 3 | A |

| | | | |
|---|---|---|---|
| 308 | ATCTACTAGTTTGCGT-1 | 3 | A |
| 309 | ATCTACTCAGTGCTGC-1 | 4 | A |
| 310 | ATCTGCCCATTCGTTT-1 | 10 | D |
| 311 | ATCTGCCGTCATGCCG-1 | 6 | B |
| 312 | ATCTGCCTCCCGCATT-1 | 3 | A |
| 313 | ATGAGGGAGAGGTAGA-1 | 10 | D |
| 314 | ATGAGGGAGTTTGCGT-1 | 2 | A |
| 315 | ATGAGGGTCCGACCAG-1 | 3 | A |
| 316 | ATGCGATCATTACAGC-1 | 3 | A |
| 317 | ATGCGATGTCAAGTTC-1 | 10 | D |
| 318 | ATGCGATGTGATCCTA-1 | 8 | D |
| 319 | ATGCGATGTTTAGGAA-1 | 8 | D |
| 320 | ATGGGAGAGTCCCACG-1 | 3 | A |
| 321 | ATGGGAGGTACCGTGC-1 | 11 | D |
| 322 | ATGGGAGTCGGAGAGT-1 | 3 | A |
| 323 | ATGGGAGTCTTAAGGC-1 | 11 | D |
| 324 | ATGGGAGTCTTCTAAC-1 | 8 | D |
| 325 | ATGTGTGCAAAGTAAC-1 | 11 | D |
| 326 | ATGTGTGGTCTCGTCT-1 | 11 | D |
| 327 | ATTACTCAGTAATCCC-1 | 3 | A |
| 328 | ATTACTCCACACACGC-1 | 2 | A |
| 330 | ATTACTCGTAAAGTCA-1 | 1 | A |
| 331 | ATTACTCGTAGCGTCC-1 | 11 | D |
| 332 | ATTATCCAGCGATGAC-1 | 3 | A |
| 333 | ATTATCCAGGAGTACC-1 | 10 | D |
| 334 | ATTATCCTCCGATACA-1 | 6 | B |
| 336 | ATTCTACAGCTAAGAT-1 | 6 | B |
| 337 | ATTCTACAGCTATGCT-1 | 5 | A |
| 338 | ATTCTACCAGCGCTTG-1 | 9 | D |
| 339 | ATTCTACGTGGAAAGA-1 | 11 | D |
| 340 | ATTCTACTCTCCCTAG-1 | 11 | D |
| 341 | ATTGGACTCGTAGCGC-1 | 8 | D |
| 342 | ATTGGTGAGGGATACC-1 | 10 | D |
| 343 | ATTGGTGCACTAAGGG-1 | 11 | D |
| 344 | ATTGGTGTCGCCAACG-1 | 11 | D |
| 345 | ATTTCTGAGTAGGTGC-1 | 7 | C |
| 346 | ATTTCTGCACACACGC-1 | 3 | A |
| 347 | ATTTCTGCACTAACTG-1 | 2 | A |
| 348 | ATTTCTGTCACACGGC-1 | 6 | B |
| 349 | CAACCAAAGTACGCGA-1 | 1 | A |
| 351 | CAACCAAGTTCGATAC-1 | 11 | D |
| 352 | CAACCTCAGACTTCCA-1 | 5 | A |

| | | | |
|---|---|---|---|
| 353 | CAACCTCCAAGAACTA-1 | 4 | A |
| 354 | CAACCTCTCAACTGCA-1 | 3 | A |
| 355 | CAACCTCTCTGAAGCT-1 | 4 | A |
| 356 | CAACTAGGTCGTCTTC-1 | 11 | D |
| 358 | CAACTAGTCGACCATA-1 | 1 | A |
| 359 | CAAGAAAAGCAAAGCC-1 | 6 | B |
| 360 | CAAGAAACATAGGACG-1 | 6 | B |
| 361 | CAAGAAAGTTTGTCTT-1 | 3 | A |
| 362 | CAAGAAATCGAACAAA-1 | 3 | A |
| 364 | CAAGATCAGGTGTTAA-1 | 6 | B |
| 366 | CAAGATCCATTACGCA-1 | 6 | B |
| 367 | CAAGATCGTGCTTAGT-1 | 4 | A |
| 368 | CAAGATCGTGTTCGAT-1 | 10 | D |
| 370 | CAAGATCTCACACGTA-1 | 8 | D |
| 371 | CAAGGCCCAAACCCTA-1 | 11 | D |
| 372 | CAAGGCCGTCACCTGG-1 | 3 | A |
| 373 | CAAGTTGTCGCGGCAT-1 | 5 | A |
| 374 | CACAAACCAAGAATCA-1 | 8 | D |
| 375 | CACAAACCAGGCCTTG-1 | 6 | B |
| 376 | CACAAACCATCAAGAA-1 | 2 | A |
| 377 | CACAAACTCGAGAGGT-1 | 5 | A |
| 378 | CACACAAGTACGTTTG-1 | 1 | A |
| 379 | CACACAAGTTGCACTA-1 | 11 | D |
| 380 | CACACCTGTACGTGCC-1 | 10 | D |
| 381 | CACACTCAGACGTCGA-1 | 11 | D |
| 382 | CACACTCAGCGTAGTG-1 | 11 | D |
| 383 | CACACTCGTTAAAGGT-1 | 5 | A |
| 384 | CACACTCTCTACAGTG-1 | 1 | A |
| 386 | CACAGGCCATAGTGTC-1 | 5 | A |
| 387 | CACAGGCGTAGTCTGT-1 | 11 | D |
| 388 | CACAGGCTCCGTCTAC-1 | 7 | C |
| 389 | CACAGTAAGTCGTACT-1 | 5 | A |
| 390 | CACATAGAGGTATCGG-1 | 6 | B |
| 391 | CACATAGAGTCGTCCG-1 | 11 | D |
| 392 | CACATAGCACAACTTG-1 | 9 | D |
| 393 | CACATAGGTACAATAG-1 | 4 | A |
| 394 | CACATAGGTTGGTGAG-1 | 5 | A |
| 395 | CACATAGTCTCAGTTT-1 | 3 | A |
| 396 | CACATTTAGCAGCGTA-1 | 3 | A |
| 397 | CACATTTAGGAATCGC-1 | 3 | A |
| 398 | CACATTTCATGGAGAC-1 | 1 | A |
| 399 | CACATTTGTCGCTTCT-1 | 10 | D |

| | | | |
|---|---|---|---|
| 400 | CACATTTGTTTCCTGC-1 | 1 | A |
| 401 | CACCACTCATCTAAGC-1 | 3 | A |
| 402 | CACCACTGTCATGTTG-1 | 11 | D |
| 403 | CACCAGGAGACAGTTA-1 | 11 | D |
| 404 | CACCAGGAGGAGTAGA-1 | 2 | A |
| 405 | CACCAGGCAATCTCTT-1 | 11 | D |
| 406 | CACCAGGCACGGCTGT-1 | 11 | D |
| 407 | CACCAGGCATTACTTC-1 | 3 | A |
| 408 | CACCTTGAGGCTCTCG-1 | 11 | D |
| 409 | CACCTTGCAGACACAG-1 | 3 | A |
| 410 | CACCTTGCAGACAGTG-1 | 7 | C |
| 411 | CACCTTGTCCGACGTG-1 | 8 | D |
| 412 | CACTCCAAGCAGACTG-1 | 3 | A |
| 413 | CACTCCAAGTGTCCAT-1 | 10 | D |
| 415 | CAGAATCTCGAAAGGC-1 | 7 | C |
| 416 | CAGAATCTCGGAATGG-1 | 2 | A |
| 417 | CAGAATCTCTACGGGC-1 | 11 | D |
| 418 | CAGAGAGAGTCGTACT-1 | 1 | A |
| 419 | CAGAGAGAGTGAAGAG-1 | 8 | D |
| 420 | CAGAGAGCACGAGCTC-1 | 8 | D |
| 421 | CAGAGAGTCCGAAGCC-1 | 8 | D |
| 422 | CAGAGAGTCCGAGATT-1 | 11 | D |
| 423 | CAGATCAAGAAGGGTA-1 | 6 | B |
| 425 | CAGATCAGTTCTGTCC-1 | 5 | A |
| 426 | CAGCAGCGTTCAAGGG-1 | 8 | D |
| 428 | CAGCAGCTCGCAACGC-1 | 1 | A |
| 429 | CAGCATAAGCGTTGCC-1 | 10 | D |
| 430 | CAGCATACAAACGGGT-1 | 5 | A |
| 431 | CAGCATACACGGAATG-1 | 9 | D |
| 432 | CAGCATAGTCCGCAGT-1 | 10 | D |
| 433 | CAGCCGAAGCGTCAAG-1 | 2 | A |
| 435 | CAGCGACAGCGCCTCA-1 | 8 | D |
| 436 | CAGCGACCAAGTCGGA-1 | 8 | D |
| 437 | CAGCTAATCCACTTCG-1 | 11 | D |
| 438 | CAGCTAATCGAAAGGC-1 | 1 | A |
| 439 | CAGCTGGAGCCACCTG-1 | 2 | A |
| 440 | CAGCTGGGTCCTTGGG-1 | 3 | A |
| 441 | CAGCTGGTCACGGTAT-1 | 5 | A |
| 442 | CAGGTGCCAGCATGTT-1 | 11 | D |
| 443 | CAGGTGCTCGAGGTGA-1 | 8 | D |
| 444 | CAGTAACAGACCACGA-1 | 5 | A |
| 445 | CAGTAACAGCACCGCT-1 | 8 | D |

| | | | |
|---|---|---|---|
| 446 | CAGTAACCACACCTAA-1 | 2 | A |
| 447 | CAGTAACCATACGTTG-1 | 11 | D |
| 448 | CAGTCCTAGTCAAGGC-1 | 3 | A |
| 449 | CAGTCCTGTCTTCACC-1 | 5 | A |
| 450 | CATATGGTCTAGACAC-1 | 4 | A |
| 451 | CATATTCCAGTAGAAT-1 | 1 | A |
| 452 | CATATTCGTCGCGCTA-1 | 3 | A |
| 453 | CATATTCGTGACCTGC-1 | 11 | D |
| 454 | CATATTCTCTTGCTTA-1 | 4 | A |
| 455 | CATCAAGTCCTACGAA-1 | 8 | D |
| 456 | CATCAAGTCTAGCCTC-1 | 3 | A |
| 457 | CATCAAGTCTCCATGC-1 | 11 | D |
| 458 | CATCAGAGTTAGTGAA-1 | 5 | A |
| 459 | CATCCACCATTACAGC-1 | 10 | D |
| 460 | CATCCACTCCTGCGTT-1 | 10 | D |
| 461 | CATCCACTCCTTGAAG-1 | 3 | A |
| 462 | CATCGAACAATGGGAC-1 | 4 | A |
| 464 | CATCGAACATTTCGTT-1 | 10 | D |
| 465 | CATCGAAGTGCTCGAC-1 | 8 | D |
| 466 | CATCGAAGTTTGGCGC-1 | 4 | A |
| 467 | CATCGGGCACCCAAAT-1 | 10 | D |
| 468 | CATCGGGTCTCAGTTT-1 | 3 | A |
| 469 | CATGACACAACTCGCG-1 | 2 | A |
| 471 | CATGACATCTCCGGAG-1 | 8 | D |
| 472 | CATGCCTGTCCGCAAC-1 | 7 | C |
| 473 | CATGCCTGTTACGGCC-1 | 3 | A |
| 474 | CATGCCTTCATGTGAC-1 | 11 | D |
| 475 | CATGGCGAGATATGGT-1 | 4 | A |
| 476 | CATGGCGCAGTCGGAA-1 | 9 | D |
| 477 | CATGGCGCATTTGCGA-1 | 6 | B |
| 478 | CATGGCGTCGAGCACC-1 | 11 | D |
| 479 | CATTATCAGCGTGAAC-1 | 3 | A |
| 480 | CATTATCCAACAGCCC-1 | 10 | D |
| 482 | CATTATCGTAAGAAGG-1 | 9 | D |
| 483 | CATTCGCAGCAAATCA-1 | 1 | A |
| 484 | CATTCGCCACGCACGT-1 | 7 | C |
| 485 | CCAATCCGTCAAGCCC-1 | 6 | B |
| 487 | CCACCTAAGCTCCTTC-1 | 10 | D |
| 488 | CCACCTAAGTGGTAGC-1 | 5 | A |
| 490 | CCACCTAGTTCGAGCC-1 | 6 | B |
| 491 | CCACGGAAGACTGTAA-1 | 3 | A |
| 493 | CCACGGACAGAAGACC-1 | 11 | D |

| | | | |
|---|---|---|---|
| 494 | CCACTACAGACAAAGG-1 | 10 | D |
| 496 | CCACTACTCTACGGCG-1 | 2 | A |
| 497 | CCACTACTCTCGGAAT-1 | 10 | D |
| 498 | CCAGCGATCTACAGTG-1 | 2 | A |
| 499 | CCATGTCTCGAGTACT-1 | 5 | A |
| 500 | CCATGTCTCGCTGAAT-1 | 11 | D |
| 501 | CCATTCGAGTGCTGCC-1 | 1 | A |
| 502 | CCATTCGCACATAAAG-1 | 8 | D |
| 503 | CCCAATCGTCGACCAC-1 | 6 | B |
| 504 | CCCAGTTAGCACGCCT-1 | 11 | D |
| 505 | CCCATACAGCCACGTC-1 | 6 | B |
| 506 | CCCATACCAACGAAAT-1 | 6 | B |
| 507 | CCCATACCACAAGCTT-1 | 3 | A |
| 508 | CCCATACCATCTTTCA-1 | 3 | A |
| 510 | CCCTCCTCAGAATTCC-1 | 3 | A |
| 511 | CCCTCCTGTATTCGCA-1 | 8 | D |
| 512 | CCGGGATCACCAATGT-1 | 1 | A |
| 513 | CCGGTAGAGCCAGTTT-1 | 6 | B |
| 514 | CCGGTAGAGCCTATGT-1 | 9 | D |
| 515 | CCGGTAGGTTCAGTAC-1 | 5 | A |
| 516 | CCGGTAGTCGAGTTGT-1 | 5 | A |
| 517 | CCGGTAGTCGCGATGC-1 | 8 | D |
| 518 | CCGTACTAGGCTCATT-1 | 11 | D |
| 519 | CCGTACTTCGGAGAGT-1 | 6 | B |
| 520 | CCGTGGAAGTAGATGT-1 | 8 | D |
| 521 | CCGTGGACACCAAAGG-1 | 9 | D |
| 522 | CCGTGGACACTGGCCA-1 | 3 | A |
| 523 | CCGTGGAGTCAACCGC-1 | 2 | A |
| 524 | CCGTTCAAGTCGCCGT-1 | 11 | D |
| 525 | CCTAAAGGTACCGTGC-1 | 9 | D |
| 526 | CCTAAAGTCAAGCCGC-1 | 1 | A |
| 527 | CCTACACAGGGATCTG-1 | 2 | A |
| 528 | CCTACACCAACCGCTG-1 | 2 | A |
| 529 | CCTACACGTGGAAGGG-1 | 8 | D |
| 530 | CCTACACTCGCGATGC-1 | 9 | D |
| 531 | CCTACACTCGTTATTC-1 | 11 | D |
| 532 | CCTACACTCTCAGATG-1 | 2 | A |
| 533 | CCTACCATCGGAGAGT-1 | 6 | B |
| 534 | CCTAGCTCAATCCGTA-1 | 1 | A |
| 535 | CCTAGCTTCATGTCAG-1 | 3 | A |
| 536 | CCTAGCTTCTACATGG-1 | 4 | A |
| 537 | CCTATTAAGCCAACAG-1 | 2 | A |

| | | | |
|---|---|---|---|
| 538 | CCTATTACAACAGCCC-1 | 2 | A |
| 539 | CCTATTACACGAGTTT-1 | 8 | D |
| 540 | CCTATTAGTATTCGTG-1 | 3 | A |
| 541 | CCTATTAGTGACCCAC-1 | 7 | C |
| 542 | CCTATTATCGAACCCG-1 | 4 | A |
| 543 | CCTCTGACATGGGCCT-1 | 5 | A |
| 544 | CCTTACGAGTGACTCT-1 | 7 | C |
| 545 | CCTTACGCAAGGTATA-1 | 3 | A |
| 546 | CCTTACGCATACGTTG-1 | 1 | A |
| 547 | CCTTACGTCATCAGCA-1 | 11 | D |
| 548 | CCTTCCCCAACCCTGG-1 | 1 | A |
| 549 | CCTTCCCCAACGTCTA-1 | 4 | A |
| 550 | CCTTCCCGTTCAGGTT-1 | 8 | D |
| 551 | CCTTCCCTCGCGGACT-1 | 10 | D |
| 553 | CCTTCGAAGTCGTCCG-1 | 11 | D |
| 554 | CCTTCGATCAGCAGCC-1 | 7 | C |
| 555 | CCTTCGATCAGTACCA-1 | 10 | D |
| 556 | CGAACATAGGCTAGTG-1 | 8 | D |
| 557 | CGAACATGTGACTGAG-1 | 10 | D |
| 558 | CGAACATTCAGTACCA-1 | 7 | C |
| 559 | CGAACATTCTGGGATT-1 | 11 | D |
| 560 | CGAATGTAGTACGACG-1 | 3 | A |
| 561 | CGAATGTCAGACAGTG-1 | 11 | D |
| 562 | CGAATGTCAGCCCATG-1 | 4 | A |
| 563 | CGAATGTGTGTCAACT-1 | 5 | A |
| 564 | CGACCTTAGGGTAACC-1 | 8 | D |
| 565 | CGACCTTAGTGTTTGC-1 | 3 | A |
| 566 | CGACCTTCAGTCGGAA-1 | 3 | A |
| 567 | CGACTTCGTGGTGCAA-1 | 3 | A |
| 568 | CGAGAAGCAAACTCAC-1 | 10 | D |
| 570 | CGAGCACAGGCGATAC-1 | 3 | A |
| 572 | CGAGCACCAGATATCC-1 | 7 | C |
| 573 | CGAGCACGTCGAGTAG-1 | 9 | D |
| 574 | CGAGCACTCAAGTTGC-1 | 8 | D |
| 575 | CGAGCACTCGGAAGAC-1 | 3 | A |
| 576 | CGAGCACTCTGATCAG-1 | 6 | B |
| 577 | CGAGCCAAGCTGGTCC-1 | 5 | A |
| 579 | CGAGCCACAAGAACTA-1 | 3 | A |
| 580 | CGAGCCACAGACGAGT-1 | 11 | D |
| 581 | CGAGCCACAGGACCAA-1 | 6 | B |
| 582 | CGAGCCACATAGTGAA-1 | 8 | D |
| 583 | CGATCGGGTACCGTGC-1 | 1 | A |

| | | | |
|---|---|---|---|
| 586 | CGATGTAAGATCGGGT-1 | 5 | A |
| 587 | CGATGTACAAGTACAA-1 | 1 | A |
| 588 | CGATGTAGTCGCCAAC-1 | 2 | A |
| 589 | CGATGTATCTCCAGCT-1 | 6 | B |
| 590 | CGATTGAAGCTAGCCC-1 | 11 | D |
| 591 | CGATTGACACCTCCAT-1 | 6 | B |
| 592 | CGATTGAGTAAGAAGG-1 | 2 | A |
| 593 | CGATTGAGTACTCCGG-1 | 3 | A |
| 594 | CGATTGAGTATTCGCA-1 | 2 | A |
| 595 | CGATTGAGTCAACTCA-1 | 1 | A |
| 596 | CGATTGAGTGCATCTA-1 | 1 | A |
| 599 | CGCCAAGAGGCTCAGA-1 | 8 | D |
| 601 | CGCGGTAGTCGGAAGT-1 | 8 | D |
| 602 | CGCGTTTCACAAGCTT-1 | 11 | D |
| 604 | CGCGTTTTCTGGATCA-1 | 8 | D |
| 606 | CGCTATCGTGATTGTC-1 | 1 | A |
| 607 | CGCTATCTCCGGCTAG-1 | 11 | D |
| 608 | CGCTATCTCGACTTCC-1 | 8 | D |
| 609 | CGCTTCACAAGCACGA-1 | 3 | A |
| 610 | CGCTTCACACGAAGGT-1 | 6 | B |
| 611 | CGCTTCAGTTCTAGTG-1 | 1 | A |
| 612 | CGGACACAGACGCTTT-1 | 10 | D |
| 613 | CGGACACGTAGTAGTA-1 | 6 | B |
| 614 | CGGACACGTCTAGGTT-1 | 11 | D |
| 615 | CGGACACTCGAGTTGT-1 | 6 | B |
| 616 | CGGACGTCACTTGCTC-1 | 2 | A |
| 618 | CGGAGCTAGGCTACGA-1 | 11 | D |
| 619 | CGGAGCTGTCCGAAGA-1 | 11 | D |
| 620 | CGGAGCTTCTAGACAC-1 | 10 | D |
| 621 | CGGAGTCAGAAACCTA-1 | 8 | D |
| 622 | CGGAGTCAGCATGGCA-1 | 9 | D |
| 623 | CGGAGTCAGCCAACAG-1 | 1 | A |
| 624 | CGGAGTCCAGAGGCAT-1 | 5 | A |
| 625 | CGGAGTCCAGCTTGAT-1 | 5 | A |
| 626 | CGGAGTCGTATCAGTC-1 | 8 | D |
| 627 | CGGAGTCGTCTTCACC-1 | 11 | D |
| 628 | CGGCTAGAGACGCACA-1 | 8 | D |
| 629 | CGGCTAGAGTAGCCGA-1 | 11 | D |
| 630 | CGGCTAGAGTAGTATG-1 | 3 | A |
| 631 | CGGCTAGGTGTCAACT-1 | 6 | B |
| 632 | CGGCTAGGTTCTTCGC-1 | 10 | D |
| 633 | CGGCTAGTCATGTGAC-1 | 8 | D |

| | | | |
|---|---|---|---|
| 634 | CGGGTCAGTCATTACG-1 | 3 | A |
| 635 | CGGTTAACACCACATA-1 | 11 | D |
| 636 | CGGTTAACATGGAGAC-1 | 9 | D |
| 637 | CGTAGCGAGGACAGAA-1 | 2 | A |
| 638 | CGTAGCGAGTACGATA-1 | 6 | B |
| 639 | CGTAGCGCAACGGGAT-1 | 3 | A |
| 640 | CGTAGGCAGGCGTTGA-1 | 3 | A |
| 641 | CGTAGGCCACTTGTTT-1 | 3 | A |
| 642 | CGTAGGCCATGAGGGT-1 | 3 | A |
| 644 | CGTCACTAGGCGCTCT-1 | 5 | A |
| 645 | CGTCACTAGTCCCACG-1 | 8 | D |
| 646 | CGTCACTCATTCCGAA-1 | 3 | A |
| 647 | CGTCACTTCGTTACCC-1 | 11 | D |
| 649 | CGTCAGGAGCAGGCTA-1 | 6 | B |
| 650 | CGTCAGGAGGACAGCT-1 | 1 | A |
| 651 | CGTCAGGCACGAGCTC-1 | 11 | D |
| 652 | CGTCCATGTAGATGTA-1 | 3 | A |
| 653 | CGTCCATTCTCCACAC-1 | 3 | A |
| 654 | CGTCTACCAACGACTT-1 | 9 | D |
| 656 | CGTCTACCACAGCTGC-1 | 10 | D |
| 657 | CGTCTACGTGGAGCTA-1 | 8 | D |
| 658 | CGTCTACTCTTAAGGC-1 | 11 | D |
| 659 | CGTGAGCTCGAATGTC-1 | 10 | D |
| 661 | CGTGTAAGTAGGTTTC-1 | 11 | D |
| 662 | CGTGTAAGTTTCCGAA-1 | 1 | A |
| 663 | CGTGTCTCAACCGCTG-1 | 6 | B |
| 664 | CGTGTCTCACAGCTAT-1 | 11 | D |
| 665 | CGTGTCTCATCCATCC-1 | 8 | D |
| 666 | CGTGTCTTCCAACAAC-1 | 6 | B |
| 667 | CGTGTCTTCCGAGTCG-1 | 9 | D |
| 668 | CGTGTCTTCGTTGCAA-1 | 8 | D |
| 670 | CGTTAGAAGGGTAACC-1 | 8 | D |
| 671 | CGTTAGAAGTGGCACA-1 | 3 | A |
| 672 | CGTTAGAGTCATCCAA-1 | 11 | D |
| 673 | CGTTAGATCAGCCTGG-1 | 3 | A |
| 674 | CGTTCTGAGCTGCGAA-1 | 1 | A |
| 675 | CGTTCTGAGGCTAGCA-1 | 4 | A |
| 676 | CGTTCTGGTTGAACAA-1 | 10 | D |
| 677 | CGTTCTGTCTCTAGTT-1 | 11 | D |
| 678 | CGTTGGGAGGCGCTCT-1 | 11 | D |
| 679 | CGTTGGGAGTCCCACG-1 | 5 | A |
| 680 | CTAACTTAGGGATGGG-1 | 10 | D |

| | | | |
|---|---|---|---|
| 681 | CTAACTTGTTGACGGA-1 | 11 | D |
| 682 | CTAACTTTCTTGGCCT-1 | 3 | A |
| 683 | CTAAGACGTGATTGTC-1 | 11 | D |
| 684 | CTAATGGCAACCTGAT-1 | 4 | A |
| 685 | CTACACCAGTTACCCA-1 | 1 | A |
| 686 | CTACACCGTACCGTGC-1 | 8 | D |
| 687 | CTACATTCACGAGATG-1 | 8 | D |
| 688 | CTACATTTCAAAGTGA-1 | 3 | A |
| 689 | CTACATTTCACGTTAG-1 | 8 | D |
| 690 | CTACCCACAGTAGGGT-1 | 10 | D |
| 691 | CTACCCAGTAGAACAT-1 | 8 | D |
| 692 | CTACCCAGTATATGAG-1 | 11 | D |
| 693 | CTACGTCGTAGTGGCA-1 | 3 | A |
| 694 | CTAGAGTTCGACAATC-1 | 9 | D |
| 695 | CTAGCCTAGGGATACC-1 | 10 | D |
| 696 | CTAGCCTTCCAACGAT-1 | 6 | B |
| 697 | CTAGCCTTCTCCGGAG-1 | 5 | A |
| 698 | CTAGCCTTCTTGGATG-1 | 3 | A |
| 699 | CTAGCCTTCTTGTCCG-1 | 11 | D |
| 700 | CTAGTGACAGTTTGCA-1 | 7 | C |
| 701 | CTAGTGAGTAGGGTGT-1 | 8 | D |
| 703 | CTCACACCATAGGTCT-1 | 2 | A |
| 704 | CTCACACGTCTCTCTG-1 | 3 | A |
| 705 | CTCACACTCAAGAGGC-1 | 3 | A |
| 706 | CTCAGAACATGGAGAC-1 | 9 | D |
| 707 | CTCAGAATCATGACCA-1 | 3 | A |
| 708 | CTCATTAGTGTTCCGG-1 | 10 | D |
| 710 | CTCATTATCTGATCAG-1 | 5 | A |
| 711 | CTCCTAGAGGAGTACC-1 | 5 | A |
| 712 | CTCCTAGCAAACCGAG-1 | 9 | D |
| 713 | CTCGAAAAGCAGGTCA-1 | 5 | A |
| 714 | CTCGAAACAAGCTACT-1 | 4 | A |
| 715 | CTCGAAACATTTGTGG-1 | 7 | C |
| 716 | CTCGAAATCTACAAGC-1 | 10 | D |
| 717 | CTCGAGGAGCGATATA-1 | 11 | D |
| 718 | CTCGAGGAGTGACATA-1 | 7 | C |
| 719 | CTCGAGGCAACTAGTC-1 | 3 | A |
| 720 | CTCGAGGCAAGAATCA-1 | 1 | A |
| 721 | CTCGAGGCAAGTGTTC-1 | 10 | D |
| 722 | CTCGAGGGTCATGTTG-1 | 6 | B |
| 723 | CTCGGAGAGAGCAATT-1 | 3 | A |
| 724 | CTCGGAGAGCTAACAA-1 | 2 | A |

| | | | |
|---|---|---|---|
| 725 | CTCGGAGTCCACTAAG-1 | 8 | D |
| 726 | CTCGGGAAGTGTTAGA-1 | 8 | D |
| 727 | CTCGGGACAAGGGTGT-1 | 7 | C |
| 728 | CTCGGGATCATATCTC-1 | 5 | A |
| 729 | CTCGTACAGAGCTTCT-1 | 8 | D |
| 730 | CTCGTACAGCTGCGAA-1 | 6 | B |
| 731 | CTCGTACCACGTACAT-1 | 3 | A |
| 732 | CTCGTACCACGTTTAG-1 | 3 | A |
| 733 | CTCGTACCATGCCATA-1 | 2 | A |
| 734 | CTCGTACGTTGCCTCT-1 | 11 | D |
| 735 | CTCGTCAAGCGGCTTC-1 | 3 | A |
| 736 | CTCGTCACAACGACGA-1 | 6 | B |
| 737 | CTCGTCACAAGGTGAC-1 | 4 | A |
| 738 | CTCGTCAGTGTCCCTT-1 | 5 | A |
| 739 | CTCGTCATCACTGCGG-1 | 6 | B |
| 740 | CTCTAATGTATCGCCG-1 | 3 | A |
| 741 | CTCTAATTCATATCTC-1 | 10 | D |
| 742 | CTCTACGAGAAAGTGG-1 | 11 | D |
| 743 | CTCTACGGTGATTGGG-1 | 10 | D |
| 744 | CTGAAGTAGTGATTGA-1 | 9 | D |
| 745 | CTGAAGTCAGACTTGT-1 | 9 | D |
| 746 | CTGAAGTGTAACGGCA-1 | 11 | D |
| 747 | CTGAAGTGTCAACCGC-1 | 2 | A |
| 748 | CTGAAGTGTCCCACAG-1 | 11 | D |
| 749 | CTGAAGTTCGTAGCGC-1 | 8 | D |
| 750 | CTGATAGAGCAGCGTA-1 | 6 | B |
| 751 | CTGATAGCAGCTCATA-1 | 3 | A |
| 752 | CTGATAGTCAGAAACA-1 | 8 | D |
| 753 | CTGATCCGTTCGTCAA-1 | 3 | A |
| 754 | CTGCCTAGTGGTCAGA-1 | 11 | D |
| 755 | CTGCCTAGTTCCACAA-1 | 7 | C |
| 756 | CTGCCTATCGTAGCGC-1 | 3 | A |
| 757 | CTGCGGAAGAGTGACC-1 | 11 | D |
| 758 | CTGCGGAAGATGTCTC-1 | 10 | D |
| 759 | CTGCGGAAGTTGTCGT-1 | 6 | B |
| 760 | CTGCGGAGTCTCCACT-1 | 11 | D |
| 761 | CTGCGGATCAGAAACA-1 | 6 | B |
| 762 | CTGCGGATCCAAAGAA-1 | 4 | A |
| 763 | CTGCGGATCCCAGAAT-1 | 8 | D |
| 764 | CTGCTGTAGTAGGCCA-1 | 3 | A |
| 765 | CTGCTGTAGTCTCGGC-1 | 1 | A |
| 766 | CTGCTGTCAACATAAG-1 | 3 | A |

| | | | |
|---|---|---|---|
| 767 | CTGGTCTAGCGATATA-1 | 1 | A |
| 768 | CTGGTCTAGGTCGGAT-1 | 6 | B |
| 770 | CTGGTCTTCTCCCTAG-1 | 3 | A |
| 771 | CTGTGCTCAATGGGAC-1 | 3 | A |
| 772 | CTGTGCTCAGACACGA-1 | 1 | A |
| 773 | CTGTGCTCATGGCTCG-1 | 2 | A |
| 774 | CTGTGCTGTGGGCCTT-1 | 3 | A |
| 775 | CTGTTTAAGAGTCTGG-1 | 8 | D |
| 776 | CTGTTTAAGCTGCCCA-1 | 11 | D |
| 777 | CTGTTTAAGGCTATCT-1 | 10 | D |
| 778 | CTGTTTACACGCCTGA-1 | 6 | B |
| 779 | CTGTTTAGTCTCTTTA-1 | 5 | A |
| 780 | CTGTTTATCACCAGCG-1 | 2 | A |
| 781 | CTTAACTAGCATCTTG-1 | 1 | A |
| 782 | CTTAACTGTGGAAAGA-1 | 3 | A |
| 783 | CTTAACTTCCTTGCAC-1 | 5 | A |
| 784 | CTTACCGAGGGCTTCC-1 | 6 | B |
| 785 | CTTACCGGTAAGTAAC-1 | 5 | A |
| 786 | CTTACCGGTCACCTGG-1 | 8 | D |
| 787 | CTTACCGGTGCTATTG-1 | 8 | D |
| 788 | CTTACCGTCACACGTA-1 | 8 | D |
| 789 | CTTACCGTCCGCGACA-1 | 4 | A |
| 790 | CTTAGGACACACGCAC-1 | 5 | A |
| 791 | CTTAGGATCTGGGCTG-1 | 11 | D |
| 792 | CTTCTCTAGCTGCCCA-1 | 10 | D |
| 793 | CTTCTCTAGGTCGGAT-1 | 11 | D |
| 794 | CTTCTCTCAATGTAGA-1 | 9 | D |
| 795 | CTTCTCTCAGCTATGT-1 | 11 | D |
| 796 | CTTCTCTCATCGCATA-1 | 2 | A |
| 797 | CTTCTCTTCCTATGAG-1 | 1 | A |
| 798 | CTTGGCTAGACAGAGA-1 | 5 | A |
| 799 | CTTGGCTAGCGCCTTG-1 | 8 | D |
| 800 | CTTGGCTAGTCTCGTA-1 | 2 | A |
| 801 | CTTGGCTGTTCTAGTG-1 | 9 | D |
| 802 | CTTTGCGGTCAAAGCG-1 | 8 | D |
| 803 | CTTTGCGTCCTCAGTC-1 | 4 | A |
| 804 | GAAACTCGTGCGAACA-1 | 11 | D |
| 805 | GAAACTCGTTCGTCAA-1 | 5 | A |
| 806 | GAAACTCTCACGCTCT-1 | 5 | A |
| 807 | GAAACTCTCACGTAGT-1 | 1 | A |
| 808 | GAAACTCTCTGGTGAT-1 | 7 | C |
| 809 | GAAATGAAGGCTCTTA-1 | 9 | D |

| | | | |
|---|---|---|---|
| 810 | GAAATGACACCCTCTA-1 | 5 | A |
| 811 | GAACATCGTACTTGCA-1 | 11 | D |
| 812 | GAACATCTCATGCGCG-1 | 2 | A |
| 813 | GAACCTACAAAGGTTA-1 | 7 | C |
| 814 | GAACCTACAATCGCGC-1 | 1 | A |
| 815 | GAACCTAGTCACATGT-1 | 10 | D |
| 816 | GAACCTAGTCGAGTAG-1 | 4 | A |
| 817 | GAACCTAGTTCGAGCC-1 | 1 | A |
| 818 | GAACGGACACTTGCAA-1 | 11 | D |
| 820 | GAACGGAGTACGTTTG-1 | 6 | B |
| 821 | GAACGGATCTTCACGC-1 | 11 | D |
| 822 | GAAGCAGAGGGAGTAA-1 | 10 | D |
| 823 | GAAGCAGCACACGCAC-1 | 5 | A |
| 824 | GAATAAGAGGCTACGA-1 | 1 | A |
| 825 | GAATAAGCAAATACGA-1 | 8 | D |
| 826 | GAATAAGCAGAGATCG-1 | 6 | B |
| 827 | GAATGAACAACCTGCG-1 | 3 | A |
| 828 | GAATGAATCTTCACAT-1 | 3 | A |
| 829 | GACACGCCAAATGATG-1 | 11 | D |
| 830 | GACAGAGAGCCTAACT-1 | 10 | D |
| 831 | GACAGAGCACAGACCC-1 | 11 | D |
| 832 | GACAGAGCAGTAATCC-1 | 10 | D |
| 833 | GACAGAGCATCGGGCT-1 | 1 | A |
| 834 | GACAGAGTCAGCCCGA-1 | 1 | A |
| 835 | GACAGAGTCCTACGAA-1 | 8 | D |
| 836 | GACAGAGTCCTGCGTT-1 | 7 | C |
| 837 | GACCAATAGACTTTCG-1 | 3 | A |
| 838 | GACCAATAGGACAGAA-1 | 11 | D |
| 839 | GACCAATTCCGGCTAG-1 | 3 | A |
| 840 | GACCTGGGTAGCCCTG-1 | 8 | D |
| 841 | GACGCGTAGTAACCCT-1 | 9 | D |
| 842 | GACGCGTCAGGTCAAG-1 | 9 | D |
| 843 | GACGGCTGTGTAATGA-1 | 2 | A |
| 844 | GACGGCTGTTGAGCAG-1 | 3 | A |
| 845 | GACGTGCGTGCGTCAC-1 | 2 | A |
| 846 | GACTAACCACCCGTGA-1 | 3 | A |
| 847 | GACTAACCAGACACAG-1 | 6 | B |
| 848 | GACTACACACAATCAC-1 | 10 | D |
| 849 | GACTGCGCAGTAGGGT-1 | 5 | A |
| 850 | GACTGCGGTGTCTGTA-1 | 3 | A |
| 851 | GACTGCGTCATATCTC-1 | 10 | D |
| 852 | GACTGCGTCGTAGCGC-1 | 2 | A |

| | | | |
|---|---|---|---|
| 853 | GAGCAGAAGAATCGTA-1 | 3 | A |
| 854 | GAGCAGAAGTTATCGC-1 | 3 | A |
| 855 | GAGCAGATCCGCCTAT-1 | 8 | D |
| 856 | GAGGTGAAGCCGAACA-1 | 5 | A |
| 857 | GAGGTGACACTAACTG-1 | 3 | A |
| 858 | GAGGTGACAGCTATGT-1 | 1 | A |
| 859 | GAGGTGACAGTGAATA-1 | 5 | A |
| 861 | GAGTCCGAGGGTTCGA-1 | 3 | A |
| 862 | GAGTCCGCATACGGTT-1 | 8 | D |
| 863 | GAGTCCGGTACTGCTA-1 | 1 | A |
| 864 | GAGTCCGGTCCTAGTA-1 | 8 | D |
| 865 | GAGTCCGTCAGACACT-1 | 1 | A |
| 866 | GATCAGTCAAGCACGA-1 | 3 | A |
| 867 | GATCAGTTCCACCCAT-1 | 1 | A |
| 868 | GATCAGTTCCGCTCTA-1 | 6 | B |
| 869 | GATCGATGTCCTACCT-1 | 9 | D |
| 870 | GATCGCGAGTCCATAC-1 | 8 | D |
| 871 | GATCGTAAGTAGGCCA-1 | 8 | D |
| 872 | GATCGTACAACCGGAA-1 | 7 | C |
| 873 | GATCGTACATACCAGT-1 | 9 | D |
| 875 | GATCTAGCAAGTGTTC-1 | 10 | D |
| 876 | GATCTAGTCACGCTCT-1 | 1 | A |
| 877 | GATGAAAAGTAATCCC-1 | 10 | D |
| 879 | GATGAAACATGCAGTT-1 | 5 | A |
| 880 | GATGAAAGTAACGGCA-1 | 6 | B |
| 881 | GATGAAAGTTCAGTAC-1 | 5 | A |
| 883 | GATGCTAGTAGGCTGA-1 | 3 | A |
| 884 | GATGCTAGTCTCTCGT-1 | 1 | A |
| 885 | GATGCTAGTTCCCTAC-1 | 10 | D |
| 886 | GATGCTAGTTGGAACG-1 | 6 | B |
| 887 | GATGCTATCACTATGG-1 | 9 | D |
| 888 | GATGCTATCATCAGCA-1 | 8 | D |
| 889 | GATTCAGAGATGTGGC-1 | 10 | D |
| 890 | GATTCAGGTTCAGTGT-1 | 10 | D |
| 891 | GATTCAGTCGACAACT-1 | 4 | A |
| 892 | GCAAACTAGCCACGTC-1 | 2 | A |
| 893 | GCAAACTCAAGCCAAG-1 | 11 | D |
| 894 | GCAAACTCACATTGGT-1 | 8 | D |
| 895 | GCAAACTTCCTACGAA-1 | 9 | D |
| 896 | GCAATCAAGGCAACAC-1 | 6 | B |
| 897 | GCAATCAGTTAGTGAA-1 | 10 | D |
| 898 | GCAATCAGTTGCCTCT-1 | 5 | A |

| | | | |
|---|---|---|---|
| 899 | GCAATCATCCCTGAGG-1 | 5 | A |
| 900 | GCAATCATCCGTTCAC-1 | 6 | B |
| 901 | GCACATAGTGAGCGAT-1 | 5 | A |
| 902 | GCACTCTAGACTTTCG-1 | 11 | D |
| 903 | GCACTCTAGCATCTTG-1 | 11 | D |
| 904 | GCACTCTAGTCCAGGA-1 | 1 | A |
| 905 | GCACTCTCAAGCATGG-1 | 4 | A |
| 906 | GCACTCTTCCACCTTG-1 | 8 | D |
| 908 | GCAGCCACATTGCATG-1 | 6 | B |
| 910 | GCAGCCATCGCTCTAC-1 | 5 | A |
| 911 | GCAGTTAAGGCCCTTG-1 | 11 | D |
| 912 | GCAGTTAGTAGCAGTG-1 | 8 | D |
| 913 | GCAGTTAGTGGAAGGG-1 | 7 | C |
| 914 | GCAGTTATCGCCCAGA-1 | 3 | A |
| 915 | GCATACAAGATCCTGT-1 | 10 | D |
| 916 | GCATACAAGGACAGAA-1 | 6 | B |
| 917 | GCATACACACGTGCGT-1 | 8 | D |
| 918 | GCATACATCAGATGTC-1 | 1 | A |
| 919 | GCATACATCCGCTCTA-1 | 8 | D |
| 920 | GCATGATCACGATAGG-1 | 3 | A |
| 921 | GCATGATCACTCACCT-1 | 10 | D |
| 922 | GCATGATCATGACCGC-1 | 8 | D |
| 923 | GCATGATGTCGCTCAG-1 | 3 | A |
| 924 | GCATGATGTTCGCGTG-1 | 1 | A |
| 925 | GCATGATTCCGTTAAG-1 | 1 | A |
| 926 | GCATGCGAGTAACCCT-1 | 1 | A |
| 927 | GCATGTATCCACAGCG-1 | 9 | D |
| 928 | GCCAAATAGTAAGTAC-1 | 3 | A |
| 930 | GCCAAATCATTTGTGG-1 | 1 | A |
| 931 | GCCAAATTCCGCGGAT-1 | 10 | D |
| 932 | GCCAAATTCTACAGTG-1 | 3 | A |
| 933 | GCCTCTAGTACGGTGA-1 | 8 | D |
| 934 | GCCTCTATCACGTAGT-1 | 3 | A |
| 935 | GCGACCACAACTCATG-1 | 2 | A |
| 936 | GCGACCAGTCCTACCT-1 | 6 | B |
| 937 | GCGAGAAAGCTTCAGT-1 | 2 | A |
| 938 | GCGCAGTAGCGATGAC-1 | 1 | A |
| 939 | GCGCAGTGTACTCGTA-1 | 1 | A |
| 940 | GCGCAGTGTGACACGA-1 | 8 | D |
| 941 | GCGCAGTGTGACCAAG-1 | 11 | D |
| 942 | GCGCCAAAGCGAGAAA-1 | 2 | A |
| 943 | GCGCCAAGTTGCCAAT-1 | 1 | A |

| | | | |
|---|---|---|---|
| 944 | GCGCCAATCCTCATCG-1 | 11 | D |
| 945 | GCGCGATAGGACTGGT-1 | 3 | A |
| 946 | GCGCGATAGGATATAC-1 | 8 | D |
| 948 | GCGGGTTAGGAGTTTA-1 | 10 | D |
| 949 | GCGGGTTAGTGTATCC-1 | 5 | A |
| 950 | GCGGGTTTCTATGCCC-1 | 3 | A |
| 951 | GCTCCTAAGATCCCGC-1 | 7 | C |
| 952 | GCTCCTAAGCCCGAAA-1 | 1 | A |
| 953 | GCTCCTATCCCAGAAT-1 | 2 | A |
| 954 | GCTCTGTAGCATGGCA-1 | 3 | A |
| 955 | GCTCTGTAGTTAGCGG-1 | 6 | B |
| 956 | GCTCTGTCAAACTCAC-1 | 6 | B |
| 957 | GCTCTGTTCACTCATT-1 | 6 | B |
| 958 | GCTGCAGAGGAGTTGC-1 | 10 | D |
| 959 | GCTGCAGGTAACGTGG-1 | 6 | B |
| 960 | GCTGCAGGTCGCGAAA-1 | 8 | D |
| 961 | GCTGCAGGTTGAACAA-1 | 6 | B |
| 962 | GCTGCGACACCTCCAT-1 | 10 | D |
| 963 | GCTGCGATCACTACAG-1 | 9 | D |
| 964 | GCTGCTTCACGATCTA-1 | 8 | D |
| 965 | GCTGCTTGTCATCTAG-1 | 8 | D |
| 966 | GCTGCTTTCAGTACAC-1 | 10 | D |
| 967 | GCTGGGTAGGCATATA-1 | 8 | D |
| 968 | GCTGGGTCAACTCCGG-1 | 3 | A |
| 969 | GCTGGGTGTGTAGTTC-1 | 5 | A |
| 970 | GCTGGGTGTGTCGCTG-1 | 4 | A |
| 971 | GCTGGGTTCTATGGTG-1 | 1 | A |
| 972 | GCTTCCAAGGGTTTCT-1 | 8 | D |
| 973 | GCTTCCATCCTACGAA-1 | 3 | A |
| 974 | GCTTCCATCTGGCCTT-1 | 8 | D |
| 975 | GCTTCCATCTTCCGAC-1 | 8 | D |
| 976 | GCTTGAACAGTGCTTA-1 | 1 | A |
| 978 | GGAAAGCAGCCAGAAC-1 | 5 | A |
| 979 | GGAAAGCCATGACTCA-1 | 8 | D |
| 980 | GGAAAGCTCCACCTTG-1 | 2 | A |
| 981 | GGAAAGCTCGTGTACA-1 | 11 | D |
| 982 | GGAACTTAGGGTCTCC-1 | 10 | D |
| 984 | GGAACTTGTCGGGAAT-1 | 6 | B |
| 985 | GGAACTTGTTTCCATT-1 | 5 | A |
| 986 | GGAACTTTCCGATACA-1 | 3 | A |
| 988 | GGACAAGAGGGATTTA-1 | 1 | A |
| 989 | GGACAAGGTGTAAAGT-1 | 2 | A |

| | | | |
|---|---|---|---|
| 990 | GGACAAGGTTCAGTAC-1 | 6 | B |
| 991 | GGACAAGTCGACGAAG-1 | 1 | A |
| 992 | GGACAAGTCGCGCACA-1 | 3 | A |
| 993 | GGACAGAAGGTATCGG-1 | 3 | A |
| 994 | GGACAGACAAACCCTA-1 | 10 | D |
| 995 | GGACAGACAGCCAGTC-1 | 1 | A |
| 996 | GGACATTAGTCAAGCG-1 | 6 | B |
| 997 | GGACGTCAGAACTTCC-1 | 1 | A |
| 998 | GGACGTCCACACGTGC-1 | 10 | D |
| 999 | GGAGCAAAGGTGCTTT-1 | 5 | A |
| 1001 | GGATGTTAGCCAGGAT-1 | 5 | A |
| 1002 | GGATGTTAGTGTACCT-1 | 3 | A |
| 1003 | GGATTACAGCCACCTG-1 | 1 | A |
| 1004 | GGATTACGTACTGTTG-1 | 8 | D |
| 1005 | GGATTACTCCAACAAC-1 | 8 | D |
| 1006 | GGATTACTCTCGGTTC-1 | 2 | A |
| 1007 | GGCAATTGTACAACGG-1 | 2 | A |
| 1008 | GGCAATTGTGCAACCC-1 | 1 | A |
| 1009 | GGCAATTTCTTGGATG-1 | 10 | D |
| 1010 | GGCCGATTCTGACGCG-1 | 2 | A |
| 1011 | GGCGACTGTTACTGAC-1 | 1 | A |
| 1012 | GGCGACTGTTCCGTCT-1 | 6 | B |
| 1013 | GGCGACTGTTGGAACG-1 | 8 | D |
| 1014 | GGCGACTTCCTTATTG-1 | 9 | D |
| 1016 | GGCTCGACAATGGCTT-1 | 5 | A |
| 1017 | GGCTCGACATACTGCA-1 | 10 | D |
| 1019 | GGCTGGTCAACCTGCG-1 | 7 | C |
| 1020 | GGCTGGTTCCTACGAA-1 | 1 | A |
| 1021 | GGCTGGTTCTGTGGCG-1 | 4 | A |
| 1022 | GGGAATGCAGATTGTC-1 | 8 | D |
| 1023 | GGGAATGCATCATGGT-1 | 8 | D |
| 1024 | GGGAATGGTGGTACAG-1 | 6 | B |
| 1025 | GGGACCTCATCCATCC-1 | 11 | D |
| 1026 | GGGACCTGTCCGGGTT-1 | 3 | A |
| 1027 | GGGACCTTCATCGACA-1 | 6 | B |
| 1028 | GGGAGATAGGCTAGCA-1 | 3 | A |
| 1029 | GGGAGATCACAGACCC-1 | 8 | D |
| 1030 | GGGAGATCAGCGCTTG-1 | 11 | D |
| 1031 | GGGAGATCATGCAGTT-1 | 3 | A |
| 1032 | GGGCACTGTGCAACCC-1 | 10 | D |
| 1033 | GGGCACTGTTCAAGGG-1 | 9 | D |
| 1034 | GGGCATCCACGCTGAC-1 | 1 | A |

| | | | |
|---|---|---|---|
| 1035 | GGGCATCGTCTCTATT-1 | 2 | A |
| 1036 | GGGCATCTCGAACCCG-1 | 3 | A |
| 1037 | GGGTCTGCATCGGTCG-1 | 1 | A |
| 1038 | GGGTCTGTCCACGATA-1 | 11 | D |
| 1039 | GGGTTGCGTGTCGCTG-1 | 3 | A |
| 1040 | GGTATTGAGGGTTCGA-1 | 6 | B |
| 1041 | GGTATTGAGGTTCCTA-1 | 8 | D |
| 1042 | GGTGAAGGTATTCGTG-1 | 2 | A |
| 1043 | GGTGCGTCACTAACTG-1 | 6 | B |
| 1044 | GGTGCGTGTGTTTCAG-1 | 1 | A |
| 1045 | GGTGTTAGTCAGTGGA-1 | 6 | B |
| 1046 | GTAACGTAGGCAGTCA-1 | 5 | A |
| 1048 | GTAACGTTCCCACAAA-1 | 8 | D |
| 1049 | GTAACTGAGCCGTCGT-1 | 2 | A |
| 1050 | GTAACTGCATAGGGCA-1 | 6 | B |
| 1051 | GTAACTGGTAGATGCG-1 | 11 | D |
| 1052 | GTAACTGGTTGAACTC-1 | 8 | D |
| 1053 | GTAACTGTCGAATGTC-1 | 11 | D |
| 1054 | GTACGTACACTGGCCA-1 | 3 | A |
| 1055 | GTACGTAGTGCGCTTG-1 | 2 | A |
| 1056 | GTACTCCGTCAGTTTG-1 | 3 | A |
| 1057 | GTACTCCGTTGGTATC-1 | 6 | B |
| 1058 | GTACTTTAGAGTAATC-1 | 6 | B |
| 1059 | GTACTTTCAGGCCGTT-1 | 10 | D |
| 1060 | GTACTTTTCAGTACAC-1 | 11 | D |
| 1061 | GTAGGCCCACGTGGCT-1 | 5 | A |
| 1062 | GTAGGCCGTGTAATGA-1 | 3 | A |
| 1063 | GTAGGCCTCCTTGCAC-1 | 10 | D |
| 1064 | GTAGGCCTCGTGCTAA-1 | 2 | A |
| 1065 | GTAGGCCTCTCGAGCG-1 | 6 | B |
| 1066 | GTAGTCAGTCATTACG-1 | 1 | A |
| 1067 | GTAGTCATCGAACCCG-1 | 2 | A |
| 1068 | GTATCTTAGGGAGTAA-1 | 3 | A |
| 1069 | GTATCTTAGGGATTTA-1 | 10 | D |
| 1070 | GTATCTTCATTTGTGG-1 | 6 | B |
| 1071 | GTATCTTGTCGCTTCT-1 | 11 | D |
| 1072 | GTATCTTGTCTCGGTG-1 | 2 | A |
| 1073 | GTATTCTCATAGGTCT-1 | 10 | D |
| 1074 | GTATTCTCATGCAGTT-1 | 4 | A |
| 1075 | GTCAAGTAGGACCAGT-1 | 6 | B |
| 1076 | GTCACAAAGTGAACAT-1 | 8 | D |
| 1077 | GTCACAAGTCAGTGGA-1 | 11 | D |

| | | | |
|---|---|---|---|
| 1078 | GTCACGGGTATTGAAG-1 | 8 | D |
| 1079 | GTCACGGTCCACAGAT-1 | 1 | A |
| 1080 | GTCATTTAGCAAAGCC-1 | 8 | D |
| 1081 | GTCATTTGTCGCTTCT-1 | 11 | D |
| 1082 | GTCATTTTCCCTTATA-1 | 2 | A |
| 1083 | GTCCTCAGTTTCCTGC-1 | 5 | A |
| 1084 | GTCCTCATCTAAGACC-1 | 11 | D |
| 1085 | GTCGGGTCACAGCTGC-1 | 9 | D |
| 1086 | GTCTCGTCAGGTTAAA-1 | 2 | A |
| 1087 | GTCTCGTCATTCCGAA-1 | 8 | D |
| 1088 | GTCTCGTGTACGTGCC-1 | 8 | D |
| 1089 | GTCTCGTTCACACGGC-1 | 2 | A |
| 1090 | GTCTCGTTCATTGCTT-1 | 8 | D |
| 1091 | GTCTTCGAGGTGCTAG-1 | 1 | A |
| 1092 | GTCTTCGAGTGTTGAA-1 | 11 | D |
| 1093 | GTCTTCGTCTGGTAGT-1 | 8 | D |
| 1094 | GTGAAGGGTCCTCGGA-1 | 3 | A |
| 1095 | GTGAAGGGTCTCGTCT-1 | 10 | D |
| 1096 | GTGCAGCAGCTGAAAT-1 | 3 | A |
| 1097 | GTGCAGCCAAAGTTAG-1 | 9 | D |
| 1098 | GTGCAGCGTCCTACCT-1 | 11 | D |
| 1099 | GTGCATACACTCTTAT-1 | 2 | A |
| 1100 | GTGCGGTCAATCGTCA-1 | 4 | A |
| 1101 | GTGCGGTCAGATCGAG-1 | 9 | D |
| 1102 | GTGCGGTGTACTCACA-1 | 3 | A |
| 1104 | GTGGGTCCACAGAGCA-1 | 3 | A |
| 1105 | GTGGGTCGTACCACTA-1 | 2 | A |
| 1106 | GTGGGTCGTTGGAACG-1 | 3 | A |
| 1107 | GTGGGTCTCCGCTTGT-1 | 11 | D |
| 1108 | GTGGGTCTCCTAGACA-1 | 8 | D |
| 1109 | GTGTGCGAGAAGGTTT-1 | 1 | A |
| 1110 | GTGTGCGAGCGCGATA-1 | 4 | A |
| 1111 | GTGTGCGAGGCTCTCG-1 | 11 | D |
| 1112 | GTGTGCGGTCGCTTCT-1 | 4 | A |
| 1113 | GTTAAGCAGTGTTTGC-1 | 3 | A |
| 1114 | GTTAAGCCAGATATCC-1 | 8 | D |
| 1115 | GTTAAGCTCCACTTAT-1 | 9 | D |
| 1116 | GTTAAGCTCTGTCGAA-1 | 11 | D |
| 1117 | GTTACAGAGTACACCT-1 | 1 | A |
| 1118 | GTTACAGTCAGGTGAG-1 | 4 | A |
| 1120 | GTTCATTTCGAGTATC-1 | 5 | A |
| 1121 | GTTCATTTCGTAGGGA-1 | 1 | A |

| | | | |
|---|---|---|---|
| 1122 | GTTCATTTCTCTTCCT-1 | 5 | A |
| 1123 | GTTCGGGTCAACAGTC-1 | 11 | D |
| 1124 | GTTCTCGCAACCCACG-1 | 5 | A |
| 1125 | GTTCTCGCACGCACCA-1 | 3 | A |
| 1126 | GTTCTCGCAGTGCTCG-1 | 1 | A |
| 1127 | GTTCTCGGTCATGTTG-1 | 11 | D |
| 1128 | GTTCTCGTCATGACCA-1 | 8 | D |
| 1129 | GTTTCTAAGGTGATTA-1 | 11 | D |
| 1130 | GTTTCTAAGTGGGATC-1 | 8 | D |
| 1131 | GTTTCTAGTCCGGTGT-1 | 6 | B |
| 1132 | GTTTCTATCAGATTCG-1 | 9 | D |
| 1133 | TAAACCGAGGCTAGAC-1 | 6 | B |
| 1134 | TAAACCGAGGTAGCCA-1 | 5 | A |
| 1135 | TAAACCGTCCAAGGGA-1 | 6 | B |
| 1136 | TAAGAGACAGCGTCAC-1 | 10 | D |
| 1137 | TAAGAGACATCTCGTC-1 | 9 | D |
| 1138 | TAAGAGAGTGTGTTCA-1 | 3 | A |
| 1139 | TAAGAGATCCAGGTTA-1 | 3 | A |
| 1140 | TAAGCGTTCATTGGAC-1 | 8 | D |
| 1141 | TAAGTGCAGGGATTCG-1 | 8 | D |
| 1142 | TAAGTGCAGGTAAGTT-1 | 8 | D |
| 1143 | TAAGTGCTCGGGTCAC-1 | 10 | D |
| 1144 | TACACGACAAGCAAGC-1 | 10 | D |
| 1145 | TACACGACACTAGGCC-1 | 3 | A |
| 1146 | TACACGATCCGACCAG-1 | 3 | A |
| 1147 | TACACGATCCGTTCAC-1 | 5 | A |
| 1148 | TACAGTGAGTCCCGAC-1 | 3 | A |
| 1149 | TACAGTGGTCAATTCG-1 | 9 | D |
| 1150 | TACAGTGGTCTAACGT-1 | 1 | A |
| 1151 | TACCTATAGTAGGTGC-1 | 10 | D |
| 1152 | TACCTATGTAAACAGT-1 | 4 | A |
| 1153 | TACCTATGTCAGTTTG-1 | 3 | A |
| 1154 | TACCTATGTTGAACAA-1 | 3 | A |
| 1155 | TACCTTAAGACTGGCA-1 | 6 | B |
| 1156 | TACCTTAAGGAGCGAG-1 | 1 | A |
| 1157 | TACCTTAAGTGAATTG-1 | 1 | A |
| 1158 | TACCTTACATTCCGAA-1 | 1 | A |
| 1159 | TACCTTATCACTCATT-1 | 1 | A |
| 1160 | TACGGATAGCTGAACG-1 | 5 | A |
| 1161 | TACGGATAGTGTTAGA-1 | 1 | A |
| 1162 | TACGGATCAACACCGC-1 | 6 | B |
| 1163 | TACGGATCAGAGGCAT-1 | 5 | A |

| | | | |
|---|---|---|---|
| 1165 | TACGGATGTGATGTTC-1 | 10 | D |
| 1166 | TACGGATTCAAGCCGC-1 | 8 | D |
| 1167 | TACGGATTCACACGGC-1 | 6 | B |
| 1168 | TACGGGCAGGTAAGTT-1 | 11 | D |
| 1169 | TACGGGCGTATCTCTT-1 | 5 | A |
| 1170 | TACGGTAAGAACAATC-1 | 10 | D |
| 1171 | TACGGTAAGTTTCCTT-1 | 3 | A |
| 1172 | TACGGTAGTGCTTTCC-1 | 3 | A |
| 1173 | TACGGTATCTCGTGGG-1 | 5 | A |
| 1174 | TACTCATAGAACAATC-1 | 10 | D |
| 1176 | TACTCATAGGCATGGT-1 | 4 | A |
| 1177 | TACTCATCAACGGCTC-1 | 6 | B |
| 1178 | TACTCGCAGAAGGTGA-1 | 3 | A |
| 1180 | TACTCGCCATGATGCT-1 | 5 | A |
| 1181 | TACTCGCTCCACGATA-1 | 6 | B |
| 1182 | TACTTGTAGTGGGTAC-1 | 10 | D |
| 1183 | TACTTGTCAGACACAG-1 | 8 | D |
| 1185 | TACTTGTGTTCGGTGC-1 | 10 | D |
| 1186 | TACTTGTTCACCGGTG-1 | 10 | D |
| 1188 | TAGACCAAGTCGATAA-1 | 9 | D |
| 1189 | TAGACCACATCCACGG-1 | 1 | A |
| 1190 | TAGACCACATGATGCT-1 | 10 | D |
| 1191 | TAGACCATCATCAGCA-1 | 7 | C |
| 1192 | TAGAGCTAGCTTTCAG-1 | 8 | D |
| 1193 | TAGAGCTAGGCTAGGT-1 | 8 | D |
| 1195 | TAGAGCTCATTGGTCA-1 | 3 | A |
| 1196 | TAGAGCTTCTCAGATG-1 | 3 | A |
| 1197 | TAGCCGGAGTCGTGTT-1 | 8 | D |
| 1198 | TAGCCGGCAACCTCAA-1 | 8 | D |
| 1199 | TAGCCGGCACCACACG-1 | 11 | D |
| 1200 | TAGCCGGCACGCTGAC-1 | 10 | D |
| 1201 | TAGCCGGGTTCCACAA-1 | 11 | D |
| 1202 | TAGCCGGTCAGCGGCT-1 | 8 | D |
| 1203 | TAGGCATAGCTAAACA-1 | 4 | A |
| 1204 | TAGGCATAGTAGGTGC-1 | 6 | B |
| 1205 | TAGGCATAGTGAACAT-1 | 8 | D |
| 1206 | TAGGCATCAATCGTCA-1 | 9 | D |
| 1207 | TAGGCATGTCGGCATC-1 | 3 | A |
| 1208 | TAGGCATGTTCAGTAC-1 | 3 | A |
| 1209 | TAGTGGTCAAGTGTTC-1 | 10 | D |
| 1210 | TAGTGGTCACACACGC-1 | 6 | B |
| 1211 | TAGTGGTGTGTGACAG-1 | 1 | A |

| | | | |
|---|---|---|---|
| 1213 | TAGTTGGCATGAGAAT-1 | 5 | A |
| 1214 | TAGTTGGCATGGCACC-1 | 1 | A |
| 1215 | TAGTTGGTCCTGATTT-1 | 5 | A |
| 1216 | TAGTTGGTCGAACCCG-1 | 9 | D |
| 1217 | TATCAGGAGTCCGTAT-1 | 11 | D |
| 1219 | TATCAGGGTTCGAGCC-1 | 7 | C |
| 1220 | TATCAGGGTTCGTCAA-1 | 5 | A |
| 1221 | TATCAGGTCTGGTAGT-1 | 3 | A |
| 1222 | TATCAGGTCTTCTTGA-1 | 11 | D |
| 1223 | TATCTCAAGACTGGGT-1 | 5 | A |
| 1224 | TATCTCAGTAGTGGCA-1 | 3 | A |
| 1225 | TATCTCAGTCCGAAGA-1 | 9 | D |
| 1226 | TATCTCAGTGACCCGT-1 | 8 | D |
| 1227 | TATGCCCAGGGCTTGA-1 | 11 | D |
| 1228 | TATGCCCTCTAGAACC-1 | 3 | A |
| 1229 | TATTACCAGCCAGGAT-1 | 4 | A |
| 1230 | TATTACCGTGCTATTG-1 | 4 | A |
| 1231 | TCAACGAAGAGCAATT-1 | 3 | A |
| 1233 | TCAACGAAGGCTGCTC-1 | 11 | D |
| 1234 | TCAACGATCTTTCGCG-1 | 6 | B |
| 1235 | TCACAAGCATTTCCAT-1 | 8 | D |
| 1236 | TCACGAAGTCAAACTC-1 | 3 | A |
| 1237 | TCACGAAGTGTCTAAC-1 | 4 | A |
| 1238 | TCAGATGAGAGCCCAA-1 | 8 | D |
| 1239 | TCAGATGTCCGATACA-1 | 5 | A |
| 1240 | TCAGCAAGTAGATGCG-1 | 11 | D |
| 1241 | TCAGCAAGTGTTTCAG-1 | 3 | A |
| 1242 | TCAGCAAGTTGGTGAG-1 | 3 | A |
| 1243 | TCAGCTCAGTTGTCGT-1 | 5 | A |
| 1245 | TCAGGATTCGGGTCAC-1 | 6 | B |
| 1246 | TCAGGTACATAGACAA-1 | 3 | A |
| 1247 | TCAGGTACATGTAGCT-1 | 10 | D |
| 1248 | TCAGGTATCAAGTGCT-1 | 8 | D |
| 1249 | TCATTACAGCCAGTTT-1 | 1 | A |
| 1250 | TCATTACCACAGACCC-1 | 3 | A |
| 1251 | TCATTACCATTCAGTG-1 | 1 | A |
| 1252 | TCATTACTCTCCACAC-1 | 8 | D |
| 1253 | TCATTTGAGTAGCCGA-1 | 8 | D |
| 1254 | TCATTTGAGTCTCGTA-1 | 9 | D |
| 1255 | TCCACACCAAGCAAGC-1 | 6 | B |
| 1256 | TCCACACGTGCTCGAC-1 | 4 | A |
| 1257 | TCCACACTCGCAACGC-1 | 6 | B |

| | | | |
|---|---|---|---|
| 1258 | TCCCGATAGAATCGTA-1 | 11 | D |
| 1259 | TCCCGATAGTGCGATG-1 | 9 | D |
| 1260 | TCCCGATGTTAAGTAG-1 | 11 | D |
| 1261 | TCGAGGCAGATCCTGT-1 | 3 | A |
| 1262 | TCGAGGCCACCGAACC-1 | 6 | B |
| 1263 | TCGAGGCCAGTGAAAT-1 | 3 | A |
| 1265 | TCGCGAGAGAGTTGGC-1 | 1 | A |
| 1266 | TCGCGAGGTGATCCTA-1 | 1 | A |
| 1267 | TCGCGAGGTTCAAGAA-1 | 1 | A |
| 1268 | TCGCGAGTCCGCTTGT-1 | 3 | A |
| 1269 | TCGCGAGTCTACGGCG-1 | 2 | A |
| 1270 | TCGCGTTAGACTGGGT-1 | 3 | A |
| 1271 | TCGCGTTAGGACGAAA-1 | 7 | C |
| 1272 | TCGCGTTCAACAACTC-1 | 3 | A |
| 1273 | TCGCGTTGTCGCGAAA-1 | 8 | D |
| 1274 | TCGCGTTGTCTAGCGC-1 | 2 | A |
| 1276 | TCGGGACAGTCAAGGC-1 | 8 | D |
| 1277 | TCGGGACGTTTCCATT-1 | 11 | D |
| 1278 | TCGGGACTCGTAAGGG-1 | 2 | A |
| 1279 | TCGGTAAGTATACACC-1 | 6 | B |
| 1280 | TCGGTAATCAGAAACA-1 | 7 | C |
| 1281 | TCGTACCCAATCCATG-1 | 9 | D |
| 1282 | TCGTACCCACGTTTAG-1 | 2 | A |
| 1283 | TCGTACCGTGTGACAG-1 | 11 | D |
| 1284 | TCGTAGACAATCCGTA-1 | 4 | A |
| 1285 | TCGTAGAGTGACTGAG-1 | 1 | A |
| 1286 | TCTATTGCAACGAAAT-1 | 11 | D |
| 1287 | TCTATTGCAGAGATCG-1 | 9 | D |
| 1288 | TCTATTGCATGCGTGC-1 | 8 | D |
| 1289 | TCTATTGGTATTCGCA-1 | 1 | A |
| 1290 | TCTATTGGTTGGCGTC-1 | 2 | A |
| 1291 | TCTATTGTCAGCCCGA-1 | 11 | D |
| 1292 | TCTATTGTCATCGACA-1 | 1 | A |
| 1293 | TCTCATAAGCAGCCCT-1 | 10 | D |
| 1294 | TCTCATAAGGGATCTG-1 | 8 | D |
| 1295 | TCTCATAGTGCACACC-1 | 10 | D |
| 1296 | TCTCATATCTAAGACC-1 | 5 | A |
| 1297 | TCTCATATCTGTTGTT-1 | 1 | A |
| 1298 | TCTCTAAAGTAAGTAC-1 | 11 | D |
| 1299 | TCTCTAAGTAGGTTTC-1 | 4 | A |
| 1300 | TCTGAGAAGATCTGCT-1 | 6 | B |
| 1301 | TCTGGAACACGTGGCT-1 | 3 | A |

| | | | |
|---|---|---|---|
| 1302 | TCTGGAATCAGCGGCT-1 | 11 | D |
| 1303 | TCTTCGGAGAGGGCTT-1 | 3 | A |
| 1304 | TCTTCGGAGCCAGGAT-1 | 1 | A |
| 1305 | TCTTCGGAGCTATGCT-1 | 10 | D |
| 1306 | TCTTCGGGTCGGATGA-1 | 3 | A |
| 1307 | TCTTTCCCACAGCTAT-1 | 11 | D |
| 1308 | TCTTTCCCACTAGGCC-1 | 5 | A |
| 1309 | TCTTTCCGTGGTGAGT-1 | 4 | A |
| 1311 | TGAAAGAAGGTACTCT-1 | 10 | D |
| 1313 | TGACAACAGCGAACTG-1 | 1 | A |
| 1314 | TGACAACAGCTCCTTC-1 | 10 | D |
| 1315 | TGACGGCAGTTACGGG-1 | 8 | D |
| 1317 | TGACGGCGTCCGATGC-1 | 1 | A |
| 1318 | TGACGGCGTGTCAACT-1 | 8 | D |
| 1319 | TGACTAGGTAGGGTGT-1 | 11 | D |
| 1320 | TGACTAGTCCTCGCTA-1 | 5 | A |
| 1321 | TGACTAGTCGGGCTTG-1 | 1 | A |
| 1323 | TGACTTTAGATTCACC-1 | 1 | A |
| 1324 | TGACTTTAGCAACGGT-1 | 4 | A |
| 1325 | TGACTTTAGGTGTGCA-1 | 6 | B |
| 1326 | TGACTTTAGTACAGTA-1 | 8 | D |
| 1327 | TGAGAGGAGTGTCCAT-1 | 10 | D |
| 1328 | TGAGAGGGTCCCACAG-1 | 2 | A |
| 1329 | TGAGAGGGTCGCGTAC-1 | 3 | A |
| 1330 | TGAGAGGTCTACGGCG-1 | 8 | D |
| 1331 | TGAGCATCATAGGTCT-1 | 9 | D |
| 1332 | TGAGCATCATCGGGCT-1 | 10 | D |
| 1333 | TGAGCATTCTGTATGG-1 | 3 | A |
| 1334 | TGAGCCGAGGCATGGT-1 | 10 | D |
| 1335 | TGAGCCGGTGCTGTAT-1 | 4 | A |
| 1336 | TGAGCCGGTTCCCTAC-1 | 10 | D |
| 1337 | TGAGGGACACCAATGT-1 | 4 | A |
| 1338 | TGATTTCTCACGATAC-1 | 5 | A |
| 1339 | TGATTTCTCCGACCAG-1 | 1 | A |
| 1340 | TGCACCTAGTACGCGA-1 | 6 | B |
| 1341 | TGCACCTTCAGCCCGA-1 | 7 | C |
| 1342 | TGCCAAAAGACAGGCT-1 | 10 | D |
| 1343 | TGCCAAACAAGTGTTC-1 | 3 | A |
| 1344 | TGCCAAACACTACATG-1 | 2 | A |
| 1345 | TGCCCATAGGATGCGT-1 | 1 | A |
| 1346 | TGCCCATAGGCCATAG-1 | 11 | D |
| 1347 | TGCCCATGTACCAGCC-1 | 11 | D |

| | | | |
|---|---|---|---|
| 1348 | TGCCCATTCCAACAAC-1 | 11 | D |
| 1349 | TGCCCTAAGCCAAACG-1 | 7 | C |
| 1351 | TGCCCTAGTCGTAATC-1 | 3 | A |
| 1352 | TGCCCTAGTGTAGTTC-1 | 4 | A |
| 1353 | TGCCCTAGTTCTAGAC-1 | 6 | B |
| 1354 | TGCGCAGAGAGGTACC-1 | 6 | B |
| 1356 | TGCGGGTCAGACAGTG-1 | 10 | D |
| 1357 | TGCGGGTCATACGTCA-1 | 8 | D |
| 1358 | TGCGGGTCATCTGTAG-1 | 6 | B |
| 1359 | TGCGGGTTCTATGGTG-1 | 9 | D |
| 1360 | TGCGTGGAGTAGTGCG-1 | 3 | A |
| 1361 | TGCTACCGTACGTCAT-1 | 7 | C |
| 1363 | TGCTGCTTCGGAGAGT-1 | 8 | D |
| 1364 | TGGACGCAGACGCTTT-1 | 8 | D |
| 1365 | TGGACGCCATCGGCGT-1 | 10 | D |
| 1367 | TGGACGCTCGAACCCG-1 | 8 | D |
| 1368 | TGGCCAGGTCCTAGTA-1 | 7 | C |
| 1369 | TGGCCAGTCAAGCCAT-1 | 5 | A |
| 1370 | TGGCGCAGTCGACCAC-1 | 5 | A |
| 1371 | TGGCGCAGTGTCCGAC-1 | 11 | D |
| 1372 | TGGCGCATCCAACCGG-1 | 5 | A |
| 1373 | TGGCGCATCCGGTTCT-1 | 10 | D |
| 1374 | TGGCTGGAGTAAGTAC-1 | 10 | D |
| 1375 | TGGCTGGCAACGCGCT-1 | 4 | A |
| 1377 | TGGGAAGAGGAATGGA-1 | 11 | D |
| 1378 | TGGGAAGCAAGGTGAC-1 | 6 | B |
| 1379 | TGGGAAGGTGGTAACG-1 | 7 | C |
| 1380 | TGGGAAGTCACTTTAC-1 | 3 | A |
| 1381 | TGGGCGTAGGCTCAGA-1 | 3 | A |
| 1382 | TGGGCGTGTTTCCTGC-1 | 3 | A |
| 1383 | TGGGCGTTCACTGTTT-1 | 9 | D |
| 1384 | TGGTTAGAGGGATGGG-1 | 3 | A |
| 1385 | TGGTTAGAGTTAGCGG-1 | 3 | A |
| 1386 | TGGTTAGTCTTATGGG-1 | 3 | A |
| 1387 | TGGTTCCAGACAAAGG-1 | 11 | D |
| 1388 | TGGTTCCAGAGTAAGG-1 | 9 | D |
| 1389 | TGGTTCCAGGTGCACA-1 | 6 | B |
| 1390 | TGGTTCCTCACACCTC-1 | 5 | A |
| 1391 | TGTATTCCAACATAAG-1 | 1 | A |
| 1392 | TGTATTCCAATCGCGC-1 | 1 | A |
| 1393 | TGTCCCACAGTCGGAA-1 | 3 | A |
| 1394 | TGTCCCAGTAGATAGT-1 | 1 | A |

| | | | |
|---|---|---|---|
| 1395 | TGTGGTACATTATCGG-1 | 6 | B |
| 1396 | TGTGGTACATTGAAGA-1 | 3 | A |
| 1397 | TGTGGTAGTTTGGGCC-1 | 7 | C |
| 1398 | TGTGGTATCAAGCCGC-1 | 11 | D |
| 1400 | TGTTCCGCATAGAGCG-1 | 2 | A |
| 1401 | TTAACTCCAGCTCATA-1 | 9 | D |
| 1402 | TTAACTCCATGGCTAT-1 | 7 | C |
| 1403 | TTAACTCTCAGTACAC-1 | 11 | D |
| 1404 | TTAACTCTCGATAGTC-1 | 9 | D |
| 1405 | TTAGGACAGAGCTGCA-1 | 1 | A |
| 1406 | TTAGGACAGTTACCCA-1 | 3 | A |
| 1407 | TTAGGACTCTGAAGTC-1 | 11 | D |
| 1408 | TTAGGCAAGGTAAGTT-1 | 11 | D |
| 1409 | TTAGGCACACCAATGT-1 | 5 | A |
| 1410 | TTAGGCAGTAACGACG-1 | 7 | C |
| 1411 | TTAGTTCAGAGCCACA-1 | 1 | A |
| 1412 | TTATGCTAGTGCCATT-1 | 5 | A |
| 1413 | TTATGCTCAGAGACGT-1 | 4 | A |
| 1414 | TTCCCAGAGGAGCGTT-1 | 5 | A |
| 1415 | TTCCCAGCACGATCTA-1 | 8 | D |
| 1417 | TTCCCAGGTTCTCTCG-1 | 3 | A |
| 1418 | TTCCCAGTCCGCCTTA-1 | 9 | D |
| 1419 | TTCGAAGGTACTCGCG-1 | 9 | D |
| 1420 | TTCGGTCAGCGATGAC-1 | 5 | A |
| 1421 | TTCGGTCAGTGAATTG-1 | 1 | A |
| 1422 | TTCGGTCCACTTGAGT-1 | 3 | A |
| 1423 | TTCGGTCCATAGCACT-1 | 11 | D |
| 1424 | TTCTACAAGAGGTAGA-1 | 8 | D |
| 1425 | TTCTACAAGCAGAGAA-1 | 11 | D |
| 1426 | TTCTACAAGTGACTCT-1 | 3 | A |
| 1427 | TTCTACAGTCGCTTCT-1 | 4 | A |
| 1428 | TTCTACAGTCTAACGT-1 | 1 | A |
| 1429 | TTCTACATCTTAGCTT-1 | 8 | D |
| 1430 | TTCTCAAAGGCACATG-1 | 4 | A |
| 1432 | TTCTCAAGTAGGGTGT-1 | 10 | D |
| 1433 | TTCTCAAGTGGCCACT-1 | 1 | A |
| 1434 | TTCTCCTAGACAGACC-1 | 1 | A |
| 1435 | TTCTCCTAGCAGCCCT-1 | 11 | D |
| 1436 | TTCTCCTCATTCGTTT-1 | 1 | A |
| 1437 | TTCTCCTTCATCGCGG-1 | 6 | B |
| 1438 | TTCTTAGAGTACTTGC-1 | 3 | A |
| 1439 | TTGAACGAGAATTGTG-1 | 4 | A |

| | | | |
|---|---|---|---|
| 1440 | TTGAACGAGCACGCCT-1 | 3 | A |
| 1441 | TTGAACGAGTGTCCAT-1 | 10 | D |
| 1442 | TTGACTTAGGGCTTCC-1 | 8 | D |
| 1443 | TTGACTTAGTCCGTAT-1 | 10 | D |
| 1444 | TTGCCGTAGAGTAATC-1 | 3 | A |
| 1445 | TTGCCGTAGTCGTGTT-1 | 8 | D |
| 1446 | TTGCCGTGTAAGCGCA-1 | 3 | A |
| 1447 | TTGCCGTGTGCACTTA-1 | 3 | A |
| 1448 | TTGCGTCAGGATGTAT-1 | 4 | A |
| 1449 | TTGCGTCTCATTTGCT-1 | 5 | A |
| 1450 | TTGGAACGTAAGGTGC-1 | 8 | D |
| 1451 | TTGGCAATCATGGTGT-1 | 5 | A |
| 1452 | TTGTAGGAGAGTACAT-1 | 8 | D |
| 1453 | TTGTAGGTCTTCTTGA-1 | 2 | A |
| 1454 | TTTACTGAGAAACGCC-1 | 6 | B |
| 1455 | TTTACTGAGACAGACC-1 | 10 | D |
| 1456 | TTTACTGAGCCCAATT-1 | 2 | A |
| 1458 | TTTACTGGTAACAATG-1 | 4 | A |
| 1459 | TTTACTGTCCACGATA-1 | 3 | A |
| 1460 | TTTATGCAGTAGTATG-1 | 2 | A |
| 1461 | TTTATGCTCTGAAGGG-1 | 1 | A |
| 1462 | TTTCCTCAGCTCCTTC-1 | 10 | D |
| 1463 | TTTCCTCCAACAGCCC-1 | 1 | A |
| 1466 | TTTGCGCAGTGCCATT-1 | 11 | D |
| 1467 | TTTGCGCCAACTCATG-1 | 6 | B |
| 1468 | TTTGCGCTCGATAAGA-1 | 9 | D |
| 1469 | TTTGCGCTCTCATCCG-1 | 1 | A |
| 1470 | TTTGGTTCAAGACGAC-1 | 11 | D |
| 1471 | TTTGGTTGTGATGTTC-1 | 10 | D |
| 1472 | TTTGGTTTCAGATTCG-1 | 11 | D |
| 1473 | TTTGTCAGTGTGTTCA-1 | 10 | D |
| 1474 | TTTGTCATCACTGCGG-1 | 6 | B |

**Supplementary Table 4 | CNV events by cluster.** Frequency of each of the 114 CNV events that passed filters across the 11 clusters and 4 groups. The start and end headers are the median values of the edges shared across all single cells for that event.

| chrom | start | end | ploidy | event | Group A | Group B | Group C | Group D | Cluster 1 | Cluster 2 | Cluster 3 | Cluster 4 | Cluster 5 | Cluster 6 | Cluster 7 | Cluster 8 | Cluster 9 | Cluster 10 | Cluster 11 |
|---|---|---|---|---|---|---|---|---|---|---|---|---|---|---|---|---|---|---|---|
| 1 | 17200001 | 120520000 | 2 | 13 | 0 | 0.726 | 0.698 | 0.002 | 0 | 0 | 0 | 0 | 0 | 0.726 | 0.698 | 0.006 | 0 | 0 | 0 |
| 1 | 87340001 | 120520000 | 2 | 16 | 0.832 | 0.017 | 0.023 | 0.812 | 0.839 | 0.92 | 0.805 | 0.813 | 0.817 | 0.017 | 0.023 | 0.878 | 0.826 | 0.692 | 0.833 |
| 1 | 800001 | 12920000 | 2 | 14 | 0.021 | 0.88 | 0.907 | 0.03 | 0.007 | 0.034 | 0.018 | 0.04 | 0.025 | 0.88 | 0.907 | 0.035 | 0.029 | 0.015 | 0.038 |
| 1 | 13780001 | 16840000 | 2 | 15 | 0.011 | 0.487 | 0.605 | 0.014 | 0 | 0 | 0.009 | 0.027 | 0.025 | 0.487 | 0.605 | 0.023 | 0.014 | 0.008 | 0.011 |
| 1 | 149860001 | 249220000 | 3 | 54 | 0.115 | 0.12 | 0.047 | 0.109 | 0.161 | 0.08 | 0.086 | 0.12 | 0.133 | 0.12 | 0.047 | 0.087 | 0.101 | 0.12 | 0.124 |
| 1 | 17200001 | 87320000 | 3 | 51 | 0.646 | 0 | 0 | 0.698 | 0.705 | 0.761 | 0.629 | 0.627 | 0.533 | 0 | 0 | 0.738 | 0.87 | 0.504 | 0.737 |
| 1 | 800001 | 12920000 | 3 | 53 | 0.806 | 0.017 | 0 | 0.798 | 0.812 | 0.807 | 0.846 | 0.693 | 0.792 | 0.017 | 0 | 0.837 | 0.797 | 0.737 | 0.806 |
| 1 | 13780001 | 16840000 | 3 | 58 | 0.577 | 0.017 | 0 | 0.571 | 0.57 | 0.216 | 0.588 | 0.573 | 0.833 | 0.017 | 0 | 0.483 | 0.246 | 0.812 | 0.602 |
| 1 | 149860001 | 249160000 | 4 | 83 | 0.547 | 0.564 | 0.535 | 0.521 | 0.537 | 0.716 | 0.593 | 0.56 | 0.342 | 0.564 | 0.535 | 0.651 | 0.594 | 0.301 | 0.532 |
| 2 | 95400001 | 243000000 | 3 | 69 | 0.637 | 0.675 | 0.605 | 0.621 | 0.591 | 0.784 | 0.67 | 0.693 | 0.492 | 0.675 | 0.605 | 0.669 | 0.696 | 0.511 | 0.629 |
| 2 | 20001 | 87060000 | 3 | 71 | 0.559 | 0.556 | 0.442 | 0.571 | 0.604 | 0.795 | 0.624 | 0.52 | 0.233 | 0.556 | 0.442 | 0.674 | 0.797 | 0.165 | 0.683 |
| 2 | 33160001 | 87060000 | 3 | 72 | 0.123 | 0.068 | 0.116 | 0.15 | 0.134 | 0.023 | 0.072 | 0.133 | 0.267 | 0.068 | 0.116 | 0.081 | 0.029 | 0.429 | 0.059 |
| 2 | 20001 | 33120000 | 3 | 73 | 0.124 | 0.12 | 0.209 | 0.125 | 0.121 | 0.023 | 0.081 | 0.133 | 0.275 | 0.12 | 0.209 | 0.058 | 0.043 | 0.368 | 0.043 |
| 3 | 60001 | 24560000 | 2 | 28 | 0.945 | 0.906 | 0.837 | 0.898 | 0.987 | 0.989 | 0.914 | 0.96 | 0.908 | 0.906 | 0.837 | 0.919 | 0.928 | 0.85 | 0.903 |
| 3 | 93520001 | 197860000 | 4 | 92 | 0.593 | 0.65 | 0.372 | 0.564 | 0.685 | 0.739 | 0.633 | 0.507 | 0.35 | 0.65 | 0.372 | 0.605 | 0.696 | 0.353 | 0.629 |
| 3 | 26660001 | 90500000 | 4 | 93 | 0.701 | 0.59 | 0.558 | 0.668 | 0.698 | 0.818 | 0.742 | 0.693 | 0.55 | 0.59 | 0.558 | 0.779 | 0.768 | 0.496 | 0.651 |
| 3 | 24560001 | 26660000 | 9 | 114 | 0.118 | 0.145 | 0.116 | 0.139 | 0.134 | 0.273 | 0.072 | 0.173 | 0.033 | 0.145 | 0.116 | 0.105 | 0.333 | 0.045 | 0.167 |
| 4 | 80001 | 49080000 | 2 | 29 | 0.801 | 0.769 | 0.791 | 0.786 | 0.866 | 0.898 | 0.814 | 0.787 | 0.633 | 0.769 | 0.791 | 0.849 | 0.928 | 0.624 | 0.79 |
| 4 | 52680001 | 190900000 | 4 | 94 | 0.478 | 0.504 | 0.442 | 0.455 | 0.523 | 0.705 | 0.493 | 0.533 | 0.192 | 0.504 | 0.442 | 0.465 | 0.609 | 0.203 | 0.57 |
| 4 | 70300001 | 190900000 | 4 | 96 | 0.044 | 0.051 | 0.093 | 0.068 | 0.034 | 0.023 | 0.05 | 0.027 | 0.075 | 0.051 | 0.093 | 0.076 | 0.087 | 0.053 | 0.065 |
| 4 | 52680001 | 69360000 | 4 | 95 | 0.184 | 0.205 | 0.279 | 0.198 | 0.174 | 0.102 | 0.163 | 0.16 | 0.308 | 0.205 | 0.279 | 0.198 | 0.13 | 0.323 | 0.134 |
| 5 | 70740001 | 180700000 | 2 | 30 | 0.677 | 0.786 | 0.651 | 0.711 | 0.752 | 0.852 | 0.719 | 0.6 | 0.425 | 0.786 | 0.651 | 0.82 | 0.855 | 0.564 | 0.661 |
| 5 | 20001 | 46400000 | 2 | 31 | 0.841 | 0.897 | 0.767 | 0.846 | 0.893 | 0.875 | 0.833 | 0.907 | 0.725 | 0.897 | 0.767 | 0.884 | 0.942 | 0.654 | 0.914 |
| 5 | 49440001 | 68820000 | 2 | 32 | 0.93 | 0.983 | 0.907 | 0.93 | 0.946 | 0.955 | 0.946 | 0.96 | 0.842 | 0.983 | 0.907 | 0.953 | 0.957 | 0.85 | 0.957 |
| 6 | 65300001 | 170920000 | 2 | 33 | 0.046 | 0.248 | 0.14 | 0.021 | 0 | 0.102 | 0.063 | 0.027 | 0.042 | 0.248 | 0.14 | 0.023 | 0.029 | 0.03 | 0.011 |
| 6 | 65300001 | 170920000 | 3 | 74 | 0.625 | 0.521 | 0.488 | 0.641 | 0.725 | 0.659 | 0.615 | 0.693 | 0.45 | 0.521 | 0.488 | 0.808 | 0.623 | 0.414 | 0.656 |
| 6 | 220001 | 58780000 | 3 | 75 | 0.083 | 0.085 | 0.093 | 0.055 | 0.04 | 0.125 | 0.081 | 0.04 | 0.133 | 0.085 | 0.093 | 0.07 | 0.058 | 0.068 | 0.032 |
| 6 | 200001 | 58760000 | 4 | 100 | 0.686 | 0.41 | 0.442 | 0.67 | 0.772 | 0.716 | 0.719 | 0.707 | 0.483 | 0.41 | 0.442 | 0.669 | 0.768 | 0.466 | 0.78 |
| 6 | 61880001 | 65320000 | 4 | 97 | 0.064 | 0.026 | 0.023 | 0.054 | 0.054 | 0.034 | 0.086 | 0.013 | 0.092 | 0.026 | 0.023 | 0.052 | 0.043 | 0.053 | 0.059 |
| 6 | 61880001 | 65300000 | 4 | 98 | 0.121 | 0.034 | 0.093 | 0.082 | 0.141 | 0.102 | 0.14 | 0.147 | 0.058 | 0.034 | 0.093 | 0.093 | 0.116 | 0.083 | 0.059 |
| 6 | 61880001 | 65280000 | 4 | 99 | 0.038 | 0.077 | 0.07 | 0.064 | 0.04 | 0.034 | 0.041 | 0.013 | 0.05 | 0.077 | 0.07 | 0.07 | 0.043 | 0.075 | 0.059 |
| 7 | 150740001 | 159120000 | 2 | 34 | 0.908 | 0.889 | 0.791 | 0.918 | 0.933 | 0.92 | 0.9 | 0.933 | 0.867 | 0.889 | 0.791 | 0.913 | 0.913 | 0.925 | 0.919 |
| 7 | 80001 | 58040000 | 4 | 101 | 0.807 | 0.778 | 0.698 | 0.814 | 0.826 | 0.898 | 0.819 | 0.813 | 0.692 | 0.778 | 0.698 | 0.89 | 0.841 | 0.692 | 0.823 |
| 7 | 76820001 | 125740000 | 4 | 102 | 0.752 | 0.709 | 0.698 | 0.746 | 0.772 | 0.841 | 0.724 | 0.773 | 0.7 | 0.709 | 0.698 | 0.744 | 0.783 | 0.707 | 0.763 |
| 7 | 63260001 | 74140000 | 4 | 103 | 0.939 | 0.923 | 0.884 | 0.912 | 0.966 | 0.966 | 0.946 | 0.907 | 0.892 | 0.923 | 0.884 | 0.936 | 0.971 | 0.872 | 0.898 |
| 7 | 143800001 | 150740000 | 4 | 104 | 0.299 | 0.35 | 0.209 | 0.304 | 0.322 | 0.284 | 0.262 | 0.453 | 0.25 | 0.35 | 0.209 | 0.25 | 0.362 | 0.278 | 0.349 |
| 7 | 144100001 | 150740000 | 4 | 105 | 0.349 | 0.265 | 0.349 | 0.316 | 0.356 | 0.239 | 0.357 | 0.307 | 0.433 | 0.265 | 0.349 | 0.366 | 0.232 | 0.361 | 0.269 |
| 7 | 144160001 | 150720000 | 4 | 106 | 0.074 | 0.06 | 0.07 | 0.066 | 0.074 | 0.08 | 0.109 | 0.013 | 0.042 | 0.06 | 0.07 | 0.047 | 0.101 | 0.068 | 0.07 |

| | | | | | | | | | | | | | | | | | | | |
|---|---|---|---|---|---|---|---|---|---|---|---|---|---|---|---|---|---|---|---|
| 7 | 126140001 | 143860000 | 6 | 113 | 0.801 | 0.726 | 0.674 | 0.777 | 0.799 | 0.886 | 0.824 | 0.84 | 0.675 | 0.726 | 0.674 | 0.802 | 0.913 | 0.692 | 0.763 |
| 8 | 46880001 | 146300000 | 3 | 76 | 0.717 | 0.752 | 0.07 | 0.005 | 0.752 | 0.875 | 0.715 | 0.747 | 0.542 | 0.752 | 0.07 | 0 | 0.014 | 0.015 | 0 |
| 8 | 12400001 | 43780000 | 3 | 79 | 0.838 | 0.863 | 0.023 | 0 | 0.893 | 0.943 | 0.814 | 0.867 | 0.717 | 0.863 | 0.023 | 0 | 0 | 0 | 0 |
| 8 | 160001 | 7000000 | 3 | 77 | 0.873 | 0.855 | 0.07 | 0 | 0.866 | 0.864 | 0.905 | 0.893 | 0.817 | 0.855 | 0.07 | 0 | 0 | 0 | 0 |
| 8 | 8080001 | 11880000 | 3 | 78 | 0.975 | 0.991 | 0.07 | 0.002 | 0.98 | 0.989 | 0.986 | 0.987 | 0.933 | 0.991 | 0.07 | 0.006 | 0 | 0 | 0 |
| 8 | 46880001 | 146300000 | 4 | 107 | 0.002 | 0.009 | 0.442 | 0.679 | 0 | 0 | 0.005 | 0 | 0 | 0.009 | 0.442 | 0.744 | 0.783 | 0.526 | 0.688 |
| 8 | 12400001 | 43780000 | 4 | 110 | 0 | 0 | 0.535 | 0.859 | 0 | 0 | 0 | 0 | 0 | 0 | 0.535 | 0.855 | 0.928 | 0.782 | 0.892 |
| 8 | 160001 | 7000000 | 4 | 108 | 0 | 0 | 0.512 | 0.862 | 0 | 0 | 0 | 0 | 0 | 0 | 0.512 | 0.866 | 0.899 | 0.827 | 0.871 |
| 8 | 8080001 | 11880000 | 4 | 109 | 0 | 0 | 0.651 | 0.946 | 0 | 0 | 0 | 0 | 0 | 0 | 0.651 | 0.936 | 0.957 | 0.932 | 0.962 |
| 9 | 200001 | 30340000 | 2 | 35 | 0.879 | 0.94 | 0.86 | 0.888 | 0.899 | 0.92 | 0.896 | 0.88 | 0.792 | 0.94 | 0.86 | 0.901 | 0.957 | 0.827 | 0.892 |
| 9 | 71040001 | 1.41E+08 | 3 | 80 | 0.286 | 0.145 | 0.279 | 0.075 | 0.765 | 0.33 | 0.036 | 0.067 | 0.258 | 0.145 | 0.279 | 0.064 | 0.13 | 0.083 | 0.059 |
| 9 | 30340001 | 38760000 | 3 | 81 | 0.06 | 0.051 | 0.302 | 0.062 | 0.02 | 0.114 | 0.045 | 0.08 | 0.083 | 0.051 | 0.302 | 0.052 | 0.116 | 0.068 | 0.048 |
| 9 | 71040001 | 141020000 | 4 | 111 | 0.467 | 0.564 | 0.442 | 0.702 | 0 | 0.455 | 0.765 | 0.733 | 0.342 | 0.564 | 0.442 | 0.756 | 0.696 | 0.602 | 0.726 |
| 9 | 30340001 | 38760000 | 4 | 112 | 0.815 | 0.838 | 0.558 | 0.838 | 0.852 | 0.739 | 0.869 | 0.787 | 0.742 | 0.838 | 0.558 | 0.814 | 0.797 | 0.835 | 0.876 |
| 10 | 49400001 | 135420000 | 2 | 4 | 0.772 | 0.778 | 0.721 | 0.741 | 0.772 | 0.864 | 0.778 | 0.8 | 0.675 | 0.778 | 0.721 | 0.733 | 0.899 | 0.586 | 0.801 |
| 10 | 120001 | 39140000 | 2 | 5 | 0.003 | 0.923 | 0.837 | 0 | 0.007 | 0 | 0 | 0 | 0.008 | 0.923 | 0.837 | 0 | 0 | 0 | 0 |
| 10 | 120001 | 7720000 | 2 | 6 | 0.058 | 0 | 0 | 0.07 | 0.054 | 0.023 | 0.077 | 0.027 | 0.075 | 0 | 0 | 0.087 | 0.058 | 0.03 | 0.086 |
| 10 | 120001 | 7640000 | 2 | 7 | 0.219 | 0 | 0 | 0.243 | 0.195 | 0.159 | 0.186 | 0.373 | 0.258 | 0 | 0 | 0.192 | 0.145 | 0.346 | 0.253 |
| 10 | 120001 | 7580000 | 2 | 8 | 0.147 | 0 | 0 | 0.161 | 0.134 | 0.125 | 0.172 | 0.107 | 0.158 | 0 | 0 | 0.174 | 0.13 | 0.128 | 0.183 |
| 10 | 33360001 | 39140000 | 2 | 9 | 0.115 | 0 | 0 | 0.114 | 0.081 | 0.08 | 0.172 | 0.067 | 0.108 | 0 | 0 | 0.14 | 0.087 | 0.075 | 0.129 |
| 10 | 33380001 | 39140000 | 2 | 10 | 0.21 | 0 | 0 | 0.207 | 0.295 | 0.182 | 0.149 | 0.227 | 0.225 | 0 | 0 | 0.14 | 0.217 | 0.308 | 0.194 |
| 10 | 33420001 | 39140000 | 2 | 11 | 0.06 | 0 | 0 | 0.057 | 0.047 | 0.057 | 0.054 | 0.053 | 0.092 | 0 | 0 | 0.087 | 0.058 | 0.023 | 0.054 |
| 10 | 42380001 | 46160000 | 2 | 12 | 0.939 | 0.949 | 0.814 | 0.932 | 0.919 | 0.875 | 0.955 | 0.96 | 0.967 | 0.949 | 0.814 | 0.942 | 0.942 | 0.932 | 0.919 |
| 10 | 7360001 | 33500000 | 3 | 43 | 0.172 | 0 | 0 | 0.173 | 0.107 | 0.216 | 0.19 | 0.173 | 0.183 | 0 | 0 | 0.186 | 0.232 | 0.09 | 0.199 |
| 10 | 7620001 | 33400000 | 3 | 44 | 0.32 | 0 | 0 | 0.305 | 0.315 | 0.341 | 0.249 | 0.44 | 0.367 | 0 | 0 | 0.25 | 0.304 | 0.459 | 0.247 |
| 10 | 7680001 | 33300000 | 3 | 45 | 0.32 | 0 | 0 | 0.314 | 0.396 | 0.205 | 0.385 | 0.24 | 0.242 | 0 | 0 | 0.366 | 0.261 | 0.18 | 0.382 |
| 11 | 54800001 | 134940000 | 3 | 46 | 0.438 | 0.444 | 0.442 | 0.541 | 0.309 | 0.705 | 0.525 | 0.56 | 0.167 | 0.444 | 0.442 | 0.529 | 0.638 | 0.338 | 0.661 |
| 11 | 81120001 | 134940000 | 3 | 52 | 0.253 | 0.077 | 0.116 | 0.193 | 0.497 | 0.125 | 0.095 | 0.2 | 0.367 | 0.077 | 0.116 | 0.256 | 0.203 | 0.263 | 0.081 |
| 11 | 200001 | 51580000 | 3 | 47 | 0.476 | 0.744 | 0.651 | 0.8 | 0 | 0.602 | 0.756 | 0.8 | 0.258 | 0.744 | 0.651 | 0.82 | 0.826 | 0.774 | 0.79 |
| 11 | 54800001 | 80780000 | 3 | 48 | 0.084 | 0.051 | 0.047 | 0.075 | 0.181 | 0.034 | 0.041 | 0.027 | 0.117 | 0.051 | 0.047 | 0.128 | 0.014 | 0.098 | 0.032 |
| 11 | 54800001 | 80400000 | 3 | 49 | 0.057 | 0.043 | 0.047 | 0.077 | 0.081 | 0.057 | 0.018 | 0.133 | 0.05 | 0.043 | 0.047 | 0.11 | 0.072 | 0.083 | 0.043 |
| 11 | 40840001 | 51580000 | 3 | 50 | 0.15 | 0.06 | 0 | 0.002 | 0.523 | 0.068 | 0 | 0 | 0.117 | 0.06 | 0 | 0.006 | 0 | 0 | 0 |
| 11 | 200001 | 40840000 | 4 | 82 | 0.24 | 0.043 | 0.023 | 0.002 | 0.785 | 0.193 | 0 | 0 | 0.192 | 0.043 | 0.023 | 0 | 0.014 | 0 | 0 |
| 12 | 37860001 | 133840000 | 3 | 55 | 0.689 | 0.709 | 0.465 | 0.738 | 0.732 | 0.795 | 0.724 | 0.667 | 0.508 | 0.709 | 0.465 | 0.837 | 0.928 | 0.556 | 0.704 |
| 12 | 200001 | 34840000 | 3 | 56 | 0.864 | 0.906 | 0.721 | 0.884 | 0.832 | 0.92 | 0.864 | 0.92 | 0.825 | 0.906 | 0.721 | 0.884 | 0.942 | 0.805 | 0.919 |
| 13 | 19280001 | 115080000 | 2 | 17 | 0.075 | 0.017 | 0.023 | 0.059 | 0.067 | 0.091 | 0.068 | 0.027 | 0.117 | 0.017 | 0.023 | 0.058 | 0.116 | 0.045 | 0.048 |
| 13 | 19280001 | 115100000 | 3 | 57 | 0.51 | 0.436 | 0.395 | 0.371 | 0.732 | 0.625 | 0.552 | 0.093 | 0.333 | 0.436 | 0.395 | 0.738 | 0.42 | 0.293 | 0.07 |
| 13 | 19420001 | 115100000 | 4 | 84 | 0.133 | 0.274 | 0.279 | 0.282 | 0.007 | 0.148 | 0.077 | 0.56 | 0.117 | 0.274 | 0.279 | 0.017 | 0.261 | 0.211 | 0.586 |
| 14 | 20200001 | 107280000 | 3 | 59 | 0.135 | 0.111 | 0.07 | 0.1 | 0.027 | 0.864 | 0 | 0.093 | 0.008 | 0.111 | 0.07 | 0 | 0.797 | 0.008 | 0 |
| 14 | 44260001 | 107280000 | 3 | 61 | 0.639 | 0.65 | 0.698 | 0.621 | 0.711 | 0.011 | 0.792 | 0.813 | 0.617 | 0.65 | 0.698 | 0.797 | 0.029 | 0.549 | 0.731 |
| 14 | 20200001 | 43820000 | 3 | 60 | 0.757 | 0.744 | 0.837 | 0.8 | 0.852 | 0 | 0.928 | 0.827 | 0.833 | 0.744 | 0.837 | 0.919 | 0.029 | 0.865 | 0.93 |
| 15 | 30940001 | 82580000 | 2 | 18 | 0.87 | 0.761 | 0.814 | 0.855 | 0.879 | 0.989 | 0.842 | 0.88 | 0.817 | 0.761 | 0.814 | 0.89 | 0.928 | 0.759 | 0.866 |

| | | | | | | | | | | | | | | | | | | | |
|---|---|---|---|---|---|---|---|---|---|---|---|---|---|---|---|---|---|---|---|
| 15 | 23980001 | 28540000 | 2 | 19 | 0.905 | 0.94 | 0.814 | 0.911 | 0.933 | 0.864 | 0.896 | 0.907 | 0.917 | 0.94 | 0.814 | 0.93 | 0.812 | 0.91 | 0.93 |
| 15 | 84920001 | 102400000 | 4 | 85 | 0.874 | 0.88 | 0.86 | 0.905 | 0.906 | 0.875 | 0.878 | 0.853 | 0.842 | 0.88 | 0.86 | 0.907 | 0.913 | 0.902 | 0.903 |
| 16 | 46440001 | 90160000 | 2 | 20 | 0.851 | 0.795 | 0.674 | 0.77 | 0.872 | 0.943 | 0.86 | 0.92 | 0.7 | 0.795 | 0.674 | 0.762 | 0.855 | 0.639 | 0.839 |
| 16 | 80001 | 16280000 | 3 | 62 | 0.711 | 0.803 | 0.326 | 0.268 | 0.819 | 0.727 | 0.828 | 0.013 | 0.783 | 0.803 | 0.326 | 0.529 | 0.261 | 0.241 | 0.048 |
| 16 | 18800001 | 32660000 | 3 | 63 | 0.159 | 0.137 | 0.116 | 0.043 | 0.101 | 0.011 | 0.154 | 0 | 0.45 | 0.137 | 0.116 | 0.064 | 0 | 0.098 | 0 |
| 16 | 80001 | 16280000 | 4 | 86 | 0.11 | 0.043 | 0.465 | 0.582 | 0 | 0.102 | 0.014 | 0.733 | 0.042 | 0.043 | 0.465 | 0.314 | 0.536 | 0.556 | 0.866 |
| 17 | 25280001 | 81100000 | 3 | 64 | 0.787 | 0.795 | 0.721 | 0.734 | 0.846 | 0.909 | 0.824 | 0.44 | 0.775 | 0.795 | 0.721 | 0.872 | 0.841 | 0.684 | 0.602 |
| 17 | 40001 | 22240000 | 3 | 65 | 0.868 | 0.897 | 0.791 | 0.82 | 0.913 | 0.943 | 0.946 | 0.573 | 0.8 | 0.897 | 0.791 | 0.93 | 0.841 | 0.789 | 0.731 |
| 17 | 25280001 | 81080000 | 4 | 87 | 0.046 | 0.017 | 0.07 | 0.075 | 0 | 0.011 | 0 | 0.333 | 0.033 | 0.017 | 0.07 | 0 | 0.058 | 0.045 | 0.172 |
| 17 | 20001 | 22240000 | 4 | 88 | 0.043 | 0.017 | 0.07 | 0.091 | 0.007 | 0.011 | 0 | 0.32 | 0.017 | 0.017 | 0.07 | 0.006 | 0.072 | 0.068 | 0.194 |
| 18 | 18540001 | 7.80E+07 | 2 | 21 | 0.006 | 0.803 | 0.791 | 0.004 | 0 | 0.023 | 0.005 | 0 | 0.008 | 0.803 | 0.791 | 0.006 | 0 | 0.008 | 0 |
| 18 | 140001 | 15200000 | 2 | 22 | 0.002 | 0.923 | 0.93 | 0.002 | 0.007 | 0 | 0 | 0 | 0 | 0.923 | 0.93 | 0 | 0 | 0.008 | 0 |
| 18 | 140001 | 10060000 | 2 | 23 | 0.086 | 0 | 0 | 0.109 | 0.081 | 0.102 | 0.086 | 0.147 | 0.042 | 0 | 0 | 0.157 | 0.116 | 0.105 | 0.065 |
| 18 | 140001 | 9940000 | 2 | 24 | 0.067 | 0 | 0 | 0.061 | 0.101 | 0.034 | 0.05 | 0.08 | 0.075 | 0 | 0 | 0.058 | 0.043 | 0.068 | 0.065 |
| 18 | 140001 | 9880000 | 2 | 25 | 0.179 | 0 | 0 | 0.205 | 0.154 | 0.114 | 0.163 | 0.253 | 0.242 | 0 | 0 | 0.163 | 0.13 | 0.293 | 0.21 |
| 18 | 140001 | 9840000 | 2 | 26 | 0.107 | 0 | 0 | 0.096 | 0.128 | 0.068 | 0.113 | 0.053 | 0.133 | 0 | 0 | 0.11 | 0.159 | 0.053 | 0.091 |
| 18 | 140001 | 9760000 | 2 | 27 | 0.089 | 0 | 0 | 0.075 | 0.081 | 0.068 | 0.081 | 0.12 | 0.108 | 0 | 0 | 0.099 | 0.087 | 0.075 | 0.048 |
| 18 | 18540001 | 78000000 | 3 | 66 | 0.812 | 0.051 | 0.047 | 0.773 | 0.859 | 0.784 | 0.842 | 0.88 | 0.675 | 0.051 | 0.047 | 0.756 | 0.899 | 0.639 | 0.839 |
| 19 | 27740001 | 59100000 | 3 | 67 | 0.848 | 0.863 | 0.814 | 0.848 | 0.872 | 0.909 | 0.891 | 0.813 | 0.717 | 0.863 | 0.814 | 0.826 | 0.87 | 0.805 | 0.892 |
| 19 | 260001 | 24600000 | 3 | 68 | 0.868 | 0.846 | 0.767 | 0.846 | 0.893 | 0.92 | 0.896 | 0.8 | 0.792 | 0.846 | 0.767 | 0.895 | 0.884 | 0.782 | 0.833 |
| 20 | 29420001 | 62900000 | 4 | 89 | 0.827 | 0.838 | 0.744 | 0.9 | 0.859 | 0.864 | 0.814 | 0.88 | 0.75 | 0.838 | 0.744 | 0.936 | 0.928 | 0.835 | 0.903 |
| 20 | 60001 | 26320000 | 4 | 90 | 0.862 | 0.88 | 0.814 | 0.88 | 0.852 | 0.886 | 0.896 | 0.84 | 0.808 | 0.88 | 0.814 | 0.907 | 0.942 | 0.797 | 0.892 |
| 21 | 15220001 | 48060000 | 3 | 70 | 0.865 | 0.769 | 0.791 | 0.852 | 0.852 | 0.875 | 0.878 | 0.947 | 0.8 | 0.769 | 0.791 | 0.843 | 0.913 | 0.827 | 0.855 |
| 22 | 16860001 | 51180000 | 4 | 91 | 0.842 | 0.829 | 0.698 | 0.82 | 0.846 | 0.909 | 0.896 | 0.853 | 0.683 | 0.829 | 0.698 | 0.849 | 0.826 | 0.699 | 0.876 |
| X | 92380001 | 155240000 | 2 | 36 | 0.655 | 0.726 | 0.488 | 0.632 | 0.812 | 0.841 | 0.756 | 0.68 | 0.125 | 0.726 | 0.488 | 0.634 | 0.797 | 0.398 | 0.737 |
| X | 300001 | 58560000 | 2 | 42 | 0.069 | 0.034 | 0.047 | 0.066 | 0.047 | 0.318 | 0.014 | 0.08 | 0.008 | 0.034 | 0.047 | 0.023 | 0.304 | 0.03 | 0.043 |
| X | 114420001 | 155240000 | 2 | 37 | 0.167 | 0.137 | 0.14 | 0.175 | 0.06 | 0.045 | 0.095 | 0.12 | 0.55 | 0.137 | 0.14 | 0.174 | 0.058 | 0.323 | 0.113 |
| X | 300001 | 31300000 | 2 | 38 | 0.783 | 0.821 | 0.651 | 0.789 | 0.812 | 0.5 | 0.878 | 0.827 | 0.75 | 0.821 | 0.651 | 0.895 | 0.522 | 0.722 | 0.839 |
| X | 61720001 | 88460000 | 2 | 39 | 0.934 | 0.957 | 0.767 | 0.927 | 0.953 | 0.966 | 0.941 | 0.893 | 0.9 | 0.957 | 0.767 | 0.948 | 0.942 | 0.887 | 0.93 |
| X | 32300001 | 58560000 | 2 | 40 | 0.848 | 0.923 | 0.698 | 0.834 | 0.913 | 0.602 | 0.932 | 0.827 | 0.808 | 0.923 | 0.698 | 0.919 | 0.638 | 0.805 | 0.849 |
| X | 92380001 | 114200000 | 2 | 41 | 0.103 | 0.068 | 0.116 | 0.123 | 0.04 | 0.045 | 0.032 | 0.08 | 0.367 | 0.068 | 0.116 | 0.128 | 0.043 | 0.248 | 0.059 |
| Y | 13460001 | 19560000 | 0 | 1 | 0.982 | 0.974 | 1 | 0.98 | 0.973 | 1 | 0.982 | 1 | 0.967 | 0.974 | 1 | 1 | 0.986 | 0.94 | 0.989 |
| Y | 20800001 | 24520000 | 0 | 2 | 1 | 1 | 1 | 1 | 1 | 1 | 1 | 1 | 1 | 1 | 1 | 1 | 1 | 1 | 1 |
| Y | 6620001 | 10080000 | 0 | 3 | 1 | 1 | 1 | 1 | 1 | 1 | 1 | 1 | 1 | 1 | 1 | 1 | 1 | 1 | 1 |

**Supplementary file 1 |** R script to perform Clustering of single cell CNV data, generating a CNV mutation matrix with polymorphic events.

```
library(GenomicRanges)
library(dplyr)

# path to outs folder
PATH_2_OUTS <- "YOUR_PATH"

# read the per cell summary
per_cell_summary <- read.table(file.path(PATH_2_OUTS,"per_cell_summary_metrics.csv"), sep=",",header =
TRUE)

# define inputs and cutoffs
cells <- per_cell_summary$cell_id[per_cell_summary$is_noisy==0]
cells <- paste0("cell_",cells)
NCELLS <- length(cells)
EVENT_QUAL = 15
EVENT_SIZE = 2e6
EVENT_FREQ = .05
CUTOFF_N <- round(NCELLS*EVENT_FREQ)

#' read bed file and subset to keep events only from
#'terminal nodes that are not flagged as noisy
get_bed <- function(path_2_outs){
  df <- read.table(file.path(path_2_outs,"node_cnv_calls.bed"),header=FALSE,sep="\t")
  names(df) <- c("chrom", "start", "end", "node", "ploidy", "qual")
  nnodes <- max(df$node)
  ncells <- nnodes/2
  df$node <- ifelse(df$node>ncells,paste0("inode_",df$node),paste0("cell_",df$node))
  return(df)
}

bed <- get_bed(PATH_2_OUTS)
bed <- subset(bed,node %in% cells)
bed.gr <- makeGRangesFromDataFrame(bed, keep.extra.columns=TRUE,
                                  ignore.strand=TRUE,
                                  seqinfo=NULL,
                                  seqnames.field="chrom",
                                  start.field="start",
                                  end.field="end",
                                  starts.in.df.are.0based=TRUE)

# apply a quality and size cut off and
# order the events going from largest to smallest
bed.gr <- bed.gr[bed.gr$qual>EVENT_QUAL]
bed.gr <- bed.gr[width(bed.gr) > EVENT_SIZE]
bed.gr <- bed.gr[order(-width(bed.gr),start(bed.gr))]

#split bed file by ploidy and chromosome
bed.gr <- split(bed.gr,bed.gr$ploidy)
```

```
split.bed.gr <- lapply(bed.gr, function(x) split(x,seqnames(x)))

# find pairwise 90% reciprocal overalps with the same ploidy
events.byploidy <- list()
for (i in seq_along(split.bed.gr)){
  gr <- split.bed.gr[[i]]
  hits <- lapply(gr, function(x) findOverlaps(x,drop.self=FALSE,drop.redundant=TRUE,type="any",select="all"))
  xhits <- lapply(seq_along(gr), function(x) gr[[x]][queryHits(hits[[x]])])
  yhits <- lapply(seq_along(gr), function(x) gr[[x]][subjectHits(hits[[x]])])
  pmax <- lapply(seq_along(gr), function(x) pmax(width(yhits[[x]]),width(xhits[[x]])))
  overlaps <- lapply(seq_along(gr), function(x) pintersect(xhits[[x]],yhits[[x]]))
  frac <- lapply(seq_along(gr), function(x) width(overlaps[[x]])/pmax[[x]])
  merge <- lapply(frac, function(x) x>=0.9)

  final <- list()
  for(j in seq_along(gr)){
    tokeep <- yhits[[j]][merge[[j]]]
    tokeep$qhit <- queryHits(hits[[j]])[merge[[j]]]
    tokeep$subhit <- subjectHits(hits[[j]])[merge[[j]]]
    #the same subject hit is being grouped multiple times and supporting different queries
    tokeep <- tokeep[!duplicated(tokeep$subhit)]
    qcount <- table(tokeep$qhit)
    tokeep <- tokeep[tokeep$qhit %in% names(qcount[qcount>CUTOFF_N])]
    final[[j]] <- tokeep
  }
  final <- do.call("c",final)
  events.byploidy[[i]] <- final
}

# generate unique labels for shared events
label.qhit <- sapply(events.byploidy, function(x) paste0(x$qhit))
label.chr <- sapply(events.byploidy, function(x) as.vector(seqnames(x)))
label <- sapply(seq_along(label.qhit), function(i) paste0(label.chr[[i]],label.qhit[[i]]))
label <- sapply(seq_along(label),function(i) paste0("ploidy",i-1,"_chr",label[[i]]))
label <- unlist(label)
tokeep <- sapply(label,function(x) grepl("ploidy\\d+_chr[XY]*\\d+",x))
label <- label[tokeep]
label <- factor(label,labels = 1:length(unique(label)))

# make a single GenomicRanges object
events.05.gr <- do.call("c",events.byploidy)
events.05.gr$event <- label
events.05.gr$qhit <- NULL
events.05.gr$subhit <- NULL
rm(events.byploidy)
```

**Supplementary file 2** | Python script to split BAM files by barcode assignment, generating a BAM file for each sub-clone.

```
import pysam
import pandas as pd
import sys
import os
import argparse

def parse_args():
  parser = argparse.ArgumentParser()
parser.add_argument('--bam', help='Input BAM file')
parser.add_argument('--out-root', help='Root directory to write output BAMs to')
parser.add_argument('--csv', help='CSV file with two columns: barcode and cluster.')
return parser.parse_args()

if __name__ == '__main__':
  args = parse_args()
df = pd.read_csv(args.csv)
assert 'barcode' in df.columns and 'cluster' in df.columns, 'Missing column identifiers'
assert os.path.exists(args.bam), 'Cannot find input BAM'
fh = pysam.Samfile(args.bam)
cluster_map = dict(zip(df.barcode, df.cluster))
os.makedirs(args.out_root)
out_handles = {}
for c in set(df.cluster):
  o = pysam.Samfile(os.path.join(args.out_root, 'cluster' + str(c) + '.bam'), 'wb', template=fh)
out_handles[c] = o
for rec in fh:
  if rec.has_tag('CB') and rec.get_tag('CB') in cluster_map:
  cluster = cluster_map[rec.get_tag('CB')]
out_handles[cluster].write(rec)
for h in out_handles.values():
  h.close()
```
